# A complete single-cell atlas of the embryonic *Drosophila* heart and the cell-type specific role of Tinman

**DOI:** 10.1101/2021.01.15.426556

**Authors:** Georg Vogler, K’Leigh Guillotte, Bill Hum, Marco Tamayo, Yoav Altman, Karen Ocorr, Rolf Bodmer

## Abstract

Heart morphogenesis is a complex process that is orchestrated during development via the interaction of different cell types and the activity of distinct gene programs within these cells. Here, we analyzed the development and differentiation of the *Drosophila* embryonic heart at the single-cell level to characterize in detail the genetic expression profiles and phenotypic differences of cardiac cell types during heart morphogenesis. We present an embryonic fly heart cell atlas at unprecedented resolution that integrates the entire catalogue of known heart cells. We identified new gene programs and cell type marker genes that allows characterization of the molecular genetics of fly cardiogenesis in granular detail. In cardioblasts we described the temporal process of cardioblast-to-cardiomyocyte differentiation. Two sets of pericardial cells, likely contributing to cardiomyocyte differentiation, are eliminated by programmed cell death at the end of embryogenesis, whereas as third set continues to shape/influence cardiac function into adulthood. To dissect the gene programs downstream of the cardiac homeodomain transcription factor *tinman* we analyzed cardiac cells with reduced levels of Tinman. Here we find that Tinman acts both as suppressor and as activator of identified direct Tin-target genes in a cell type-dependent manner. We also found a developmental switch that alters the fate of pericardial cells towards wing heart cell fate and identified an entire wing heart gene program suppressed by Tinman in pericardial cells. Lastly, we find that in pericardial cells Tinman controls a Wingless/WNT receptor switch through selective activation and repression of *frizzled* and *frizzled2*, respectively. This study paves the way for investigating other core cardiogenic genes in delineating cardiac regulatory networks.

## INTRODUCTION

Congenital heart disease (CHD) is the most frequent birth defect among newborns and the leading cause of infant death or illness. While massive genome sequencing efforts have exponentially expanded the number of potential genes involved in human disease, including CHD ^1–3^, we still lack a fundamental mechanistic understanding how most of these genes and variants contribute to disease ^4^. In addition, most candidate genes are likely to act in concert with other genes in a patient-specific genetic background; and the study of such gene-gene interactions has not been established yet at a scale required for most diseases and conditions ^5,6^. Model systems that allow to systematically identify genetic interactions with high throughput are typically based on cell assays of organisms with a simple genetic architecture such as yeast or *Drosophila* S2 cell cultures ^7–9^. For genetic interactions within a multi-cellular organism, the fruit fly *Drosophila* has always been an important model organism (for example *NOTCH,* see ^10^). The high degree of genetic conservation between flies and humans, including the presence of orthologous organ systems (including the heart) and conservation of many developmental and molecular-genetic mechanisms make *Drosophila* highly relevant for the study of human disease. This sets the stage for new approaches that integrate patient-specific genomics with an amenable *Drosophila* model to understand fundamental genetic pathways likely at play in the disease process (e.g. ^11,12^). The prospect to study genetic interactions at the single-cell level in a multicellular organism is a logical next step in this important endeavor.

The formation of the heart is a finely controlled process of specification and differentiation events thought to be under tight control of cardiac transcription factors and signaling pathways, in vertebrates as well as in the fruit fly ^13–15^. The segmental *Drosophila* embryonic heart consists of 104 cardioblasts (CBs), distributed over eight segments (T3, A1-A7), as well as several types of pericardial cells ^16–18^. Post-specification the CBs express a common set of transcription factors (TFs), including Tail-up, H15 and Mef2 ^19–21^. However, within each segment there are defined subsets of CBs that in addition express a combination of other cardiac TFs (Tinman and/or Ladybird; Doc1/2/3/Svp) ^22–26^. CBs are flanked by ∼140 pericardial cells (PCs) that are also expressing a common set of TFs (Hand, Zfh1), as well as other TF combinations that create different types of PCs, including Tin, Odd and Eve ^27–31^.

Of these transcription factors, *tinman* is expressed in the mesoderm and then restricted to the cardiac mesoderm prior to CB subtype diversification ^22,23^. The cardiac specificity and DNA sequence-specific binding of the homeodomain-containing Tinman protein has allowed for detailed ChIP-on chip characterization of cardiac Tinman target genes ^32–34^, in parallel to several groups defining Tinman-dependent regulatory networks by bioinformatical FACS-assisted characterization of chromatin states of cardiac cells and machine learning approaches ^35,36^. Nevertheless, the regulatory gene networks involved are still incompletely understood, in part because these could not be done in a cell specific manner.

To obtain a comprehensive picture of the logic and landscape of the gene networks within cardiac subtypes, it is now possible to perform genome wide RNA sequencing at the single cell level (scRNAseq). We therefore sought to create a transcriptional atlas of the embryonic fly heart to identify the full set of CB- and PC-expressed genes potentially involved in cardiac subtype differentiation and heart-tube morphogenesis and maturation. This will allow us to examine co-expression data of transcription factors and target genes at the single cell level to identify comprehensive gene regulatory networks.

Here, we present an atlas of the embryonic *Drosophila* heart during the last stages of embryonic development, based on FACS-sorted scRNAseq of heart cells. We present a high-resolution map of cardiac cell types as well as other systematically recovered cell types of the late developing fly embryo, including cell type- and cell subtype-specific marker genes. Our analysis uncovers distinct biological processes active during tissue differentiation and maturation, such as combinatorial expression of a suite of cell adhesion molecules, signaling pathways and transcription factors. Our data also show that two pericardial cell types undergo programmed cell death at the end of heart morphogenesis settling their hitherto unknown fate.

Mutations in cardiac transcription factors and their interactors are often the root cause of congenital heart diseases ^37–43^. It is thus necessary to understand their downstream regulatory networks controlling cardiac gene expression, which is a highly complex endeavor. Analyzing heart development by an orthogonal approach, such as using *Drosophila* and to study conserved pathways, can reveal general blueprints for such networks ^40,44^, e.g. downstream of Tinman. Due to the organ-specific expression of Tinman in the dorsal vessel it was possible to determine its binding to regulatory DNA via chromatin immunoprecipitation ^32–34^, and in combination with motif and chromatin analysis and machine learning many potential direct Tinman target genes have been revealed ^35,36^. However, without experimental validation of cell type-specific expression, e.g. by *in situ* ^45^ cardiac specificity of Tinman target genes was not comprehensively resolved to the cardiac sub-cell type level (e.g., CB vs PC). Furthermore, the dependency of enhancer activity on Tinman was tested only for a few genes ^32^, thus if Tinman is an activator or repressor for most of these targets is not clear. In this study we provide a cell-type specific map of Tinman activity in the heart.

Because the cardiac transcription factor Tinman is a key factor of heart development, necessary for both specification and differentiation of cardiac cell types, we employed mesoderm-specific RNAi-mediated knockdown of *tinman* to study changes in gene expression at the single-cell level. This allowed us to determine at cellular resolution whether Tinman is an activator or repressor for its reported target genes. Most notably, as a specific example, we report that Tinman directs two biological processes: it suppresses the wing-heart program in cardiac cell types and controls a Frizzled/Frizzled-2 receptor switch in Even-skipped pericardial cells.

## RESULTS

### Embryonic transcriptome atlas of *Hand*-reporter expressing cells

To comprehensively describe the cellular diversity of the *Drosophila* embryonic heart as it undergoes morphogenesis and differentiation we performed scRNA-seq on cardiac cells of mid-to late-stage embryos. The cardiac transcription factor Hand has been found to be strongly expressed in all cardiac cell types as well as a few other tissues ^46,47^. To be able to fluorescently sort heart cells prior to sequencing we made use of a 513bp enhancer inside the *Hand* gene that fully recapitulates *Hand* expression ^28^ and created a *Hand*::DsRed^express^ (RFP) fluorescent reporter line (Hand^RFP^). In line with previous reports, major tissues labeled by RFP include the heart, lymph gland and pericardial cells, cells anterior to the heart (wing hearts, see below), cells of the ventral nerve chord and proventriculus (**Figure 1A**). Two-hour collections of embryos from two wildtype genetic backgrounds (4 GD and 5 KK replicates ^48^, **Figure 1B**) were sorted (**Supplemental Figure 1A**), processed with 10X Chromium v3.2 followed by stringent bioinformatic analysis (see <u>Methods</u>). We recovered a total of 92k cells (GD: 32k, KK: 60k), which clustered to all expected cell types based on their marker gene expressions (Unsupervised uniform Manifold Approximation and Projection (UMAP) clustering, **Figure 1C**, **Supplemental Table 1**). In addition, we identified clusters with gene expression characteristic for myoblasts/muscle cells ^49^ and plasmatocytes ^50^, both of which have not been reported to express Hand^RFP^. Cells of the brain and ventral nerve chord include different subtypes of neuroblasts and ganglion mother cells ^51^, glia cells ^52^ and neurons ^53^ (**Supplemental Figure 2**). Our single-cell dataset thus contains all expected cell types based on *Hand* reporter gene expression.

**Figure 1.**
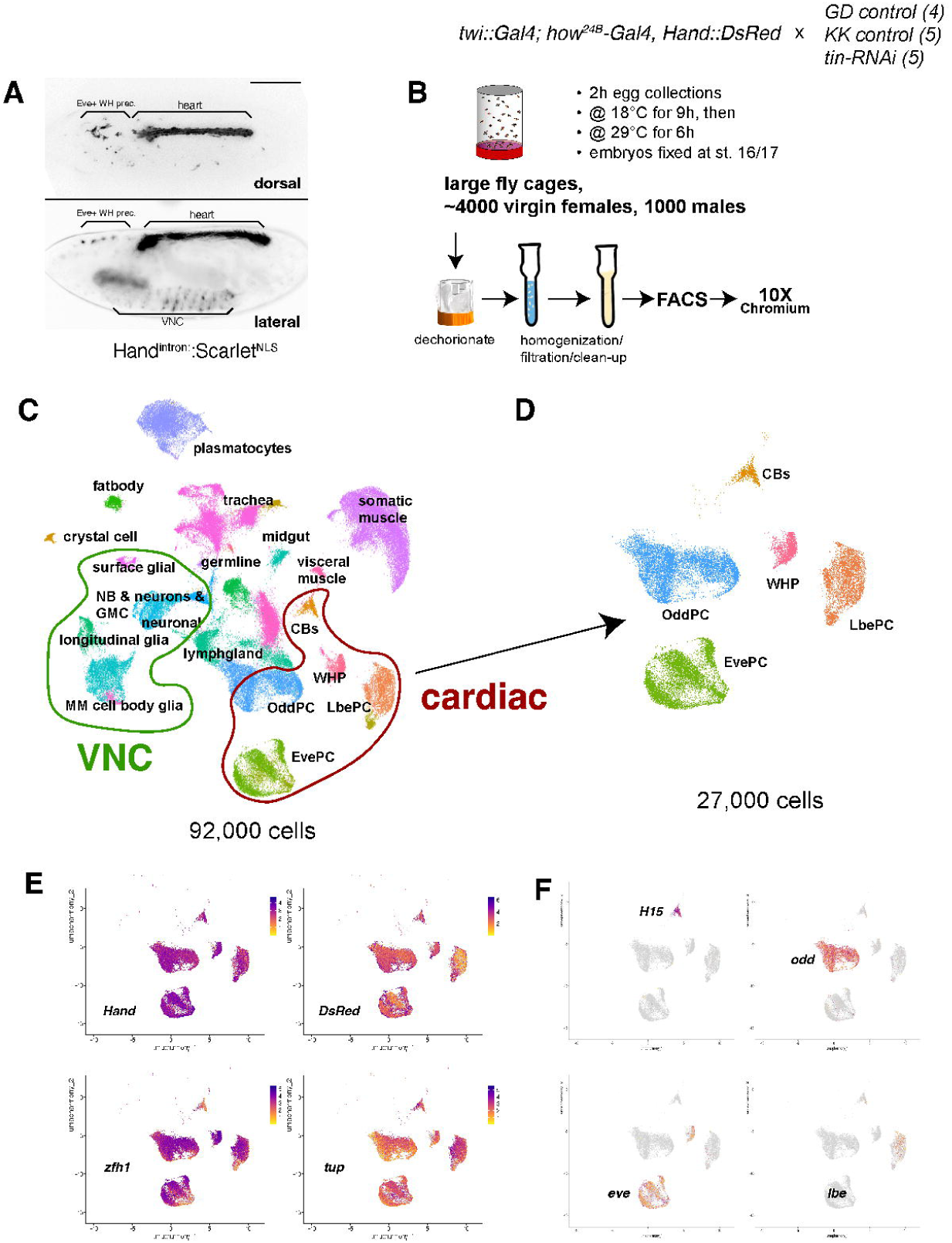

### All cardiac cell types at single-cell resolution

The strongest expression of Hand^RFP^ is seen in the cells of the dorsal heart (**Figure 1A**), which is comprised of several cell types: cardioblasts (CBs) that develop into cardiomyocytes, Odd-skipped-positive pericardial cells (*odd*-PCs) that become nephrocytes ^54^, Tinman-positive pericardial cells (TPCs) that co-express *even*-*skipped* (*Eve-*PCs, dorsally located) ^55,56^ or ventral TPCs expressing *ladybird-early* (*Lbe-PCs* ^30^). The anterior-most population of *HandRFP*-positive cells are wing-heart precursors ^57^. We were able to identify each of these cell types in the scRNA-seq dataset relying on a few known marker genes in combination with *Hand* expression (**Figures 1F, 2**). Cardiac cells contribute to about one third of all sorted cells (27k of 92k cells) across all samples. In addition, we detect expression of other known pan-cardiac genes *tail-up* (*tup*) ^21^ and *Zn finger homeodomain 1* (*zfh1*) ^31^ in all cardiac cells as expected (**Figure 1E**), although the latter more prominently in the PCs.

**Figure 2.**
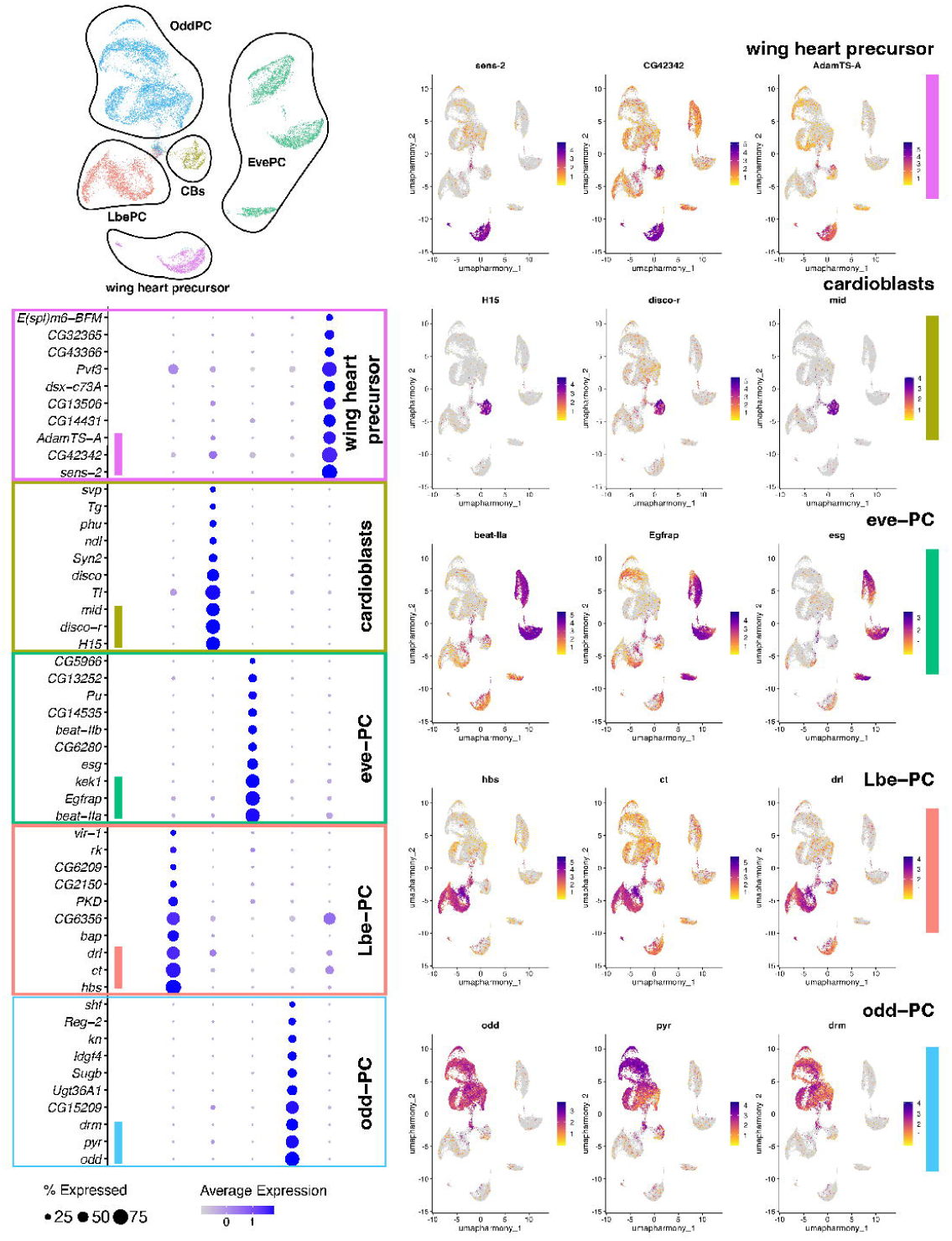

We next focused on identifying novel or previously unrecognized cell type-specific transcription factors, signaling pathways and cell adhesion molecules that might contribute to the assembly and terminal differentiation of the embryonic heart in addition to many of the known cardiac genes. For each cardiac cluster we determined genes of annotated pathways (e.g. wingless/WNT) and transcription factors that are highly expressed in one cell type versus all cardiac cells (**Supplemental Table 2**), which will be described in the following sections. Lastly, we performed GO term enrichment analysis using cell type-specific marker gene expression (with an average expression log2FC > 0.5 and FDR < 0.05 and expressed in more than 30 percent of cells of the respective tissue; **Supplemental Table 3**). Interestingly, the GO term “cell surface receptor signaling pathway” was highly enriched in all three cardiac cell types and “motor neuron axon guidance” stood out in Eve- and Lbe/Tin-PCs.

### Cardioblast differentiation at the single-cell level

The 104 cardioblasts (CBs) of the segmental Drosophila embryonic heart are distributed over eight segments (T3, A1-A7) ^16–18^. Among all cardiac cell types, CBs are the most studied. Post-specification, the CBs express a common set of transcription factors (TFs), including *tail-up*, *H15* and *Mef2* ^19–21^ and within each segment CBs are subdivided through a combination of other cardiac TFs (*tinman* and/or *ladybird-early*; *dorsocross-1/-2/-3*; *seven-up*)^22–26^. Along the anterior-posterior axis the heart has two compartments, the anterior aorta (characterized e.g. by *mthl5* ^58^) and the posterior heart proper (expresses e.g. *Mp* ^59^). While strongly labeled by Hand^RFP^, CBs proved difficult to isolate and their number in the scRNA-seq dataset is not stochiometric to what we expect compared to other cardiac cells such a EvePCs and LbePCs. We obtained a total of 799 CBs which clustered into four compartments (**Figure 3**): into non-overlapping subsets of *tinman*-positive and *seven-up*-positive CBs, and within each subset into non-overlapping clusters of *mthl5-*positive of the aorta (anterior CBs) and *Mp*-positive cells of the heart proper (posterior CBs). When we focused on the *svp*-positive cluster (cells that become the inflow tracts of the heart (‘ostia’)), we identified several new genes with unreported expression (**Supplemental Figure 3**): *Wnt6*, another member of the Wingless(*wg*)/WNT signaling pathway known to be active in the fly heart proper (such as *wg*, *Wnt4* ^60–63)^ is expressed in the same cells as *wg*. In addition, we find expression of neurotransmitter/receptor genes such as Ion transporter peptide (ITP), *serotonine receptor 7* (*5-HT7*), *Shaker cognate b* (*Shab*), *SK* channel and glutamate receptor *GluRIIA* strongly co-expressed in *wg*-positive ostia cells (**Supplemental Figure 3**).

**Figure 3.**
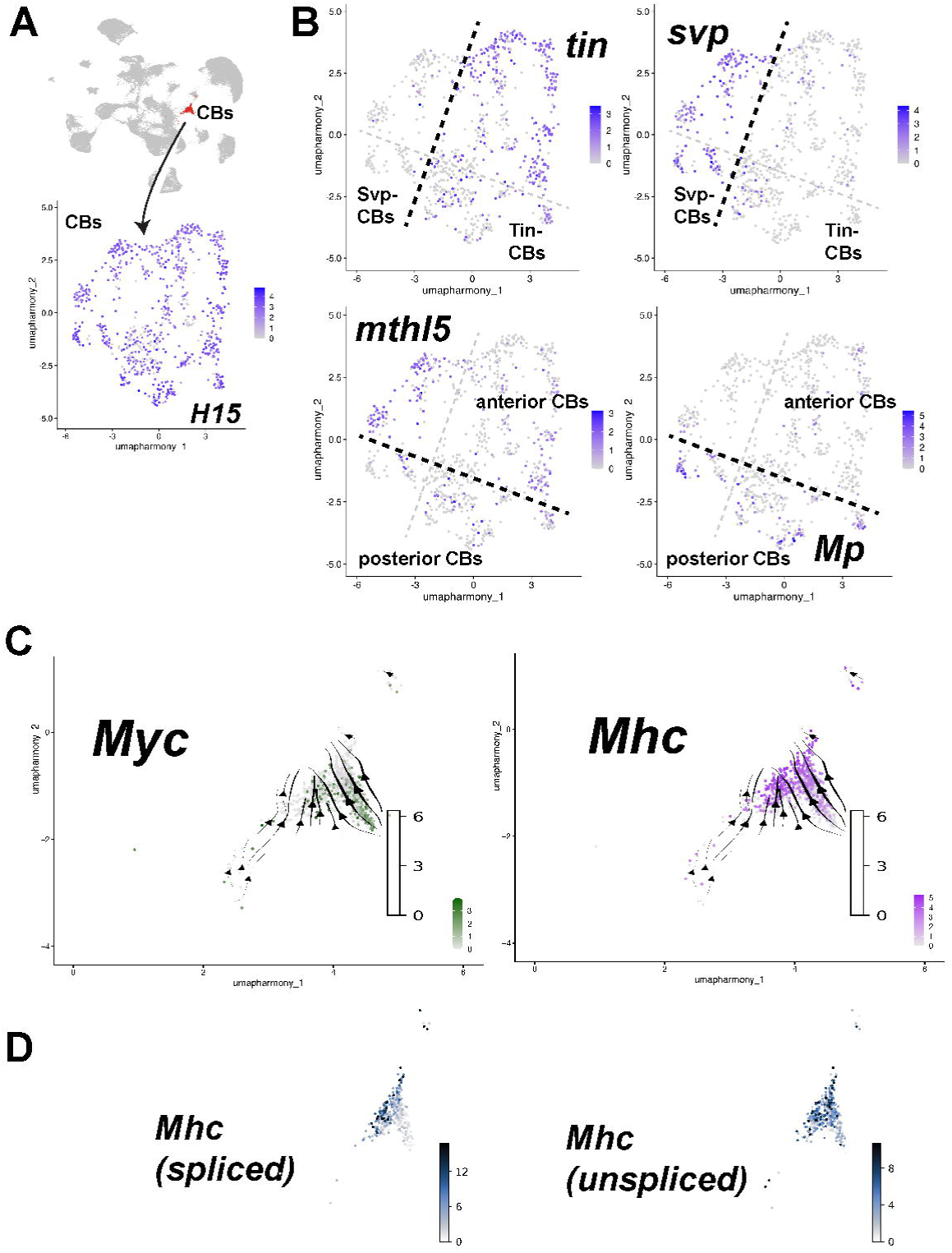

Since we collected embryos over a 2-hour window we assumed that we should be able to infer temporal aspects within cell types using RNA velocity as proxy ^64,65^. When we ran this analysis on CBs, we found trajectories from early (less differentiated) CBs to differentiated cardiomyocytes (CMs) with high expression of sarcomeric genes such as *Mhc* and *bt* (**Figure 3, Supplemental Figure 3**). For example, the onset of expression of the main muscle structural gene Mhc (by relative increased amounts of unspliced *Mhc*) occurs before we observe the full expression level of the spliced *Mhc* transcript. This expression of sarcomeric genes is preceded by ribosomal biogenesis genes likely induced by *Myc*, displaying an inverse pattern of expression along the velocity trajectory compared to spliced *Mhc* (**Figure 3C, D**). scRNA-seq of developing CBs therefore not only captures novel molecular-genetic factors but also temporal dynamics during cardioblast-to-cardiomyocyte differentiation.

### A complete pericardial cell atlas reveals new mechanisms during heart development

During early heart development cardiac lineages undergo cell divisions giving rise to CBs as well as pericardial cells and a subset of muscle precursors ^30,66^. During heart morphogenesis, ∼140 pericardial cells (PCs) flanking the CBs are also expressing a common set of TFs (*Hand*, *zfh1*), as well as other TF combinations that create different types of PCs, including Tin, Odd and Eve ^27–31^. While some cell types persist into adulthood (cardiomyocytes, nephrocytes, wing hearts), the fate of Eve- and Lbe-PCs has been unclear. In contrast to CBs, we were able to harvest plenty of PCs for scRNA-seq and present a complete pericardial cell atlas of the *Drosophila* embryo.

#### Odd-skipped Pericardial Cells

From the cardiogenic mesoderm two lineages emerge: an Odd-positive progenitor and a mixed CB/PC lineage both of which produce OddPCs, the precursors cells that differentiate into the larval and adult nephrocytes ^54,66,67^. In addition, the anterior dorsal mesoderm produces the Odd-positive lymph gland (LG) ^68,69^. We identified 13,600 *odd*-positive cells belonging to these two major groups in our scRNA-seq datasets: nephrocytes that co-express the collagen-processing enzyme *lonely heart* (*loh*) ^70^, and the anteriorly located lymph gland cells characterized by co-expression of the transcription factors *serpent (srp)* ^71^ and *Collier*/*knot* (*kn*) ^72^(**Figure 4**). New marker genes that discriminate between OddPCs and LG are *hamlet* and *PCNA* (LG markers ^45^), as well as *CadN* (**Supplemental Figure 4**). Odd-positive cells and hemocytes are also both positive for the GATA transcription factor *pannier*, *pnr*.

**Figure 4.**
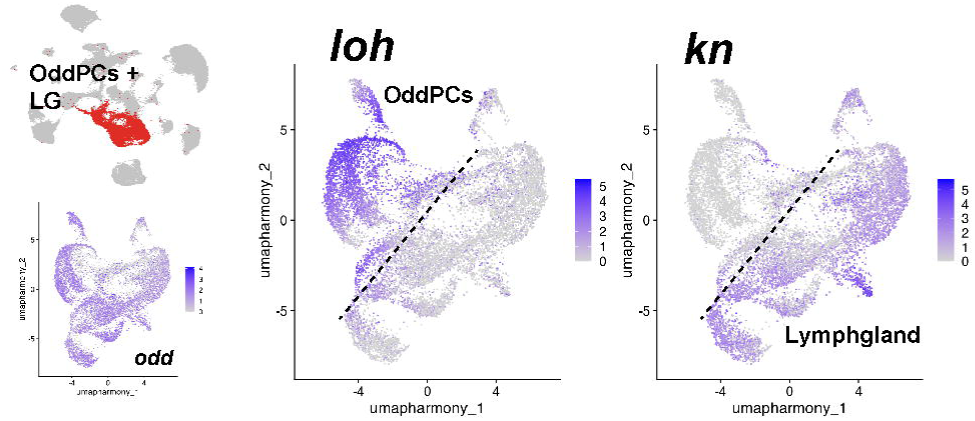

Aside from these three factors the main differences between subgroups of these cells are homeotic gene expression patterns (**Supplemental Figure 4**), indicating minor differences among cell types along the AP axis. With respect to signaling pathways, OddPCs show expression of *Pvf2* (PDGF- and VEGF-receptor related signaling), and both FGF ligands *pyramus* (*pyr*) and *thisbe* (*ths*).

#### Even-skipped Pericardial Cells

Even-skipped Pericardial Cells (EvePCs) are characterized by expression of the transcription factor Even-skipped. They were the first pericardial cell type to be described, located alongside cardioblasts throughout embryonic heart development ^22,23,55^. EvePCs originate from a bipotent precursor in the cardiac mesoderm that is giving rise to an EvePC and a founder cell of the dorsal acute 1 (DA1) muscle ^73,74^. The regulation of cardiac *eve* expression has been analyzed in detail ^75,76^ and embryos lacking mesodermal *eve* expression show normal cardiac development, with reduced number of larval PCs and reduced larval heart rate ^56^. Eve-positive pericardial cells differentiate into three groups towards the end of embryogenesis: EvePCs that remain closely associated with the heart tube, EvePCs that descent towards the outflow tract of the heart ^77^ and the anterior located wing heart precursor cells (WHPs, see next section)^57^.

Hox gene expression of these Eve-positive subgroups ^77^ showed that they split into three distinct groups characterized by expression of (a) anterior Hox genes *Antp* (EvePC + WHP) and *Ubx* (EvePCs), (b) posterior Hox genes *abd-A* and *Abd-B* (EvePCs) and (c) a group of cells devoid of Hox gene expression (**Figure 5A,B**). This Hox-negative subgroup is characterized by expression of the Iroquois genes *araucan* (*ara*) and *caupolican* (*caup*) (**Figure 5C, D**). Four Ara/Caup/Eve-positive cells are located medially to the anterior-most EvePCs, with the two anterior cells expressing higher levels of Ara/Caup compared to the trailing pair (**Figure 5C, D**). This population of *ara/caup/eve*-positive cells has previously been described to be localized in the thoracic segments ^78^. Together with our observation of absence of Hox gene expression clearly identifies them as part of the cardiac outflow tract, the outflow hanging structure as described by ^77^. This is a *bona fide* example of how data derived from single cell data can perfectly align with well-described spatial data from morphological analysis.

**Figure 5.**
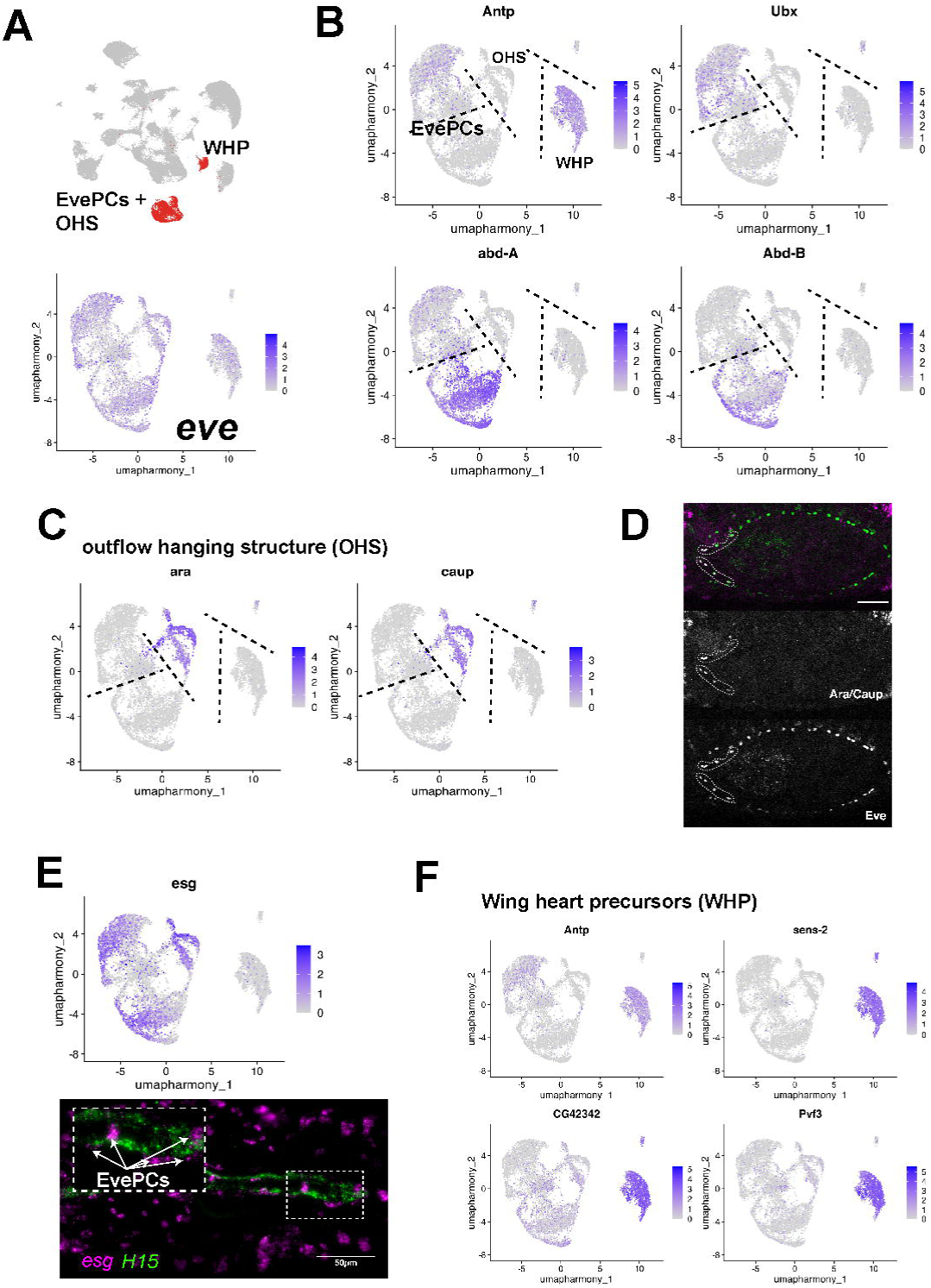

We next determined gene groups that are specifically expressed in EvePCs versus other cell types (**Supplemental Table 1**), namely components of the EGF signaling pathway (*EGFR adaptor protein*, *Egfrap*; *kekkon 1*, *kek1*; *vein*, *vn*; *rhomboid*, *rho*) and the wingless receptor *frizzled 2* (*fz2*). EvePCs and WHPs shared several cell adhesion genes (*rolling pebbles*, *rols*; *Teiresias*, *tei; Lachesin*, *Lac*), as did EvePCs and LbePCs (see below) (*scab*, *scb*; *shotgun*, *shg; Lachesin*, *Lac;* **Supplemental Figure 5**). Also shared expression between Eve- and LbePCs is the G-protein coupled receptor (GPCR) *Corazonine receptor* (*CrzR*) (**Supplemental Figure 5**). Interestingly, we found the Snail-type zinc finger transcription factor *escargot* (*esg*) to be specifically expressed in EvePCs (**Figure 5E**), including cells of the outflow hanging structure. *esg* is strongly expressed in the heart-anchoring cells ^79^, and overexpression of *esg* was found to increase the number of *eve*-positive cells of the heart ^80^.

Other genes with differential gene expression within the Eve cluster are the zinc-finger transcription factor *teashirt* (*tsh*), the actin-binding protein *formin3* (*form3*), and the Netrin-receptor *unc-5* (all anterior genes) and *CG13252* (posterior expression) (**Supplemental Figure 5C**), and the immunoglobulin proteins of the *beaten path* family (specifically *beat-IIb* and *beat-IIa*) that are involved in motoneuronal axon guidance ^81^. It is interesting to note that many are involved in cell adhesion and/or motility, suggesting a role in cardiac morphogenesis.

#### Wing heart precursor cells

The intronic *Hand* enhancer was shown to be active in an anterior population of cells (**Figure 1A** and ^82^) that are part of the EvePC lineage and that become wing hearts, a structure of the thorax necessary for clearing and maturation of the wing ^57,83^. Indeed, *eve*-positive cells in our dataset cluster into two main groups, a *tinman*-positive EvePCs cluster and a *tinman*-negative, *Antp*-positive wing heart precursor (WHP) cluster (**Figure 5B, Supplemental Figure 6A**). Our scRNA-seq data contained a cluster of cells characterized by the co-expression of the anterior hox gene *Antennapedia* (*Antp*) and *eve*, indicating that these are indeed WHPs (**Figure 2**, **Supplemental Figure 6A**). Upon further inspection of marker genes for WHPs we found several new WHP genes: most notably they express the conserved zinc-finger transcription factor *senseless-2* (*sens-2*; *GFI1/1B ortholog*), which is also expressed in the surface glia cluster (SPG, **Supplemental Figure 6A**, and ^84^), as well as the collagen type IX/XIII gene *CG42342* and the PDGF signaling ligand *Pvf3* (**Figure 5F**).

#### Ladybird-Pericardial Cells

During heart development a third population of pericardial cells that are Ladybird/Tinman-positive (LbePCs) is found alongside the heart, but its developmental function has yet to be determined ^85^. From the Tin-positive cardiac mesoderm two groups of cells arise: Tin/Lb-positive CBs and Tin/Lb-positive PCs ^30,66,85^. At the end of heart morphogenesis, LbePCs localize ventrally to the forming heart tube (**Fig. 4A-C**), with high expression levels of the cell adhesion receptor Neurotactin (Nrt), as well as the cardiac transcription factors Zfh1 ^31^ and Tin (**Figure 6**). In our single-cell data we capture over 4000 LbePCs that allows us to identify new genes to describe this cell in greater detail (**Supplemental Table 1**). LbePCs express the two homeobox genes *tinman* and *ladybird early* (along with *zfh1*), but in addition we find that two other homeodomain transcription factors, *cut* (*ct*) and *bagpipe* (*bap*), are also strongly expressed in LbePCs throughout heart morphogenesis and heart closure (**Figure 6**). With respect to signaling pathways, LbePCs express *Pvf2* and *Pvf3* as well as their receptor *Pvr*, the Notch agonist *hibris* (*hbs*), the WNT ligand *Wnt4* and the WNT receptor *frizzled* (*fz*) (**Supplemental Figure 6B**).

**Figure 6.**
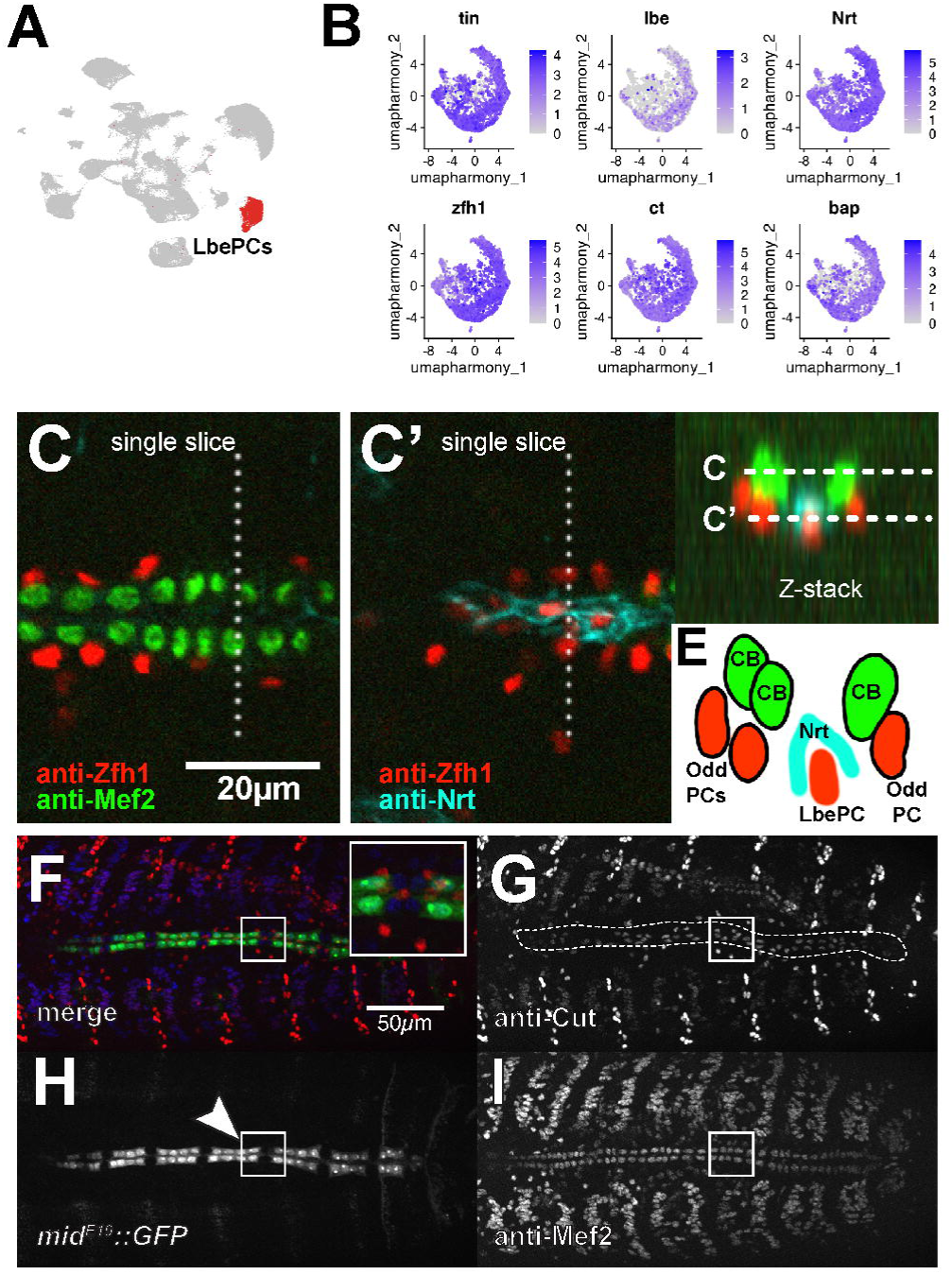

Furthermore, we find that the GPCR *Corazonine receptor* (*CrzR*) is also expressed in LbePCs as well as EvePCs already during dorsal closure of the heart and again throughout morphogenesis (**Supplemental Figure 5**). We also tested for the expression of the cell adhesion molecule and septate junction protein *Lachesin* (*Lac*), which is necessary for the integrity of the embryonic heart ^86^. *Lac* is expressed in the heart already at stage 15 (as well as in tissues outside the heart such as the developing trachea, **Supplemental Figure 5B**). However, especially during the final stages of heart morphogenesis *Lac* expression is increased especially in the EvePCs of the heart proper (**Supplemental Figure 5B**, arrowheads), indicating a region-specific function for Lac.

### Cell death – the sad fate of Eve- and LbePCs

Programmed cell death (PCD) is a common mechanism during animal development and organogenesis ^87,88^, including during mammalian heart development ^89^. The *Drosophila* heart was shown to become partially histolyzed during metamorphosis, when the posterior-most cardiomyocytes disappear while cells of the larval aorta partially remodel to form the adult heart ^90–92^. Cell death during embryonic or early larval fly heart development has not been observed, but while OddPCs and cardioblasts persist into larval and adult stages, the fate of both and Eve- and LbePCs has been unclear. Here, we find that both cell types upregulate a cell death gene program at the end of development, indicating that after completion of heart morphogenesis both EvePCs and LbePCs are terminally removed (**Figure 7**). We first noticed that both cell types have specific subclusters characterized by strong expression of the *Drosophila* cell death genes *head involution defective* (*hid*) and *reaper* (*rpr*) (**Figure 7**). Upon comparison of all upregulated genes in the *rpr-*positive clusters of both Eve- and LbePCs we find they share common signatures of phagosome formation, oxidative phosphorylation and indeed of apoptosis (**Supplemental Table 4**). Since we captured cells over a two-hour time window we wondered if the dying population of cells would be found in the oldest, most differentiated clusters, hinting at PCD as their final step of differentiation. We used ScVelo to calculate overall RNA velocities ^64^ and determined that these *hid*/*rpr*-positive cells indeed are at the distal end of the trajectories for both cell types (**Figure 7B**), as we would have expected based on the *in situ* expression of *reaper* in late PCs (**Figure 7A**). Next, we confirmed the expression of *rpr* by HCR *in situ* and co-labeling for *Hand* and *bap* (**Figure 7C**) and found bursts of *rpr* transcript in the hearts of very old (stage 17, gut fully developed) embryos, coinciding with *bap* transcript (**Figure 7C**) and spatially with dying posterior EvePCs based on their position (**Figure 7C**).

**Figure 7.**
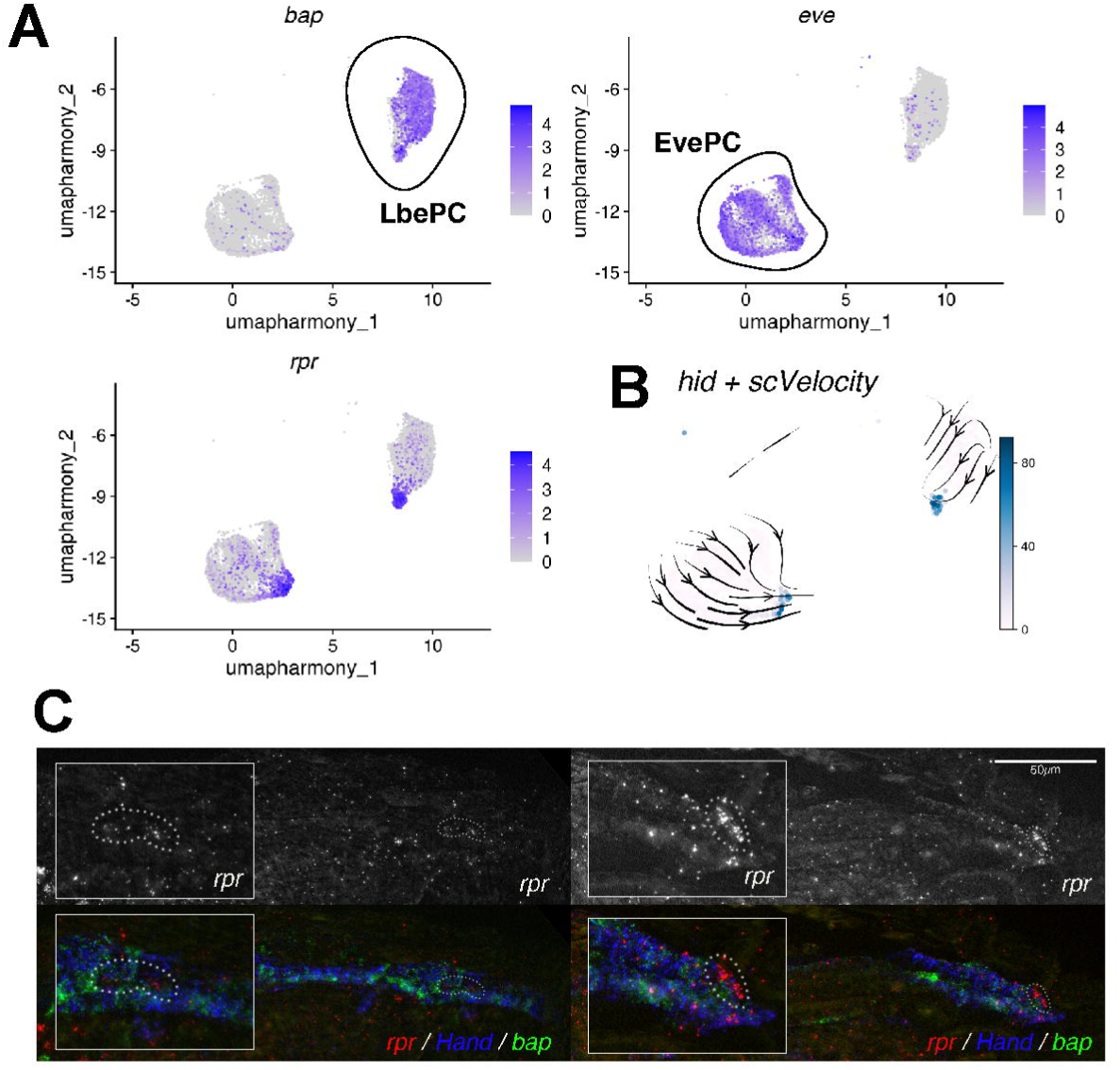

### RNAi-mediated knockdown of the cardiac TF *tinman* disrupts heart morphogenesis

The homeobox gene *tinman* is a conserved cardiac transcription factor ^22,23,93^ and critical for the development of visceral and cardiac mesoderm and their derivatives such as the gut musculature, dorsal somatic muscles and heart. Loss of *tinman* results in failure to form a heart ^22,23^ due to specification defects in the early (pre-cardiac) mesoderm hindering the analysis of *tin* function later in heart development. Analysis of *tin^LOF^* is thus limited, either to constitutive *tin* mutant analysis focusing on mesodermal targets and *tin*^LOF^ cell fates ^94^ and genetic interaction studies with *tin^LOF^* sensitized background ^95^. An alternative approach is using an elegant genetic rescue construct combined with *tin^LOF^* that lacks *tin* expression only in the developing heart, but early mesoderm-wide expresses of *tin* is unaffected ^63^. However, to avoid the required genotyping of sorted cells from F_1_ embryos after single-cell sequencing we instead relied on tissue-specific knockdown of *tin* using RNAi ^48,96^ driven by a strong stage- and tissue-specific driver line combination of *twist*-Gal4 with *how^24B^*-Gal4 ^97,98^. Loss of cardiac *tinman* was shown to dramatically increase the number of *Doc3*-positive cardioblasts, and pericardial cells appeared disorganized ^63^. During pupal and adult stages the heart proper histolyzes and disappears, followed by premature death.

We first phenotypically analyzed hearts from *twi;24B*-driven *tin*-RNAi embryos to identify morphological or specification defects caused by reduction of Tin. We found less Tinman protein in CB nuclei (**Figure 8A, B**), loss of Neurotactin localization (**Figure 8C-D’**), and expansion of Svp-positive cardioblasts (**Figure 8E, F**) in *tin*-RNAi hearts, confirming knockdown efficacy. Of note, the penetrance of *tin*-RNAi was not complete, and we observed phenotypic variability within each embryonic heart.

**Figure 8.**
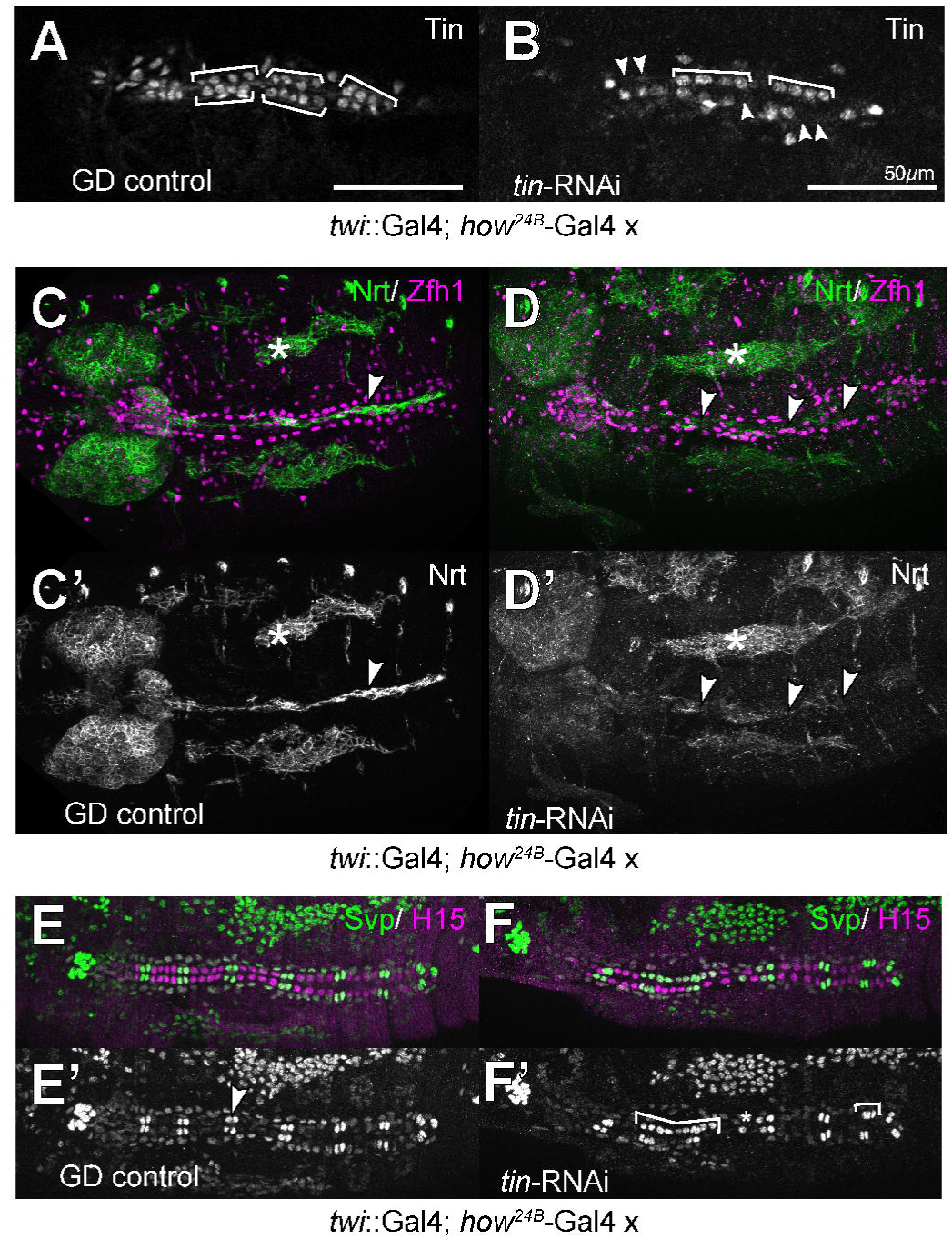

We next sorted cells from *tin*-RNAi embryos and obtained 38k cells from five replicates, and in parallel 27k cells of control embryos (GD background, four replicates) following bioinformatic analysis. We calculated for each cell type the number of differentially expressed genes (DEGs) between *tin*-RNAi and controls (**Supplemental Table 5**) using pseudo-bulk counts with edgeR ^99^. Of note, two genes, *Hsp70Bb* and *GstE11* were overexpressed in all tissues in the *tin*-RNAi background: *Hsp70Bb* likely as part of the UAS-*tin*^RNAi^ minimal promoter ^100^, and *GstE11* potentially close to the UAS-*tin*^RNAi^ insertion site (GstE11 is endogenously expressed in many tissues, **Supplemental Figure 1C,D**).

To control for specificity of *tin^RNAi^*, we reasoned that because *tin* is expressed only in a few tissues (e.g. not expressed in nervous system) *tin*-RNAi should not affect the *Hand^RFP^*-positive, but *tin*-negative cell types. We therefore tested if the number of significantly changed genes (FDR p-value < 0.05) is higher in normally *tin* expressing cell types, independent of the number of recovered cells. This was indeed the case, even for CBs despite the low numbers of cells recovered (**Supplemental Table 6**; see below). This *tin*-dependency of the number of DEGs also indicates that overexpression of *GstE11* did not alter gene expression patterns *per se* (see e.g. visceral mesoderm).

### Cardioblast gene expression changes

Our scRNAseq dataset contains 259 wild-type and 188 *tin-*RNAi cardioblasts, less than one-fifth of each pericardial subtype and likely due to specific difficulties in separating heart cells from their extracellular matrix. This low cell number and the associated reduced statistical power resulted in 41 differentially expressed genes (DEGs) between control and *tin*-RNAi (with FDR < 0.05; **Figure 9A, B**, **Supplemental Table 5**). A direct indication of altered Tinman activity is the downregulation of *tincar* (*tinc*), a transmembrane protein expressed in Tin+ CBs and repressed by Seven-up ^101^ that was upregulated in *tin-*RNAi embryonic hearts (**Figure 8**). We also identified the homeodomain transcription factor *Six4* ^102^ to be expressed in wild-type CBs and downregulated upon *tin* knockdown (**Figure 9C-D’**). Other dysregulated transcription factors include the muscle genes *lame duck* (*lmd*) and *spalt major* (*salm*), both of which were upregulated in *tin-*RNAi (**Supplemental Table 5**). Since we found an overrepresentation of DEGs in *tin*-positive, compared to *tin*-negative, cell types (see above), we decided in the case of CBs to relax the FDR threshold and to consider all DEGs with FDR <0.5, resulting in 109 potential DEGs. We then performed GO term analyses for enrichment of biological processes (**BP**), molecular function (**MF**) and **KEGG** pathways among these DEGs. For BPs, we found enrichment of genes belonging to cell surface receptor signaling pathway (18/761, p=0.003) as top GO term, followed by heparan sulfate proteoglycan biosynthetic process due to downregulation of 2/6 genes (*sulfateless - sfl*, *sugarless - sgl*). For MF, there was a significant enrichment for transcription factors (transcription cis-regulatory region binding, p=0.04) with 12 genes, including Six4, *salm* and *lmd* (see above). The Krüppel target gene *knockout* (*ko*) is another transcription factor prominently expressed in the heart ^103^ that is downregulated by *tin*-RNAi. Of note, most genes from these GO terms are significantly dysregulated with FDR below 0.05 (**Supplemental Table 7**).

**Figure 9.**
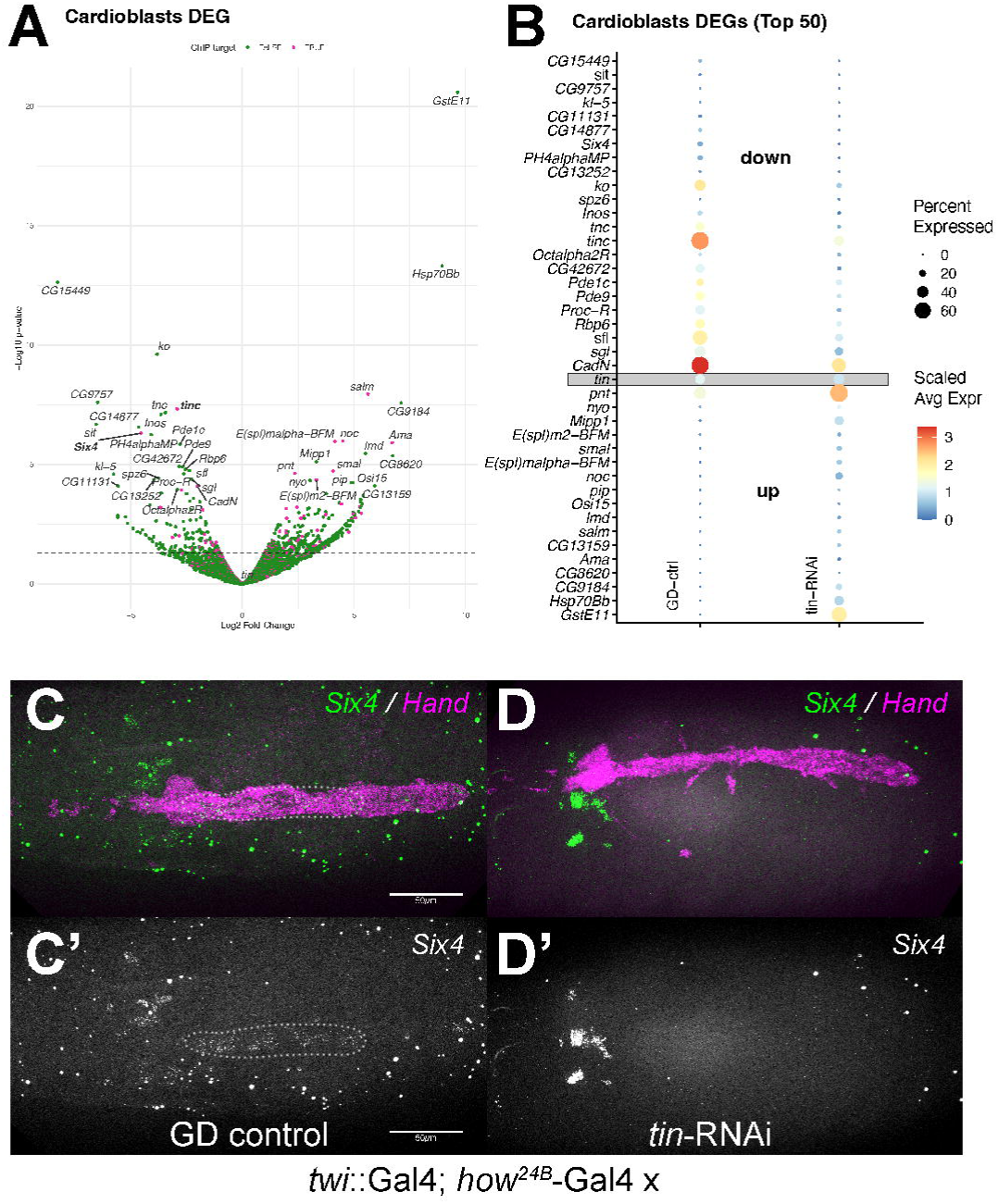

### Eve- and Lbe-PC gene expression changes

In our dataset we recovered a total of 1481 Eve-PCs and 1334 Lbe-PCs from 4 GD control samples, versus 3695 Eve-PCs and 1817 Lbe-PCs from 5 *tin-*RNAi samples (**Supplemental Table 8**). Differential gene expression analysis with edgeR resulted in 1015 DEGs for EvePCs and 1278 DEGs for LbePCs with FDR < 0.05 (**Figure 10**, **Supplemental Table 5**). 395 genes were shared between both cell types, including 363 with identical change in directionality of expression (185 up/up and 178 down/down, **Supplemental Figure 9A, B**). This indicates that despite their molecular-genetic differences knockdown of *tin* affects a common subset of genes in Eve- and Lbe-PCs.

**Figure 10.**
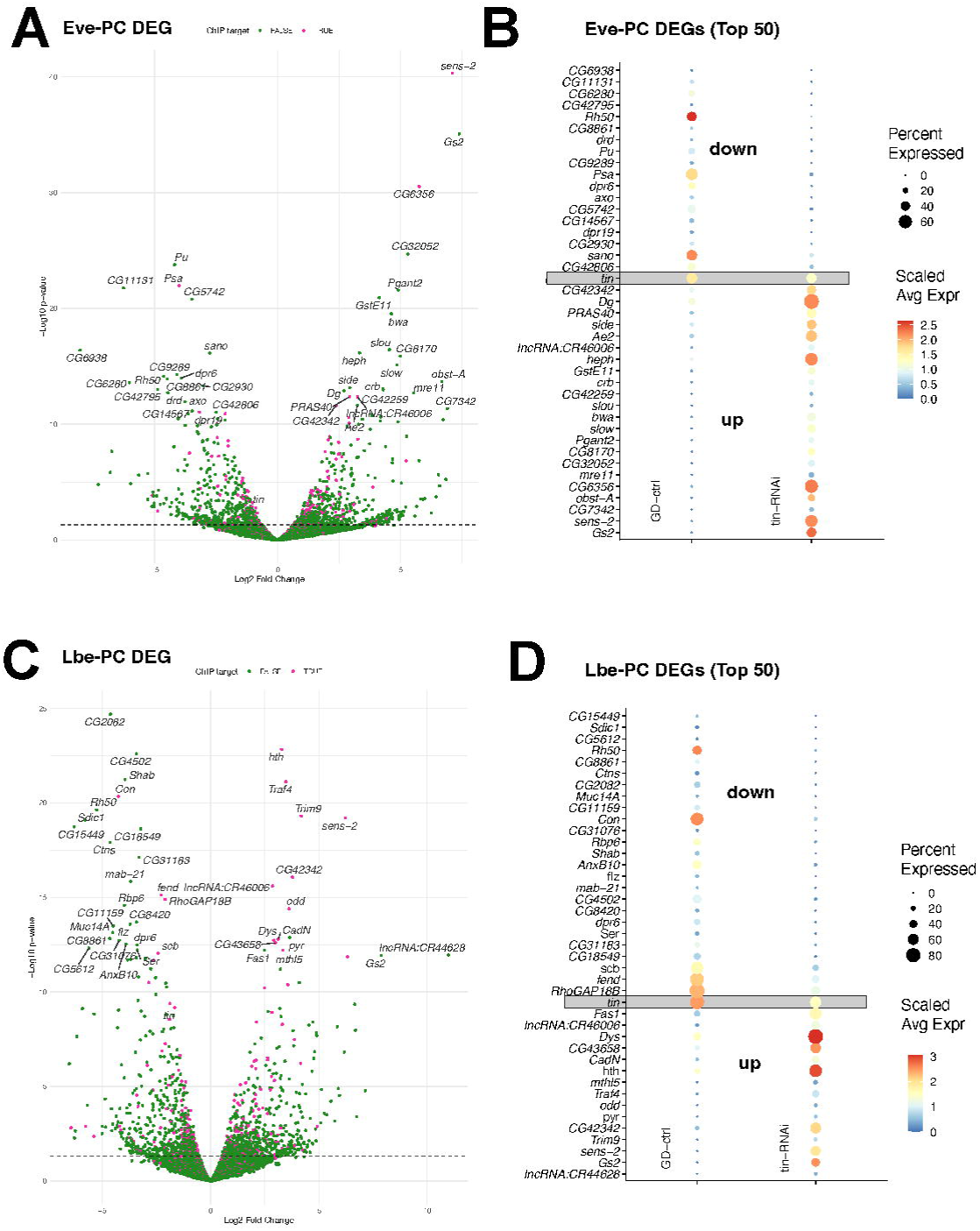

Next, we performed GO term analysis on DEGs for all cell types (**Supplemental Table 7**). DEGs of EvePCs and LbePCs are highly enriched for genes involved in motor neuron axon guidance, cell surface receptor signaling and homophilic cell adhesion via plasma membrane adhesion molecules (all biological processes, **Supplemental Table 7**). Both cell types upregulate genes involved in muscle targeting of motoneurons ^81,104–106^, specifically the Sidestep Ig-domain proteins *side, side-IV* (and in EvePCs also *side-V*), downregulate the transmembrane protein *forked end* (*fend*), and switch the expression of *side*-interacting genes from *beat-IIa* (EvePC) and *beat-IIb* (Eve- and LbePCs) to *beat-IIIc* (**Supplemental Figure 10**). In addition, the collagen XV/XVIII *Multiplexin* (*Mp*, expressed in the embryonic nervous system and the heart proper ^59,107^) is strongly upregulated upon *tin-*RNAi. *Capricious* (*caps*) is a cell adhesion molecule involved in axonal targeting of muscles and is expressed in both neuron and muscle subsets ^108^, including the Eve-positive aCC, RP2 and U neurons of the ventral nerve chord as well as the Eve-lineage derived DA1 muscles. In both Eve- and LbePCs *tin-*RNAi causes significant upregulation of *caps* expression, indicating that Tinman suppresses the expression of non-PC genes that are normally expressed e.g. in Eve-positive neurons and muscle cells.

Common changes of other cell adhesion molecules of both cell types are downregulation of *shotgun* (*shg*, *Drosophila* E-Cadherin), upregulation of *Cadherin 96Ca* (*Cad96Ca*) and the IgLON ortholog *CG13506*. In addition, LbePCs with knockdown of *tin* upregulate *Cadherin 87A* (*Cad87A*) and *Cadherin-N* (*CadN*), which is normally expressed only in CBs among cardiac cells. LbePCs show a change in integrin signaling: downregulation of *scab* (*scb, alphaPS3*) and *Integrin alphaPS4*, (*ItgaPS4*) and upregulation of *multiple edematous wings* (*mew*, *alphaPS1*) and *Tiggrin* (*Tig* integrin ligand). They also show upregulation of *Fasciclin 1* (*Fas1*), and downregulation of *Fasciclin 3* (*Fas3*), as well as upregulation of *Dystroglycan* (*Dg*), altogether resulting in an altered cell adhesion code compared to wildtype. It will thus be interesting to see if their combined manipulation will alter cardiac morphology.

### scRNA-seq of *tinman* uncovers direct and indirect Tinman targets

Enhancer occupancy studies are an invaluable tool to identify how transcription factors regulate target gene expression and to delineate gene regulatory networks. This has previously been applied to Tinman mainly via chromatin immunoprecipitation ^32–34^, with complementary approaches such as open/closed chromatin analysis and machine learning ^35,36^ and functional validation by enhancer/reporter gene studies. This revealed many potential direct Tinman target genes, but not in a cell-type specific fashion and not differentiating, e.g. between CBs and LbePCs. Furthermore, only a few target genes were functionally tested, e.g. by examining their expression in a cardiac-specific mutant (*tin^ABD^*; *tin^LOF^*) ^32,63^. It is therefore unclear for most genes controlled by an enhancer bound by Tinman if Tinman acts as activator or repressor.

We have found that cardiac cells have significantly more DEGs upon *tin*-RNAi compared to non-cardiac, *tin*-negative cell-types (**Supplemental Table 9**) indicating direct or indirect dependency of gene expression on Tinman. Focusing on marker genes for each cell-type (determined by *FindAllMarkers* default parameters, followed by filtering for average log2FC > 0.5, adjusted p-value < 0.05, and expressed in more than 30% of cells per cell-type) we calculated for each tissue the odds of a cell marker being downregulated upon *tin*-RNAi. For Lbe-PCs, 68 marker genes were downregulated and 2 upregulated (odds ratio 37.6; Eve-PCs: 53 down, 1 up, odds ratio 72.4; **Supplemental Table 9**). While these two cell-types were the two most significant, the only other cell types that show significant changes in marker gene downregulation are odd-PCs and CBs. This indicates that Tinman is necessary for the expression of these marker genes and thus maintaining cellular identity.

We wondered whether up- or downregulation in one cell-type caused the opposite misregulation in another cell-type. Interestingly, only 17 genes showed such differential regulation (**Supplemental Table 10**), suggesting that in general each cell-type is uniquely regulated. Across all cell types, 171 DEGs out of 1447 genes have been shown to bind Tinman ^32,34^. With respect to dysregulation of motoneuronal axon guidance genes such as the Sidestep and BEAT gene families that we found upregulated in EvePC and LbePCs, all have been shown to bind Tinman during embryonic development (**Supplemental Figure 11**). Thus, the role of Tinman extends beyond activating cardiac gene programs but apparently to also orchestrates suppression of e.g. motor axon guidance programs.

### *Tinman* suppresses a wing heart program in Eve- and LbePCs

The zinc-finger transcription factor *senseless-2* (*sens-2*) is the most-upregulated transcription factor in *tin*-RNAi EvePCs and LbePCs, indicating that Tinman directly suppresses *sens-2* in these cells (**Figures 10, 11A, Supplemental Table 5**). Among dorsal *eve-*positive cells, *sens-2* is specifically expressed in the wing heart precursors (WHP, **Figures 5F, 11A**). We directly confirmed the ectopic expression of *sens-2* in *Hand*-positive cells of the entire heart in *tin*-RNAi embryos (**Figure 11A**), in addition to wild-type expression of *sens-2* in the anterior-most *Hand*-positive wing heart precursor cells that are *tin*-negative and thus not affected by the knock-down (**Figure 11A**). Both ChIPseq datasets ^32,34^ show binding of Tinman inside the fourth intron of *sens-2* (**Supplemental Figure 11**), and it was shown to have a classifier-predicted functional cardiac enhancer ^35^ active in CBs and PCs, which suggests that Tin directly represses *sens-2* expression.

**Figure 11.**
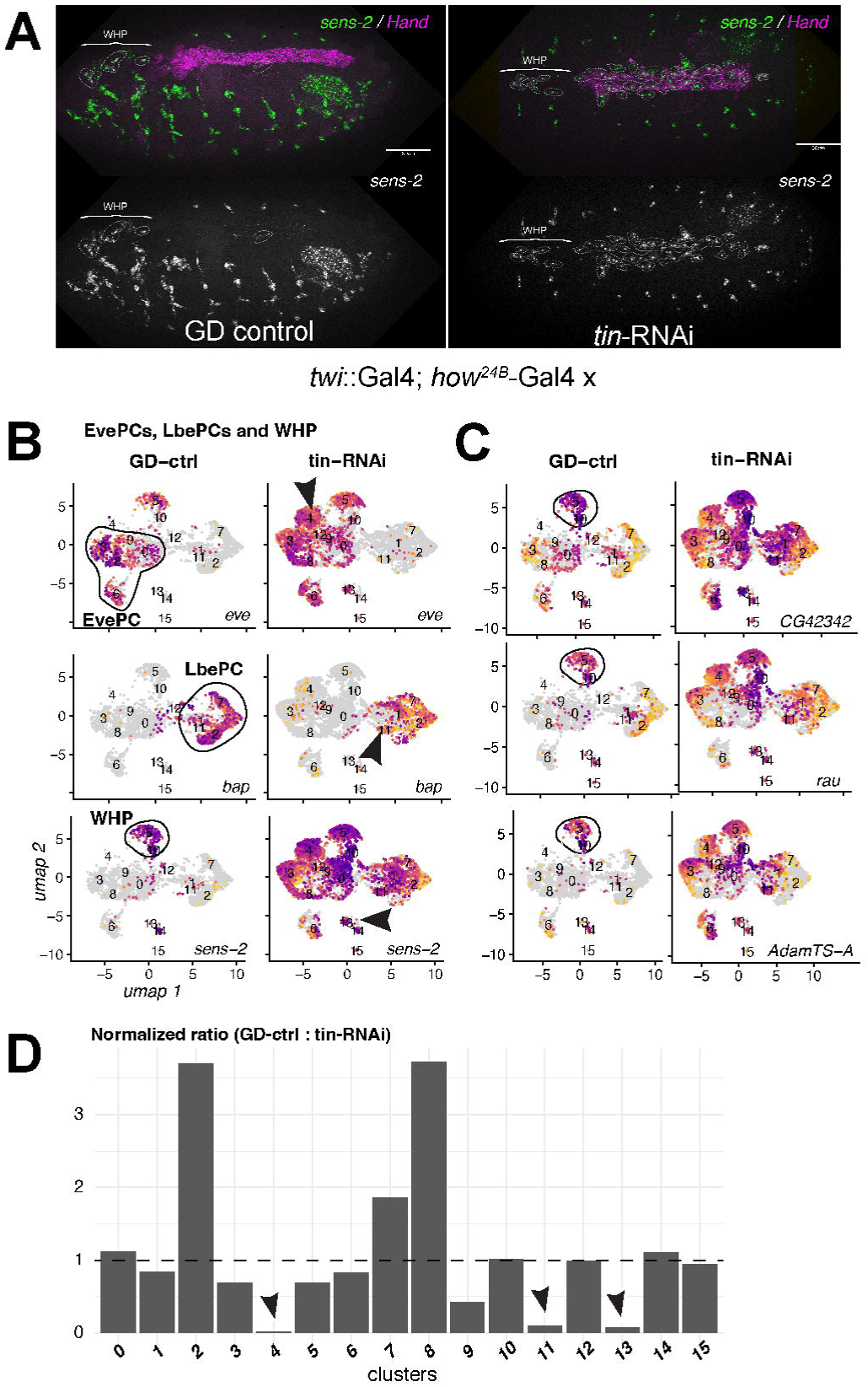

We next analyzed if *tin* repressed only the expression of *sens-2* or if we find other WHP genes to be ectopically expressed. Comparing the joint UMAP of EvePCs, LbePCs and WHP in control and *tin*-RNAi (**Figure 11B, C**) we find broad expression of *CG42342*, *rau* and *AdamTS-A* in *tin*-RNAi in all cells, while they are normally expressed only in WHP cells (**Supplemental Figure 7C**). This indicates that Tinman restricts the expression of a wing heart gene program in cells not meant to become WHPs. Supporting this hypothesis, it was previously shown that these cells do not express *tinman* later in development, and that overexpression of Tinman in *Hand*-positive cells abolishes the formation of WHP cells ^57^.

We also noticed that the relative population of WHP cells in *tin*-RNAi was significantly increased (**Figure 11D, Supplemental Figure 7A, B**), and specific clusters where absent in wild-type (clusters 4, 11, 13) or diminished in *tin*-RNAi (clusters 2, 8, **Figure 11D**, **Supplemental Figure 7A**). We speculate that these cells are likely incompletely transformed Eve-/LbePCs at their original position (**Figure 11A**), that would eventually co-cluster along wildtype WHPs. Indeed, we find cells in the WHP cluster in *tin*-RNAi but express posterior Hox gene markers *Ubx* and *Abd-B* (**Supplemental Figure 12**). We therefore conclude that there is a wing heart gene program that is repressed by Tinman, either directly, or indirectly through other TFs such as *sens-2*.

### Loss of *tinman* causes a receptor switch in EvePCs

During embryonic heart development the Wingless/WNT signaling pathway is crucial for induction of the cardiac mesoderm, and specification and differentiation of cardiac cells by canonical and non-canonical signaling mechanisms ^13,109–113^. *wingless* (*wg*) and several paralogs are expressed in embryonic cardiac cell types: *wg* and *Wnt6* in Svp-positive cardioblasts, *Wnt4* in CBs (see also ^61,62^), as well as in LbePCs and an OddPC subset (this study), and *Wnt5* at low levels in all cells (**Supplemental Figure 13**).

Wingless/WNT ligands act via the Frizzled receptor gene family that includes *frizzled* and *frizzled-2* (*fz* and *fz2*), both of which are necessary for heart formation in *Drosophila* ^114^. In our wild-type single cell dataset, both Fz receptors are expressed in specific cardiac cell-types: CBs and WHPs express both receptors, OddPCs and LbePCs only express *fz*, and EvePCs express only *fz2* (**Figure 12A, Supplemental Figure 14**). Among *eve*-positive cells, EvePCs can be identified by the expression of *esg* (see above, and **Figure 12B**), which co-express *fz2* but not *fz*, confirming the scRNAseq observation. Following knockdown of *tinman* with RNAi we noticed a striking change in *frizzled* receptor expression specifically in EvePCs: *fz* becomes strongly induced, while *fz2* is downregulated in these cells. The only other cell type that shows significant upregulation of *fz* are WHPs, but here we do not find changes in *fz2*. We validated this receptor switch *in situ* and indeed observed induction of *fz* in *esg*-expressing EvePCs, while at the same time *fz2* is largely gone (**Figure 12B-C’’**). Of note, *fz* and *fz2* have been shown to bind Tinman ^32,34^, indicating that in the context of EvePCs Tinman acts as suppressor of *fz* and an activator/maintaining factor of *fz2*.

**Figure 12.**
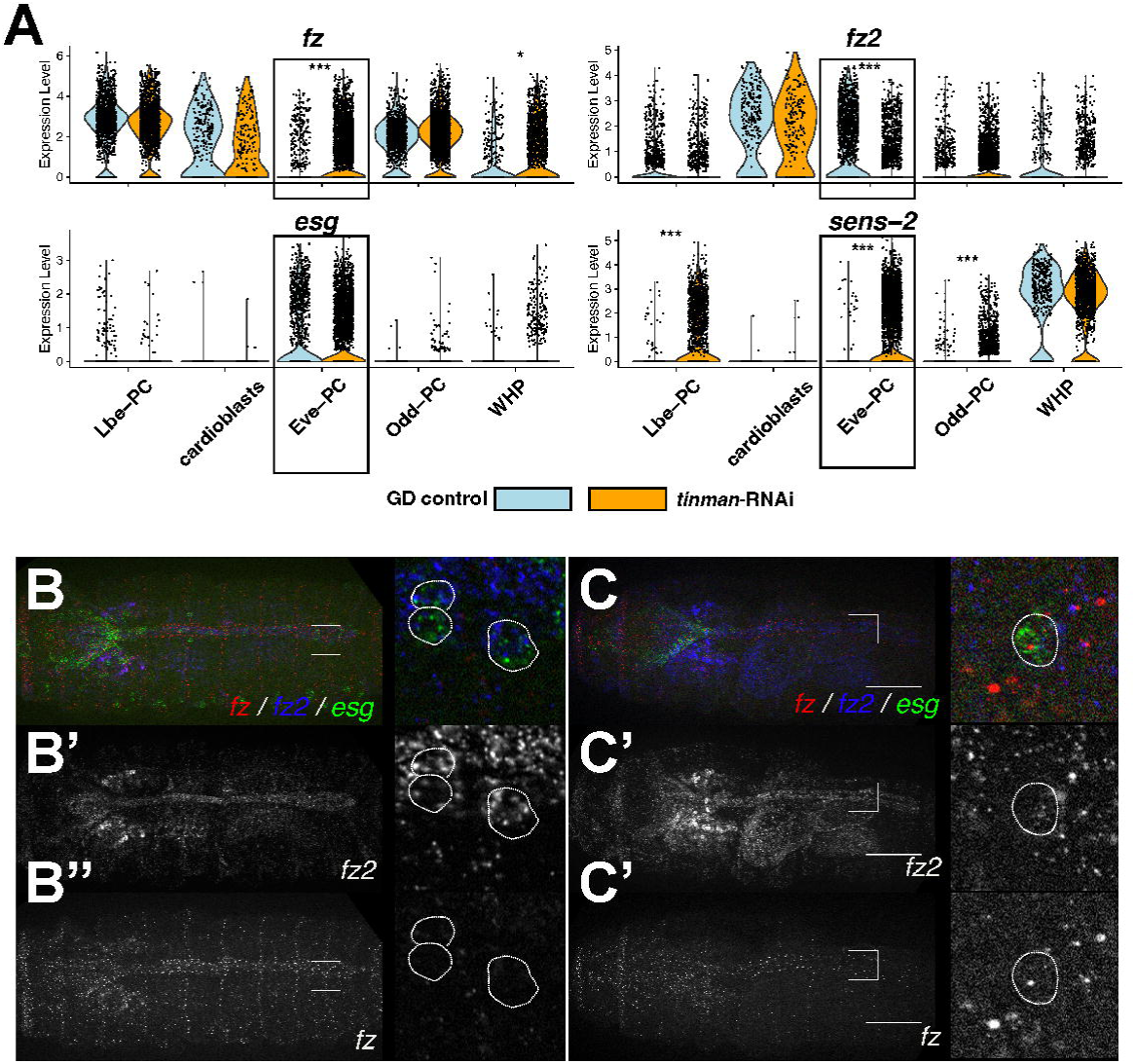

Taken together, by analyzing cell-specific effects of *tinman*-RNAi during fly cardiogenesis we identified specific gene programs controlled by Tinman either directly or indirectly that are critical for heart morphogenesis and differentiation, beyond what is known on Tin during early cardiogenesis. This functionally annotated single-cell atlas of the *Drosophila* heart thus is an important resource to further explore gene regulatory networks and the role of individual cardiac cell types during heart development.

## DISCUSSION

Our study presents a comprehensive single-cell RNA sequencing atlas of the developing *Drosophila* embryonic heart, using a *Hand* reporter gene to isolate and analyze 92,000 cells across multiple cell types including cardioblasts, various pericardial cell populations (Odd-PCs, Eve-PCs, Lbe-PCs), and wing heart precursors. We identified novel marker genes and signaling pathways for each cardiac cell type, discovered that Eve-PCs and Lbe-PCs undergo programmed cell death at the end of embryonic development, and revealed temporal dynamics during cardioblast-to-cardiomyocyte differentiation. Through functional analysis using RNAi-mediated knockdown of the master cardiac transcription factor Tinman, we demonstrated that Tinman not only activates cardiac-specific gene programs but also actively suppresses inappropriate gene expression, particularly neuronal/motor neuron axon guidance genes and wing heart developmental programs in non-wing heart cardiac cells. Additionally, we found that Tinman controls a receptor switch in Eve-PCs, maintaining *frizzled-2* expression while suppressing *frizzled* expression, which likely has implications for Wingless/WNT signaling during heart morphogenesis. This work provides unprecedented molecular detail of cardiac cell diversity and the regulatory networks controlling heart development in *Drosophila*.

The rise of highly resolved RNA sequencing has caused an ever-growing number of single-cell atlases from a large range of organisms, model systems, tissues and experimental and disease-states. This allowed to identify the cellular diversity and their states inside of different organs including the human ^115,116^, murine ^117,118^, zebrafish^119,120^ and fly hearts ^121^.

We identified programmed cell death of Eve-PCs and Lbe-PCs at the completion of heart morphogenesis, overturning the assumption that all *Drosophila* cardiac cell types persist beyond embryogenesis. RNA velocity provides evidence that these cells represent the most differentiated state, indicating that their elimination is a regulated terminal program rather than a developmental accident. We have previously found that the EvePCs influence cardiac morphogenesis and function ^56,122^. The occurrence of apoptosis during mammalian heart development is well established ^89,123–125^, but whether similar transient cell populations exist that contribute to heart morphogenesis has not been investigated in the *Drosophila* heart model and is thus an exciting possibility.

Our analysis of Tinman reveals a dual activator–repressor role in maintaining cardiac identity. In addition to activating cardiac programs, Tinman directly represses inappropriate neuronal and wing heart gene expression, as shown by ectopic marker induction upon Tinman knockdown and ChIP-seq evidence of direct binding. This active suppression explains how a broadly expressed transcription factor preserves cell-type specificity, highlighting repression as a central principle of lineage maintenance. The Tin ortholog NKX2-5 has also been shown to both activate and repress specific target genes ^126–131^, and is likely that NKX2-5 also controls specific programs. Interestingly, during endothelial-to-hematopoietic transition (EHT), NKX2-5-positive angioblasts differentiate into hematopoietic cells ^132,133^, which is controlled by GFI1/1B. Since GFI1 is critical for angioblast differentiation and we found its ortholog *sens-2* likely involved in wing heart precursor differentiation, we speculate that these tissues might share a common regulatory network. Previous studies on the role of *tinman* post-cardiac specification indicate distinct de-repression of genes normally expressed in Tin-negative ostia precursors (e.g. *Dorsocross-3* ^63^). In contrast to their findings, we also observe ectopic expression of Seven-up, either because of ectopic expression in Tin-CBs or due to misspecification during lineage determination.

We also uncover a Wingless receptor switch in Eve-PCs upon Tinman loss, where *frizzled-2* is replaced by *frizzled*. Because these receptors activate distinct downstream pathways ^134,135^, this switch likely fine-tunes cellular responses to morphogen gradients. Although it has been found that both, *fz* and *fz-2* can act redundantly during development ^114^, the distinct control of Fz receptor expression by Tinman and the specific expression pattern suggests such a fine-tuning activity during heart morphogenesis and differentiation. Together with the cell-type–specific distribution of Wingless ligands and Frizzled receptors, these findings reveal a complex signaling landscape that coordinates cardiac morphogenesis.

Temporal analysis delineates the transition from cardioblasts to differentiated cardiomyocytes, marked by a shift from ribosomal to sarcomeric gene expression. The inverse correlation between Myc-driven ribosomal genes and contractile muscle genes likely reflects a metabolic switch from growth and proliferation to assembly of the contractile apparatus. Myc has several functions during myoblast differentiation and muscle growth ^136–138^ and they likely depend on the specific muscle-cell type (e.g. smooth muscle vs. cardiomyocyte). Our data provides a clear basic model of how Myc expression precedes ribosome expression which in turn precedes sarcomeric gene expression, thus reflecting a cardiomyocyte differentiation pathway.

The identification of novel marker genes for each cardiac subtype provides tools for functional studies and uncovers unexpected pathways, including axon guidance programs enriched in cardiac populations and neurotransmitter receptor expression in ostia cells of the myocardium. These observations suggest shared molecular logic between neural and cardiac development, with potential roles for neuronal guidance and neuromodulation genes in heart morphogenesis. Cell adhesion and axon guidance molecules play important roles during tissue morphogenesis and homeostasis ^139,140^. In *Drosophila,* the role of the Slit-Robo pathway in heart development was established first ^141–143^, and was later found to be active also in vertebrate heart development ^144^.

Another well known axon guidance pathway, Netrin-Unc5, is expressed during heart development in both flies ^145–148^ and mice ^149^. Of note, while functional analysis of *Fasciclin 3* (*Fas3*) showed that it is necessary for robust cell matching during CB dorsal closure ^150^ our dataset suggests that an intricate interplay of different cardiac cell types, each expressing a unique combination of cell adhesion molecules that is at play during cardiac morphogenesis. Especially the BEAT/side pathway ^139,140^ is likely a novel player during fly heart development. This lets us hypothesize that cardiac morphogenesis is controlled by appropriate cell matching of all cardiac cell types, and programmed cell death of two cardiac subtypes, Eve- and Lbe-PCs, functions as a developmental timer indicating the end of dorsal heart closure signaling for terminal cardiomyocyte differentiation.

Despite technical limitations—including low cardioblast recovery, coarse temporal resolution, and incomplete, but nevertheless informative Tinman knockdown—our dataset provides a foundation for future studies. Key questions include whether blocking programmed cell death alters heart function, whether analogous transient populations exist in mammals, and how receptor switching contributes to signaling specificity.

In sum, single-cell analysis reveals unanticipated complexity in the *Drosophila* heart, exposing new paradigms of transcription factor function, transient cell populations, and signaling dynamics that parallel vertebrate systems. These insights establish a framework for dissecting the gene regulatory networks that govern cardiac development across species.

## METHODS

- METHODS DETAILS

Reporter gene construction and fly lines
Single cell isolation, encapsulation and RNA sequencing

We adapted the protocols to prepare a cell suspension from primary from ^151,152^. Embryos were dechorionated using 4% bleach for 3 min and transferred into a 1ml Dounce homogenizer containing 500ml Accumax (Innovative Cell Technologies). Embryos were homogenized on ice using 10 strokes (‘tight’ pestle) and incubated at room temperature for 15min. The suspension was vortexed briefly every 5min to resuspend any cells that have settled. The suspension was then poured over a 100µm cell strainer and the flow-through collected in a 50ml Falcon tube. The Dounce homogenizer was rinsed with another 0.5ml of Accumax and added to the 100µm filter. Next, we poured the filtered suspension through a 30µm MACS filter (Miltenyi Biotech) and the flow-through collected in a 15ml Falcon tube. Cells were centrifuged for 3min at 300g and the cell pellet was washed once with 1.5ml of Accumax (or Schneider’s medium or 1% BSA/1xDPBS during optimization steps) and stored on ice.

Cells were sorted on a SORP FACSAriaIIu (BD Biosciences, San Jose) with 100 µm nozzle, 23 psi sheath pressure, and 32.7 kHz drop frequency and FACSDiVa v8.0.3 software. Standard excitation and filters were used for both GFP and DAPI. Autofluorescence was assessed with 561 nm excitation and 610/20BP emission filter. Both sample chamber and collection tubes were kept at 4°C for the duration of the sort. FACSDiVa and FlowJo v10 were used for data analysis.

### Single cell analysis pipeline using R

Single-cell RNA sequencing data from multiple samples were processed using a comprehensive computational pipeline (R/Seurat v5^153^). Raw sequencing data from Cell Ranger (10x Genomics) outputs were imported as filtered feature-barcode matrices in HDF5 format, with genes showing zero expression across all cells removed during initial processing. SoupX ^154^ was applied to each sample individually to estimate and remove ambient RNA contamination, using pre-clustering with SCTransform normalization followed by PCA, UMAP, and graph-based clustering to guide the contamination estimation. Doublet detection was performed using scDblFinder^155^.

Quality control metrics (QC) were calculated for each cell, including mitochondrial and ribosomal gene percentages using Drosophila-specific gene annotations, and cell cycle scores were computed using UCell with custom Drosophila cell cycle gene sets derived from published literature. Cells identified as doublets were separated into distinct objects for potential downstream analysis, while singlet cells were retained for the main analysis pipeline. All samples underwent SCTransform normalization using the glmGamPoi method with v2 flavor, regressing out ribosomal percentage, mitochondrial percentage, and cell cycle difference scores.

Integration of multiple samples was performed using two complementary approaches to ensure robust results. FastMNN integration was applied to the first 30 principal components to correct for batch effects while preserving biological variation, followed by Harmony integration using the same dimensional space. Both integration methods generated separate UMAP embeddings for visualization and downstream analysis. Graph-based clustering was performed to identify cell populations.

### Sub-clustering and sub-cell type identification

Cell type annotation was performed through a combination of known marker gene expression and differential gene expression analysis. Cluster analyses and markers used for cell type annotation in Supplemental Methods Table **1**.

### Differential Analysis of *tinman* RNAi cells

Differential expression analysis was conducted using a pseudobulk approach, employing edgeR ^99^ for more robust statistical inference by aggregating expression data across biological replicates within each condition and cluster combination. Genes with FDR < 0.05 were considered significant (exception: cardioblasts).

### Gene Ontology Enrichment Analysis

DEGs were filtered to retain only genes with false discovery rate (FDR) < 0.05. Gene ontology (GO) enrichment analysis was performed using the gprofiler2 package ^156^. Enrichment testing was conducted against three ontology databases: Gene Ontology Molecular Function (GO:MF), Gene Ontology Biological Process (GO:BP), and Kyoto Encyclopedia of Genes and Genomes (KEGG) pathways. Multiple testing correction was applied using the false discovery rate (FDR) method, with evidence codes retained for detailed annotation. To reduce redundancy in enriched GO terms, semantic similarity-based simplification was performed using the compEpiTools package (maximum overlap threshold of 0.1 for biological process ontology terms), retaining the most representative terms while removing highly similar or redundant annotations.

### Immunohistochemistry

- HCR in situ

HCR RNA in situ hybridization was done according to the manufacturer’s protocol for *Drosophila* embryos (Molecular Instruments). Probes were synthesized for the following genes (using the provided GenBank accession numbers): *CG6415* (NM_135597.3); *H15* (NM_135082); *CrzR* (NM_140314.4); *Lac* (NM_078989.3). YFP/GFP probe *d2eGFP* was obtained from the manufacturer. Custom HCR probes were designed using (https://zenodo.org/records/15537771), and probe-pair oligos were synthesized by IDT’s Pooled DNA oligos service (IDT, USA) for the following transcripts: *bap* (NM_169958), *eve* (NM_078946), *fz* (NM_080073), *fz2* (NM_079431), *Hand* (NM_135526), *H15* (NM_135082).

Images were acquired using a C-Apochromat 40×/1.2 NA water immersion objective lens on an AxioImager Z1 equipped with an Apotome.2 (all Carl Zeiss) and a OrcaFlash4.0LT camera (Hamamatsu), with ZEN software 2.3 software (Carl Zeiss). Image processing was done using ImageJ/FIJI ^157^.

## ACKNOWLEDGMENTS

We appreciate the support of the Sanford Burnham Prebys Shared Resources, including Amy Cortez, Ziyi Sang, and Benji Portillo (Flow Cytometry), and Rebecca Porritt (Genomics).

## DECLARATION OF INTERESTS

The authors declare no competing interests.

---

**Supplemental Figure 1.**
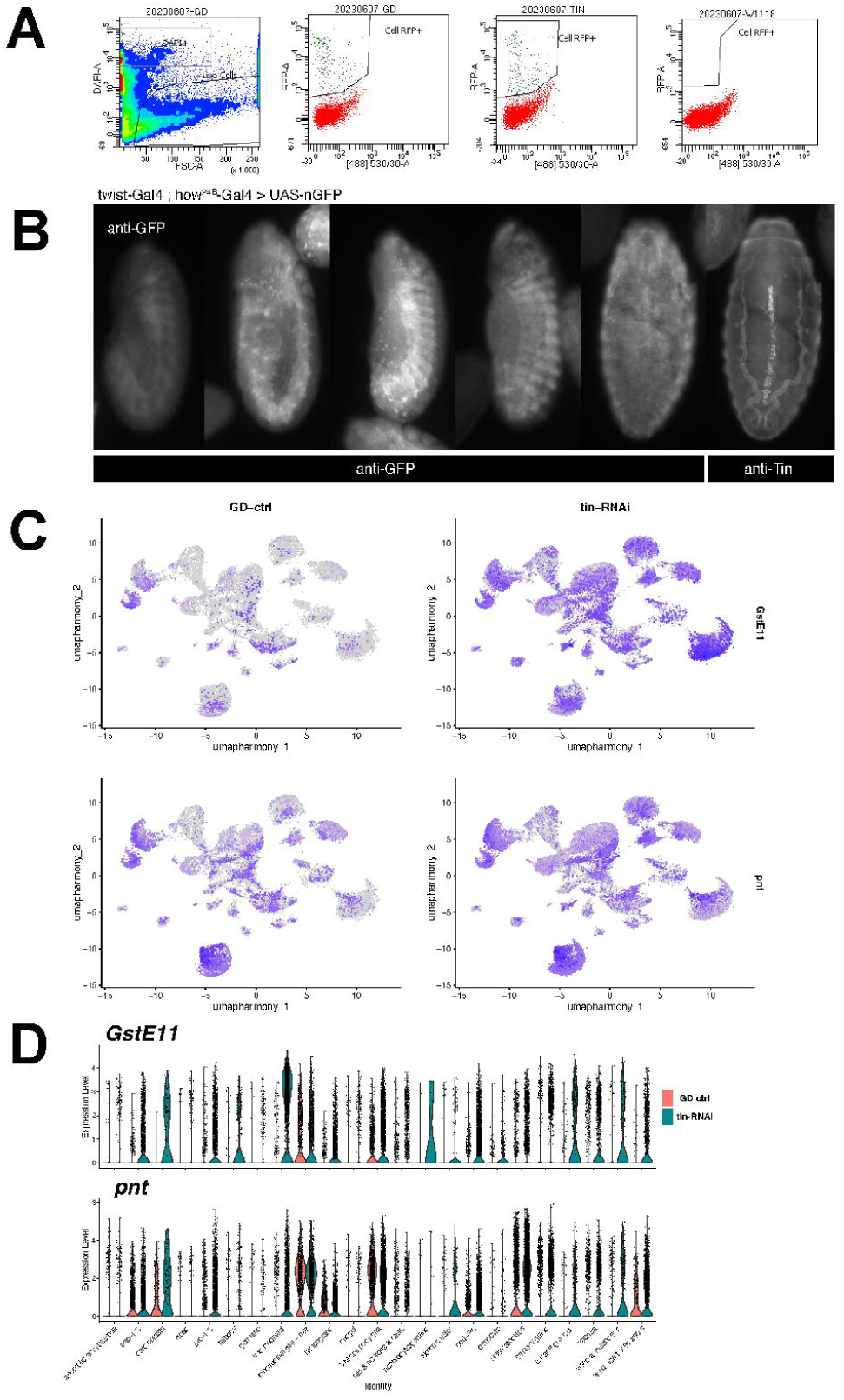

**Supplemental Figure 2.**
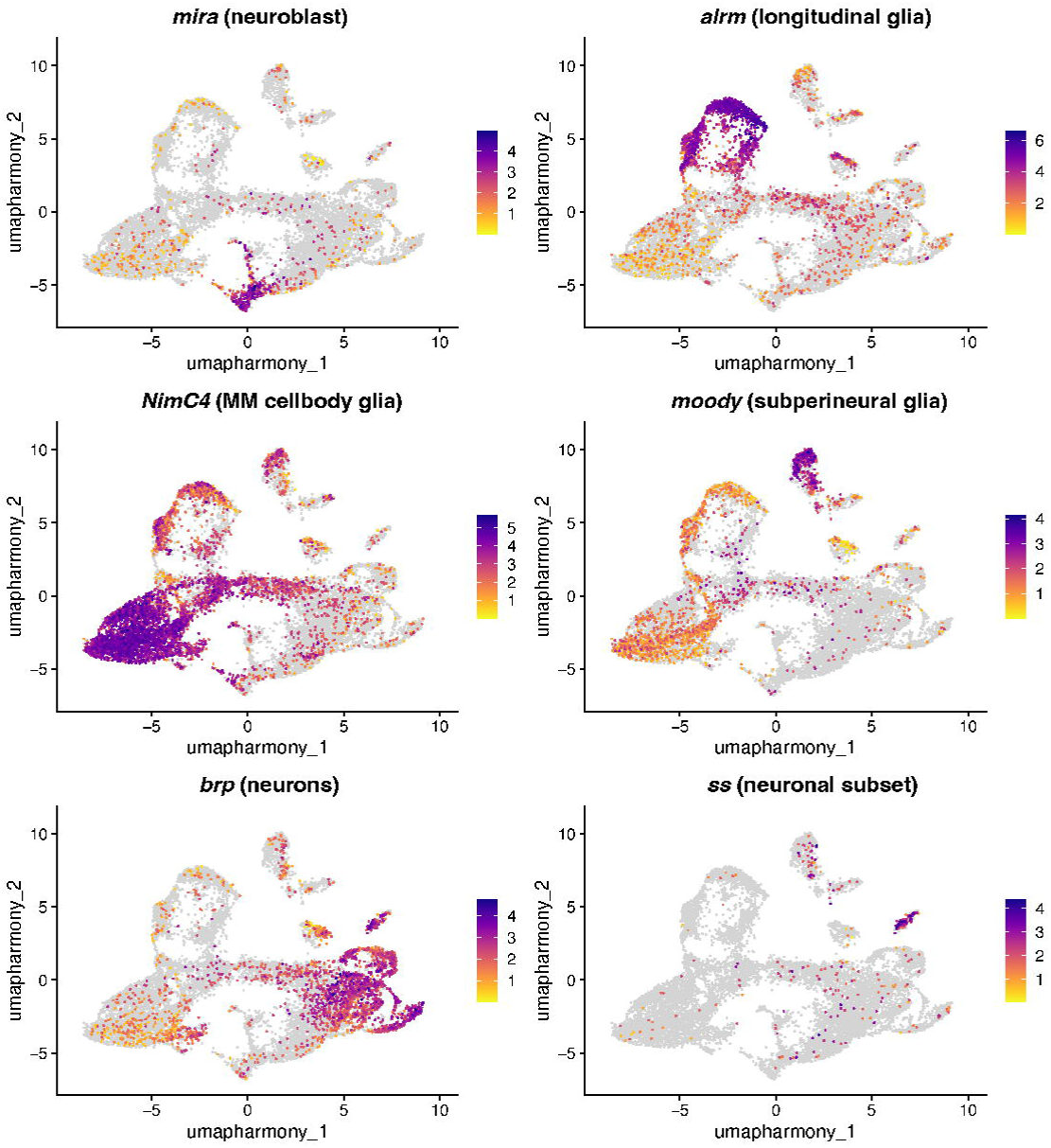

**Supplemental Figure 3.**
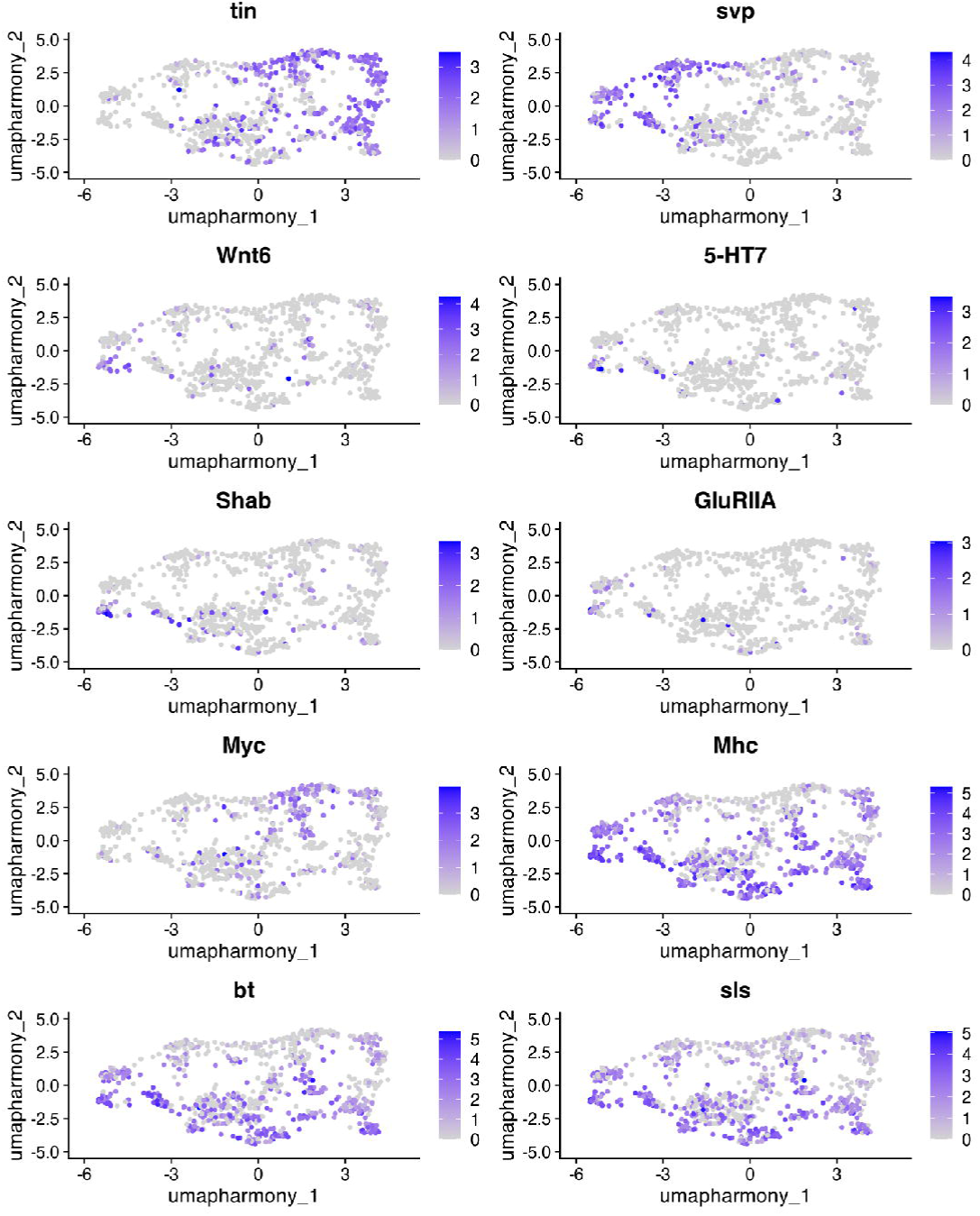

**Supplemental Figure 4.**
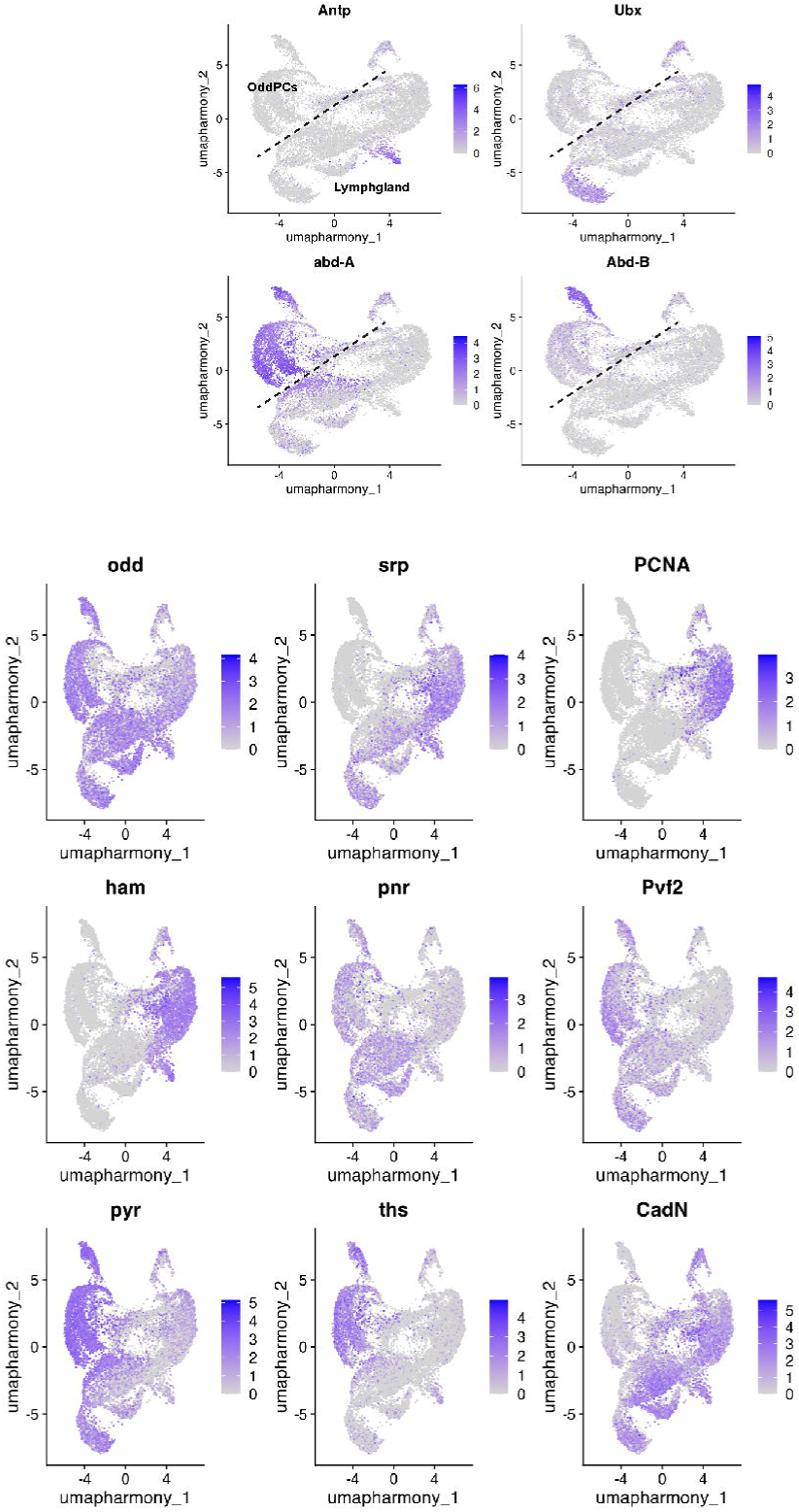

**Supplemental Figure 5.**
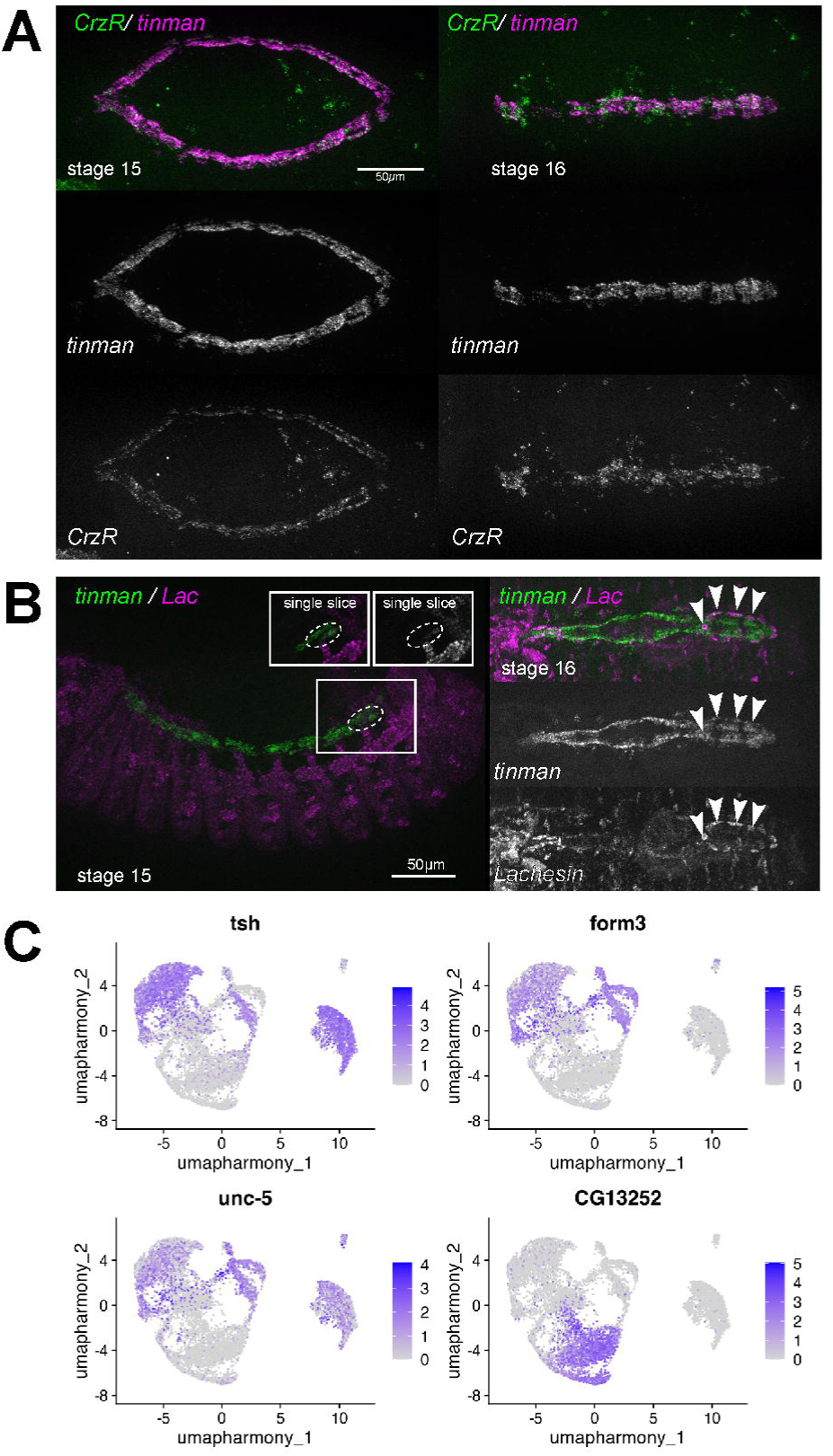

**Supplemental Figure 6.**
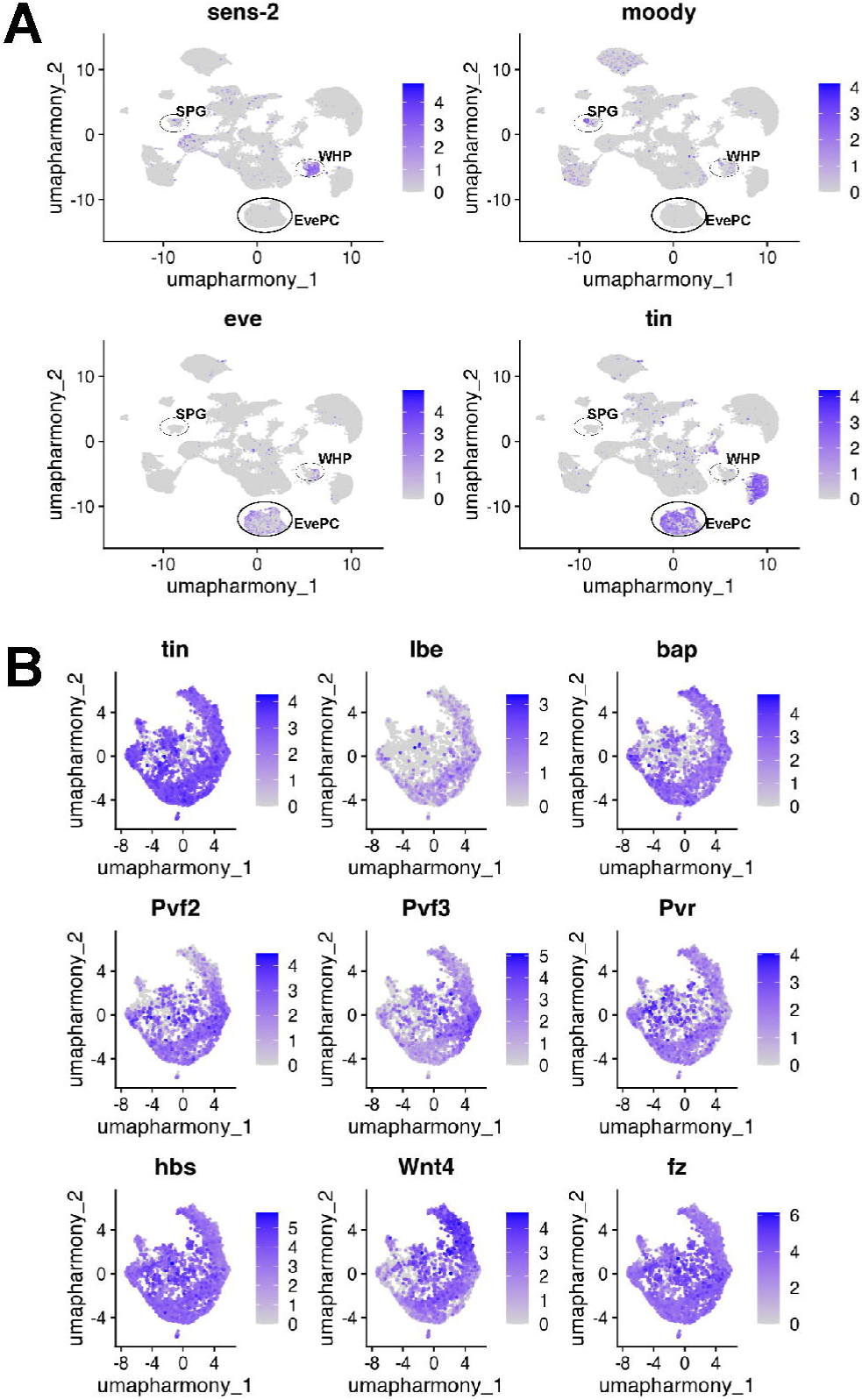

**Supplemental Figure 7.**
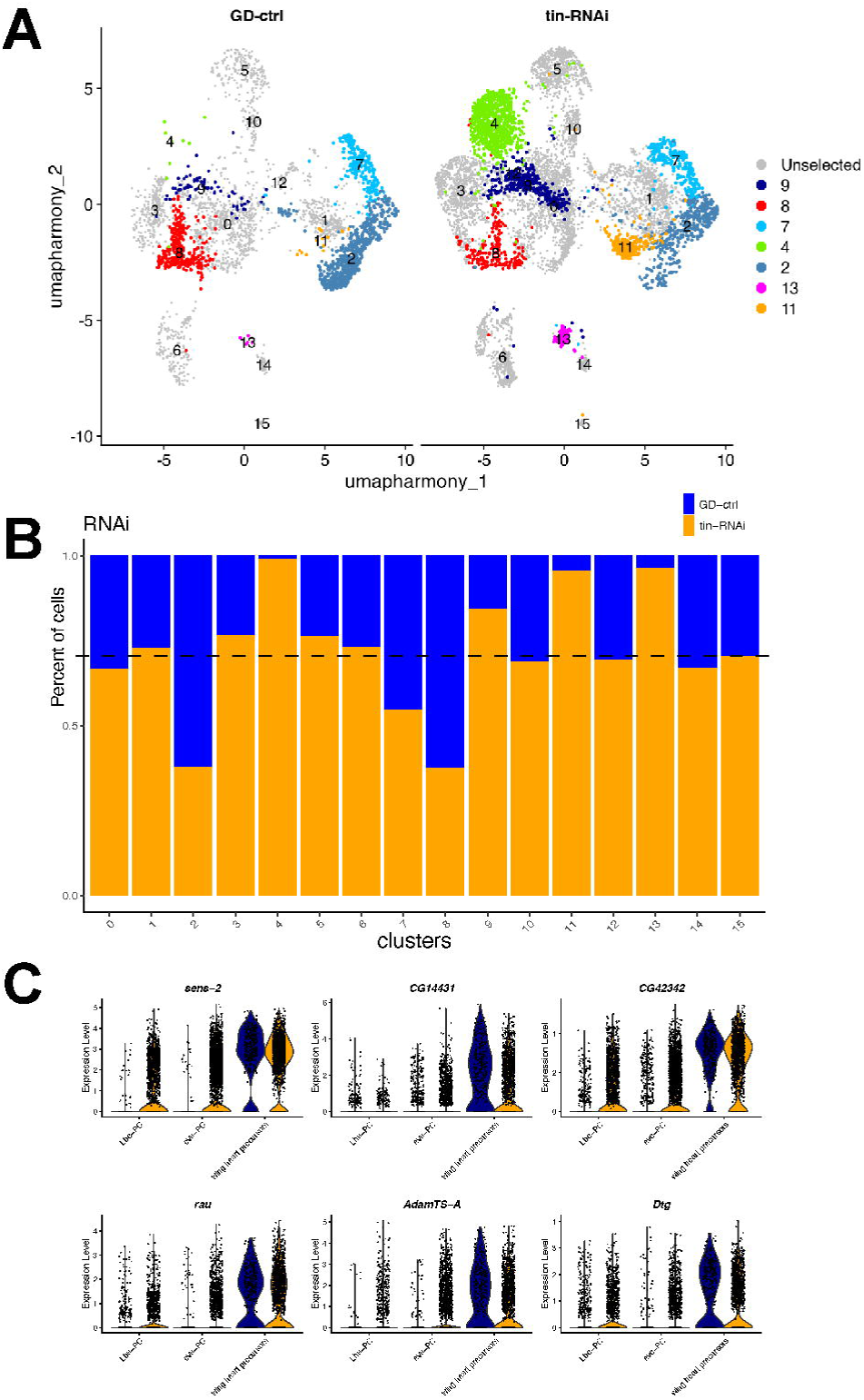

**Supplemental Figure 8.**
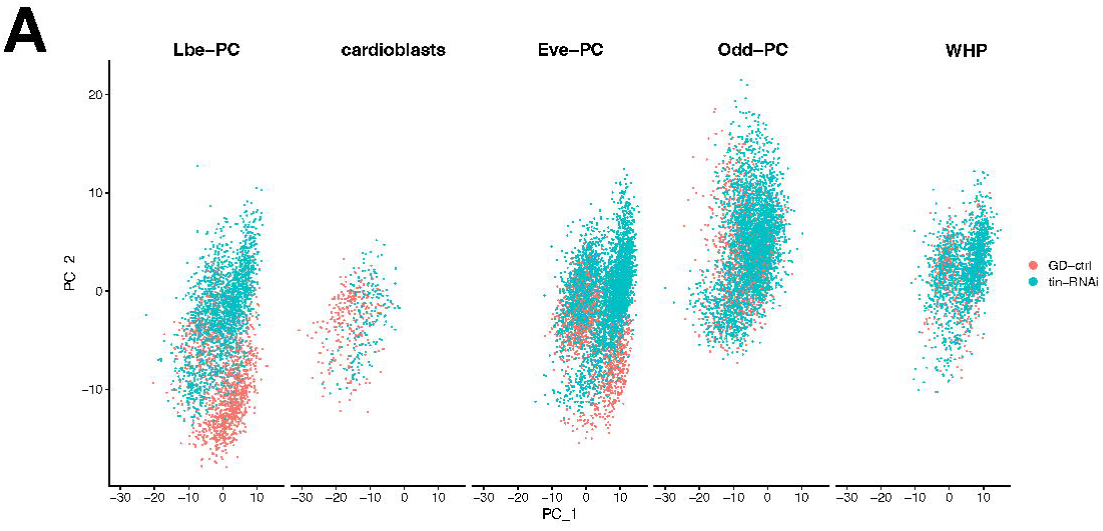

**Supplemental Figure 9.**
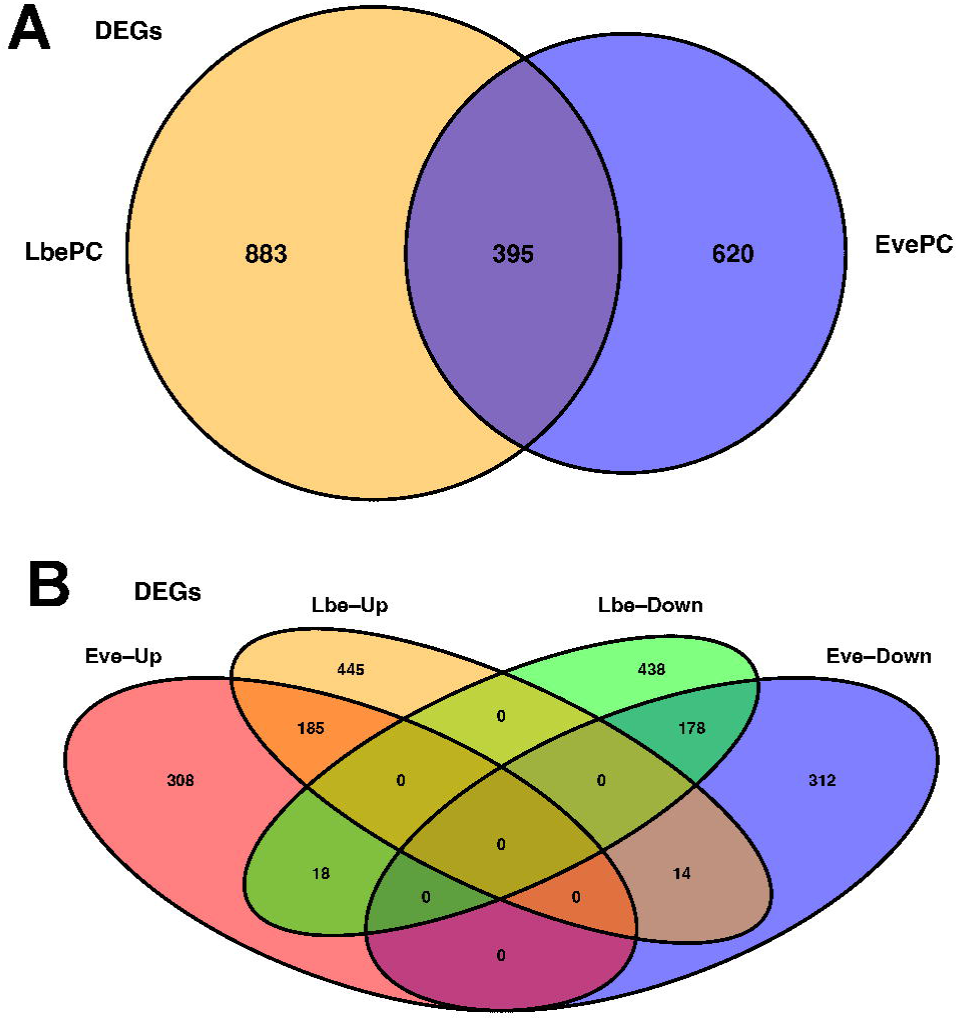

**Supplemental Figure 10.**
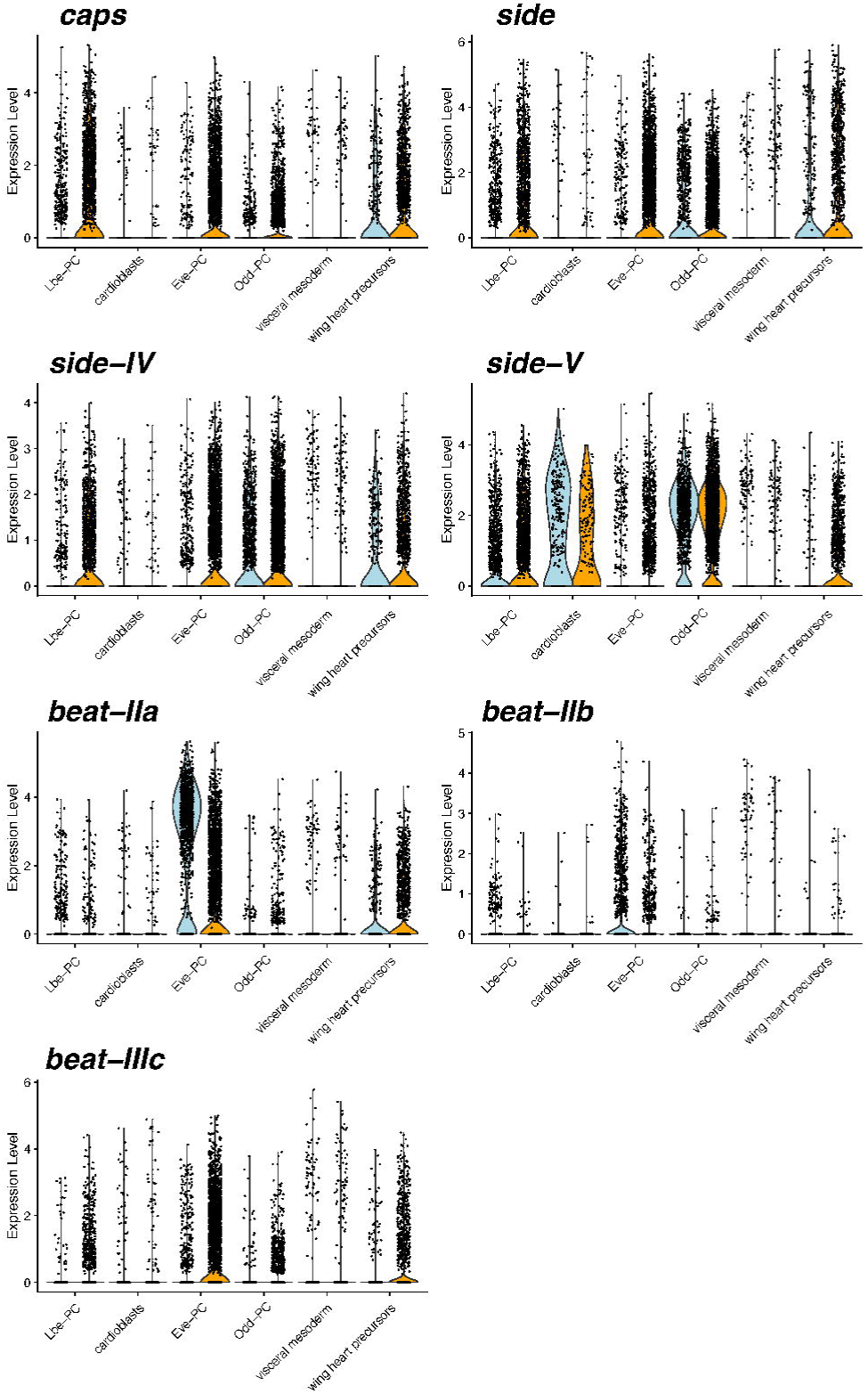

**Supplemental Figure 11.**
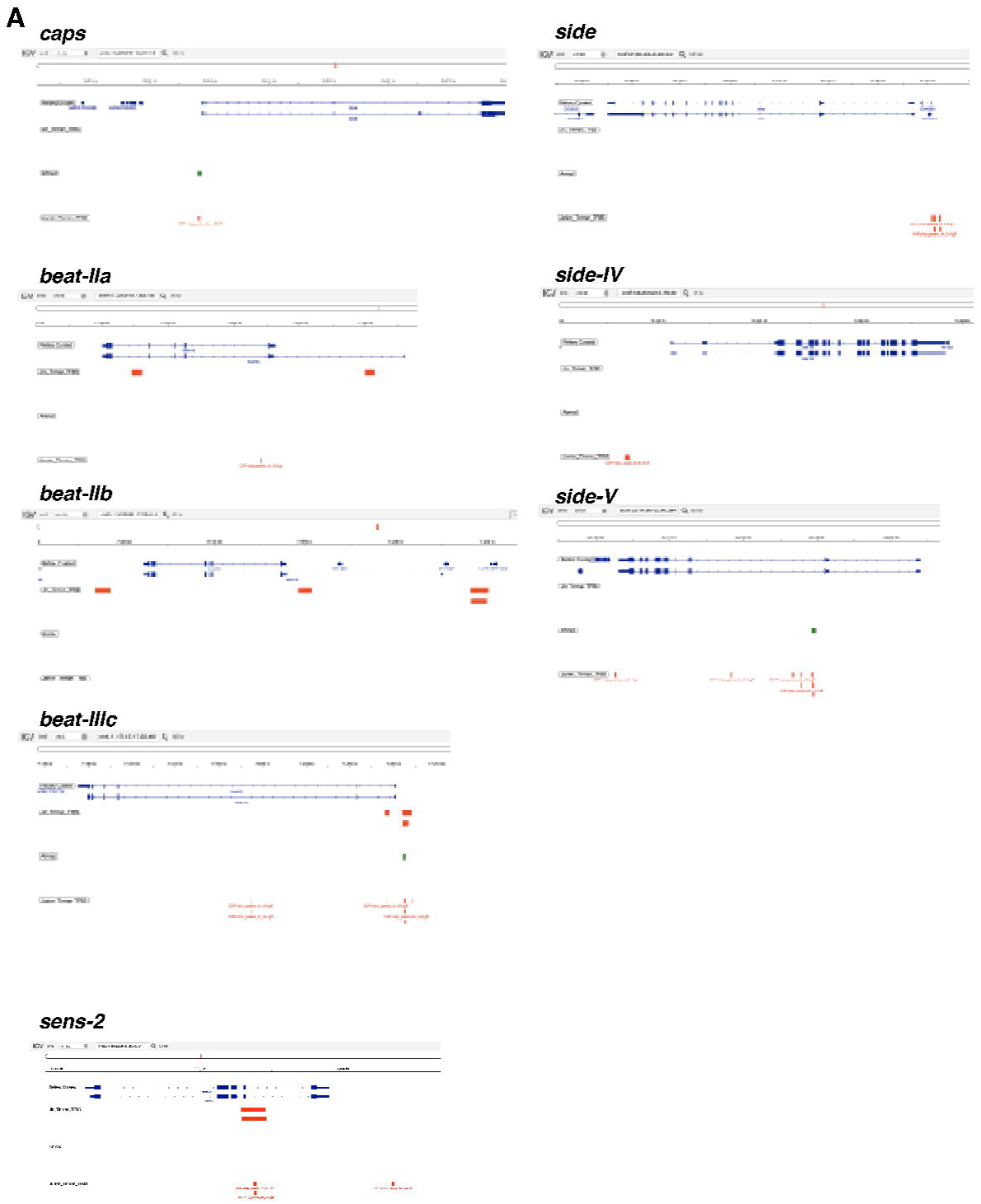

**Supplemental Figure 12.**
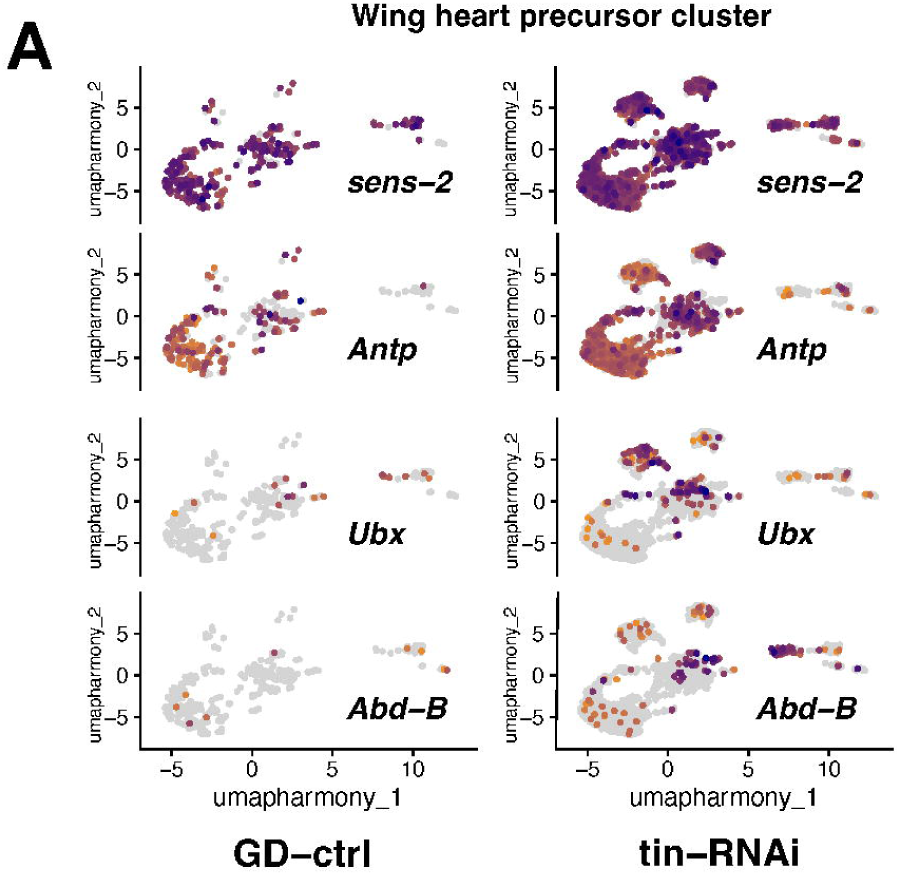

**Supplemental Figure 13.**
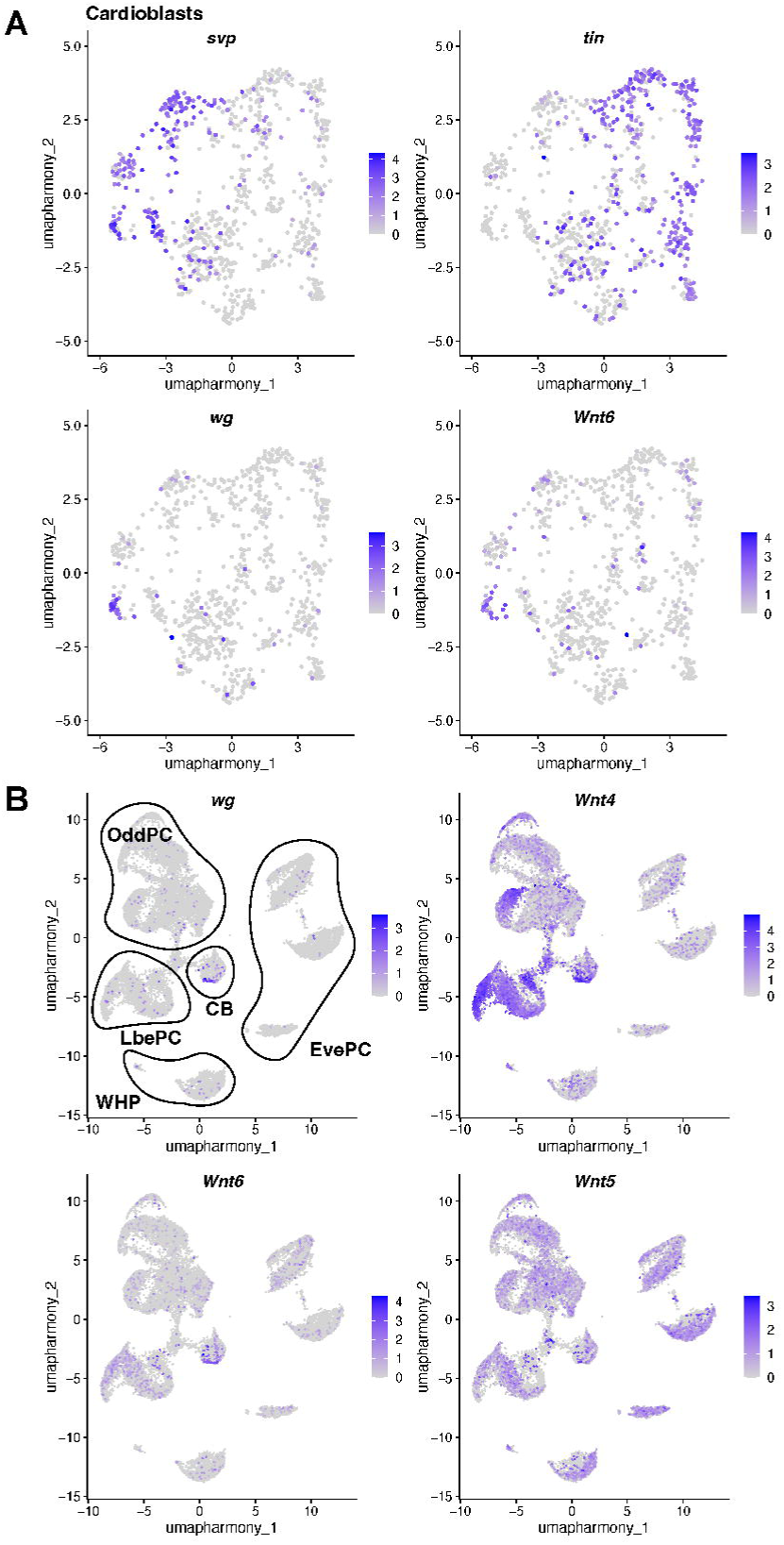

**Supplemental Figure 14.**
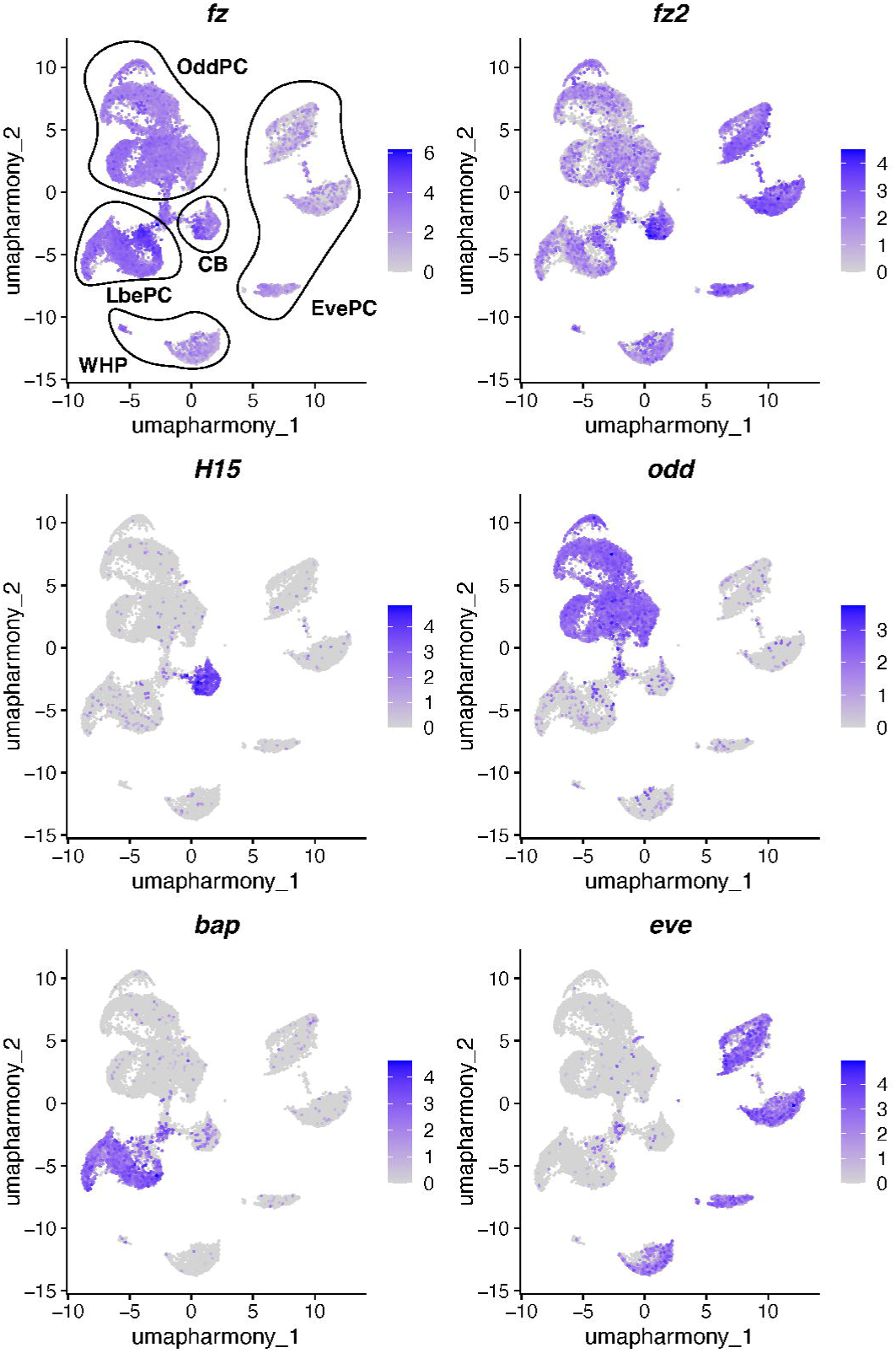

## Notes

### Competing Interest Statement

The authors have declared no competing interest.

### Summary of Updates

The updated version of this manuscript contains the addition of new cell types due to a different report construct, a more in-depth characterization of wild-type cells and we added functional data testing the consequences of tinman RNAi-mediated knockdown on gene expression.

### Funding

- National Heart Lung and Blood Institute

