## Supplemental Table 3 for "A complete single-cell atlas of the embryonic *Drosophila* heart and the cell-type specific role of Tinman"

### 1.1. Gene Ontology Enrichment Analysis: odd-PC

Significant GO terms (p < 0.05) with associated genes

| GO Term | Source | P-value | Genes |
| --- | --- | --- | --- |
| cytoplasmic translation | GO:BP | 2.65e-71 | *eIF3m,eIF3c,eIF3i,RpL24-like,eIF3e,eIF3g1,eIF3k,eIF3l,RpL22,RpL30,Rack1,RpL38,RpS5a,RpS14a,RpL40,RpL4,RpL23A,RpL11,RpL32,RpS20,RpS24,RpS2,RpS29,RpL35A,RpL34b,RpS10b,sta,RpLP1,RpS3,RpS28b,RpL27,RpL35,RpL12,RpS21,RpL39,RpS19a,RpL13,RpL10Ab,RpL9,RpL13A,RpL18A,RpS15Aa,RpL18,RpS4,RpS7,RpS27A,RpL24,RpL7,RpLP2,RpS27,RpL36A,RpS12,RpL23,RpL7A,RpS18,RpS13,RpS8,RpS9,RpS25,RpL26,RpL31,RpS26,RpS16,RpL28,RpL14,RpL36,RpL27A,RpL19,RpS15,RpS23,RpL41* |
| animal organ morphogenesis | GO:BP | 6.59e-07 | *odd,drm,mew,rst,tsh,tup,numb,pnut,Traf4,prc,cindr,Egfr,N,pnr,sn,fz,bab2,RhoGAP15B,Mkp3,Moe,Pvf2,noc,Cdep,jar,stg,drk,Dg,robo1,Dys,Rok,Septin1,lolal,gbb,Asap,kn,CG43658,shot,Vang,Myo31DF,Btk,daw,Rac2,Pal1,nmo,aux,CSN5,srp,sds22,Tak1* |
| formation of cytoplasmic translation initiation complex | GO:BP | 1.27e-05 | *eIF3m,eIF3c,eIF3i,eIF3e,eIF3g1,eIF3k,eIF3l* |
| dorsal closure | GO:BP | 2.50e-05 | *tup,Traf4,Egfr,N,Sec61alpha,Inx3,jar,Rok,gbb,shot,Btk,Rac2,srp,Tak1* |
| ribosomal small subunit assembly | GO:BP | 4.54e-04 | *RpS5a,RpS14a,sta,RpS28b,RpS19a,RpS27,RpS15* |
| germ cell migration | GO:BP | 7.09e-04 | *mim,wun2,Tre1,wun,zfh1,abd-A,srp* |
| embryonic heart tube development | GO:BP | 7.98e-04 | *Hand,tup,numb,pnr,robo1,srp* |
| motor neuron axon guidance | GO:BP | 9.58e-04 | *tup,N,zfh1,Ten-a,Ptp4E,robo1,Sulf1,cher,Rac2,side* |
| asymmetric protein localization involved in cell fate determination | GO:BP | 0.002 | *Galphai,Traf4,Klp98A,jar,egr* |
| imaginal disc-derived wing vein specification | GO:BP | 0.003 | *Egfr,N,Mkp3,Dg,Dys,gbb,kn,nmo* |
| ommatidial rotation | GO:BP | 0.003 | *Egfr,N,fz,Vang,nmo,Tak1* |
| mesodermal cell fate specification | GO:BP | 0.003 | *pyr,ths,abd-A* |
| muscle cell cellular homeostasis | GO:BP | 0.004 | *N,zfh1,Dg,robo1,Dys,Rack1* |
| cell fate commitment involved in pattern specification | GO:BP | 0.005 | *tup,numb,Egfr,N,fz,abd-A* |
| protein localization to endoplasmic reticulum | GO:BP | 0.006 | *CG5885,Sec61alpha,Spase22-23,Spase25,Sec61beta,CG8860* |
| glial cell migration | GO:BP | 0.006 | *pyr,numb,N,Pvf2,ths,CSN5* |
| head involution | GO:BP | 0.006 | *tsh,tup,Sec61alpha,Btk,chrb,Rac2* |
| cell surface receptor signaling pathway | GO:BP | 0.008 | *pyr,mew,wdp,apt,tsh,numb,Traf4,Egfr,lin-28,N,Sarm,fz,Mkp3,Pvf2,noc,E(spl)malpha-BFM,E(spl)mbeta-HLH,drk,Toll-7,Ptp4E,robo1,ths,Hsp83,gbb,Sulf1,Vang,Dcp-1,Lmpt,daw,Rac2,egr,nmo,aux,Tak1,Rack1,RpS12* |
| post-translational protein targeting to membrane, translocation | GO:BP | 0.009 | *Sec61alpha,Sec61beta,CG8860* |
| muscle cell fate specification | GO:BP | 0.009 | *numb,abd-A,kn* |
| germ cell repulsion | GO:BP | 0.01 | *wun2,wun* |
| myoblast migration | GO:BP | 0.01 | *pyr,ths* |
| tracheal outgrowth, open tracheal system | GO:BP | 0.012 | *Egfr,drk,lolal,Rac2* |
| SRP-dependent cotranslational protein targeting to membrane, translocation | GO:BP | 0.012 | *Sec61alpha,Sec61beta,CG8860* |
| cold acclimation | GO:BP | 0.012 | *Hsp83,Hsp23,Hsp26* |
| negative regulation of lamellocyte differentiation | GO:BP | 0.012 | *N,cher,CSN5* |
| chaperone cofactor-dependent protein refolding | GO:BP | 0.013 | *mrj,DnaJ-1,Torsin,Hsp70Ab,CG32641* |
| translational elongation | GO:BP | 0.013 | *eEF1beta,Rack1,eEF5,RpLP1,RpLP2* |
| germ-band shortening | GO:BP | 0.017 | *tup,Egfr,Rac2,srp* |
| signal peptide processing | GO:BP | 0.017 | *Spase22-23,Spase25,Spp* |
| 'de novo' protein folding | GO:BP | 0.02 | *mrj,DnaJ-1,Torsin,Hsp70Ab,CG32641* |
| midgut development | GO:BP | 0.022 | *mew,tsh,fz,abd-A,srp* |
| pericardial nephrocyte differentiation | GO:BP | 0.022 | *pyr,numb,pnr* |
| regulation of glycolytic process | GO:BP | 0.022 | *N,Dg* |
| larval fat body development | GO:BP | 0.022 | *gbb,Torsin* |
| regulation of asymmetric cell division | GO:BP | 0.022 | *numb,loco* |
| cortical microtubule organization | GO:BP | 0.022 | *Moe,shot* |
| axon ensheathment in central nervous system | GO:BP | 0.022 | *Galphai,loco* |
| gastrulation | GO:BP | 0.024 | *pyr,Traf4,T48,stg,Rok,ths,abd-A,srp* |
| salivary gland morphogenesis | GO:BP | 0.024 | *mew,fz,Pvf2,lolal,Btk,Rac2,srp* |
| regulation of myoblast fusion | GO:BP | 0.026 | *rst,Pax,Rok* |
| ectopic germ cell programmed cell death | GO:BP | 0.032 | *wun2,Tre1,wun* |
| negative regulation of cell population proliferation | GO:BP | 0.036 | *wdp,numb,Xrp1,Hsp83,CG6770,CSN5,sds22* |
| maintenance of epithelial integrity, open tracheal system | GO:BP | 0.038 | *mew,Egfr,Mkp3* |
| imaginal disc-derived leg joint morphogenesis | GO:BP | 0.038 | *odd,drm,N* |
| positive regulation of neuroblast proliferation | GO:BP | 0.04 | *N,E(spl)mbeta-HLH,Hsp83,daw* |
| positive regulation of wound healing | GO:BP | 0.045 | *Egfr,Rok,Rac2* |
| structural constituent of ribosome | GO:MF | 6.60e-54 | *RpL24-like,RpL22,RpL30,Rack1,RpL38,RpS5a,RpS14a,RpL40,RpL4,RpL23A,RpL11,RpL32,RpS20,RpS24,RpS2,RpS29,RpL35A,RpL34b,RpS10b,sta,RpLP1,RpS3,RpS28b,RpL27,RpL35,RpL12,RpS21,RpL39,RpS19a,RpL13,RpL10Ab,RpL9,RpL13A,RpL18A,RpS15Aa,RpL18,RpS4,RpS7,RpS27A,RpL24,RpL7,RpLP2,RpS27,RpL36A,RpS12,RpL23,RpL7A,RpS18,RpS13,RpS8,RpS9,RpS25,RpL26,RpL31,RpS26,RpS16,RpL28,RpL14,RpL36,RpL27A,RpL19,RpS15,RpS23,RpL41* |
| structural molecule activity | GO:MF | 7.40e-25 | *prc,Dg,RpL24-like,Dys,RpL22,shot,beta'COP,His4r,RpL30,Rack1,RpL38,RpS5a,RpS14a,RpL40,RpL4,RpL23A,RpL11,RpL32,RpS20,RpS24,RpS2,RpS29,RpL35A,RpL34b,RpS10b,sta,RpLP1,RpS3,RpS28b,RpL27,RpL35,RpL12,RpS21,RpL39,RpS19a,RpL13,RpL10Ab,RpL9,RpL13A,RpL18A,RpS15Aa,RpL18,RpS4,RpS7,RpS27A,RpL24,RpL7,RpLP2,RpS27,RpL36A,RpS12,RpL23,RpL7A,RpS18,RpS13,RpS8,RpS9,RpS25,RpL26,RpL31,RpS26,RpS16,RpL28,RpL14,RpL36,RpL27A,RpL19,RpS15,RpS23,RpL41* |
| RNA binding | GO:MF | 4.31e-12 | *apt,CG42458,lin-28,SC35,Non1,Rsf1,Pep,Pabp2,eIF3c,eIF3i,eIF2A,SmD2,eIF3g1,eIF3k,AGO2,RpL22,ytr,LSm7,Rpp25,su(f),trsn,Pop5,RpL30,CG10418,RpS5a,RpS14a,RpL4,RpL23A,RpL11,eEF5,ps,RpS20,RpS2,RpS10b,RpS3,RpL35,RpL12,RpS19a,RpL13,RpL10Ab,RpL9,RpL13A,RpL18,RpS4,RpL24,RpL7,RpS27,RpL23,RpL7A,RpS18,RpS13,RpS9,RpL26,RpS26,RpS16,RpL14,RpL19,RpS15* |
| protein binding | GO:MF | 2.96e-06 | *form3,side-V,pyr,mim,drm,mew,rst,Hand,wdp,apt,18w,tsh,tup,numb,Galphai,pnut,Traf4,CG7702,cindr,Egfr,N,Lrch,pnr,bif,Sarm,Septin2,sn,mrj,fz,pns,Fim,CG45263,bab2,olf186-M,Moe,Pvf2,mlt,Klp98A,side-IV,loco,IP3K2,CG12384,noc,CG10353,Hsp27,E(spl)malpha-BFM,cib,E(spl)mbeta-HLH,slbo,Cdep,jar,kud,drk,Pabp2,CG10011,eIF3m,Toll-7,eIF3c,Sh3beta,eIF3i,Ten-a,Dg,RhoGAP93B,CG40228,Ptp4E,Gclm,pix,eIF2A,CG1888,robo1,CG13551,Dys,Rok,Svil,Septin1,Idgf4,CG7029,ths,AGO2,eIF3l,lolal,Prosbeta4,udd,Hsp83,gbb,abd-A,Asap,shot,CG42709,Vang,Dcp-1,HP4,CG8209,Myo31DF,CG7484,Btk,CG14696,Taf13,MED22,MED11,Roc2,Nop17l,CG2147,beta'COP,DnaJ-1,Polr2J,dpr17,cher,daw,LSm7,His4r,Rac2,HP1b,CG1129,egr,Pal1,nmo,su(f),aux,CSN5,Tctp,trsn,dgt2,srp,sds22,side,Rtnl1,Tak1,Hsp23,Rack1,Hsp70Ab,CG5961,CG10418,e(y)2,CG32641,Hsp26,RpL40,RpL11,RpS10b,RpLP1,RpS27A,RpL23,RpS26,Pde1c* |
| rRNA binding | GO:MF | 5.37e-06 | *RpS5a,RpS14a,RpL23A,RpL11,RpL12,RpL9,RpS4,RpL23,RpS18,RpS13,RpS9* |
| nucleic acid binding | GO:MF | 7.34e-05 | *odd,drm,Hand,apt,CG42458,tsh,tup,Galphai,lin-28,pnr,bab2,noc,Xrp1,E(spl)mbeta-HLH,SC35,slbo,Non1,Rsf1,zfh1,Pep,Pabp2,eIF3m,eIF3c,eIF3i,eIF2A,eIF3e,SmD2,eIF3g1,eIF3k,AGO2,eIF3l,lolal,CG3224,abd-A,kn,RpL22,Polr2K,ytr,eEF1beta,Polr2J,LSm7,His4r,Rpp25,su(f),trsn,srp,Pop5,RpL30,CG10418,e(y)2,RpS5a,RpS14a,RpL4,RpL23A,RpL11,eEF5,ps,RpS20,RpS2,RpS10b,RpS3,RpL35,RpL12,RpS19a,RpL13,RpL10Ab,RpL9,RpL13A,RpL18,RpS4,RpL24,RpL7,RpS27,RpL23,RpL7A,RpS18,RpS13,RpS9,RpL26,RpS26,RpS16,RpL14,RpL19,RpS15* |
| protein domain specific binding | GO:MF | 0.001 | *mim,rst,cindr,N,CG12384,Sh3beta,Dys,lolal,Hsp83,Dcp-1,e(y)2* |
| translation factor activity, RNA binding | GO:MF | 0.002 | *eIF3m,eIF3c,eIF3i,eIF2A,eIF3e,eIF3g1,eIF3k,eIF3l,eEF1beta,eEF5* |
| ribosome binding | GO:MF | 0.002 | *Sec61alpha,pix,eIF2A,eIF3k,Rack1,eEF5,sta,RpS21* |
| actin binding | GO:MF | 0.002 | *form3,mim,pnut,bif,sn,Fim,Moe,cib,jar,Dys,Svil,shot,Myo31DF,cher* |
| translation initiation factor activity | GO:MF | 0.002 | *eIF3m,eIF3c,eIF3i,eIF2A,eIF3e,eIF3g1,eIF3k,eIF3l* |
| translation regulator activity | GO:MF | 0.004 | *eIF3m,eIF3c,eIF3i,eIF2A,eIF3e,eIF3g1,eIF3k,eIF3l,eEF1beta,Rack1,eEF5* |
| ribonucleoprotein complex binding | GO:MF | 0.006 | *Sec61alpha,pix,eIF2A,eIF3k,Rack1,eEF5,sta,RpLP1,RpS21* |
| growth factor receptor binding | GO:MF | 0.01 | *pyr,Pvf2,drk,Idgf4,ths* |
| growth factor activity | GO:MF | 0.01 | *pyr,Pvf2,ths,gbb,daw* |
| ubiquitin protein ligase binding | GO:MF | 0.018 | *pnut,Traf4,E(spl)malpha-BFM,Septin1,CG14696,Tak1,RpL40,RpS27A* |
| ubiquitin-like protein ligase binding | GO:MF | 0.02 | *pnut,Traf4,E(spl)malpha-BFM,Septin1,CG14696,Tak1,RpL40,RpS27A* |
| protein-containing complex binding | GO:MF | 0.03 | *mew,Galphai,bif,sn,Fim,Sec61alpha,jar,pix,eIF2A,Dys,eIF3k,Svil,Myo31DF,cher,aux,Rack1,e(y)2,eEF5,sta,RpLP1,RpS21* |
| mRNA binding | GO:MF | 0.037 | *apt,lin-28,SC35,Pabp2,eIF2A,eIF3g1,ytr,su(f),RpS5a,RpS14a,ps,RpL35,RpL13A,RpL24,RpS26* |
| cytoskeletal protein binding | GO:MF | 0.038 | *form3,mim,pnut,bif,sn,Fim,Moe,mlt,Klp98A,cib,Cdep,jar,Dys,Svil,shot,Myo31DF,cher,dgt2,Hsp26,RpL23* |
| inositol-1,4,5-trisphosphate 3-kinase activity | GO:MF | 0.044 | *IP3K1,IP3K2* |
| inositol tetrakisphosphate kinase activity | GO:MF | 0.044 | *IP3K1,IP3K2* |
| actin filament binding | GO:MF | 0.046 | *bif,sn,Fim,jar,Dys,Svil,Myo31DF,cher* |

### 1.2. Gene Ontology Enrichment Analysis: germline

Significant GO terms (p < 0.05) with associated genes

| GO Term | Source | P-value | Genes |
| --- | --- | --- | --- |
| regulation of metabolic process | GO:BP | 2.61e-27 | *CycB,bru1,wisp,smg,CycA,BigH1,retn,Marf1,me31B,gnu,zld,Nup358,Top2,PCNA,pgc,tral,CycE,ovo,CG46385,Incenp,lok,CG9705,CycB3,Su(var)2-10,Uba2,msl-1,Lsd-2,srl,rin,sqd,CNBP,e(y)3,brat,geminin,Mtor,G9a,ago,kis,Su(var)2-HP2,woc,Usp7,Dek,Tl,Gbs-70E,Usp14,Hrb87F,east,CtBP,Hsp83,Fkbp39,Dp,gro,wech,Hsc70-3,E(bx),dco,Rpn2,Unr,Aef1,MED26,Jarid2,mnb,rdx,CG2926,Rox8,chn,Gyf,Pdcd4,hang,Sumo,Desat1,Ssdp,Paip2,Hsc70-4,jigr1,Kdm5,Mitf,PAN3,Cpsf6,Tudor-SN,Myc,eIF4G1,mts,cnc,dom,UbcE2H,osa,Lk6,gus,Nipped-B,stau,TER94,spoon,Ndf,Brd7-9,SRPK,Cul1,hoip,psq,TfIIFalpha,Usp10,Pp2C1,Ars2,Sod1,kra,Non2,CkIIalpha,Nup153,gw,upSET,gcl,sov,mask,Sin3A,CG8677,sesB,nej,heph,Ufd4,Not1,CG1677,Hrb98DE,Hel25E,l(3)80Fj,Ntf-2,nsl1,Nedd4,bin3,BtbVII,Nipped-A,E(Pc),Helz,mei-P26,Rpt2,Mi-2,PlexA,Rpt6,CG32767,wcy,dap,Gug,CkIalpha,skd,Pits,CycT,d4,cact,da,tou,eIF3b,cg,Mnt,sina* |
| nucleobase-containing compound metabolic process | GO:BP | 1.31e-18 | *bru1,wisp,smg,exu,BigH1,retn,Marf1,me31B,gnu,zld,Nup358,fs(1)Ya,Top2,PCNA,pgc,CycE,ovo,PyK,CG46385,RnrS,dUTPase,Gapdh2,Caf1-180,CG10254,Nop60B,RnrL,lok,CG9705,mt:ATPase6,Su(var)2-10,Uba2,Snp,awd,srl,sqd,e(y)3,brat,geminin,Mtor,G9a,ago,kis,woc,Dek,lost,Tl,nop5,Hrb87F,east,CtBP,Dp,ncm,His2Av,gro,E(bx),Aef1,Nap1,MED26,Jarid2,mnb,CG2926,Rox8,chn,Pdcd4,hang,SmD3,Sumo,Top1,Ssdp,Hsc70-4,jigr1,Idh,Kdm5,LysRS,Mitf,PAN3,Cpsf6,Tudor-SN,Myc,cnc,dom,Vha55,Mdh1,osa,Gapdh1,Nipped-B,SNRPG,Vha68-2,spoon,Ndf,Brd7-9,SRPK,Uev1A,hoip,psq,Aps,TfIIFalpha,Sem1,Hex-A,Usp10,Polr2A,mod,Ars2,Non2,Nup153,gw,upSET,gcl,sov,mask,Sin3A,CG8677,sesB,nej,heph,Not1,CG1677,Ahcy,Hrb98DE,Hel25E,nsl1,blw,bin3,e(r),BtbVII,Nipped-A,E(Pc),Ref1,Pde11,mei-P26,Mi-2,Nop56,PlexA,CG32767,dap,Gug,CkIalpha,skd,Pits,ben,CycT,d4,cact,da,tou,cg,Pglym78,Mnt* |
| negative regulation of biosynthetic process | GO:BP | 1.90e-14 | *bru1,smg,BigH1,retn,Marf1,me31B,zld,Nup358,Top2,pgc,tral,ovo,CG46385,CG9705,sqd,e(y)3,brat,Mtor,G9a,kis,Su(var)2-HP2,Usp7,east,CtBP,Hsp83,gro,wech,Hsc70-3,E(bx),Unr,Rox8,chn,Gyf,Pdcd4,Paip2,Hsc70-4,PAN3,Tudor-SN,dom,Cul1,psq,Usp10,Pp2C1,Ars2,kra,gw,upSET,gcl,sov,Sin3A,CG8677,heph,Not1,Hrb98DE,bin3,Helz,Mi-2,CG32767,Gug,Pits,d4,cact,Mnt* |
| regulation of RNA metabolic process | GO:BP | 1.00e-12 | *bru1,smg,BigH1,retn,Marf1,me31B,zld,pgc,ovo,CG46385,lok,CG9705,Su(var)2-10,Uba2,srl,sqd,e(y)3,Mtor,G9a,kis,woc,Tl,Hrb87F,CtBP,Dp,gro,E(bx),Aef1,MED26,Jarid2,mnb,CG2926,Rox8,chn,Pdcd4,hang,Sumo,Ssdp,jigr1,Kdm5,Mitf,PAN3,Cpsf6,Myc,cnc,dom,osa,Nipped-B,Ndf,Brd7-9,SRPK,psq,TfIIFalpha,Usp10,Non2,Nup153,gw,upSET,gcl,sov,mask,Sin3A,CG8677,nej,Not1,CG1677,Hrb98DE,Hel25E,nsl1,BtbVII,Nipped-A,E(Pc),mei-P26,Mi-2,CG32767,Gug,skd,Pits,CycT,d4,cact,da,tou,cg,Mnt* |
| catabolic process | GO:BP | 1.14e-11 | *smg,Jafrac1,26-29-p,Marf1,Pxt,me31B,zld,Ggt-1,PyK,CG46385,dUTPase,Gapdh2,CG9705,CG32473,CG7840,Uba1,Lsd-2,sqd,ago,Usp7,Usp14,Fkbp39,dco,Rpn2,rdx,Prosalpha4,Gyf,Ubqn,Rpn5,Echs1,Desat1,Hsc70-4,Mitf,PAN3,Cpsf6,Prosalpha5,Tudor-SN,Myc,mts,UbcE2H,Gapdh1,gus,Ubc4,TER94,Prosbeta5,ctp,Cul1,Prosalpha6,Aps,Prosalpha7,Sem1,Hex-A,Usp10,Sod1,CkIIalpha,gw,gcl,Prosbeta6,mask,sesB,Ufd4,Not1,CG1677,Rpn11,Ahcy,Nedd4,Prosalpha3,Prosbeta2,stx,Rpt2,Rpt6,wcy,CkIalpha,Prosbeta4,Prosbeta1,Pglym78,sina* |
| positive regulation of smoothened signaling pathway | GO:BP | 2.04e-09 | *Su(var)2-10,Tnpo,tws,Usp7,dco,Sumo,mts,cno,CkIIalpha,nej,apolpp,gish,CkIalpha* |
| NLS-bearing protein import into nucleus | GO:BP | 9.90e-08 | *Pen,Nup358,Rcc1,Tnpo,Fs(2)Ket,Nup153,Kap-alpha3* |
| negative regulation of signal transduction | GO:BP | 1.93e-05 | *Cam,Su(var)2-10,Uba2,srl,Tao,Usp7,Tl,CtBP,His2Av,gro,E(bx),dco,mnb,rdx,hang,Ptp61F,Myc,mts,TER94,Cul1,wap,gcl,Sin3A,sesB,nej,Ufd4,Nedd4,Gug,CkIalpha,cact* |
| cell surface receptor signaling pathway | GO:BP | 4.63e-05 | *CycE,Su(var)2-10,Uba2,Tnpo,tws,srl,Tao,Usp7,Tl,CtBP,Hsp83,gro,E(bx),dco,for,rdx,CG45050,Sumo,Ssdp,Ptp61F,Myc,mts,osa,TER94,Cul1,cno,Uev1A,Usp10,wap,CkIIalpha,gcl,mask,Sin3A,nej,apolpp,Nedd4,gish,PlexA,Gug,CkIalpha,skd,ben,cact,sina* |
| peptidyl-serine phosphorylation | GO:BP | 5.90e-05 | *awd,Tlk,dco,mbt,mnb,Lk6,SRPK,CkIIalpha,gish,CkIalpha* |
| positive regulation of transcription by RNA polymerase II | GO:BP | 8.03e-05 | *zld,ovo,lok,srl,e(y)3,Mtor,Tl,CtBP,E(bx),chn,Ssdp,Kdm5,Myc,cnc,Ndf,TfIIFalpha,Nup153,sov,mask,CG8677,nej,nsl1,CG32767,skd,CycT,d4,da,tou* |
| animal organ morphogenesis | GO:BP | 1.03e-04 | *CycE,msk,ovo,Cam,rin,CNBP,Tao,kis,Hrb87F,CtBP,dco,mbt,rdx,chn,spdi,Mitf,mts,osa,baz,gus,Nipped-B,ctp,cno,hoip,psq,Moe,mask,nej,heph,Hrb98DE,Nedd4,gish,dap,wge,Gug,skd,dsd,p120ctn,da,cg,sdk,tth,sina* |
| positive regulation of canonical Wnt signaling pathway | GO:BP | 1.71e-04 | *tws,CtBP,dco,Ssdp,mts,nej,apolpp,gish,skd* |
| RNA splicing, via transesterification reactions | GO:BP | 1.81e-04 | *bru1,exu,sqd,lost,Hrb87F,ncm,CG2926,Rox8,SmD3,Hsc70-4,Cpsf6,dom,SNRPG,SRPK,hoip,mod,Ars2,CG1677,Hrb98DE,Hel25E,Ref1* |
| protein polyubiquitination | GO:BP | 3.00e-04 | *CG10254,Uba1,CG2924,rdx,UbcE2H,Ubc4,Cul1,Uev1A,gcl,Ufd4,Nedd4,ben* |
| ventral cord development | GO:BP | 3.26e-04 | *Nup358,CycE,Rcc1,glu,brat,ago,Surf6,Galphai,dap,CycT* |
| negative regulation of cell population proliferation | GO:BP | 4.95e-04 | *polo,tws,brat,ago,Hsp83,Rox8,Ptp61F,mts,dom,osa* |
| negative regulation of smoothened signaling pathway | GO:BP | 6.70e-04 | *dco,rdx,mts,TER94,Cul1,Nedd4,Gug,CkIalpha* |
| regulation of cyclin-dependent protein serine/threonine kinase activity | GO:BP | 7.72e-04 | *CycB,CycA,CycE,CycB3,dap* |
| post-translational protein modification | GO:BP | 8.12e-04 | *CycE,CG10254,Su(var)2-10,Uba2,Uba1,ago,Usp7,Usp14,Tlk,CG2924,rdx,Sumo,UbcE2H,Ubc4,Cul1,Uev1A,Usp10,gcl,Ufd4,Rpn11,Nedd4,mei-P26,ben,sina* |
| meiotic cell cycle | GO:BP | 8.23e-04 | *dhd,wisp,mtrm,Top2,polo,Incenp,lok,Ran,east,Kdm5,Nipped-B,SRPK,endos,baf,Cul1,enc,mei-P26,dap,fwd* |
| regulation of chromatin binding | GO:BP | 8.25e-04 | *CycB,CycA,CycB3* |
| nuclear pore complex assembly | GO:BP | 8.25e-04 | *Nup358,Ran,emb* |
| pronuclear migration | GO:BP | 8.25e-04 | *wisp,alphaTub67C,polo* |
| sperm aster formation | GO:BP | 8.25e-04 | *wisp,alphaTub67C,polo* |
| sperm DNA decondensation | GO:BP | 1.00e-03 | *dhd,Nlp,Nap1,Nph* |
| larval somatic muscle development | GO:BP | 0.001 | *polo,Tl,chn,Kdm5,Sin3A,nsl1,Gug,skd* |
| habituation | GO:BP | 0.001 | *me31B,G9a,for,wcy* |
| negative regulation of neuroblast proliferation | GO:BP | 0.002 | *polo,tws,brat,mts,osa* |
| negative regulation of canonical Wnt signaling pathway | GO:BP | 0.002 | *CtBP,gro,TER94,Cul1,Sin3A,nej,CkIalpha* |
| positive regulation of stem cell differentiation | GO:BP | 0.002 | *bru1,CycA,Pdcd4* |
| pronuclear fusion | GO:BP | 0.002 | *wisp,fs(1)Ya,polo* |
| positive regulation of Toll signaling pathway | GO:BP | 0.002 | *Uba2,tws,for,Sumo,mts* |
| ribose phosphate metabolic process | GO:BP | 0.002 | *PyK,Gapdh2,mt:ATPase6,awd,Vha55,Gapdh1,Vha68-2,Aps,Hex-A,sesB,blw,Pde11,Pglym78* |
| protein K63-linked ubiquitination | GO:BP | 0.002 | *CG10254,Uev1A,Ufd4,ben* |
| positive regulation of organ growth | GO:BP | 0.003 | *kis,mnb,Myc,wap* |
| positive regulation of proteasomal ubiquitin-dependent protein catabolic process | GO:BP | 0.003 | *dco,UbcE2H,TER94,Cul1,Ufd4,CkIalpha,sina* |
| regulation of chromosome organization | GO:BP | 0.004 | *pgc,tws,Mtor,Usp7,Cul1,Moe,Non2,Mi-2* |
| regulation of pole plasm oskar mRNA localization | GO:BP | 0.004 | *exu,cnc,stau,TER94,enc* |
| regulation of alternative mRNA splicing, via spliceosome | GO:BP | 0.004 | *bru1,sqd,Hrb87F,CG2926,Rox8,Cpsf6,dom,Hrb98DE,Hel25E* |
| deoxyribonucleotide biosynthetic process | GO:BP | 0.005 | *RnrS,dUTPase,RnrL* |
| miRNA-mediated gene silencing by inhibition of translation | GO:BP | 0.005 | *me31B,gw* |
| regulation of metaphase plate congression | GO:BP | 0.005 | *Mtor,east* |
| determination of adult lifespan | GO:BP | 0.006 | *Jafrac1,Hsp27,Trxr-1,mt:ATPase6,kis,Prosalpha5,cnc,Prosalpha6,Sod1,Sin3A,sesB,Sam-S,fwd* |
| negative regulation of insulin receptor signaling pathway | GO:BP | 0.006 | *srl,Tl,Ptp61F,mts,Cul1* |
| stress granule assembly | GO:BP | 0.007 | *me31B,tral,rin* |
| oocyte karyosome formation | GO:BP | 0.007 | *lok,Kdm5,SRPK,baf,enc* |
| negative regulation of hippo signaling | GO:BP | 0.008 | *Usp7,mnb,mts,Cul1,wap,CkIalpha* |
| glycolytic process | GO:BP | 0.008 | *PyK,Gapdh2,Gapdh1,Hex-A,Pglym78* |
| protein refolding | GO:BP | 0.009 | *Hsp27,Droj2,Hsc70-3,Hsc70-4,Hsp60A* |
| nucleoside diphosphate catabolic process | GO:BP | 0.009 | *PyK,Gapdh2,Gapdh1,Hex-A,Pglym78* |
| negative regulation of organ growth | GO:BP | 0.009 | *Tao,Hsc70-3,Rox8,Ptp61F* |
| regulation of establishment of planar polarity | GO:BP | 0.009 | *dco,baz,cno,gish* |
| negative regulation of receptor signaling pathway via JAK-STAT | GO:BP | 0.009 | *Su(var)2-10,Uba2,E(bx),Ptp61F* |
| somatic stem cell population maintenance | GO:BP | 0.011 | *CycE,His2Av,dom,da* |
| muscle cell cellular homeostasis | GO:BP | 0.011 | *Cam,mt:ATPase6,Gyf,TER94,sesB* |
| positive regulation of nuclear-transcribed mRNA poly(A) tail shortening | GO:BP | 0.012 | *smg,Marf1,gw* |
| regulation of autophagy of mitochondrion | GO:BP | 0.012 | *ago,Sod1,mask* |
| positive regulation of cytoplasmic mRNA processing body assembly | GO:BP | 0.012 | *PAN3,stau* |
| male germline ring canal formation | GO:BP | 0.012 | *polo,fwd* |
| meiotic spindle midzone assembly | GO:BP | 0.012 | *polo,Incenp* |
| 3'-UTR-mediated mRNA destabilization | GO:BP | 0.012 | *Marf1,gw* |
| eggshell chorion gene amplification | GO:BP | 0.013 | *PCNA,CycE,geminin,Dp* |
| negative regulation of phosphatidylinositol 3-kinase/protein kinase B signal transduction | GO:BP | 0.015 | *Ptp61F,mts,Cul1* |
| positive regulation of hippo signaling | GO:BP | 0.015 | *Tao,Hsc70-3,dco,Rox8,Ptp61F* |
| protein autophosphorylation | GO:BP | 0.019 | *awd,Tlk,dco,mnb,Lk6* |
| peripheral nervous system development | GO:BP | 0.022 | *alphaTub67C,CycE,chn,SmD3,egh,GlcAT-P* |
| mRNA export from nucleus | GO:BP | 0.023 | *sqd,Mtor,Sem1,Hel25E,Ref1* |
| bicoid mRNA localization | GO:BP | 0.024 | *exu,cnc,stau* |
| antimicrobial humoral immune response mediated by antimicrobial peptide | GO:BP | 0.024 | *wisp,Uba2,Tl,Sumo,Cul1,Ntf-2,cact* |
| positive regulation of receptor signaling pathway via JAK-STAT | GO:BP | 0.029 | *CycE,CG45050,mask* |
| neuroblast fate determination | GO:BP | 0.029 | *CycE,brat,stau,da* |
| germarium-derived oocyte fate determination | GO:BP | 0.029 | *baz,BicD,enc,dap* |
| nucleoside triphosphate biosynthetic process | GO:BP | 0.03 | *mt:ATPase6,awd,Vha68-2,sesB,blw* |
| short-term memory | GO:BP | 0.032 | *G9a,kis,for,Nipped-B* |
| positive regulation of mRNA splicing, via spliceosome | GO:BP | 0.033 | *Hrb87F,Hrb98DE* |
| positive regulation of cytoplasmic translation | GO:BP | 0.033 | *CNBP,spoon* |
| juvenile hormone mediated signaling pathway | GO:BP | 0.033 | *Nup358,Fkbp39* |
| neuron cellular homeostasis | GO:BP | 0.04 | *mt:ATPase6,srl,Gyf,sesB* |
| imaginal disc-derived wing vein morphogenesis | GO:BP | 0.042 | *msk,osa,heph,Nedd4,Gug* |
| regulation of syncytial blastoderm mitotic cell cycle | GO:BP | 0.045 | *grp,geminin* |
| follicle cell of egg chamber-cell adhesion | GO:BP | 0.045 | *egh,baz* |
| dendritic spine morphogenesis | GO:BP | 0.045 | *Sumo,p120ctn* |
| negative regulation of translational initiation | GO:BP | 0.045 | *Unr,Paip2* |
| positive regulation of tumor necrosis factor-mediated signaling pathway | GO:BP | 0.045 | *Uev1A,ben* |
| male germ-line stem cell population maintenance | GO:BP | 0.045 | *His2Av,Kdm5,dom* |
| protein sumoylation | GO:BP | 0.045 | *Su(var)2-10,Uba2,Sumo* |
| protein binding | GO:MF | 1.92e-31 | *CycB,bru1,wisp,Pen,smg,Jafrac1,yl,CycA,exu,mtrm,retn,me31B,gnu,zld,Nup358,fs(1)Ya,Top2,pgc,tral,feo,CycE,msk,ovo,Hsp27,CG8223,Drep2,dUTPase,CG42232,Caf1-180,Cam,polo,Incenp,lok,CycB3,Trxr-1,glu,Df31,cue,Su(var)2-10,Uba2,msl-1,Tnpo,Mapmodulin,awd,tws,Lsd-2,Ran,srl,emb,rin,e(y)3,brat,stai,geminin,Tao,Mtor,G9a,ago,kis,LpR2,woc,Usp7,Dek,Tl,Droj2,ced-6,Gbs-70E,Tlk,Nlp,CtBP,Hsp83,Fkbp39,Dp,ncm,His2Av,gro,wech,Hsc70-3,E(bx),dco,Unr,Nap1,mbt,CCT2,mnb,rdx,chn,CG45050,Gyf,Sumo,Ubqn,Desat1,Ssdp,Paip2,Hsc70-4,Ptp61F,jigr1,Kdm5,spdi,Mitf,PAN3,Tudor-SN,Myc,eIF4G1,mts,cnc,dom,osa,Lk6,baz,gus,Nipped-B,Fs(2)Ket,Arpc2,stau,Ubc4,TER94,spoon,Brd7-9,Prosbeta5,Spn,Nph,ctp,Vap33,Hsp60A,Cul1,cno,Uev1A,psq,TfIIFalpha,Sem1,CCT5,Hex-A,Dhit,Pp2C1,toc,BicD,mod,Dlic,Ars2,Sod1,kra,Moe,wap,CCT3,CkIIalpha,gw,upSET,gcl,mask,Sin3A,CG8677,bif,Kap-alpha3,Phb2,nej,Hsc70Cb,apolpp,CanA-14F,Ufd4,Not1,CCT4,Rpn11,CCT1,l(3)80Fj,Ntf-2,nsl1,Nedd4,Prosalpha3,SKIP,bin3,BtbVII,Nipped-A,stx,Pde11,unc-13,Helz,mei-P26,CCT6,Mi-2,Galphai,milt,PlexA,Rpt6,CG32767,wcy,dap,CG11267,Gug,CkIalpha,Prosbeta4,skd,dsd,ben,Prosbeta1,CG13917,p120ctn,CycT,d4,cact,da,tou,eIF3b,sdk,Mnt,sina* |
| nucleic acid binding | GO:MF | 2.42e-09 | *bru1,wisp,smg,exu,mtrm,BigH1,retn,Marf1,me31B,zld,Top2,PCNA,tral,msk,ovo,Nop60B,CG9705,Snp,msl-1,srl,rin,sqd,CNBP,brat,eEF2,Mtor,kis,Su(var)2-HP2,Capr,Dek,nop5,Hrb87F,Nlp,Fkbp39,Dp,ncm,His2Av,E(bx),Unr,Aef1,D1,Jarid2,Rox8,chn,CG45050,Gyf,hang,SmD3,Top1,Ssdp,Paip2,jigr1,Kdm5,LysRS,Mitf,PAN3,Cpsf6,Tudor-SN,Myc,eIF4G1,cnc,dom,osa,stau,RpS5b,SNRPG,spoon,Surf6,Ndf,Nph,baf,hoip,psq,TfIIFalpha,CG14478,Polr2A,mod,Nup153,gw,mask,nej,heph,Hrb98DE,Hel25E,bin3,vig2,enc,BtbVII,Ref1,Helz,Mi-2,Galphai,Nop56,CG32767,da,tou,eIF3b,cg,eIF4H1,Mnt* |
| purine nucleotide binding | GO:MF | 3.45e-09 | *wisp,alphaTub67C,me31B,Top2,PyK,Gapdh2,polo,betaTub56D,RnrL,lok,rdog,grp,glu,Uba2,Uba1,awd,Ran,eEF2,Tao,kis,Droj2,Tlk,CtBP,Hsp83,Hsc70-3,dco,mbt,for,CCT2,mnb,Hsc70-4,Idh,LysRS,PAN3,dom,UbcE2H,Vha55,Lk6,Gapdh1,Ubc4,TER94,Vha68-2,Ndf,SRPK,Hsp60A,cno,CCT5,Hex-A,Dlic,CCT3,CkIIalpha,Hsc70Cb,CCT4,CCT1,Hel25E,blw,CCT6,gish,Rpt2,Mi-2,Sam-S,Galphai,Rpt6,CG11267,CkIalpha,ben,Gs1* |
| ATP-dependent protein folding chaperone | GO:MF | 3.63e-09 | *Hsp83,Hsc70-3,CCT2,Hsc70-4,Hsp60A,CCT5,CCT3,Hsc70Cb,CCT4,CCT1,CCT6,CG11267* |
| ATP binding | GO:MF | 4.48e-09 | *wisp,me31B,Top2,PyK,polo,RnrL,lok,rdog,grp,glu,Uba2,Uba1,awd,Tao,kis,Droj2,Tlk,Hsp83,Hsc70-3,dco,mbt,for,CCT2,mnb,Hsc70-4,LysRS,PAN3,dom,UbcE2H,Vha55,Lk6,Ubc4,TER94,Vha68-2,SRPK,Hsp60A,cno,CCT5,Hex-A,Dlic,CCT3,CkIIalpha,Hsc70Cb,CCT4,CCT1,Hel25E,blw,CCT6,gish,Rpt2,Mi-2,Sam-S,Rpt6,CG11267,CkIalpha,ben,Gs1* |
| adenyl ribonucleotide binding | GO:MF | 5.27e-09 | *wisp,me31B,Top2,PyK,polo,RnrL,lok,rdog,grp,glu,Uba2,Uba1,awd,Tao,kis,Droj2,Tlk,Hsp83,Hsc70-3,dco,mbt,for,CCT2,mnb,Hsc70-4,LysRS,PAN3,dom,UbcE2H,Vha55,Lk6,Ubc4,TER94,Vha68-2,SRPK,Hsp60A,cno,CCT5,Hex-A,Dlic,CCT3,CkIIalpha,Hsc70Cb,CCT4,CCT1,Hel25E,blw,CCT6,gish,Rpt2,Mi-2,Sam-S,Rpt6,CG11267,CkIalpha,ben,Gs1* |
| nucleotide binding | GO:MF | 4.66e-08 | *wisp,alphaTub67C,me31B,Top2,PyK,Gapdh2,polo,betaTub56D,RnrL,lok,rdog,grp,Trxr-1,glu,Uba2,Uba1,awd,Ran,eEF2,Tao,kis,Droj2,Tlk,CtBP,Hsp83,Hsc70-3,dco,mbt,for,CCT2,mnb,Hsc70-4,Idh,LysRS,PAN3,dom,UbcE2H,Vha55,Lk6,Gapdh1,Ubc4,TER94,Vha68-2,Ndf,SRPK,Hsp60A,cno,CCT5,Hex-A,Dlic,CCT3,CkIIalpha,Hsc70Cb,CCT4,CCT1,Hel25E,blw,CCT6,gish,Rpt2,Mi-2,Sam-S,Galphai,Rpt6,CG11267,CkIalpha,ben,Gs1* |
| mRNA binding | GO:MF | 5.29e-08 | *bru1,smg,Marf1,me31B,Top2,tral,CG9705,srl,rin,sqd,CNBP,brat,Hrb87F,Unr,Rox8,hang,Cpsf6,eIF4G1,stau,RpS5b,mod,heph,Hrb98DE,Ref1,eIF3b,eIF4H1* |
| RNA binding | GO:MF | 5.29e-08 | *bru1,wisp,smg,exu,Marf1,me31B,Top2,tral,msk,Nop60B,CG9705,srl,rin,sqd,CNBP,brat,Capr,nop5,Hrb87F,Nlp,ncm,Unr,Rox8,hang,SmD3,LysRS,PAN3,Cpsf6,Tudor-SN,eIF4G1,stau,RpS5b,SNRPG,spoon,Surf6,Nph,hoip,mod,gw,mask,heph,Hrb98DE,Hel25E,bin3,vig2,Ref1,Helz,Nop56,eIF3b,eIF4H1* |
| histone binding | GO:MF | 5.39e-08 | *CG8223,Caf1-180,Df31,Mapmodulin,e(y)3,kis,Dek,Nlp,E(bx),Nap1,dom,Brd7-9,Nph,upSET,CG8677,Mi-2,d4* |
| protein folding chaperone | GO:MF | 5.67e-08 | *Hsp83,Hsc70-3,CCT2,Hsc70-4,Hsp60A,CCT5,CCT3,Hsc70Cb,CCT4,CCT1,CCT6,CG11267* |
| translation repressor activity | GO:MF | 3.12e-07 | *smg,Marf1,me31B,brat,wech,Unr,Gyf,Paip2,heph* |
| anion binding | GO:MF | 5.59e-07 | *wisp,alphaTub67C,me31B,Top2,PyK,polo,betaTub56D,RnrL,lok,rdog,grp,Trxr-1,glu,Uba2,Uba1,awd,Ran,eEF2,Tao,kis,Droj2,Tlk,Hsp83,Hsc70-3,dco,mbt,for,CCT2,mnb,Hsc70-4,LysRS,PAN3,dom,UbcE2H,Vha55,Lk6,baz,Ubc4,TER94,Vha68-2,CG8036,SRPK,Hsp60A,cno,CCT5,Hex-A,Dlic,Moe,CCT3,CkIIalpha,Hsc70Cb,apolpp,CCT4,CCT1,Hel25E,blw,CCT6,gish,Rpt2,Mi-2,Sam-S,Galphai,Rpt6,CG11267,CkIalpha,ben,Gs1* |
| mRNA 3'-UTR binding | GO:MF | 1.28e-06 | *bru1,smg,CG9705,sqd,brat,Hrb87F,Unr,Rox8,stau,heph,Hrb98DE* |
| chromatin binding | GO:MF | 2.18e-06 | *BigH1,zld,Top2,PCNA,Rcc1,glu,msl-1,e(y)3,Mtor,kis,woc,Nlp,Nap1,osa,Nipped-B,Ndf,Nph,Nup153,upSET,Sin3A,nej,nsl1,Mi-2,wcy,wge* |
| small molecule binding | GO:MF | 8.65e-06 | *wisp,yl,alphaTub67C,Pxt,me31B,zld,Nup358,Top2,ovo,PyK,Cyp6a19,RnrS,dUTPase,Gapdh2,Cam,polo,betaTub56D,RnrL,lok,rdog,grp,Trxr-1,glu,CG32473,cue,Su(var)2-10,Uba2,Snp,Uba1,awd,Ran,CNBP,e(y)3,brat,eEF2,Tao,G9a,kis,LpR2,woc,Droj2,Tlk,CtBP,Hsp83,wech,Hsc70-3,E(bx),dco,Rpn2,Aef1,mbt,for,CCT2,mnb,CG2926,chn,CG45050,hang,Desat1,Hsc70-4,Idh,Kdm5,LysRS,PAN3,mts,dom,UbcE2H,Vha55,Lk6,Gapdh1,baz,Ubc4,TER94,Vha68-2,CG8036,Ndf,SRPK,Hsp60A,cno,CCT5,Hex-A,Pp2C1,Polr2A,Dlic,Sod1,Moe,CCT3,CkIIalpha,Nup153,upSET,sov,CG8677,nej,Hsc70Cb,apolpp,CanA-14F,Ufd4,CCT4,CG1677,Rpn11,CCT1,GlcAT-P,Hel25E,blw,Pde11,unc-13,Helz,mei-P26,CCT6,gish,Rpt2,Mi-2,Sam-S,Galphai,Rpt6,CG11267,CkIalpha,ben,Gs1,d4,tou,cg,sina* |
| carbohydrate derivative binding | GO:MF | 9.76e-06 | *wisp,alphaTub67C,me31B,Top2,PyK,polo,betaTub56D,RnrL,lok,rdog,grp,glu,Uba2,Uba1,awd,Ran,eEF2,Tao,kis,Droj2,Tlk,Hsp83,Hsc70-3,dco,mbt,for,CCT2,mnb,Hsc70-4,LysRS,PAN3,dom,UbcE2H,Vha55,Lk6,Ubc4,TER94,Vha68-2,SRPK,Hsp60A,cno,CCT5,Hex-A,Dlic,CCT3,CkIIalpha,Hsc70Cb,CCT4,CCT1,Hel25E,blw,CCT6,gish,Rpt2,Mi-2,Sam-S,Galphai,PlexA,Rpt6,CG11267,CkIalpha,ben,Gs1* |
| translation regulator activity | GO:MF | 2.21e-05 | *smg,Marf1,me31B,CNBP,brat,eEF2,wech,Unr,Gyf,Paip2,eIF4G1,heph,eIF3b,eIF4H1* |
| ATP hydrolysis activity | GO:MF | 3.65e-05 | *me31B,Top2,rdog,glu,kis,Hsp83,Hsc70-3,CCT2,Hsc70-4,dom,TER94,Vha68-2,Hsp60A,CCT5,CCT3,Hsc70Cb,CCT4,CCT1,Hel25E,CCT6,Rpt2,Mi-2,Rpt6* |
| molecular adaptor activity | GO:MF | 3.91e-05 | *Marf1,zld,Su(var)2-10,srl,e(y)3,CtBP,gro,wech,E(bx),MED26,mnb,rdx,Rox8,Kdm5,Cul1,BicD,wap,gcl,Sin3A,CG8677,nej,Not1,CG1677,milt,Gug,skd,Pits,d4* |
| enzyme binding | GO:MF | 3.99e-05 | *tral,msk,CycB3,Uba2,msl-1,Tnpo,emb,Mtor,ago,Gbs-70E,mbt,rdx,Sumo,Ptp61F,Myc,Lk6,Fs(2)Ket,Ubc4,Cul1,cno,Uev1A,Pp2C1,BicD,mask,nej,l(3)80Fj,nsl1,wcy,dsd,ben,sina* |
| unfolded protein binding | GO:MF | 3.99e-05 | *Hsp27,Droj2,Hsp83,Hsc70-3,CCT2,Hsc70-4,CCT5,CCT3,CCT4,CCT1,CCT6,CG11267* |
| protein-macromolecule adaptor activity | GO:MF | 5.44e-05 | *Marf1,Su(var)2-10,srl,e(y)3,CtBP,gro,wech,E(bx),MED26,mnb,rdx,Rox8,Kdm5,Cul1,BicD,wap,gcl,Sin3A,CG8677,nej,CG1677,milt,Gug,skd,Pits,d4* |
| mRNA regulatory element binding translation repressor activity | GO:MF | 1.02e-04 | *smg,me31B,Unr,Gyf,Paip2,heph* |
| DNA-binding transcription factor binding | GO:MF | 1.45e-04 | *retn,Su(var)2-10,CtBP,Dp,gro,E(bx),nej,bin3,Gug,cact,da,tou* |
| pyrophosphatase activity | GO:MF | 2.43e-04 | *me31B,Top2,dUTPase,betaTub56D,rdog,glu,Ran,eEF2,kis,Hsp83,Hsc70-3,CCT2,Hsc70-4,dom,TER94,Vha68-2,Hsp60A,Aps,CCT5,CCT3,Hsc70Cb,CCT4,CCT1,Hel25E,CCT6,Rpt2,Mi-2,Galphai,Rpt6* |
| hydrolase activity, acting on acid anhydrides | GO:MF | 2.72e-04 | *me31B,Top2,dUTPase,betaTub56D,rdog,glu,Ran,eEF2,kis,Hsp83,Hsc70-3,CCT2,Hsc70-4,dom,TER94,Vha68-2,Hsp60A,Aps,CCT5,CCT3,Hsc70Cb,CCT4,CCT1,Hel25E,CCT6,Rpt2,Mi-2,Galphai,Rpt6* |
| hydrolase activity, acting on acid anhydrides, in phosphorus-containing anhydrides | GO:MF | 2.72e-04 | *me31B,Top2,dUTPase,betaTub56D,rdog,glu,Ran,eEF2,kis,Hsp83,Hsc70-3,CCT2,Hsc70-4,dom,TER94,Vha68-2,Hsp60A,Aps,CCT5,CCT3,Hsc70Cb,CCT4,CCT1,Hel25E,CCT6,Rpt2,Mi-2,Galphai,Rpt6* |
| transcription factor binding | GO:MF | 2.85e-04 | *retn,ovo,Su(var)2-10,srl,CtBP,Dp,gro,E(bx),TfIIFalpha,nej,bin3,Gug,cact,da,tou* |
| cyclin-dependent protein serine/threonine kinase regulator activity | GO:MF | 3.11e-04 | *CycB,CycA,CycE,CycB3,dap,CycT* |
| nuclear import signal receptor activity | GO:MF | 3.63e-04 | *Pen,Tnpo,Fs(2)Ket,Kap-alpha3,Ntf-2* |
| ribonucleoside triphosphate phosphatase activity | GO:MF | 4.14e-04 | *me31B,Top2,betaTub56D,rdog,glu,Ran,eEF2,kis,Hsp83,Hsc70-3,CCT2,Hsc70-4,dom,TER94,Vha68-2,Hsp60A,CCT5,CCT3,Hsc70Cb,CCT4,CCT1,Hel25E,CCT6,Rpt2,Mi-2,Galphai,Rpt6* |
| protein-containing complex binding | GO:MF | 6.09e-04 | *BigH1,Marf1,zld,eEF2,Mtor,LpR2,Usp14,His2Av,Unr,Jarid2,Ubqn,osa,Arpc2,Ubc4,Ndf,Spn,TfIIFalpha,BicD,kra,bif,CG1677,Rpn11,l(3)80Fj,Galphai,eIF4H1* |
| nuclear localization sequence binding | GO:MF | 9.57e-04 | *Pen,Tnpo,Fs(2)Ket,Nup153,Kap-alpha3* |
| protein kinase regulator activity | GO:MF | 0.002 | *CycB,CycA,gnu,CycE,Incenp,CycB3,mnb,l(3)80Fj,dap,CycT* |
| cysteine-type endopeptidase activator activity involved in apoptotic process | GO:MF | 0.002 | *26-29-p,RnrS,RnrL,lok* |
| DNA binding | GO:MF | 0.003 | *mtrm,BigH1,retn,zld,Top2,PCNA,ovo,msl-1,Mtor,kis,Su(var)2-HP2,Dek,Hrb87F,Fkbp39,Dp,His2Av,E(bx),Aef1,D1,Jarid2,chn,CG45050,hang,Top1,Ssdp,jigr1,Kdm5,Mitf,Myc,cnc,dom,osa,Surf6,Ndf,baf,psq,TfIIFalpha,Polr2A,mod,Nup153,nej,Hrb98DE,BtbVII,Mi-2,Galphai,CG32767,da,tou,cg,Mnt* |
| RNA polymerase II-specific DNA-binding transcription factor binding | GO:MF | 0.004 | *Su(var)2-10,E(bx),nej,bin3,Gug,cact* |
| proteasome binding | GO:MF | 0.004 | *Usp14,Ubqn,Ubc4,Rpn11* |
| kinase regulator activity | GO:MF | 0.004 | *CycB,CycA,gnu,CycE,Incenp,CycB3,mnb,l(3)80Fj,dap,CycT* |
| transcription coregulator activity | GO:MF | 0.004 | *Su(var)2-10,srl,e(y)3,CtBP,gro,E(bx),MED26,mnb,Sin3A,CG8677,nej,Gug,skd,Pits,d4* |
| protein serine/threonine kinase activity | GO:MF | 0.005 | *polo,lok,grp,Tao,Tlk,dco,mbt,for,mnb,Lk6,SRPK,CkIIalpha,gish,CkIalpha* |
| cysteine-type endopeptidase regulator activity involved in apoptotic process | GO:MF | 0.006 | *26-29-p,RnrS,RnrL,lok* |
| histone acetyltransferase activity | GO:MF | 0.006 | *e(y)3,CG8677,nej,d4* |
| ubiquitin conjugating enzyme activity | GO:MF | 0.008 | *CG10254,CG2924,UbcE2H,Ubc4,ben* |
| ribonucleoside-diphosphate reductase activity, thioredoxin disulfide as acceptor | GO:MF | 0.008 | *RnrS,RnrL* |
| single-stranded RNA binding | GO:MF | 0.009 | *exu,CNBP,Hrb87F,PAN3,Hrb98DE,eIF4H1* |
| threonine-type endopeptidase activity | GO:MF | 0.01 | *Prosbeta5,Prosbeta2,Prosbeta1* |
| peptide-lysine-N-acetyltransferase activity | GO:MF | 0.012 | *e(y)3,CG8677,nej,d4* |
| enzyme regulator activity | GO:MF | 0.012 | *CycB,26-29-p,CycA,gnu,Nup358,PCNA,CycE,Rcc1,RnrS,Cam,Incenp,RnrL,lok,CycB3,tws,Rpn2,mnb,Kdm5,mts,endos,Dhit,Gdi,l(3)80Fj,PlexA,dap,CycT* |
| signal sequence binding | GO:MF | 0.013 | *Pen,Tnpo,Fs(2)Ket,Nup153,Kap-alpha3* |
| catalytic activity | GO:MF | 0.018 | *dhd,wisp,Jafrac1,26-29-p,alphaTub67C,CycA,Pxt,me31B,Nup358,Top2,Ggt-1,CycE,PyK,Cyp6a19,CG46385,RnrS,dUTPase,Gapdh2,CG10254,polo,Nop60B,betaTub56D,RnrL,lok,rdog,grp,Akr1B,CycB3,Trxr-1,glu,CG32473,CG7840,mt:ATPase6,Su(var)2-10,Uba2,Snp,Uba1,awd,Ran,rin,e(y)3,eEF2,Tao,G9a,kis,Usp7,Tl,Usp14,Mgstl,Tlk,CtBP,Hsp83,Fkbp39,GstD1,Hsc70-3,dco,Jarid2,CG2924,mbt,for,CCT2,mnb,egh,Top1,Echs1,Desat1,Hsc70-4,Ptp61F,Idh,Kdm5,LysRS,Tudor-SN,mts,dom,UbcE2H,Mdh1,Lk6,Gapdh1,Nipped-B,Ubc4,TER94,Vha68-2,CG8036,Ndf,Prosbeta5,SRPK,Hsp60A,Prosalpha6,Aps,CCT5,Hex-A,Usp10,Pp2C1,Polr2A,Sod1,CCT3,CkIIalpha,CG3655,CG8677,nej,Hsc70Cb,CanA-14F,Ufd4,Not1,CCT4,Rpn11,CCT1,Ahcy,GlcAT-P,Hel25E,Nedd4,blw,bin3,Prosbeta2,Pde11,Helz,mei-P26,CCT6,gish,Rpt2,Mi-2,mt:ND1,Sam-S,Galphai,Rpt6,Gug,CkIalpha,Prosbeta4,Pits,ben,Prosbeta1,Gs1,CycT,d4,fwd,Pglym78,sina* |
| poly(G) binding | GO:MF | 0.019 | *Hrb87F,Hrb98DE* |
| glyceraldehyde-3-phosphate dehydrogenase (NAD+) (phosphorylating) activity | GO:MF | 0.019 | *Gapdh2,Gapdh1* |
| peptidyl-cysteine S-nitrosylase activity | GO:MF | 0.019 | *Gapdh2,Gapdh1* |
| disordered domain specific binding | GO:MF | 0.019 | *Hsp83,ctp* |
| protein domain specific binding | GO:MF | 0.025 | *Tl,Hsp83,gro,ctp,Vap33,cno,psq,Nedd4* |
| NAD binding | GO:MF | 0.031 | *Gapdh2,CtBP,Idh,Gapdh1,Ndf* |
| satellite DNA binding | GO:MF | 0.032 | *Top2,D1* |
| histone reader activity | GO:MF | 0.032 | *Kdm5,CG8677* |
| identical protein binding | GO:MF | 0.032 | *exu,Drep2,dUTPase,Trxr-1,Tl,CtBP,rdx,jigr1,ctp,psq,Sod1,da,sina* |
| ubiquitin-like protein ligase binding | GO:MF | 0.032 | *msl-1,rdx,Sumo,Ubc4,Cul1,dsd,ben* |
| mitogen-activated protein kinase binding | GO:MF | 0.032 | *Lk6,Pp2C1* |
| peptidase activator activity | GO:MF | 0.032 | *26-29-p,RnrS,RnrL,lok* |
| transcription corepressor activity | GO:MF | 0.035 | *e(y)3,CtBP,gro,Sin3A,Gug,Pits* |
| transferase activity | GO:MF | 0.036 | *wisp,CycA,Nup358,Ggt-1,CycE,PyK,CG46385,Gapdh2,CG10254,polo,lok,grp,CycB3,Su(var)2-10,awd,e(y)3,Tao,G9a,Mgstl,Tlk,GstD1,dco,CG2924,mbt,for,mnb,egh,UbcE2H,Lk6,Gapdh1,Ubc4,CG8036,SRPK,Hex-A,Polr2A,CkIIalpha,CG3655,CG8677,nej,Ufd4,GlcAT-P,Nedd4,bin3,mei-P26,gish,Sam-S,CkIalpha,Pits,ben,CycT,d4,fwd,sina* |
| enzyme activator activity | GO:MF | 0.039 | *26-29-p,gnu,Nup358,PCNA,RnrS,Incenp,RnrL,lok,Dhit,Gdi,PlexA,CycT* |
| phosphotransferase activity, alcohol group as acceptor | GO:MF | 0.042 | *PyK,polo,lok,grp,Tao,Tlk,dco,mbt,for,mnb,Lk6,SRPK,Hex-A,CkIIalpha,gish,CkIalpha,fwd* |
| protein kinase activity | GO:MF | 0.042 | *polo,lok,grp,Tao,Tlk,dco,mbt,for,mnb,Lk6,SRPK,CkIIalpha,gish,CkIalpha* |
| antioxidant activity | GO:MF | 0.044 | *Jafrac1,Pxt,Trxr-1,Mgstl,GstD1,Sod1* |
| histone chaperone activity | GO:MF | 0.046 | *CG8223,dom* |
| DNA topoisomerase activity | GO:MF | 0.046 | *Top2,Top1* |
| protein-RNA adaptor activity | GO:MF | 0.046 | *Marf1,Rox8* |
| histone acetyltransferase binding | GO:MF | 0.046 | *Mtor,nsl1* |
| ubiquitin-specific protease binding | GO:MF | 0.046 | *ago,Myc* |

### 1.3. Gene Ontology Enrichment Analysis: MM cell body glia

Significant GO terms (p < 0.05) with associated genes

| GO Term | Source | P-value | Genes |
| --- | --- | --- | --- |
| tail-anchored membrane protein insertion into ER membrane | GO:BP | 3.39e-07 | *EMC7,EMC6,EMC5,EMC3,EMC1,EMC4,EMC8-9* |
| proton transmembrane transport | GO:BP | 4.20e-06 | *Vha68-2,VhaPPA1-1,Vha44,Vha14-1,Vha26,VhaM9.7-a,Vha13,VhaM9.7-b,Vha55,ATP6AP2,VhaAC39-1,Atpalpha,Vha36-1,Vha16-1,VhaSFD,VhaM9.7-c* |
| glycolytic process | GO:BP | 2.64e-05 | *Tpi,Eno,Gapdh2,Pglym78,Pgk,Ald1,Pfk,PyK,Pgi* |
| nucleotide catabolic process | GO:BP | 2.91e-05 | *Tpi,Eno,Pde9,Gapdh2,Pglym78,Pgk,Ald1,Pfk,PyK,Pgi,CG17224* |
| nucleoside diphosphate catabolic process | GO:BP | 3.28e-05 | *Tpi,Eno,Gapdh2,Pglym78,Pgk,Ald1,Pfk,PyK,Pgi* |
| signal release | GO:BP | 5.34e-05 | *nemy,CDase,Arl8,Snx3,opm,Syb,Arf6,sky,InR,Rap1,eca,NFAT,alphaSnap,CHOp24,Cadps,p24-1,mth,AdipoR* |
| organic acid metabolic process | GO:BP | 1.08e-04 | *Gabat,Tpi,Eno,e,Ssadh,Ldh,Gapdh2,Pglym78,CDase,Acbp2,Pgk,sgl,Gdh,Ald1,Mdh1,yip2,Gcat,Pfk,Baldspot,CG7920,CG1640,PyK,CG1764,Fatp1,CG4598,Etfb,Pgi,Idh,Mtpbeta,CG1440,Got1,AdipoR,Acbp1* |
| hemocyte migration | GO:BP | 1.75e-04 | *Pvf3,Ras85D,Mtl,shg,Rac2,Rho1,Rap1* |
| glucose homeostasis | GO:BP | 6.35e-04 | *Tpi,Eno,Gapdh2,Ald1,Pfk,PyK,InR,Pgi,AdipoR* |
| intracellular monoatomic ion homeostasis | GO:BP | 6.77e-04 | *zyd,nrv2,nrv1,Vha68-2,VhaAC45,Fer1HCH,ClC-c,Vha55,VhaAC39-1,Fer2LCH,BI-1,CG10470,Atpalpha,CG17593,Vha16-1* |
| terminal branching, open tracheal system | GO:BP | 7.60e-04 | *VhaPPA1-1,Vha26,Vha13,VhaAC39-1,Ras85D,Zyx,Rheb* |
| glucose catabolic process | GO:BP | 7.83e-04 | *Eno,Pfk,PhKgamma* |
| dorsal closure | GO:BP | 0.001 | *Rab5,Rab11,Ras85D,emc,Mtl,rho,spi,Ggamma1,Inx3,Rac2,Rho1,scrib,Rap1* |
| ribose phosphate metabolic process | GO:BP | 0.001 | *Tpi,Eno,Pde9,Aprt,Gapdh2,Prps,Pglym78,Pgk,Vha68-2,Ald1,Pfk,Vha55,PyK,Ac13E,Pgi,CG2246* |
| cell surface receptor signaling pathway | GO:BP | 0.001 | *SoxN,Wnt4,sli,GlyP,Akap200,Pvf3,pnt,Psn,sgl,fng,Tom,spin,edl,stumps,Prosap,for,Rgl,htl,ATP6AP2,spg,Ras85D,tow,shv,boca,Sema1a,myo,shg,rho,sty,par-1,spi,InR,Rala,rau,Rac2,mav,Rho1,pigs,Rap1,sima,lqf,GEFmeso,mth,AdipoR,Rheb,E(spl)mbeta-HLH,Tsp86D,Mnr* |
| membrane fusion | GO:BP | 0.001 | *aus,CDase,Arl8,Rab5,Snx6,Rab11,Syb,Vti1b,Rab7,alphaSnap,Bet1* |
| ommatidial rotation | GO:BP | 0.001 | *pnt,Ras85D,shg,sty,spi,Frl,Rap1* |
| positive regulation of epidermal growth factor receptor signaling pathway | GO:BP | 0.001 | *edl,Rgl,Rala,rau,GEFmeso* |
| animal organ morphogenesis | GO:BP | 0.002 | *Wnt4,zfh2,nrv2,Dll,sli,ct,pnt,fng,Tom,Sdb,Rab5,ATP6AP2,Nrg,Rab11,spg,Atpalpha,Ras85D,emc,Mtl,tow,RhoGAP100F,Rbfox1,myo,shg,rho,sty,par-1,Arf6,spi,Ggamma1,14-3-3epsilon,Amph,corto,Rala,EMC3,rau,Rac2,mav,svr,Rho1,Frl,scrib,Rap1,alphaSnap,GEFmeso,Septin1,Zyx* |
| dsRNA transport | GO:BP | 0.003 | *Sap-r,Vha16-1,AP-2mu,VhaSFD,Rab7,CG8671* |
| endoplasmic reticulum to Golgi vesicle-mediated transport | GO:BP | 0.003 | *KdelR,boca,opm,Sar1,eca,CHOp24,cni,p24-1,Bet1,epsilonCOP* |
| NADH metabolic process | GO:BP | 0.003 | *Eno,Gdh,Mdh1,Pfk* |
| apical protein localization | GO:BP | 0.004 | *Vha26,shg,Ggamma1,Rab35,Rheb* |
| organelle fusion | GO:BP | 0.004 | *aus,CDase,Arl8,Rab5,Snx6,Rab11,Syb,Vti1b,Rab7,alphaSnap,Bet1* |
| endoplasmic reticulum unfolded protein response | GO:BP | 0.004 | *BI-1,Hsc70-3,Xbp1,Der-1,Atf6* |
| stem cell fate commitment | GO:BP | 0.008 | *Ras85D,rho,spi* |
| photoreceptor cell fate determination | GO:BP | 0.008 | *Ras85D,rho,spi* |
| viral process | GO:BP | 0.008 | *Rab5,Rab11,Mvl,Sar1,Rab35,Rab7,Vps29* |
| long-term memory | GO:BP | 0.009 | *fabp,for,vsg,ATP6AP2,Tob,svr,be,CG4612,mld* |
| gastrulation | GO:BP | 0.01 | *sli,sgl,stumps,htl,Ras85D,shg,spi,Ggamma1,Rho1,Rab35* |
| 5-phosphoribose 1-diphosphate biosynthetic process | GO:BP | 0.011 | *Prps,CG2246* |
| endosomal lumen acidification | GO:BP | 0.011 | *Vha68-2,ClC-c* |
| negative regulation of lipophagy | GO:BP | 0.011 | *ifc,Rheb* |
| canonical glycolysis | GO:BP | 0.011 | *Eno,Pfk* |
| iron ion import across plasma membrane | GO:BP | 0.011 | *Fer1HCH,Fer2LCH* |
| detoxification of iron ion | GO:BP | 0.011 | *Fer1HCH,Fer2LCH* |
| positive regulation of P-type sodium:potassium-exchanging transporter activity | GO:BP | 0.011 | *nrv2,nrv1* |
| positive regulation of wound healing | GO:BP | 0.011 | *Mtl,InR,Rac2,Rho1* |
| gamma-aminobutyric acid catabolic process | GO:BP | 0.011 | *Gabat,Ssadh* |
| female courtship behavior | GO:BP | 0.011 | *shep,Nrg* |
| lymph gland plasmatocyte differentiation | GO:BP | 0.012 | *pnt,htl,Ras85D* |
| glial cell migration | GO:BP | 0.013 | *sli,Pvf3,spin,htl,Rho1,orion* |
| leg disc proximal/distal pattern formation | GO:BP | 0.016 | *Dll,pnt,Ras85D,rho* |
| endoplasmic reticulum calcium ion homeostasis | GO:BP | 0.016 | *BI-1,CG10470,CG17593* |
| exosomal secretion | GO:BP | 0.016 | *Rab11,Rab35,Rab7* |
| positive regulation of stem cell proliferation | GO:BP | 0.016 | *Rgl,Rala,GEFmeso* |
| dorsal closure, elongation of leading edge cells | GO:BP | 0.016 | *Rab11,Mtl,Rac2* |
| glycosylation | GO:BP | 0.017 | *fng,NANS,Pgant5,CG31690,Ostgamma,Dad1,CG33303,kud,Ost48,CG33774,OstDelta,Stt3B* |
| wing disc dorsal/ventral pattern formation | GO:BP | 0.017 | *pnt,Psn,fng,AnxB9,14-3-3epsilon,drpr,Rab7* |
| protein N-linked glycosylation via asparagine | GO:BP | 0.019 | *Ostgamma,CG33303,Ost48,Stt3B* |
| potassium ion import across plasma membrane | GO:BP | 0.019 | *Irk3,nrv2,nrv1,Atpalpha* |
| epidermal cell differentiation | GO:BP | 0.021 | *ATP6AP2,Mtl,tow,par-1,Rho1,scrib,Rap1* |
| determination of adult lifespan | GO:BP | 0.021 | *GLaz,Tpi,GlyP,pnt,Sod1,Atpalpha,Ras85D,myo,14-3-3epsilon,InR,VhaSFD,Mtpbeta,mth,mld* |
| trans-synaptic signaling | GO:BP | 0.021 | *Csas,nemy,CDase,Ace,Pgk,cpo,Arl8,Sod1,Manf,Ten-a,Rbp,Ent2,Atpalpha,Syb,Arf6,sky,Rap1,be,NFAT,alphaSnap,Cadps,mth* |
| extracellular vesicle biogenesis | GO:BP | 0.022 | *Rab11,Rab35,Rab7* |
| intracellular sodium ion homeostasis | GO:BP | 0.022 | *nrv2,nrv1,Atpalpha* |
| determination of genital disc primordium | GO:BP | 0.022 | *pnt,rho,spi* |
| intracellular potassium ion homeostasis | GO:BP | 0.022 | *nrv2,nrv1,Atpalpha* |
| intercellular transport | GO:BP | 0.022 | *ogre,Inx2,Inx3* |
| sodium ion export across plasma membrane | GO:BP | 0.022 | *nrv2,nrv1,Atpalpha* |
| signal peptide processing | GO:BP | 0.022 | *Spase22-23,Spase25,twr* |
| catabolic process | GO:BP | 0.022 | *CG40470,Gabat,GLaz,Tpi,Eno,Pde9,Ssadh,aus,GlyP,Gapdh2,Cp1,Pglym78,CDase,Ace,olf413,Rchy1,Pgk,Arl8,Sod1,Gdh,Ald1,AP-2alpha,CG10365,yip2,Gcat,Pfk,Lsd-2,Tango5,EMC6,CG1640,Hexo2,BI-1,GlyS,PyK,Ras85D,CG12177,Lamp1,CtsB,Ntan1,PhKgamma,InR,Vti1b,CG4598,CG12384,Etfb,Xbp1,Pgi,Rab7,sima,Der-1,lqf,ifc,Mtpbeta,CtsF,sordd1,CG1440,Rheb,EndoA,CG17224* |
| Malpighian tubule bud morphogenesis | GO:BP | 0.024 | *ct,Kr* |
| myoblast fate specification | GO:BP | 0.024 | *htl,Ras85D* |
| protein retention in ER lumen | GO:BP | 0.024 | *KdelR,CG11857* |
| intracellular sequestering of iron ion | GO:BP | 0.024 | *Fer1HCH,Fer2LCH* |
| NAD catabolic process | GO:BP | 0.024 | *Eno,Pfk* |
| L-leucine import across plasma membrane | GO:BP | 0.024 | *mnd,CD98hc* |
| negative regulation of synaptic vesicle exocytosis | GO:BP | 0.024 | *Rap1,NFAT* |
| female germ-line stem cell population maintenance | GO:BP | 0.028 | *fng,fax,shg,InR,AdipoR* |
| gluconeogenesis | GO:BP | 0.028 | *Tpi,Pgk,Pgi* |
| mushroom body development | GO:BP | 0.031 | *Dll,ced-6,Dscam1,Nrg,myo,Frl,orion,robl* |
| intestinal stem cell homeostasis | GO:BP | 0.035 | *Ras85D,Hsc70-3,InR,Xbp1,PHGPx* |
| lymph gland crystal cell differentiation | GO:BP | 0.035 | *pnt,htl,Ras85D* |
| mesoderm migration involved in gastrulation | GO:BP | 0.035 | *sli,sgl,htl* |
| epithelial cell proliferation involved in Malpighian tubule morphogenesis | GO:BP | 0.035 | *pnt,rho,spi* |
| motor neuron axon guidance | GO:BP | 0.037 | *Wnt4,CG42327,Ten-a,for,Nrg,Sema1a,Rac2,Rho1* |
| retrograde transport, endosome to Golgi | GO:BP | 0.038 | *Snx3,Snx6,Snx1,Vti1b,Vps29* |
| amino acid catabolic process | GO:BP | 0.039 | *Gabat,Ssadh,Gdh,Gcat,CG1640,Etfb,CG1440* |
| fructose 1,6-bisphosphate metabolic process | GO:BP | 0.04 | *Ald1,Pfk* |
| branched-chain amino acid transport | GO:BP | 0.04 | *mnd,CD98hc* |
| outflow tract morphogenesis | GO:BP | 0.04 | *sli,shg* |
| negative regulation of cardioblast cell fate specification | GO:BP | 0.04 | *pnt,lqf* |
| biological process involved in interspecies interaction between organisms | GO:BP | 0.041 | *Npc2a,gb,St3,GlyP,Cyp6a20,Dscam1,vir-1,Fer1HCH,Rab5,htl,Fer2LCH,Spn27A,Nrg,Rab11,Ras85D,Mvl,Mtl,RNASEK,shg,loco,drpr,Rala,Rac2,Sar1,Rho1,Rab35,Rab7,sima,NUCB1,cathD,Vps29* |
| cell adhesion involved in heart morphogenesis | GO:BP | 0.042 | *nrv2,Nrg,Ggamma1* |
| engulfment of apoptotic cell | GO:BP | 0.042 | *NimC4,drpr,Rac2* |
| female gonad development | GO:BP | 0.042 | *Wnt4,ct,InR* |
| R3/R4 cell fate commitment | GO:BP | 0.042 | *pnt,Rala,scrib* |
| positive regulation of ERK1 and ERK2 cascade | GO:BP | 0.044 | *spg,Ras85D,spi,Rap1* |
| glutathione metabolic process | GO:BP | 0.048 | *GstE1,GstE2,GstD1,CG10365,GstE11,GstE12* |
| negative regulation of apoptotic signaling pathway | GO:BP | 0.05 | *pnt,BI-1,Ras85D,CG2918* |
| liquid clearance, open tracheal system | GO:BP | 0.05 | *Vha26,Vha13,Zyx,colt* |
| calcium ion transmembrane transport | GO:BP | 0.05 | *zyd,pain,BI-1,Cnx99A,CG10470,Nmda1,fwe* |
| septate junction assembly | GO:BP | 0.05 | *nrv2,Nrg,Atpalpha,loco,scrib* |
| proton-transporting ATPase activity, rotational mechanism | GO:MF | 4.24e-10 | *Vha68-2,VhaPPA1-1,Vha44,Vha14-1,Vha26,VhaM9.7-a,Vha13,VhaM9.7-b,Vha55,VhaAC39-1,Vha36-1,Vha16-1,VhaSFD,VhaM9.7-c* |
| ATPase-coupled monoatomic cation transmembrane transporter activity | GO:MF | 8.85e-10 | *Vha68-2,VhaPPA1-1,Vha44,Vha14-1,Vha26,VhaM9.7-a,Vha13,VhaM9.7-b,Vha55,VhaAC39-1,Atpalpha,Vha36-1,Vha16-1,VhaSFD,VhaM9.7-c* |
| membrane insertase activity | GO:MF | 2.62e-07 | *EMC7,EMC6,EMC5,EMC3,EMC1,EMC4,EMC8-9* |
| ATPase-coupled transmembrane transporter activity | GO:MF | 1.92e-05 | *Vha68-2,VhaPPA1-1,Vha44,Vha14-1,Vha26,VhaM9.7-a,Vha13,VhaM9.7-b,Vha55,VhaAC39-1,Atpalpha,Vha36-1,Vha16-1,Hsc70-3,VhaSFD,VhaM9.7-c* |
| transporter activity | GO:MF | 9.93e-05 | *Npc2a,zyd,mnd,Irk3,CG3168,nebu,gb,hoe1,Gat,CG32407,bumpel,nemy,kumpel,CD98hc,Picot,CG30344,CG5888,ogre,Vha68-2,mrva,VhaPPA1-1,Vha44,spin,CG18549,Vha14-1,Vha26,pain,VhaM9.7-a,Sfxn1-3,Vha13,Ent1,ClC-c,Mtp,Orp8,VhaM9.7-b,EMC5,Vha55,VhaAC39-1,BI-1,CG10470,Ent2,Atpalpha,ATP8A,Vha36-1,Fatp1,Mvl,Vha16-1,Hsc70-3,Inx2,VhaSFD,CG6356,Nmda1,Inx3,fwe,CG11537,VhaM9.7-c,colt* |
| inorganic cation transmembrane transporter activity | GO:MF | 1.42e-04 | *zyd,Irk3,Gat,bumpel,kumpel,CG5888,Vha68-2,VhaPPA1-1,Vha44,Vha14-1,Vha26,pain,VhaM9.7-a,Vha13,VhaM9.7-b,EMC5,Vha55,VhaAC39-1,BI-1,CG10470,Atpalpha,Vha36-1,Mvl,Vha16-1,VhaSFD,Nmda1,fwe,VhaM9.7-c* |
| proton transmembrane transporter activity | GO:MF | 2.67e-04 | *Vha68-2,VhaPPA1-1,Vha44,Vha14-1,Vha26,VhaM9.7-a,Vha13,VhaM9.7-b,Vha55,VhaAC39-1,Vha36-1,Vha16-1,VhaSFD,VhaM9.7-c* |
| transmembrane transporter activity | GO:MF | 5.47e-04 | *zyd,mnd,Irk3,CG3168,nebu,gb,hoe1,Gat,bumpel,nemy,kumpel,CD98hc,Picot,CG30344,CG5888,ogre,Vha68-2,mrva,VhaPPA1-1,Vha44,spin,CG18549,Vha14-1,Vha26,pain,VhaM9.7-a,Sfxn1-3,Vha13,Ent1,ClC-c,VhaM9.7-b,EMC5,Vha55,VhaAC39-1,BI-1,CG10470,Ent2,Atpalpha,Vha36-1,Fatp1,Mvl,Vha16-1,Hsc70-3,Inx2,VhaSFD,CG6356,Nmda1,Inx3,fwe,CG11537,VhaM9.7-c,colt* |
| GDP binding | GO:MF | 8.94e-04 | *Arl8,Ras85D,Rala,Rap1,Rheb* |
| active transmembrane transporter activity | GO:MF | 0.001 | *zyd,Gat,bumpel,nemy,kumpel,Picot,Vha68-2,VhaPPA1-1,Vha44,Vha14-1,Vha26,VhaM9.7-a,Vha13,VhaM9.7-b,Vha55,VhaAC39-1,Atpalpha,Vha36-1,Mvl,Vha16-1,Hsc70-3,VhaSFD,VhaM9.7-c* |
| monoatomic cation transmembrane transporter activity | GO:MF | 0.001 | *zyd,Irk3,Gat,bumpel,kumpel,CG5888,Vha68-2,VhaPPA1-1,Vha44,Vha14-1,Vha26,pain,VhaM9.7-a,Vha13,VhaM9.7-b,Vha55,VhaAC39-1,BI-1,CG10470,Atpalpha,Vha36-1,Mvl,Vha16-1,VhaSFD,Nmda1,fwe,VhaM9.7-c* |
| GTPase activity | GO:MF | 0.002 | *Arl8,Rab5,Rab11,Ras85D,Mtl,EMC10,Arf6,Rala,Rac2,Sar1,Rho1,Rab35,Rap1,Rab7,Septin1,Rheb* |
| guanyl nucleotide binding | GO:MF | 0.004 | *Arl8,Gdh,for,Rab5,Rab11,Ras85D,Mtl,EMC10,Arf6,Rala,Rac2,Sar1,Rho1,Rab35,Rap1,Rab7,Septin1,Rheb* |
| guanyl ribonucleotide binding | GO:MF | 0.004 | *Arl8,Gdh,for,Rab5,Rab11,Ras85D,Mtl,EMC10,Arf6,Rala,Rac2,Sar1,Rho1,Rab35,Rap1,Rab7,Septin1,Rheb* |
| lipid binding | GO:MF | 0.004 | *fabp,Npc2a,NimC4,Acbp2,Snx3,Mtp,Orp8,CG3246,Snx6,Snx1,AnxB9,AP-2mu,Amph,drpr,sky,AnxB11,MSBP,lqf,gce,Acbp1,EndoA* |
| cargo receptor activity | GO:MF | 0.005 | *santa-maria,LRP1,CG40006,opm,drpr,eca,CHOp24,p24-1* |
| disulfide oxidoreductase activity | GO:MF | 0.005 | *CG11007,SelT,Alr,Grx1,CG7484,ERp60,CaBP1* |
| GTP binding | GO:MF | 0.007 | *Arl8,Gdh,Rab5,Rab11,Ras85D,Mtl,EMC10,Arf6,Rala,Rac2,Sar1,Rho1,Rab35,Rap1,Rab7,Septin1,Rheb* |
| monoatomic ion transmembrane transporter activity | GO:MF | 0.007 | *zyd,Irk3,hoe1,Gat,bumpel,kumpel,CG5888,Vha68-2,VhaPPA1-1,Vha44,Vha14-1,Vha26,pain,VhaM9.7-a,Sfxn1-3,Vha13,ClC-c,VhaM9.7-b,Vha55,VhaAC39-1,BI-1,CG10470,Atpalpha,Vha36-1,Mvl,Vha16-1,VhaSFD,Nmda1,fwe,VhaM9.7-c* |
| identical protein binding | GO:MF | 0.007 | *Liprin-gamma,Tpi,GlyP,Sod1,Psn,Gdh,Ten-a,stumps,Dscam1,Pfk,boca,shg,corto,InR,rau,fwe,Septin1,Khc-73* |
| signaling receptor binding | GO:MF | 0.019 | *Wnt4,GLaz,sli,Akap200,Pvf3,wake,CG10365,CG8507,Spn,stumps,Prosap,ATP6AP2,shv,Sema1a,myo,spi,mav,ITP,orion,CG30423,spri* |
| protein homodimerization activity | GO:MF | 0.027 | *Liprin-gamma,Tpi,GlyP,Sod1,Psn,Ten-a,Dscam1,shg,corto,rau,Septin1,Khc-73* |
| ribose phosphate diphosphokinase activity | GO:MF | 0.028 | *Prps,CG2246* |
| 2-hydroxyglutarate dehydrogenase activity | GO:MF | 0.028 | *Ldh,L2HGDH* |
| oxidoreductase activity, acting on a sulfur group of donors | GO:MF | 0.029 | *CG11007,SelT,Alr,Grx1,CG7484,ERp60,CaBP1* |
| gap junction channel activity | GO:MF | 0.044 | *ogre,Inx2,Inx3* |
| protein binding | GO:MF | 0.045 | *wrapper,CG5758,SoxN,mgl,Wnt4,CG42613,Liprin-gamma,GLaz,LRP1,Fas1,nrv1,Tpi,Nrt,sli,CG32354,zormin,GlyP,Gbs-76A,Wdfy2,Akap200,Pvf3,pnt,CG5888,CG14879,Arl8,Sod1,wake,Psn,ced-6,Gdh,CG14441,Tom,Vha26,CG10365,edl,Clc,Ten-a,pain,sel,CG8507,CG31690,Spn,stumps,Sdb,Gp93,Prosap,Rbp,Vha13,Dscam1,Pfk,CG42673,Lsd-2,Rgl,Rab5,CG1637,htl,ATP6AP2,TBCB,BI-1,Cnx99A,Nrg,Rab11,spg,mtgo,Ras85D,emc,Mtl,shv,RhoGAP100F,boca,AnxB9,Lapsyn,Calr,opm,Syb,Sema1a,Rbfox1,fax,Hsc70-3,myo,LRR,shg,Tob,PhKgamma,par-1,GCS2beta,spi,CG7484,Ggamma1,loco,14-3-3epsilon,Amph,drpr,CG11999,corto,InR,Rala,rau,Vti1b,EMC2A,CG12384,Etfb,stai,Rac2,mav,CG2918,Sar1,EMC1,Rho1,CG3061,Frl,CG4393,pigs,lbk,scrib,CLIP-190,Lamtor3,Rap1,MSBP,fwe,sima,Der-1,EMC8-9,CG2765,alphaSnap,HnRNP-K,lqf,Alg-2,ITP,GEFmeso,cib,mrj,jvl,cni,kud,Septin1,Khc-73,slim,sordd1,TfIIA-S,NKAIN,orion,CG30423,mth,Zyx,gce,Rheb,EndoA,Bet1,CG14818,E(spl)mbeta-HLH,robl,Arpc5,epsilonCOP,spri* |
| hormone binding | GO:MF | 0.048 | *mgl,InR,MSBP,gce* |

### 1.4. Gene Ontology Enrichment Analysis: bap-PC

Significant GO terms (p < 0.05) with associated genes

| GO Term | Source | P-value | Genes |
| --- | --- | --- | --- |
| animal organ morphogenesis | GO:BP | 5.25e-20 | *hbs,tin,drl,tup,shg,NetB,fng,scrib,Wnt4,ct,prc,Pvf2,fz,tsh,ec,Fas2,N,ed,unc-5,GEFmeso,Lac,pnut,RhoGAP71E,cindr,emc,stg,Nhe2,Nedd4,nuf,msi,Pvr,Egfr,Ten-m,Btk,corto,pk,kibra,numb,ttk,ci,sqh,sbb,kuz,gbb,sfl,Arf6,Rab6,crol,skd,Cka,S,yrt,sog,cic,arm,Src64B,Vang,Rho1,step,Rac1,ics,siz,rdx,sgg,sano,Ype,Doa,CG42788,ftz-f1,trol* |
| cell surface receptor signaling pathway | GO:BP | 7.11e-16 | *hbs,drl,shg,apt,fng,scb,E(spl)m3-HLH,edl,Wnt4,Pvf2,fz,E(spl)malpha-BFM,tsh,Fas2,N,Ptp10D,ed,unc-5,GEFmeso,spz6,E(spl)mbeta-HLH,Sulf1,lin-28,nkd,Nedd4,E(spl)m2-BFM,Pvr,Egfr,wdp,numb,ttk,ci,Eip63E,sqh,RalGPS,kuz,ken,PRAS40,Sema2a,gbb,sfl,Pvf3,crol,skd,sog,Mob4,cic,arm,Src64B,Vang,Sema1b,CG45050,Rho1,Roc1a,CycY,step,Rac1,ics,Ubr3,rdx,sgg,Doa,trol* |
| motor neuron axon guidance | GO:BP | 1.67e-09 | *tup,NetB,Wnt4,Fas2,fend,N,Ptp10D,unc-5,Sulf1,zfh1,Ten-m,ko,Sdc,Rho1,Rac1,trol* |
| dorsal closure | GO:BP | 1.23e-08 | *tup,scb,scrib,N,ed,Mcr,emc,Pvr,Egfr,Btk,gbb,Cka,yrt,arm,Rho1,step,Rac1* |
| glial cell migration | GO:BP | 1.56e-08 | *NetB,Pvf2,N,unc-5,Pvr,numb,kuz,Pvf3,orion,Rho1,Rac1* |
| negative regulation of signal transduction | GO:BP | 3.79e-08 | *apt,fng,edl,E(spl)malpha-BFM,Fas2,Ptp10D,ed,Sulf1,nkd,Nedd4,Egfr,wdp,numb,ci,RapGAP1,sqh,scyl,chrb,ken,PRAS40,Hr4,raskol,crol,Cka,sog,Mob4,Src64B,Rho1,Roc1a,ics,rdx,sgg,trol* |
| head involution | GO:BP | 2.51e-07 | *tup,shg,tsh,Mcr,emc,Btk,scyl,chrb,yrt,Rac1* |
| ommatidial rotation | GO:BP | 6.33e-07 | *hbs,shg,fz,ec,N,Egfr,pk,S,Vang* |
| wing disc dorsal/ventral pattern formation | GO:BP | 7.06e-07 | *fng,drpr,N,Sulf1,nuf,sbb,AnxB9,tara,skd,cic,Su(Tpl)* |
| germ cell migration | GO:BP | 1.42e-06 | *tin,shg,Tre1,Hmgcr,zfh1,Fpps,mim,wun,abd-A* |
| regulation of tube length, open tracheal system | GO:BP | 3.69e-06 | *fz,Fas2,Mcr,Egfr,pasi2,sqh,Rho1,sano* |
| spiracle morphogenesis, open tracheal system | GO:BP | 6.19e-06 | *ct,Egfr,Btk,ci,sqh,Trf2,Rho1* |
| dorsal appendage formation | GO:BP | 7.61e-06 | *N,ed,Pax,emc,Egfr,ttk,Klc,S,cic,Rac1* |
| salivary gland morphogenesis | GO:BP | 1.18e-05 | *tin,drl,shg,Wnt4,Pvf2,fz,Pvr,Btk,Src64B,Rho1,Rac1* |
| segment polarity determination | GO:BP | 1.32e-05 | *fz,nkd,Egfr,ci,AGO1,sfl,arm,rdx,sgg* |
| imaginal disc-derived wing vein specification | GO:BP | 1.69e-05 | *N,GEFmeso,Egfr,corto,gbb,sfl,S,sog,cic,step* |
| cell fate commitment involved in pattern specification | GO:BP | 2.06e-05 | *tup,scrib,fz,N,nuf,Egfr,numb,abd-A* |
| larval heart development | GO:BP | 2.15e-05 | *loh,scb,Hand,prc* |
| septate junction assembly | GO:BP | 3.04e-05 | *scrib,Lac,bou,Mcr,CG44325,pasi2,crok,wun* |
| positive regulation of border follicle cell migration | GO:BP | 4.57e-05 | *shg,Pvr,Egfr,ttk,sqh,kuz,slbo,Rac1* |
| striated muscle cell differentiation | GO:BP | 9.54e-05 | *hbs,tin,Zasp52,Pax,Nedd4,tmod,mspo,Arf6,Rho1,Rac1,siz* |
| gastrulation | GO:BP | 1.06e-04 | *bap,shg,T48,RhoGAP71E,stg,sqh,sfl,sog,arm,Rho1,abd-A* |
| hemocyte migration | GO:BP | 1.15e-04 | *shg,Pvf2,Pvr,Pvf3,Rho1,Rac1* |
| positive regulation of epidermal growth factor receptor signaling pathway | GO:BP | 1.42e-04 | *edl,GEFmeso,RalGPS,Src64B,step* |
| positive regulation of neuroblast proliferation | GO:BP | 2.95e-04 | *E(spl)m3-HLH,N,E(spl)mbeta-HLH,Cka,Mob4,trol* |
| protein localization to adherens junction | GO:BP | 4.93e-04 | *Shrm,arm,Rho1* |
| formation of a compartment boundary | GO:BP | 4.93e-04 | *ct,N,sqh* |
| positive regulation of catalytic activity | GO:BP | 5.55e-04 | *hbs,drl,RhoGAP18B,scf,Pvr,Evi5,RapGAP1,rk,RalGPS,Cka,Rho1,step,Rac1,RhoGAP93B* |
| determination of digestive tract left/right asymmetry | GO:BP | 8.32e-04 | *tin,shg,CG45050,Rac1* |
| regulation of myoblast fusion | GO:BP | 8.32e-04 | *hbs,Pax,mspo,Rho1* |
| cell elongation involved in imaginal disc-derived wing morphogenesis | GO:BP | 8.32e-04 | *shg,sqh,arm,Rho1* |
| germ-line stem-cell niche homeostasis | GO:BP | 0.001 | *Wnt4,fz,N,lin-28,arm* |
| embryonic digestive tract morphogenesis | GO:BP | 0.001 | *tin,gbb,Rac1* |
| positive regulation of axon guidance | GO:BP | 0.002 | *fz,pk,Vang,Rac1* |
| dendrite guidance | GO:BP | 0.002 | *drl,tup,NetB,ct,Sema2a* |
| embryonic heart tube development | GO:BP | 0.002 | *tin,tup,Hand,Hmgcr,numb* |
| establishment of epithelial cell apical/basal polarity | GO:BP | 0.002 | *scrib,Egfr,yrt* |
| heart formation | GO:BP | 0.002 | *Nedd4,ci,arm* |
| behavioral response to ethanol | GO:BP | 0.002 | *apt,CrzR,scb,Fas2,Egfr,Arf6,S* |
| regulation of metabolic process | GO:BP | 0.002 | *hbs,tin,drl,bap,CG9650,tup,apt,bru2,E(spl)m3-HLH,edl,Hand,ct,tsh,N,Pax,E(spl)mbeta-HLH,CG9932,lin-28,luna,emc,Sox14,zfh1,Nedd4,msi,scf,Pvr,Egfr,corto,BNIP3,kibra,numb,ttk,ci,Hesr,CG12769,REPTOR-BP,tna,sbb,ko,tapas,CG42672,AGO1,Pur-alpha,ken,Glut4EF,tara,Mes2,PRAS40,CG18766,Trf2,Hr4,CG6701,CG17124,Arf6,crol,skd,Cka,slbo,CG31855,cic,arm,Su(Tpl),Rho1,Roc1a,CycY,imd,step,Rac1,Adf1,ics,CG10803,PI31,abd-A,rdx,ifc,sgg,Doa,ftz-f1,chinmo,AGO3* |
| epidermal cell differentiation | GO:BP | 0.002 | *scrib,fz,pk,sqh,Vang,Rho1,Rac1* |
| positive regulation of axon extension | GO:BP | 0.003 | *fz,pk,Vang,Rac1* |
| peripheral nervous system development | GO:BP | 0.003 | *ct,N,Egfr,ttk,Rho1,Rac1,abd-A* |
| positive regulation of stem cell proliferation | GO:BP | 0.005 | *GEFmeso,lin-28,RalGPS* |
| gonadal mesoderm development | GO:BP | 0.005 | *tin,shg,abd-A* |
| dorsal closure, spreading of leading edge cells | GO:BP | 0.005 | *Egfr,Rho1,Rac1* |
| chaeta morphogenesis | GO:BP | 0.005 | *hbs,tup,N,emc,sgg,Ype* |
| female germ-line stem cell population maintenance | GO:BP | 0.005 | *shg,fng,N,AGO1,gbb* |
| negative regulation of epidermal growth factor receptor signaling pathway | GO:BP | 0.006 | *edl,Fas2,Ptp10D,ed,Sulf1* |
| neuroblast development | GO:BP | 0.006 | *N,numb,arm,abd-A* |
| germ-band shortening | GO:BP | 0.006 | *tup,Egfr,yrt,Rac1* |
| regulation of RNA metabolic process | GO:BP | 0.006 | *tin,bap,CG9650,tup,apt,bru2,E(spl)m3-HLH,edl,Hand,ct,tsh,N,E(spl)mbeta-HLH,CG9932,lin-28,luna,emc,Sox14,zfh1,scf,kibra,ttk,ci,Hesr,CG12769,REPTOR-BP,tna,sbb,ko,tapas,AGO1,Pur-alpha,ken,Glut4EF,Mes2,CG18766,Trf2,Hr4,crol,skd,slbo,cic,arm,Su(Tpl),Adf1,abd-A,Doa,ftz-f1,chinmo,AGO3* |
| zonula adherens assembly | GO:BP | 0.006 | *shg,scrib,arm* |
| determination of genital disc primordium | GO:BP | 0.006 | *Egfr,S,abd-A* |
| homophilic cell adhesion via plasma membrane adhesion molecules | GO:BP | 0.006 | *hbs,shg,fz,Fas2,ed,Lac* |
| positive regulation of JNK cascade | GO:BP | 0.007 | *Pvr,Cka,Alg-2,Rac1,rdx* |
| somatic stem cell population maintenance | GO:BP | 0.007 | *shg,zfh1,arm,sgg* |
| axonal fasciculation | GO:BP | 0.007 | *Nrt,Fas2,zfh1,Rac1* |
| midgut development | GO:BP | 0.007 | *scb,fz,tsh,emc,abd-A* |
| negative regulation of smoothened signaling pathway | GO:BP | 0.007 | *Sulf1,Nedd4,sqh,Roc1a,rdx,sgg* |
| ventral furrow formation | GO:BP | 0.008 | *shg,T48,arm,Rho1* |
| regulation of synaptic transmission, cholinergic | GO:BP | 0.008 | *bou,crok,CG6583,CG9336* |
| semaphorin-plexin signaling pathway | GO:BP | 0.008 | *Sema2a,Sema1b,trol* |
| branch fusion, open tracheal system | GO:BP | 0.008 | *shg,ed,ttk,arm* |
| pericardial nephrocyte differentiation | GO:BP | 0.008 | *tin,numb,kuz* |
| establishment of planar polarity of embryonic epithelium | GO:BP | 0.009 | *Shrm,Rho1* |
| neuroblast fate specification | GO:BP | 0.009 | *N,nkd* |
| establishment or maintenance of actin cytoskeleton polarity | GO:BP | 0.009 | *Shrm,Rho1* |
| compartment boundary maintenance | GO:BP | 0.009 | *N,sqh* |
| positive regulation of ARF protein signal transduction | GO:BP | 0.009 | *step,siz* |
| Malpighian tubule stellate cell differentiation | GO:BP | 0.009 | *hbs,tsh* |
| regulation of actomyosin structure organization | GO:BP | 0.009 | *Nedd4,Rho1,step,mtm* |
| sensory organ precursor cell fate determination | GO:BP | 0.009 | *fz,N,nuf,numb* |
| negative regulation of locomotion | GO:BP | 0.009 | *scb,Sema2a,Sema1b,Rac1* |
| positive regulation of transcription by RNA polymerase II | GO:BP | 0.01 | *tin,bap,tup,apt,Hand,tsh,N,CG9932,Sox14,ttk,ci,CG12769,ko,skd,slbo,arm,Su(Tpl),Adf1,abd-A,ftz-f1* |
| epithelial cell proliferation involved in Malpighian tubule morphogenesis | GO:BP | 0.01 | *N,Egfr,S* |
| regulation of locomotor rhythm | GO:BP | 0.01 | *Fas2,Rho1,Rac1* |
| positive regulation of mitotic cell cycle | GO:BP | 0.011 | *N,stg,ci,crol,sgg* |
| transposable element silencing by piRNA-mediated mRNA destabilization | GO:BP | 0.016 | *tapas,AGO3* |
| negative regulation of JUN kinase activity | GO:BP | 0.016 | *ics,sgg* |
| synaptic target attraction | GO:BP | 0.016 | *NetB,Ten-m* |
| embryonic anterior midgut (ectodermal) morphogenesis | GO:BP | 0.016 | *tin,Rac1* |
| imaginal disc-derived wing hair site selection | GO:BP | 0.016 | *fz,Rac1* |
| epithelial to mesenchymal transition | GO:BP | 0.016 | *zfh1,rk* |
| maintenance of epithelial integrity, open tracheal system | GO:BP | 0.016 | *shg,Lac,Egfr* |
| axon midline choice point recognition | GO:BP | 0.016 | *drl,Nedd4,Rac1,RhoGAP93B* |
| positive regulation of neuron remodeling | GO:BP | 0.019 | *Sox14,orion,Roc1a* |
| positive regulation of wound healing | GO:BP | 0.019 | *Egfr,Rho1,Rac1* |
| positive regulation of receptor signaling pathway via JAK-STAT | GO:BP | 0.019 | *shg,lin-28,CG45050* |
| R7 cell development | GO:BP | 0.019 | *Ten-m,ttk,Rab6* |
| negative regulation of canonical Wnt signaling pathway | GO:BP | 0.019 | *Sulf1,nkd,Rho1,Roc1a,sgg* |
| protein autoubiquitination | GO:BP | 0.022 | *Rnf11,Ubr3,rdx* |
| synaptic target inhibition | GO:BP | 0.022 | *Wnt4,Sulf1,Sema2a* |
| muscle cell fate determination | GO:BP | 0.023 | *tup,N* |
| notum cell fate specification | GO:BP | 0.023 | *tup,Egfr* |
| symmetric stem cell division | GO:BP | 0.023 | *shg,lin-28* |
| juvenile hormone mediated signaling pathway | GO:BP | 0.023 | *Chd64,ftz-f1* |
| imaginal disc-derived wing vein morphogenesis | GO:BP | 0.024 | *emc,Nedd4,Egfr,gbb,sog* |
| wing disc anterior/posterior pattern formation | GO:BP | 0.026 | *fng,Sulf1,ci* |
| positive regulation of voltage-gated potassium channel activity | GO:BP | 0.026 | *crok,CG6583,CG9336* |
| tracheal outgrowth, open tracheal system | GO:BP | 0.031 | *Egfr,ttk,Rac1* |
| imaginal disc-derived male genitalia morphogenesis | GO:BP | 0.031 | *Fas2,N,Pvr* |
| positive regulation of muscle organ development | GO:BP | 0.033 | *gbb,abd-A* |
| maternal specification of dorsal/ventral axis, oocyte, soma encoded | GO:BP | 0.033 | *sog,cic* |
| amnioserosa maintenance | GO:BP | 0.033 | *tup,yrt* |
| stem cell fate commitment | GO:BP | 0.033 | *Egfr,S* |
| photoreceptor cell fate determination | GO:BP | 0.033 | *Egfr,S* |
| second mitotic wave involved in compound eye morphogenesis | GO:BP | 0.033 | *N,Egfr* |
| positive regulation of G1/S transition of mitotic cell cycle | GO:BP | 0.033 | *N,ci* |
| negative regulation of axon extension involved in axon guidance | GO:BP | 0.033 | *Sema2a,Sema1b* |
| protein localization involved in establishment of planar polarity | GO:BP | 0.033 | *Shrm,Rho1* |
| negative regulation of fibroblast growth factor receptor signaling pathway | GO:BP | 0.033 | *Ptp10D,trol* |
| ventral cord development | GO:BP | 0.034 | *tin,Pvr,kuz,18w,gbb,Rac1* |
| negative regulation of cell adhesion | GO:BP | 0.04 | *N,rk,arm* |
| mushroom body development | GO:BP | 0.04 | *drl,Fas2,Src64B,orion,ftz-f1,chinmo* |
| cell competition in a multicellular organism | GO:BP | 0.04 | *shg,drpr,Rac1* |
| chitin-based larval cuticle pattern formation | GO:BP | 0.04 | *fz,arm,sgg* |
| negative regulation of receptor signaling pathway via JAK-STAT | GO:BP | 0.04 | *apt,wdp,ken* |
| positive regulation of insulin receptor signaling pathway | GO:BP | 0.04 | *lin-28,Mob4,step* |
| negative regulation of vascular endothelial growth factor receptor signaling pathway | GO:BP | 0.042 | *Ptp10D,PRAS40* |
| muscle cell fate specification | GO:BP | 0.042 | *numb,abd-A* |
| hemocyte development | GO:BP | 0.042 | *zfh1,Rac1* |
| maintenance of epithelial cell apical/basal polarity | GO:BP | 0.042 | *Rho1,trol* |
| positive regulation of fibroblast growth factor receptor signaling pathway | GO:BP | 0.042 | *Src64B,trol* |
| dendrite self-avoidance | GO:BP | 0.044 | *Fas2,Vang,Rho1* |
| cell projection assembly | GO:BP | 0.049 | *sprt,shg,N,pnut,Pvr,Egfr,pico,mim,Unc-115a,Rho1,Rac1,mtm* |
| negative regulation of biosynthetic process | GO:BP | 0.049 | *tin,apt,bru2,edl,ct,tsh,N,E(spl)mbeta-HLH,CG9932,lin-28,emc,zfh1,msi,Egfr,corto,numb,ttk,ci,sbb,tapas,AGO1,Hr4,CG6701,crol,cic,arm,abd-A,AGO3* |
| genital disc morphogenesis | GO:BP | 0.05 | *Fas2,N,Pvr* |
| regulation of potassium ion transmembrane transport | GO:BP | 0.05 | *crok,CG6583,CG9336* |
| protein binding | GO:MF | 6.13e-14 | *hbs,CG45263,drl,Zasp52,sprt,tup,shg,apt,NetB,CG4393,Oatp74D,scb,scrib,E(spl)m3-HLH,RhoGAP18B,edl,drpr,Nrt,Wnt4,Hand,Pvf2,fz,E(spl)malpha-BFM,tsh,ec,Fas2,N,Ptp10D,Kal1,ed,Con,unc-5,GEFmeso,Lac,spz6,pnut,E(spl)mbeta-HLH,cindr,Mcr,CG17839,gukh,sick,emc,Hmgcr,nkd,Nedd4,pns,Septin2,Pvr,Egfr,Hex-A,Ten-m,wdp,Btk,corto,Evi5,pico,pk,BNIP3,kibra,pain,numb,tmod,ttk,ci,Shrm,Hesr,CG40228,Eip63E,Sh3beta,sqh,rk,REPTOR-BP,Dlc90F,GMF,CG2082,sbb,CG3402,kuz,AnxB9,Act5C,CG42672,CG11882,AGO1,Pur-alpha,ken,Mlc-c,slim,Rab4,Klc,CG18766,Trf2,18w,Sema2a,gbb,CG1888,mim,Magi,NAT1,CG11658,Snap29,RabGGTa,Pvf3,miple2,Rab6,CG9003,skd,Cka,CG42748,Unc-115a,yrt,slbo,sog,mtgo,cib,cic,Meltrin,arm,Src64B,Ppa,Vang,CG14483,Sema1b,Su(Tpl),Ca-beta,CG45050,orion,Rho1,Roc1a,CycY,imd,udd,step,Alg-2,Rac1,mtm,ics,siz,Ubr3,ReepA,PI31,abd-A,rdx,Ras64B,sgg,l(3)05822,CG42788,Roc2,Chd64,ftz-f1,chinmo,RhoGAP93B,AGO3,Svil,trol* |
| protein homodimerization activity | GO:MF | 3.51e-05 | *drl,shg,Lac,pnut,Septin2,Ten-m,corto,ttk,ci,Shrm,REPTOR-BP,Dlc90F,mim,step,rdx* |
| identical protein binding | GO:MF | 5.76e-05 | *drl,shg,Lac,pnut,Septin2,Ten-m,corto,BNIP3,ttk,ci,Shrm,REPTOR-BP,Dlc90F,Pur-alpha,CG1888,mim,Meltrin,step,rdx* |
| transcription regulator activity | GO:MF | 2.72e-04 | *tin,bap,CG9650,tup,apt,E(spl)m3-HLH,edl,Hand,ct,tsh,N,Pax,E(spl)mbeta-HLH,CG9932,luna,emc,Sox14,zfh1,ttk,ci,Hesr,CG12769,REPTOR-BP,tna,sbb,ko,Pur-alpha,ken,Glut4EF,Trf2,Hr4,crol,skd,slbo,cic,arm,Adf1,abd-A,ftz-f1* |
| DNA-binding transcription factor activity | GO:MF | 2.90e-04 | *tin,bap,CG9650,tup,apt,E(spl)m3-HLH,edl,Hand,ct,tsh,E(spl)mbeta-HLH,CG9932,luna,Sox14,zfh1,ttk,ci,Hesr,CG12769,REPTOR-BP,ko,Pur-alpha,ken,Glut4EF,Trf2,Hr4,crol,slbo,cic,Adf1,abd-A,ftz-f1* |
| transcription cis-regulatory region binding | GO:MF | 3.24e-04 | *tin,bap,CG9650,tup,apt,E(spl)m3-HLH,Hand,ct,E(spl)mbeta-HLH,CG9932,luna,Sox14,zfh1,ttk,ci,Hesr,CG12769,ko,Pur-alpha,ken,Glut4EF,Trf2,Hr4,crol,slbo,cic,Su(Tpl),Adf1,abd-A,Chd64,ftz-f1* |
| RNA polymerase II transcription regulatory region sequence-specific DNA binding | GO:MF | 3.24e-04 | *tin,bap,CG9650,tup,apt,E(spl)m3-HLH,Hand,ct,E(spl)mbeta-HLH,CG9932,luna,Sox14,zfh1,ttk,ci,Hesr,CG12769,ko,Pur-alpha,ken,Glut4EF,Trf2,Hr4,crol,slbo,cic,abd-A,Chd64,ftz-f1* |
| cytoskeletal protein binding | GO:MF | 3.24e-04 | *Zasp52,sprt,shg,pnut,gukh,Nedd4,Ten-m,tmod,Shrm,sqh,GMF,CG2082,AnxB9,Mlc-c,Klc,mim,Rab6,Unc-115a,yrt,cib,Rho1,ReepA,Chd64,Svil* |
| protein domain specific binding | GO:MF | 3.39e-04 | *N,cindr,nkd,Nedd4,pico,ttk,Sh3beta,Dlc90F,CG42672,mim,imd* |
| cell adhesion molecule binding | GO:MF | 3.39e-04 | *hbs,CG45263,scb,Nrt,Fas2,ed,cindr,CG2082,arm,siz* |
| sequence-specific double-stranded DNA binding | GO:MF | 5.16e-04 | *tin,bap,CG9650,tup,apt,E(spl)m3-HLH,Hand,ct,E(spl)mbeta-HLH,CG9932,luna,Sox14,zfh1,ttk,ci,Hesr,CG12769,ko,Pur-alpha,ken,Glut4EF,Trf2,Hr4,crol,slbo,cic,Su(Tpl),Adf1,abd-A,Chd64,ftz-f1* |
| protein dimerization activity | GO:MF | 5.16e-04 | *drl,shg,scb,E(spl)m3-HLH,Hand,Lac,pnut,E(spl)mbeta-HLH,emc,Septin2,Ten-m,corto,ttk,ci,Shrm,Hesr,REPTOR-BP,Dlc90F,gbb,mim,step,rdx,ftz-f1* |
| growth factor activity | GO:MF | 5.16e-04 | *Pvf2,spz6,gbb,Pvf3,miple2,sog* |
| DNA-binding transcription factor activity, RNA polymerase II-specific | GO:MF | 0.001 | *tin,bap,tup,apt,E(spl)m3-HLH,edl,Hand,ct,tsh,E(spl)mbeta-HLH,CG9932,luna,Sox14,zfh1,ttk,ci,Hesr,CG12769,ko,Pur-alpha,Hr4,crol,slbo,cic,Adf1,abd-A,ftz-f1* |
| sequence-specific DNA binding | GO:MF | 0.001 | *tin,bap,CG9650,tup,apt,E(spl)m3-HLH,edl,Hand,ct,E(spl)mbeta-HLH,CG9932,luna,Sox14,zfh1,ttk,ci,Hesr,CG12769,ko,Pur-alpha,ken,Glut4EF,Trf2,Hr4,crol,slbo,cic,Su(Tpl),Adf1,abd-A,Chd64,ftz-f1* |
| actin binding | GO:MF | 0.002 | *Zasp52,sprt,pnut,gukh,Shrm,GMF,mim,Rab6,Unc-115a,cib,Rho1,Chd64,Svil* |
| double-stranded DNA binding | GO:MF | 0.002 | *tin,bap,CG9650,tup,apt,E(spl)m3-HLH,Hand,ct,E(spl)mbeta-HLH,CG9932,luna,Sox14,zfh1,ttk,ci,Hesr,CG12769,ko,Pur-alpha,ken,Glut4EF,Trf2,Hr4,crol,slbo,cic,Su(Tpl),Adf1,abd-A,Chd64,ftz-f1* |
| cis-regulatory region sequence-specific DNA binding | GO:MF | 0.002 | *tin,bap,CG9650,tup,E(spl)m3-HLH,E(spl)mbeta-HLH,CG9932,luna,Sox14,zfh1,ttk,ci,Hesr,CG12769,ken,Glut4EF,Hr4,crol,slbo,Su(Tpl),abd-A,Chd64,ftz-f1* |
| RNA polymerase II cis-regulatory region sequence-specific DNA binding | GO:MF | 0.004 | *tin,bap,CG9650,tup,E(spl)m3-HLH,E(spl)mbeta-HLH,CG9932,luna,Sox14,zfh1,ttk,ci,Hesr,CG12769,ken,Glut4EF,Hr4,crol,slbo,abd-A,Chd64,ftz-f1* |
| enzyme binding | GO:MF | 0.004 | *drl,RhoGAP18B,E(spl)malpha-BFM,ec,N,GEFmeso,pnut,pns,Evi5,pico,kibra,ci,Shrm,CG40228,AGO1,Magi,Cka,CG42748,arm,Su(Tpl),Rho1,CycY,Rac1,rdx* |
| receptor ligand activity | GO:MF | 0.005 | *Wnt4,Pvf2,spz6,Sema2a,gbb,Pvf3,miple2,sog,Sema1b,orion* |
| signaling receptor activator activity | GO:MF | 0.007 | *Wnt4,Pvf2,spz6,Sema2a,gbb,Pvf3,miple2,sog,Sema1b,orion* |
| cytokine receptor binding | GO:MF | 0.01 | *Pvf2,gbb,Pvf3,orion* |
| GTPase activity | GO:MF | 0.014 | *pnut,Septin4,Septin2,Rab4,Arf6,Rab6,CG2017,Rho1,Non1,Rac1,Ras64B* |
| signaling receptor binding | GO:MF | 0.019 | *scb,Wnt4,Pvf2,ed,spz6,Nedd4,numb,Sema2a,gbb,Pvf3,miple2,sog,Src64B,Vang,Sema1b,orion* |
| kinase binding | GO:MF | 0.021 | *drl,kibra,ci,Shrm,Cka,arm,Rho1,CycY,Rac1* |
| phospholipid binding | GO:MF | 0.027 | *sprt,drpr,AnxB10,crok,AnxB9,mim,AnxB11,CG6583,step,CG9336,Svil* |
| GTPase regulator activity | GO:MF | 0.041 | *RhoGAP18B,GEFmeso,RhoGAP71E,pns,Evi5,RapGAP1,RalGPS,raskol,step,siz,RhoGAP93B* |
| cadherin binding | GO:MF | 0.041 | *CG2082,arm,siz* |
| NEDD8 ligase activity | GO:MF | 0.041 | *Roc1a,Roc2* |
| oxidoreductase activity, acting on a sulfur group of donors, disulfide as acceptor | GO:MF | 0.041 | *GILT1,GILT2,GILT3* |
| vascular endothelial growth factor receptor binding | GO:MF | 0.041 | *Pvf2,Pvf3* |
| calcium-dependent phospholipid binding | GO:MF | 0.041 | *AnxB10,AnxB9,AnxB11* |
| GPI anchor binding | GO:MF | 0.048 | *crok,CG6583,CG9336* |

### 1.5. Gene Ontology Enrichment Analysis: eve-PC

Significant GO terms (p < 0.05) with associated genes

| GO Term | Source | P-value | Genes |
| --- | --- | --- | --- |
| animal organ morphogenesis | GO:BP | 3.33e-15 | *tin,tinc,rho,Lac,shg,tup,vn,Timp,esg,prc,sinu,eve,pnut,ptc,dally,cdi,mbc,app,fz2,robo2,Myo31DF,Cont,corto,Grip,alphaSnap,cora,RhoGAP15B,Rab6,sqh,cv-2,kibra,p120ctn,ics,aPKC,emc,Ras85D,Nhe2,Rac1,Nrx-IV,Nedd4,dlg1,sano,Rip11,RhoGEF2,pyd,yrt,Septin1,Rap1,Src64B,Arf1,sbb,lolal,14-3-3epsilon,Ggamma1,SCAR,kuz,Cyfip,Rho1,M6,cindr,stck,gbb,Hem,Fhos,klar,Ced-12,faf,p38a,parvin,spg,dnr1,brk,trol,Gbeta13F,Jra,Dif,ftz-f1,Shc,sfl,tsh* |
| septate junction assembly | GO:BP | 2.04e-14 | *wun,Lac,cold,CG44325,pasi2,crok,sinu,Mcr,pck,CG9628,crim,kune,pasi1,Cont,loco,Nrx-IV,dlg1,Tsf2* |
| dorsal closure | GO:BP | 9.99e-10 | *scb,rho,tup,sinu,Mcr,mbc,alpha-Cat,cora,emc,Ras85D,Rac1,Nrx-IV,dlg1,pyd,yrt,Rap1,Ggamma1,Rho1,stck,gbb,Jra,Sac1* |
| cell surface receptor signaling pathway | GO:BP | 2.08e-09 | *Egfrap,scb,rho,shg,vn,apt,kek1,Sema5c,ptc,RalGPS,Prosap,dally,cdi,mbc,fz2,wdp,robo2,ken,Ptp4E,sqh,cv-2,ics,aPKC,Akap200,Ras85D,Ilp6,UbcE2M,spz,Roc1a,Rac1,Nedd4,dlg1,ckn,Cirl,Eip63E,Sirt1,Rap1,Mob4,Src64B,Ehbp1,gcl,for,spz6,PRL-1,kuz,CycY,Rho1,stck,gbb,Ced-12,faf,mth,spg,brk,Pi3K68D,par-6,trol,aph-1,CG11155,Gbeta13F,CG11523,Sac1,Dif,Shc,Socs44A,sfl,tsh* |
| establishment of glial blood-brain barrier | GO:BP | 3.97e-08 | *pasi2,sinu,pck,kune,pasi1,Cont,loco,cora,Nrx-IV* |
| cell adhesion involved in heart morphogenesis | GO:BP | 5.02e-08 | *Lac,prc,sinu,Cont,cora,Nrx-IV,Ggamma1,Gbeta13F* |
| striated muscle cell differentiation | GO:BP | 4.91e-07 | *tin,kon,rols,eve,mspo,DAAM,Pax,mbc,Pak3,Tm2,Rac1,Nedd4,Zasp52,SCAR,Rho1,Hem,Ced-12* |
| motor neuron axon guidance | GO:BP | 3.31e-06 | *beat-IIa,tup,fend,eve,dally,fz2,beat-IIb,Ptp4E,zfh1,spz,Rac1,for,Rho1,Sdc,trol* |
| cortical actin cytoskeleton organization | GO:BP | 4.09e-06 | *Fim,loco,Rac1,vib,mtm,RhoGEF2,Abi,SCAR,Cyfip,Rho1,Hem,Fhos,Pi3K68D* |
| positive regulation of filopodium assembly | GO:BP | 4.58e-06 | *kon,DAAM,pico,aPKC,Rac1,Abi,SCAR,Cyfip,par-6* |
| negative regulation of signal transduction | GO:BP | 1.05e-05 | *Egfrap,apt,kek1,ptc,Prosap,dally,cdi,wdp,ken,Ptp4E,sqh,ics,aPKC,Spred,Ras85D,UbcE2M,Roc1a,Nedd4,dlg1,Ufd4,Rap1,Mob4,Src64B,gcl,PRL-1,RapGAP1,Rho1,stck,Hr4,dnr1,brk,Pi3K68D,trol,Gbeta13F,CG17985,Sac1,Socs44A* |
| germ cell migration | GO:BP | 4.93e-05 | *tin,wun2,wun,shg,Tre1,trx,zfh1,abd-A,faf* |
| head involution | GO:BP | 1.00e-04 | *shg,tup,Mcr,emc,Rac1,pyd,yrt,stck,tsh* |
| larval heart development | GO:BP | 1.36e-04 | *scb,loh,Hand,prc* |
| positive regulation of voltage-gated potassium channel activity | GO:BP | 2.16e-04 | *crok,CG31676,crim,CG6583,CG9336,CG9338* |
| biological process involved in interspecies interaction between organisms | GO:BP | 2.82e-04 | *wun,shg,GILT1,Mcr,drpr,18w,loco,Rab6,aPKC,Rab14,Vps60,Ras85D,UbcE2M,spz,NUCB1,Rac1,Rab7,CG13551,dlg1,CG5390,Src64B,Vps29,Arf1,spz6,RNASEK,Rho1,CG12744,CG4968,Atg18a,faf,p38a,dnr1,AP-1-2beta,CG40160,Jra,Sec10,CG17985,Dif,TSG101,Shc,Hsp27* |
| viral process | GO:BP | 3.94e-04 | *Rab14,Vps60,Roc1a,Rab7,Vps29,AP-1-2beta,Jra,Sec10,TSG101* |
| cell competition in a multicellular organism | GO:BP | 5.92e-04 | *shg,mbc,drpr,spz,Rac1,brk* |
| regulation of tube length, open tracheal system | GO:BP | 7.11e-04 | *pasi2,Mcr,kune,pasi1,sqh,sano,Rho1* |
| regulation of potassium ion transmembrane transport | GO:BP | 0.001 | *crok,CG31676,crim,CG6583,CG9336,CG9338* |
| regulation of synaptic transmission, cholinergic | GO:BP | 0.001 | *crok,CG31676,crim,CG6583,CG9336,CG9338* |
| gastrulation | GO:BP | 0.001 | *shg,eve,T48,sqh,Ras85D,RhoGEF2,abd-A,Ggamma1,Rho1,Gbeta13F,ric8a,sfl* |
| symbiont entry into host cell | GO:BP | 0.001 | *Rab14,Vps60,Rab7,Vps29,AP-1-2beta,Sec10,TSG101* |
| protein refolding | GO:BP | 0.001 | *CG14207,Hsp70Ab,Hsp70Bc,Hsp68,Hsp27,Hsp23,Hsp26* |
| midgut development | GO:BP | 0.002 | *scb,vn,fz2,emc,Gp93,abd-A,tsh* |
| tracheal outgrowth, open tracheal system | GO:BP | 0.003 | *Timp,Ras85D,Rac1,lolal,Shc* |
| apical protein localization | GO:BP | 0.004 | *shg,aPKC,Ggamma1,par-6,Gbeta13F* |
| regulation of monoatomic ion transmembrane transport | GO:BP | 0.004 | *crok,CG31676,crim,CG6583,CG9336,CG9338,Cnx99A,Nedd4,Ca-Ma2d* |
| embryonic digestive tract morphogenesis | GO:BP | 0.004 | *tin,Rac1,gbb* |
| cell elongation involved in imaginal disc-derived wing morphogenesis | GO:BP | 0.004 | *shg,sqh,RhoGEF2,Rho1* |
| regulation of myoblast fusion | GO:BP | 0.004 | *eve,mspo,Pax,Rho1* |
| determination of digestive tract left/right asymmetry | GO:BP | 0.004 | *tin,shg,Myo31DF,Rac1* |
| positive regulation of border follicle cell migration | GO:BP | 0.005 | *shg,vn,mbc,sqh,slbo,Rac1,kuz* |
| chaperone cofactor-dependent protein refolding | GO:BP | 0.006 | *mrj,DnaJ-1,jdp,Hsp70Ab,Hsp70Bc,Hsp68* |
| maintenance of epithelial integrity, open tracheal system | GO:BP | 0.008 | *rho,Lac,shg,M6* |
| hemocyte migration | GO:BP | 0.008 | *shg,Ras85D,Rac1,Rap1,Rho1* |
| 'de novo' protein folding | GO:BP | 0.009 | *mrj,DnaJ-1,jdp,Hsp70Ab,Hsp70Bc,Hsp68* |
| trans-synaptic signaling | GO:BP | 0.01 | *apt,crok,CG31676,crim,CG6583,be,CG9336,alphaSnap,Snap25,beta-Spec,CG9338,Snap29,dlg1,Rap1,Src64B,Arf1,scramb2,Blos1,Ca-beta,gbb,mth,CG11155,Sec10,ric8a* |
| positive regulation of substrate adhesion-dependent cell spreading | GO:BP | 0.01 | *stck,Fhos* |
| germ cell repulsion | GO:BP | 0.01 | *wun2,wun* |
| positive regulation of sarcomere organization | GO:BP | 0.01 | *DAAM,Tm2* |
| cell projection assembly | GO:BP | 0.01 | *kon,shg,vn,pnut,DAAM,sprt,pico,aPKC,Rac1,mtm,Unc-115a,Rap1,Abi,SCAR,Cyfip,Rho1,Hem,parvin,par-6* |
| positive regulation of neuron remodeling | GO:BP | 0.01 | *Sox14,Roc1a,orion,Arf1* |
| vacuole organization | GO:BP | 0.011 | *Rab14,Rab7,Atg17,Svip,Atg3,Atg8a,Atg18a,Lamtor1* |
| positive regulation of fibroblast growth factor receptor signaling pathway | GO:BP | 0.011 | *dally,Src64B,trol* |
| regulation of myosin II filament organization | GO:BP | 0.011 | *RhoGEF2,Ggamma1,Gbeta13F* |
| embryonic heart tube development | GO:BP | 0.012 | *tin,tup,Hand,robo2,Ggamma1* |
| cell adhesion mediated by integrin | GO:BP | 0.012 | *scb,Abi,SCAR,Cyfip,Hem* |
| imaginal disc-derived wing vein morphogenesis | GO:BP | 0.012 | *rho,vn,dally,emc,Ras85D,Nedd4,gbb* |
| cellular response to growth factor stimulus | GO:BP | 0.014 | *dally,Ptp4E,cv-2,Ras85D,Src64B,PRL-1,gbb,faf,brk,trol,Shc,sfl* |
| leg disc proximal/distal pattern formation | GO:BP | 0.016 | *rho,vn,Ras85D,tsh* |
| dorsal closure, spreading of leading edge cells | GO:BP | 0.016 | *Ras85D,Rac1,Rho1* |
| negative regulation of smoothened signaling pathway | GO:BP | 0.016 | *ptc,sqh,UbcE2M,Roc1a,Nedd4,Gbeta13F,Sac1* |
| axon midline choice point recognition | GO:BP | 0.016 | *robo2,beta-Spec,Rac1,Nedd4,Sac1* |
| gonadal mesoderm development | GO:BP | 0.016 | *tin,shg,abd-A* |
| positive regulation of sevenless signaling pathway | GO:BP | 0.016 | *Rap1,Src64B,spg* |
| wing disc dorsal/ventral pattern formation | GO:BP | 0.019 | *dally,drpr,Su(Tpl),Rab7,AnxB9,sbb,14-3-3epsilon* |
| positive regulation of TORC1 signaling | GO:BP | 0.021 | *Ras85D,Lamtor4,Lamtor1,RagC-D,Shc* |
| behavioral response to ethanol | GO:BP | 0.021 | *scb,rho,apt,CrzR,Akap200,dlg1,Sirt1* |
| zonula adherens assembly | GO:BP | 0.022 | *shg,aPKC,par-6* |
| positive regulation of lamellipodium assembly | GO:BP | 0.022 | *aPKC,SCAR,par-6* |
| determination of genital disc primordium | GO:BP | 0.022 | *rho,ptc,abd-A* |
| positive regulation of catalytic activity | GO:BP | 0.023 | *RhoGAP18B,RalGPS,mbc,RhoGAP15B,Evi5,Rac1,Atg17,Abi,RapGAP1,Rho1,Ced-12,spg,aph-1,CG4853* |
| actin filament network formation | GO:BP | 0.023 | *Fim,Fhos* |
| nerve maturation | GO:BP | 0.023 | *Cont,Nrx-IV* |
| determination of adult lifespan | GO:BP | 0.027 | *CG42663,miple2,Ras85D,NLaz,Indy,Sirt1,VhaSFD,14-3-3epsilon,Atg8a,p38a,mth,Hsp68,Hsp27,Hsp26* |
| negative regulation of receptor signaling pathway via JAK-STAT | GO:BP | 0.027 | *apt,wdp,ken,Socs44A* |
| regulation of epithelial cell migration, open tracheal system | GO:BP | 0.028 | *ptc,robo2,Pu* |
| semaphorin-plexin signaling pathway | GO:BP | 0.028 | *Sema5c,dally,trol* |
| membrane protein ectodomain proteolysis | GO:BP | 0.028 | *Timp,kuz,aph-1* |
| synaptic vesicle fusion to presynaptic active zone membrane | GO:BP | 0.028 | *alphaSnap,Snap25,Snap29* |
| dorsal closure, amnioserosa morphology change | GO:BP | 0.028 | *alpha-Cat,Rac1,Sac1* |
| tracheal pit formation in open tracheal system | GO:BP | 0.028 | *rho,Timp,Rho1* |
| ventral cord development | GO:BP | 0.03 | *tin,Timp,robo2,18w,loco,Rac1,kuz,gbb* |
| imaginal disc-derived wing vein specification | GO:BP | 0.031 | *rho,corto,cv-2,Ras85D,gbb,Shc,sfl* |
| germ-band extension | GO:BP | 0.036 | *eve,RhoGEF2,Rho1* |
| glycophagy | GO:BP | 0.036 | *Atg17,Atg3,Atg18a* |
| positive regulation of JNK cascade | GO:BP | 0.037 | *mbc,Alg-2,Rac1,Ced-12,ALiX* |
| autophagy of mitochondrion | GO:BP | 0.037 | *Fis1,Atg17,Atg3,Atg8a,Atg18a* |
| preblastoderm mitotic cell cycle | GO:BP | 0.039 | *luna,Pu* |
| regulation of basement membrane organization | GO:BP | 0.039 | *Ndg,Ehbp1* |
| establishment or maintenance of cytoskeleton polarity involved in gastrulation | GO:BP | 0.039 | *Ggamma1,Gbeta13F* |
| epithelial to mesenchymal transition | GO:BP | 0.039 | *esg,zfh1* |
| muscle cell development | GO:BP | 0.039 | *kon,rols,DAAM,Tm2,Rac1,Nedd4,Zasp52* |
| posterior midgut invagination | GO:BP | 0.039 | *RhoGEF2,Rho1* |
| embryonic anterior midgut (ectodermal) morphogenesis | GO:BP | 0.039 | *tin,Rac1* |
| outflow tract morphogenesis | GO:BP | 0.039 | *shg,robo2* |
| positive phototaxis | GO:BP | 0.039 | *dlg1,M6* |
| activation of GTPase activity | GO:BP | 0.04 | *mbc,Evi5,RapGAP1,Ced-12,spg* |
| ectopic germ cell programmed cell death | GO:BP | 0.042 | *wun2,wun,Tre1* |
| neuron migration | GO:BP | 0.042 | *robo2,SCAR,Hem* |
| negative regulation of catalytic activity | GO:BP | 0.042 | *kek1,Timp,ics,Spn77Bc,endos,Spn31A,par-6* |
| cell fate commitment involved in pattern specification | GO:BP | 0.043 | *rho,tup,vn,dlg1,abd-A* |
| regulation of circadian sleep/wake cycle, sleep | GO:BP | 0.044 | *crok,CG31676,crim,CG6583,CG9336,CG9338* |
| positive regulation of ERK1 and ERK2 cascade | GO:BP | 0.045 | *vn,Ras85D,Rap1,spg* |
| positive regulation of hippo signaling | GO:BP | 0.048 | *ptc,kibra,pyd,Src64B,14-3-3epsilon* |
| cellular response to amino acid stimulus | GO:BP | 0.05 | *Lamtor4,Lamtor1,RagC-D* |
| protein binding | GO:MF | 3.84e-16 | *Egfrap,beat-IIa,kon,scb,Lac,shg,rols,RabGGTa,tup,CG42709,vn,apt,pain,RhoGAP18B,kek1,Timp,Hand,sinu,tei,Mcr,pnut,Septin2,DAAM,Sema5c,ptc,sprt,CG32354,Prosap,dally,Lrch,mbc,Act42A,Kal1,fz2,Pur-alpha,wdp,robo2,ken,drpr,Myo31DF,Cont,CG40228,Fim,jvl,corto,beat-IIb,Ptp4E,Wdr62,Grip,18w,GMF,Alg-2,CG14535,pns,alpha-Cat,pico,alphaSnap,loco,cora,Rab6,sqh,cib,Skp2,cv-2,Sh3beta,kibra,miple2,p120ctn,ics,trx,aPKC,Mlc-c,slbo,Akap200,CG3402,emc,CG30423,Pak3,CG42748,Spred,Gp93,CG11658,Ras85D,Snap25,Tm2,CG13252,beta-Spec,Ilp6,Su(Tpl),spz,Evi5,Roc1a,CG14207,Drep1,mrj,RhoGDI,Rac1,Snap29,Cnx99A,CG5742,CG11882,Fis1,CG13551,Nedd4,dlg1,ckn,Pat1,mtm,NAT1,orion,Ufd4,Eip63E,cuff,Rip11,Sirt1,RhoGEF2,Atg17,Unc-115a,pyd,yrt,Septin1,Rap1,Src64B,Ppa,Ehbp1,AnxB9,Zasp52,sbb,CG12868,lolal,abd-A,DnaJ-1,gcl,stau,14-3-3epsilon,spz6,EndoB,CG5721,CG17839,Abi,PRL-1,Atg8a,Spec2,jdp,Con,Sap30,Act5C,Ras64B,Arpc5,Ggamma1,SCAR,kuz,CycY,slim,Pde11,Cyfip,Vinc,Rho1,Blos1,Ca-beta,Rab4,His3.3B,CycK,cindr,stck,Klc,Cln7,Cortactin,CG4968,Atg18a,Hsp70Ab,sowah,gbb,Hem,Fhos,CG17754,klar,AGO1,Vps28,Gbeta5,Ced-12,faf,mth,parvin,chinmo,spg,CG14894,CG43102,CG14995,brk,AP-1-2beta,Pi3K68D,par-6,a,trol,Sou,CG9426,CG16952,CG5708,Gbeta13F,CG11523,ALiX,Setd3,His4r,Jra,Sec10,Cmtr1,Syx5,ric8a,CG6607,RagC-D,Dif,TSG101,TfIIEbeta,CG5445,fax,ftz-f1,Shc,Hsp70Bc,Hsp68,Hsp27,tsh,Hsp23,Hsp26* |
| actin binding | GO:MF | 6.09e-08 | *pnut,DAAM,sprt,Myo31DF,Fim,GMF,alpha-Cat,cora,Rab6,cib,Akap200,Tm2,beta-Spec,Unc-115a,Zasp52,Arpc5,SCAR,Vinc,Rho1,Cortactin,Fhos,CG17754,parvin,CG9426,Setd3* |
| phospholipid binding | GO:MF | 4.43e-06 | *crok,CG31676,crim,ptc,sprt,CG6583,drpr,Myo31DF,AnxB10,CG9336,RhoGAP15B,AnxB11,beta-Spec,CG9338,vib,AnxB9,EndoB,Snx21,Atg18a,CG8176,Pi3K68D,CG30392,CG1902* |
| cytoskeletal protein binding | GO:MF | 6.79e-06 | *shg,pnut,DAAM,sprt,Myo31DF,Fim,GMF,CG14535,alpha-Cat,cora,Rab6,sqh,cib,aPKC,Mlc-c,Akap200,Tm2,beta-Spec,Nedd4,Unc-115a,yrt,AnxB9,Zasp52,Arpc5,SCAR,Vinc,Rho1,Klc,Cortactin,Fhos,CG17754,klar,parvin,CG9426,Setd3,Hsp26* |
| GPI anchor binding | GO:MF | 5.23e-04 | *crok,CG31676,crim,CG6583,CG9336,CG9338* |
| phosphatidylinositol binding | GO:MF | 0.001 | *crok,CG31676,crim,ptc,CG6583,Myo31DF,CG9336,RhoGAP15B,beta-Spec,CG9338,vib,Snx21,Atg18a,Pi3K68D,CG1902* |
| enzyme binding | GO:MF | 0.002 | *Egfrap,RhoGAP18B,Timp,pnut,DAAM,mbc,CG40228,pns,pico,kibra,trx,Pak3,CG42748,Spred,Su(Tpl),Evi5,Rac1,CG13551,dlg1,ckn,Rip11,Atg17,Septin1,Rap1,Atg8a,Spec2,CycY,Cyfip,Rho1,AGO1,spg,Pi3K68D,par-6,CG6607,Shc* |
| enzyme regulator activity | GO:MF | 0.002 | *RhoGAP18B,Timp,Mcr,RalGPS,mbc,Kal1,pns,loco,RhoGAP15B,Pu,Ras85D,CG17272,Evi5,Drep1,Spn77Bc,RhoGDI,Rac1,CG5390,endos,RhoGEF2,Atg17,Lamtor4,RapGAP1,Spn31A,CycY,CycK,Ced-12,spg,CG43102,par-6,CG40160,Lamtor1,CG4853,CG11523,ric8a,Oda,Socs44A* |
| unfolded protein binding | GO:MF | 0.002 | *Gp93,CG14207,mrj,Cnx99A,DnaJ-1,jdp,Hsp70Ab,Hsp70Bc,Hsp68,Hsp27,Hsp23,Hsp26* |
| lipid binding | GO:MF | 0.002 | *crok,CG31676,crim,ptc,sprt,CG6583,drpr,Myo31DF,AnxB10,CG9336,RhoGAP15B,AnxB11,beta-Spec,CG9338,vib,AnxB9,EndoB,Snx21,Atg18a,CG8176,Pi3K68D,CG30392,CG1902* |
| molecular adaptor activity | GO:MF | 0.003 | *pnut,Septin2,Pax,Prosap,AP-2sigma,alphaSnap,kibra,emc,Snap25,Snap29,ckn,Pat1,Sirt1,Atg17,Septin1,sbb,Septin4,gcl,Lamtor4,Sap30,tna,Gbeta5,brk,AP-1-2beta,aph-1,Lamtor1,CG5708,Gbeta13F,Setd3,Syx5* |
| GTPase activity | GO:MF | 0.005 | *pnut,Septin2,Rab6,Rab14,Ras85D,Rac1,Rab7,CG2017,Septin1,Rap1,Arf1,Septin4,Ras64B,Rho1,Rab4,RagC-D* |
| G protein activity | GO:MF | 0.008 | *Ras85D,Rap1,Ras64B,Rab4* |
| guanyl nucleotide binding | GO:MF | 0.009 | *pnut,Septin2,Rab6,Rab14,Pu,Ras85D,Rac1,Rab7,CG2017,Septin1,Rap1,Arf1,Septin4,for,Ras64B,Rho1,Rab4,RagC-D* |
| signaling receptor binding | GO:MF | 0.009 | *Egfrap,scb,vn,kek1,Sema5c,ptc,Prosap,Grip,cv-2,miple2,Akap200,CG30423,Ilp6,spz,CG5742,Nedd4,orion,RhoGEF2,Src64B,spz6,Atg8a,gbb,Shc* |
| GTPase regulator activity | GO:MF | 0.009 | *RhoGAP18B,RalGPS,mbc,pns,loco,RhoGAP15B,Evi5,RhoGDI,RhoGEF2,Lamtor4,RapGAP1,Ced-12,spg,CG43102,Lamtor1,CG4853,ric8a* |
| guanyl ribonucleotide binding | GO:MF | 0.009 | *pnut,Septin2,Rab6,Rab14,Pu,Ras85D,Rac1,Rab7,CG2017,Septin1,Rap1,Arf1,Septin4,for,Ras64B,Rho1,Rab4,RagC-D* |
| guanyl-nucleotide exchange factor activity | GO:MF | 0.01 | *RalGPS,mbc,pns,RhoGEF2,Lamtor4,Ced-12,spg,CG43102,Lamtor1,CG4853,ric8a* |
| actin filament binding | GO:MF | 0.011 | *Myo31DF,Fim,alpha-Cat,Akap200,Tm2,beta-Spec,Unc-115a,Arpc5,Vinc,Cortactin,Fhos* |
| SH3 domain binding | GO:MF | 0.014 | *Sh3beta,Abi,cindr,Ced-12,Pi3K68D* |
| GTP binding | GO:MF | 0.014 | *pnut,Septin2,Rab6,Rab14,Pu,Ras85D,Rac1,Rab7,CG2017,Septin1,Rap1,Arf1,Septin4,Ras64B,Rho1,Rab4,RagC-D* |
| small GTPase binding | GO:MF | 0.016 | *RhoGAP18B,DAAM,mbc,pns,Pak3,Evi5,Rip11,Spec2,Cyfip,spg,Pi3K68D,CG6607* |
| beta-catenin binding | GO:MF | 0.018 | *shg,alpha-Cat,Vinc,Sec10* |
| GTPase binding | GO:MF | 0.018 | *RhoGAP18B,DAAM,mbc,pns,Pak3,Evi5,Rip11,Spec2,Cyfip,spg,Pi3K68D,CG6607* |
| kinase binding | GO:MF | 0.018 | *Egfrap,kibra,Spred,Rac1,dlg1,Atg17,Rap1,Atg8a,CycY,Rho1,par-6,Shc* |
| cell adhesion molecule binding | GO:MF | 0.029 | *scb,rols,tei,Cont,alpha-Cat,p120ctn,pyd,Abi,cindr* |
| growth factor activity | GO:MF | 0.034 | *vn,miple2,spz,spz6,gbb* |
| protein homodimerization activity | GO:MF | 0.04 | *Lac,shg,pnut,Septin2,robo2,corto,trx,Pak3,spz,Septin1,lolal,Syx5* |
| protein kinase binding | GO:MF | 0.042 | *Egfrap,Spred,Rac1,Atg17,Rap1,Atg8a,CycY,Rho1,par-6,Shc* |

### 1.6. Gene Ontology Enrichment Analysis: dying eve-PC

Significant GO terms (p < 0.05) with associated genes

| GO Term | Source | P-value | Genes |
| --- | --- | --- | --- |
| animal organ morphogenesis | GO:BP | 2.70e-16 | *prc,Timp,robo2,cv-2,trol,klar,ftz-f1,shg,tup,Grip,cdi,pyd,dally,tin,tinc,dnr1,rdx,sbb,app,Fhos,Lac,Nrg,fz2,kuz,par-1,pk,kibra,nmo,hid,ptc,how,mbc,scrib,corto,Dys,Mob2,bun,spen,dlg1,rho,cindr,lola,shot,mam,stg* |
| cell surface receptor signaling pathway | GO:BP | 6.78e-08 | *apt,robo2,cv-2,Sema5c,trol,shg,Egfrap,scb,cdi,dally,nkd,rdx,Prosap,fz2,Dyrk2,kuz,par-1,ken,kek1,nmo,RalGPS,ptc,mbc,Eip63E,spen,dlg1,hppy,rho,HDAC4,mam,wdp,Ptp10D* |
| striated muscle cell differentiation | GO:BP | 3.36e-07 | *kon,mspo,rols,tin,DAAM,Pax,how,mbc,Zasp52,Tm2,hts* |
| septate junction assembly | GO:BP | 4.16e-07 | *CG44325,wun,Lac,Nrg,CG9628,scrib,dlg1,Mcr* |
| negative regulation of signal transduction | GO:BP | 1.26e-06 | *apt,trol,Egfrap,cdi,dally,dnr1,nkd,rdx,Prosap,par-1,ken,kek1,nmo,ptc,dlg1,hppy,RapGAP1,scyl,wdp,rdgA,Ptp10D* |
| larval heart development | GO:BP | 2.40e-06 | *loh,prc,scb,Hand* |
| motor neuron axon guidance | GO:BP | 1.16e-05 | *beat-IIa,trol,tup,dally,fend,Nrg,fz2,Sdc,Ptp10D* |
| muscle cell development | GO:BP | 1.76e-05 | *kon,rols,DAAM,how,Zasp52,Dys,Tm2,hts* |
| wing disc dorsal/ventral pattern formation | GO:BP | 5.81e-05 | *dally,sbb,nmo,AnxB9,drpr,Su(Tpl),CG14073* |
| sarcomere organization | GO:BP | 6.66e-05 | *kon,rols,DAAM,how,Tm2,hts* |
| dorsal closure | GO:BP | 8.40e-05 | *tup,scb,pyd,mbc,scrib,dlg1,rho,shot,Mcr* |
| head involution | GO:BP | 8.75e-05 | *shg,tup,pyd,hid,Mcr,scyl* |
| receptor clustering | GO:BP | 1.95e-04 | *Grip,scrib,Fur1,dlg1* |
| segment polarity determination | GO:BP | 2.23e-04 | *dally,nkd,rdx,fz2,ptc,AGO1* |
| mushroom body development | GO:BP | 5.24e-04 | *robo2,chinmo,ftz-f1,DAAM,Nrg,bun,shot* |
| germ cell migration | GO:BP | 6.95e-04 | *shg,wun,wun2,tin,trx* |
| branch fusion, open tracheal system | GO:BP | 0.001 | *shg,pyd,form3,shot* |
| positive regulation of border follicle cell migration | GO:BP | 0.001 | *shg,tai,kuz,par-1,mbc* |
| germ cell repulsion | GO:BP | 0.001 | *wun,wun2* |
| neurotransmitter receptor transport postsynaptic membrane to endosome | GO:BP | 0.001 | *Grip,scrib* |
| positive regulation of sarcomere organization | GO:BP | 0.001 | *DAAM,Tm2* |
| neurotransmitter receptor transport, endosome to postsynaptic membrane | GO:BP | 0.001 | *Grip,scrib* |
| central complex development | GO:BP | 0.001 | *robo2,Sema5c,Nrg* |
| embryonic heart tube development | GO:BP | 0.002 | *robo2,tup,tin,Hand* |
| semaphorin-plexin signaling pathway | GO:BP | 0.002 | *Sema5c,trol,dally* |
| establishment of imaginal disc-derived wing hair orientation | GO:BP | 0.002 | *app,par-1,pk,scrib* |
| cell adhesion involved in heart morphogenesis | GO:BP | 0.003 | *prc,Lac,Nrg* |
| pole plasm protein localization | GO:BP | 0.003 | *par-1,scrib,dlg1* |
| anterior/posterior axis specification, follicular epithelium | GO:BP | 0.004 | *scrib,dlg1* |
| actin filament network formation | GO:BP | 0.004 | *Fhos,Fim* |
| maintenance of epithelial integrity, open tracheal system | GO:BP | 0.004 | *shg,Lac,rho* |
| behavioral response to ethanol | GO:BP | 0.004 | *apt,scb,dlg1,hppy,rho* |
| epidermal cell differentiation | GO:BP | 0.005 | *app,par-1,pk,scrib,spen* |
| imaginal disc-derived wing vein specification | GO:BP | 0.006 | *cv-2,nmo,corto,Dys,rho* |
| outflow tract morphogenesis | GO:BP | 0.006 | *robo2,shg* |
| cell fate commitment involved in pattern specification | GO:BP | 0.007 | *tup,scrib,dlg1,rho* |
| positive regulation of hippo signaling | GO:BP | 0.007 | *pyd,kibra,ptc,hppy* |
| positive regulation of transcription elongation by RNA polymerase II | GO:BP | 0.008 | *Hand,trx,Su(Tpl)* |
| positive regulation of actin filament polymerization | GO:BP | 0.008 | *chinmo,DAAM,IRSp53* |
| muscle cell fate determination | GO:BP | 0.009 | *tup,how* |
| actin crosslink formation | GO:BP | 0.009 | *IRSp53,Fim* |
| establishment or maintenance of polarity of larval imaginal disc epithelium | GO:BP | 0.009 | *scrib,dlg1* |
| protein localization to endosome | GO:BP | 0.009 | *Grip,scrib* |
| endosome to plasma membrane protein transport | GO:BP | 0.009 | *Grip,scrib* |
| negative regulation of receptor signaling pathway via JAK-STAT | GO:BP | 0.01 | *apt,ken,wdp* |
| cell competition in a multicellular organism | GO:BP | 0.01 | *shg,mbc,drpr* |
| negative regulation of fibroblast growth factor receptor signaling pathway | GO:BP | 0.013 | *trol,Ptp10D* |
| regulation of RNA metabolic process | GO:BP | 0.014 | *apt,CG9650,chinmo,Glut4EF,ftz-f1,ps,tup,tin,Hand,sbb,luna,tai,CG9932,Pur-alpha,ken,kibra,how,AGO1,bun,Smr,spen,bru2,lola,REPTOR,HDAC4,mam,trx,Su(Tpl),CG14073* |
| imaginal disc-derived wing vein morphogenesis | GO:BP | 0.016 | *dally,Dys,spen,rho* |
| positive regulation of fibroblast growth factor receptor signaling pathway | GO:BP | 0.017 | *trol,dally* |
| protein localization to synapse | GO:BP | 0.017 | *Grip,scrib* |
| regulation of short-term neuronal synaptic plasticity | GO:BP | 0.017 | *Snap25,Dys* |
| gonadal mesoderm development | GO:BP | 0.021 | *shg,tin* |
| oenocyte development | GO:BP | 0.021 | *how,rho* |
| cardioblast cell fate specification | GO:BP | 0.021 | *robo2,fz2* |
| axon midline choice point recognition | GO:BP | 0.021 | *robo2,lola,shot* |
| positive regulation of filopodium assembly | GO:BP | 0.021 | *kon,DAAM,shot* |
| positive regulation of canonical Wnt signaling pathway | GO:BP | 0.023 | *dally,nkd,Eip63E,spen* |
| germarium-derived oocyte fate determination | GO:BP | 0.023 | *par-1,AGO1,hts* |
| neuroblast fate determination | GO:BP | 0.023 | *kuz,spen,mam* |
| determination of genital disc primordium | GO:BP | 0.025 | *ptc,rho* |
| larval visceral muscle development | GO:BP | 0.025 | *klar,mbc* |
| zonula adherens assembly | GO:BP | 0.025 | *shg,scrib* |
| heterophilic cell-cell adhesion via plasma membrane cell adhesion molecules | GO:BP | 0.027 | *beat-IIa,scb,glec* |
| tracheal pit formation in open tracheal system | GO:BP | 0.03 | *Timp,rho* |
| regulation of epithelial cell migration, open tracheal system | GO:BP | 0.03 | *robo2,ptc* |
| pericardial nephrocyte differentiation | GO:BP | 0.03 | *tin,kuz* |
| basal protein localization | GO:BP | 0.03 | *scrib,dlg1* |
| membrane protein ectodomain proteolysis | GO:BP | 0.03 | *Timp,kuz* |
| ommatidial rotation | GO:BP | 0.032 | *shg,pk,nmo* |
| female germ-line stem cell population maintenance | GO:BP | 0.032 | *shg,fax,AGO1* |
| regulation of metabolic process | GO:BP | 0.035 | *apt,CG9650,Timp,chinmo,Glut4EF,ftz-f1,ps,tup,tin,Hand,dnr1,rdx,sbb,luna,tai,CG9932,Pur-alpha,par-1,ken,kek1,Pax,kibra,hid,how,corto,Mob2,AGO1,bun,Smr,spen,CG32369,bru2,rho,lola,REPTOR,HDAC4,mam,trx,Su(Tpl),CG14073,Atg17,sm* |
| determination of digestive tract left/right asymmetry | GO:BP | 0.035 | *shg,tin* |
| regulation of myoblast fusion | GO:BP | 0.035 | *mspo,Pax* |
| positive regulation of JNK cascade | GO:BP | 0.036 | *rdx,mbc,hppy* |
| R3/R4 cell fate commitment | GO:BP | 0.04 | *scrib,lola* |
| ectopic germ cell programmed cell death | GO:BP | 0.04 | *wun,wun2* |
| anterior Malpighian tubule development | GO:BP | 0.04 | *rols,mbc* |
| axon ensheathment | GO:BP | 0.04 | *Nrg,how* |
| R8 cell development | GO:BP | 0.04 | *pyd,kibra* |
| establishment of thoracic bristle planar orientation | GO:BP | 0.046 | *spen* |
| miRNA-mediated gene silencing by mRNA destabilization | GO:BP | 0.046 | *AGO1* |
| regulation of Roundabout signaling pathway | GO:BP | 0.046 | *kuz* |
| regulation of Ral protein signal transduction | GO:BP | 0.046 | *RalGPS* |
| negative regulation of semaphorin-plexin signaling pathway | GO:BP | 0.046 | *dally* |
| basement membrane assembly involved in embryonic body morphogenesis | GO:BP | 0.046 | *Ndg* |
| postsynaptic density assembly | GO:BP | 0.046 | *Prosap* |
| regulation of non-canonical Wnt signaling pathway | GO:BP | 0.046 | *Prosap* |
| anterior mRNA localization involved in anterior/posterior axis specification | GO:BP | 0.046 | *jvl* |
| macropinocytosis | GO:BP | 0.046 | *rudhira* |
| cGMP transport | GO:BP | 0.046 | *w* |
| xanthine transport | GO:BP | 0.046 | *w* |
| positive regulation of phagocytosis, engulfment | GO:BP | 0.046 | *drpr* |
| cGMP catabolic process | GO:BP | 0.046 | *Pde9* |
| positive regulation of extracellular matrix assembly | GO:BP | 0.046 | *loh* |
| negative regulation of pole plasm oskar mRNA localization | GO:BP | 0.046 | *klar* |
| negative regulation of membrane protein ectodomain proteolysis | GO:BP | 0.046 | *Timp* |
| tryptophan transport | GO:BP | 0.046 | *w* |
| testicular fusome organization | GO:BP | 0.046 | *hts* |
| oocyte localization involved in germarium-derived egg chamber formation | GO:BP | 0.048 | *shg,jvl* |
| negative regulation of imaginal disc growth | GO:BP | 0.048 | *scrib,dlg1* |
| ventral cord development | GO:BP | 0.05 | *Timp,robo2,tin,kuz* |
| protein binding | GO:MF | 5.79e-09 | *beat-IIa,kon,apt,Timp,rols,robo2,pain,cv-2,Sema5c,trol,chinmo,klar,RhoGAP18B,ftz-f1,shg,Egfrap,jvl,tup,CG17839,Grip,scb,pyd,dally,Hand,nkd,DAAM,rdx,Prosap,sbb,Fhos,Lac,Nrg,IRSp53,fax,fz2,tai,Pur-alpha,kuz,par-1,pk,ken,kek1,kibra,nmo,form3,Snap25,hid,ptc,AnxB9,mbc,Eip63E,scrib,Zasp52,corto,Dys,Mob2,AGO1,bun,Smr,Fim,CG32369,dlg1,cindr,tei,LRP1,gukh,lola,shot,His3.3B,REPTOR,HDAC4,Mcr,drpr,Tm2,wdp,trx,Septin2,Su(Tpl),Wdr62,CG14073,Lrch,sick,rdgA,hts,Atg17,Ptp10D* |
| actin binding | GO:MF | 0.002 | *DAAM,Fhos,form3,Zasp52,Dys,Fim,gukh,shot,Tm2,hts* |
| protein homodimerization activity | GO:MF | 0.012 | *robo2,shg,rdx,Lac,corto,bun,trx,Septin2* |
| protein dimerization activity | GO:MF | 0.012 | *robo2,ftz-f1,shg,scb,Hand,rdx,Lac,tai,corto,bun,His3.3B,REPTOR,trx,Septin2* |
| cell adhesion molecule binding | GO:MF | 0.014 | *rols,scb,pyd,Nrg,cindr,tei* |
| molecular adaptor activity | GO:MF | 0.014 | *rdx,Prosap,sbb,tai,Pax,kibra,Snap25,Smr,gukh,shot,mam,Septin2,CG14073,Atg17* |
| cytoskeletal protein binding | GO:MF | 0.027 | *klar,shg,DAAM,Fhos,form3,AnxB9,Zasp52,Dys,Fim,gukh,shot,Tm2,hts* |
| transcription regulator activity | GO:MF | 0.036 | *apt,CG9650,Glut4EF,ftz-f1,tup,tin,Hand,sbb,luna,tai,CG9932,Pur-alpha,ken,Pax,Smr,lola,REPTOR,mam,trx,CG14073* |
| identical protein binding | GO:MF | 0.038 | *robo2,shg,rdx,Lac,Pur-alpha,corto,bun,trx,Septin2* |
| enzyme binding | GO:MF | 0.047 | *Timp,RhoGAP18B,Egfrap,DAAM,rdx,kibra,hid,mbc,Mob2,AGO1,dlg1,trx,Su(Tpl),Atg17* |

### 1.7. Gene Ontology Enrichment Analysis: trachea

Significant GO terms (p < 0.05) with associated genes

| GO Term | Source | P-value | Genes |
| --- | --- | --- | --- |
| animal organ morphogenesis | GO:BP | 5.80e-30 | *dpy,exp,uif,verm,kmr,ed,ImpE1,bbg,tyn,sano,Myo10A,grn,RhoGEF64C,grh,caps,Fas2,cv-c,baz,Ubx,heph,dlp,zfh2,Amph,fz2,CadN,mbl,hth,NetB,ec,Dys,sdk,Hs6st,wb,Myo81F,rdx,Nrg,EcR,kis,lola,bun,tkv,Fhos,Mob2,Egfr,mam,spen,mew,sli,fra,Drak,foxo,psq,Cip4,Src42A,scrib,Moe,RhoGAP19D,elB,Mmp2,Ten-m,siz,l(2)gl,sfl,sgg,rl,cic,app,Gug,Hipk,pan,Diap1,Nipped-B,Pka-C1* |
| cell surface receptor signaling pathway | GO:BP | 6.73e-10 | *apolpp,uif,ed,CG41520,grh,Trim9,Fas2,Tl,dlp,fz2,Pvf3,rdx,kek5,tkv,sima,Egfr,mam,spen,Gp150,mew,sli,Ptp10D,fra,foxo,Src42A,Dyrk2,InR,elB,Mmp2,Gprk1,Sarm,for,l(2)gl,sfl,sgg,rl,cic,Gug,Hipk,pan,FER,Diap1,E(bx),Pka-C1* |
| motor neuron axon guidance | GO:BP | 5.05e-09 | *vvl,grn,caps,Fas2,beat-IIIc,fz2,NetB,Nrg,Ptp10D,fra,Sdc,Mmp2,Ten-m,for* |
| negative regulation of signal transduction | GO:BP | 7.20e-08 | *uif,ed,Fas2,cv-c,Tl,RapGAP1,rdx,dnc,kek5,Strn-Mlck,Egfr,Ptp10D,foxo,Src42A,Eip75B,Mmp2,Gprk1,Sarm,l(2)gl,sgg,rdgA,Gug,Hipk,raskol,mnb,Diap1,E(bx),Pka-C1* |
| chitin-based cuticle development | GO:BP | 4.05e-06 | *dpy,dsx-c73A,Gasp,verm,mgl,tyn,serp,Cht7,stw,grh,fz2,sdt,EcR,CrebA,l(2)gl,sgg,pan* |
| regulation of metabolic process | GO:BP | 2.72e-05 | *CG6118,vvl,exp,CG41520,grn,bi,grh,bru3,CG46385,CG8312,Ubx,Pdp1,ab,heph,Tl,zfh2,mbl,hth,Gbs-70E,sm,tai,rdx,EcR,kis,lola,bun,sima,CrebA,Mob2,Egfr,mam,Tis11,spen,Tet,lilli,foxo,psq,l(3)80Fj,Samuel,InR,CG32369,brat,elB,Eip75B,tral,CG17124,eIF4G1,eIF4EHP,Hsc70-3,SNF4Agamma,l(2)gl,Parp,gpp,mld,sgg,Hers,rl,cic,Gug,Nipped-A,Hipk,pan,CG14073,mnb,Sec61gamma,PAN3,Bicra,Diap1,E(bx),Nipped-B,Gyf,Pka-C1* |
| regulation of tube length, open tracheal system | GO:BP | 1.27e-04 | *verm,serp,sano,grh,Fas2,Egfr* |
| zonula adherens assembly | GO:BP | 1.56e-04 | *baz,sdt,scrib,kst* |
| segment polarity determination | GO:BP | 1.79e-04 | *dlp,fz2,rdx,Egfr,sfl,sgg,pan* |
| chitin-based larval cuticle pattern formation | GO:BP | 2.25e-04 | *fz2,CrebA,l(2)gl,sgg,pan* |
| ecdysis, chitin-based cuticle | GO:BP | 2.25e-04 | *Cht7,Cht6,Hmgcr,EcR,Eip75B* |
| regulation of RNA metabolic process | GO:BP | 2.26e-04 | *CG6118,vvl,exp,grn,bi,grh,bru3,CG46385,CG8312,Ubx,Pdp1,ab,Tl,zfh2,mbl,hth,tai,EcR,kis,lola,bun,sima,CrebA,mam,Tis11,spen,Tet,lilli,foxo,psq,Samuel,elB,Eip75B,mld,Hers,rl,cic,Gug,Nipped-A,pan,CG14073,mnb,PAN3,Bicra,E(bx),Nipped-B* |
| imaginal disc-derived wing vein morphogenesis | GO:BP | 2.66e-04 | *cv-c,heph,Dys,Egfr,spen,Gug,Hipk* |
| cell fate commitment involved in pattern specification | GO:BP | 4.21e-04 | *uif,Ubx,EcR,Egfr,scrib,l(2)gl* |
| positive regulation of organ growth | GO:BP | 6.74e-04 | *kis,InR,Hipk,mnb* |
| peripheral nervous system development | GO:BP | 7.22e-04 | *vvl,Trim9,hth,EcR,Egfr,spen,fra* |
| positive regulation of border follicle cell migration | GO:BP | 7.22e-04 | *tai,kis,spri,Egfr,InR,Diap1* |
| negative regulation of biosynthetic process | GO:BP | 9.94e-04 | *grn,bru3,CG46385,Ubx,ab,heph,EcR,kis,Egfr,Tis11,foxo,psq,Samuel,brat,elB,Eip75B,tral,eIF4EHP,Hsc70-3,gpp,Hers,cic,Gug,pan,CG14073,PAN3,Diap1,E(bx),Gyf* |
| dorsal closure | GO:BP | 0.001 | *ed,Myo10A,cv-c,tkv,Egfr,Src42A,scrib,l(2)gl,FER* |
| synaptic target inhibition | GO:BP | 0.001 | *Tl,beat-IIIc,fz2,sli* |
| olfactory learning | GO:BP | 0.001 | *Fas2,dnc,Egfr,sgg,mnb,Pka-C1* |
| short-term memory | GO:BP | 0.001 | *Fas2,dnc,kis,for,Nipped-B* |
| positive regulation of transcription by RNA polymerase II | GO:BP | 0.001 | *vvl,grn,grh,CG8312,Pdp1,Tl,hth,tai,EcR,sima,CrebA,mam,Tet,lilli,foxo,Eip75B,mld,pan,E(bx)* |
| chitin-based embryonic cuticle biosynthetic process | GO:BP | 0.002 | *dpy,tyn,grh,EcR* |
| peptidyl-threonine phosphorylation | GO:BP | 0.002 | *bt,Dyrk2,Hipk,mnb* |
| homophilic cell adhesion via plasma membrane adhesion molecules | GO:BP | 0.002 | *Fas3,ed,caps,Fas2,CadN,sdk* |
| negative regulation of epidermal growth factor receptor signaling pathway | GO:BP | 0.002 | *ed,Fas2,Ptp10D,Src42A,Gug* |
| peripheral nervous system neuron axonogenesis | GO:BP | 0.002 | *Trim9,fra* |
| muscle cell cellular homeostasis | GO:BP | 0.002 | *uif,mbl,Dys,Fhos,Gyf* |
| regulation of establishment of planar polarity | GO:BP | 0.003 | *kmr,ed,baz,Hipk* |
| apposition of dorsal and ventral imaginal disc-derived wing surfaces | GO:BP | 0.003 | *dpy,tyn,mam,mew* |
| long-term memory | GO:BP | 0.003 | *pxb,EcR,Mob2,uex,Ptp10D,for,mld* |
| dorsal appendage formation | GO:BP | 0.003 | *ed,bun,Egfr,jvl,rl,cic* |
| behavioral response to ethanol | GO:BP | 0.003 | *Fas2,pxb,Gbs-70E,dnc,Egfr,rl* |
| positive regulation of hippo signaling | GO:BP | 0.003 | *ed,Hsc70-3,l(2)gl,Hipk,Pka-C1* |
| spiracle morphogenesis, open tracheal system | GO:BP | 0.004 | *RhoGEF64C,cv-c,Egfr,pan* |
| tracheal pit formation in open tracheal system | GO:BP | 0.004 | *cv-c,Mmp2,rl* |
| negative regulation of cell growth | GO:BP | 0.006 | *Tl,sima,foxo,cic* |
| trachea morphogenesis | GO:BP | 0.006 | *exp,verm,Fhos* |
| mesodermal cell fate determination | GO:BP | 0.006 | *mam,pan* |
| wax biosynthetic process | GO:BP | 0.006 | *CG5065,CG8306* |
| positive regulation of smoothened signaling pathway | GO:BP | 0.006 | *apolpp,dlp,Gprk1,Hipk,Pka-C1* |
| R3/R4 cell fate commitment | GO:BP | 0.007 | *grh,lola,scrib* |
| anterior Malpighian tubule development | GO:BP | 0.007 | *Ubx,rols,tkv* |
| axon midline choice point recognition | GO:BP | 0.007 | *Trim9,lola,sli,fra* |
| gastrulation | GO:BP | 0.007 | *baz,Ubx,tkv,mam,sli,sfl,pan* |
| determination of adult lifespan | GO:BP | 0.008 | *sm,EcR,kis,bun,Egfr,alpha-Man-Ia,foxo,InR,mld,rl* |
| maintenance of epithelial integrity, open tracheal system | GO:BP | 0.009 | *cv-c,Egfr,mew* |
| specification of animal organ identity | GO:BP | 0.01 | *Ubx,hth* |
| digestive tract mesoderm development | GO:BP | 0.01 | *wb,sli* |
| netrin-activated signaling pathway | GO:BP | 0.01 | *Trim9,fra* |
| synaptic target attraction | GO:BP | 0.01 | *NetB,Ten-m* |
| positive regulation of wound healing | GO:BP | 0.01 | *Egfr,InR,rl* |
| R7 cell development | GO:BP | 0.01 | *CadN,lola,Ten-m* |
| cellular response to growth factor stimulus | GO:BP | 0.011 | *grh,dlp,kek5,tkv,Ptp10D,Mmp2,sfl,rl* |
| imaginal disc-derived leg morphogenesis | GO:BP | 0.011 | *RhoGEF64C,Ubx,hth,Drak,RhoGAP19D,Gug* |
| positive regulation of gene expression | GO:BP | 0.012 | *vvl,CG46385,Tl,lola,Egfr,Parp,gpp,Hipk,pan,Nipped-B,Pka-C1* |
| cholesterol homeostasis | GO:BP | 0.012 | *EcR,InR,SNF4Agamma* |
| negative regulation of cell population proliferation | GO:BP | 0.015 | *pxb,foxo,scrib,brat,Mmp2,l(2)gl* |
| positive regulation of canonical Wnt signaling pathway | GO:BP | 0.015 | *apolpp,dlp,spen,Hipk,Diap1* |
| intestinal stem cell homeostasis | GO:BP | 0.015 | *bun,Src42A,InR,Hsc70-3* |
| establishment of epithelial cell planar polarity | GO:BP | 0.016 | *l(2)gl,sgg* |
| chitin-based cuticle attachment to epithelium | GO:BP | 0.016 | *dpy,tyn* |
| establishment or maintenance of polarity of larval imaginal disc epithelium | GO:BP | 0.016 | *scrib,Moe* |
| protein localization to endosome | GO:BP | 0.016 | *scrib,Moe* |
| midgut development | GO:BP | 0.017 | *cv-c,Ubx,fz2,mew* |
| tracheal outgrowth, open tracheal system | GO:BP | 0.017 | *grh,Egfr,Mmp2* |
| positive regulation of peptidoglycan recognition protein signaling pathway | GO:BP | 0.019 | *lola,sick,Parp* |
| apical protein localization | GO:BP | 0.019 | *baz,sdt,l(2)gl* |
| glial cell migration | GO:BP | 0.02 | *Pvf3,NetB,sli,fra* |
| imaginal disc-derived wing vein specification | GO:BP | 0.02 | *Dys,Egfr,sfl,rl,cic* |
| peptidyl-serine phosphorylation | GO:BP | 0.02 | *bt,Dyrk2,Hipk,Tlk,mnb* |
| lipid phosphorylation | GO:BP | 0.021 | *rdgA,CG31140* |
| positive regulation of muscle organ development | GO:BP | 0.021 | *Ubx,tkv* |
| adult chitin-based cuticle development | GO:BP | 0.021 | *verm,serp* |
| establishment or maintenance of polarity of embryonic epithelium | GO:BP | 0.021 | *sdt,scrib* |
| secretory granule localization | GO:BP | 0.021 | *unc-13,Cadps* |
| negative regulation of fibroblast growth factor receptor signaling pathway | GO:BP | 0.021 | *Ptp10D,Mmp2* |
| regulation of cell projection organization | GO:BP | 0.021 | *vvl,Trim9,cv-c,baz,fz2,CadN,NetB,lola,fra* |
| establishment of epithelial cell apical/basal polarity | GO:BP | 0.021 | *Egfr,scrib* |
| septate junction assembly | GO:BP | 0.021 | *CG9628,Nrg,scrib,l(2)gl* |
| asymmetric protein localization involved in cell fate determination | GO:BP | 0.022 | *uif,baz,l(2)gl* |
| germ-band shortening | GO:BP | 0.022 | *EcR,Egfr,InR* |
| protein autophosphorylation | GO:BP | 0.023 | *bt,InR,Tlk,mnb* |
| regulation of membrane potential in photoreceptor cell | GO:BP | 0.028 | *Moe,SK* |
| positive regulation of feeding behavior | GO:BP | 0.028 | *foxo,mnb* |
| negative regulation of compound eye retinal cell programmed cell death | GO:BP | 0.028 | *Egfr,Diap1* |
| regulation of ecdysteroid metabolic process | GO:BP | 0.028 | *Eip75B,eIF4EHP* |
| germ-line stem-cell niche homeostasis | GO:BP | 0.028 | *fz2,tai,InR* |
| imaginal disc eversion | GO:BP | 0.028 | *ImpE1,Mmp2* |
| ganglion mother cell fate determination | GO:BP | 0.028 | *grh,brat* |
| toll-like receptor signaling pathway | GO:BP | 0.028 | *Tl,Sarm* |
| positive regulation of cell size | GO:BP | 0.029 | *Pvf3,InR,rl,Nipped-B* |
| biological process involved in interspecies interaction between organisms | GO:BP | 0.033 | *vvl,nur,Tl,RhoU,mtd,Nrg,lola,sick,sima,spen,foxo,Src42A,Eip75B,Mmp2,Sarm,Parp,rl,FER,E(bx)* |
| negative regulation of synaptic assembly at neuromuscular junction | GO:BP | 0.033 | *Mob2,Cip4,Src42A,sgg* |
| dorsal closure, spreading of leading edge cells | GO:BP | 0.035 | *Egfr,Src42A* |
| entry into diapause | GO:BP | 0.035 | *foxo,InR* |
| diacylglycerol metabolic process | GO:BP | 0.035 | *rdgA,CG31140* |
| microtubule anchoring | GO:BP | 0.043 | *Msp300,Moe* |
| rhabdomere membrane biogenesis | GO:BP | 0.043 | *Amph,Moe* |
| lateral inhibition | GO:BP | 0.043 | *mam,Mmp2* |
| intercellular transport | GO:BP | 0.043 | *shakB,Inx2* |
| apical constriction | GO:BP | 0.046 | *dyl,tyn,Shrm* |
| mushroom body development | GO:BP | 0.05 | *Fas2,ab,Nrg,EcR,bun* |
| neuroblast fate determination | GO:BP | 0.05 | *mam,spen,brat* |
| protein binding | GO:MF | 2.91e-14 | *CG6118,apolpp,Fas3,ImpL1,exp,uif,verm,ed,mgl,ImpE1,bbg,serp,CG41520,Myo10A,bt,CG43333,SKIP,CG15080,grh,Trim9,caps,Fili,bru3,Fas2,baz,Ubx,ab,osp,Tl,dlp,toc,rols,Jupiter,Amph,fz2,CadN,sdt,RhoU,hth,CG32264,Pvf3,Shrm,NetB,ec,Dys,sdk,Zasp52,Gbs-70E,Hmgcr,Myo81F,tutl,smash,Msp300,Wdr62,tai,rdx,Nrg,kek5,EcR,kis,lola,bun,spri,sick,tkv,sima,Fhos,Strn-Mlck,Mob2,Egfr,uex,unc-13,Gp150,mew,Spn,sli,Ptp10D,fra,lilli,foxo,psq,l(3)80Fj,Cip4,Samuel,Src42A,stx,scrib,InR,zormin,Moe,CG32369,jvl,brat,RhoGAP19D,elB,Gprk1,tral,eIF4G1,Sarm,Ten-m,eIF4EHP,kst,Hsc70-3,SNF4Agamma,SK,gukh,siz,l(2)gl,Parp,Chd64,sgg,rl,rdgA,cic,CG13917,Gug,CG42748,Nipped-A,Hipk,pan,Evi5,Tlk,CG14073,mnb,FER,PAN3,Diap1,E(bx),Nipped-B,Gyf,Pka-C1* |
| cell adhesion molecule binding | GO:MF | 1.62e-06 | *Fas3,ed,CG43333,CG15080,Fas2,rols,CadN,sdk,Nrg,mew,Moe,siz* |
| protein kinase activity | GO:MF | 1.52e-05 | *bt,tkv,Strn-Mlck,Egfr,CG31145,Drak,Src42A,Dyrk2,Nuak,InR,Gprk1,for,sgg,rl,Hipk,Tlk,mnb,FER,Pka-C1* |
| actin binding | GO:MF | 1.52e-05 | *Myo10A,osp,CG32264,Shrm,Dys,Zasp52,Myo81F,Msp300,Fhos,Spn,zormin,Moe,kst,gukh,Chd64* |
| phosphotransferase activity, alcohol group as acceptor | GO:MF | 5.38e-05 | *bt,tkv,Strn-Mlck,Egfr,CG31145,Drak,Src42A,Dyrk2,Nuak,InR,Gprk1,for,sgg,rl,rdgA,Hipk,CG31140,Tlk,mnb,FER,Pka-C1* |
| protein serine/threonine kinase activity | GO:MF | 1.39e-04 | *bt,tkv,Strn-Mlck,CG31145,Drak,Dyrk2,Nuak,Gprk1,for,sgg,rl,Hipk,Tlk,mnb,Pka-C1* |
| kinase activity | GO:MF | 2.33e-04 | *bt,sdt,tkv,Strn-Mlck,Egfr,CG31145,Drak,Src42A,Dyrk2,Nuak,InR,Gprk1,SNF4Agamma,for,sgg,rl,rdgA,Hipk,CG31140,Tlk,mnb,FER,Pka-C1* |
| actin filament binding | GO:MF | 3.12e-04 | *Myo10A,osp,Shrm,Dys,Myo81F,Msp300,Fhos,Spn,kst,Chd64* |
| cytoskeletal protein binding | GO:MF | 3.12e-04 | *apolpp,Myo10A,osp,toc,Jupiter,CG32264,Shrm,Dys,Zasp52,Myo81F,smash,Msp300,Fhos,Spn,zormin,Moe,Ten-m,kst,gukh,l(2)gl,Chd64* |
| DNA-binding transcription factor binding | GO:MF | 4.64e-04 | *tai,EcR,lilli,Sarm,rl,cic,Gug,Hipk,pan,E(bx)* |
| transferase activity, transferring phosphorus-containing groups | GO:MF | 5.25e-04 | *bt,CG46385,sdt,tkv,Strn-Mlck,Egfr,CG31145,Drak,Src42A,Dyrk2,Nuak,InR,Gprk1,SNF4Agamma,for,Parp,sgg,rl,rdgA,Hipk,CG31140,Tlk,mnb,FER,Pka-C1* |
| protein tyrosine kinase activity | GO:MF | 9.56e-04 | *Egfr,Src42A,Dyrk2,InR,Hipk,mnb,FER* |
| transcription coregulator binding | GO:MF | 0.001 | *Ubx,hth,tai,EcR,pan* |
| transcription factor binding | GO:MF | 0.001 | *Ubx,hth,tai,EcR,lilli,Sarm,rl,cic,Gug,Hipk,pan,E(bx)* |
| kinase binding | GO:MF | 0.001 | *RhoU,Shrm,Msp300,spri,Mob2,l(3)80Fj,SNF4Agamma,l(2)gl,rl,pan* |
| DNA-binding transcription activator activity, RNA polymerase II-specific | GO:MF | 0.002 | *vvl,grh,Pdp1,hth,EcR,sima,foxo,mld,pan* |
| DNA-binding transcription activator activity | GO:MF | 0.002 | *vvl,grh,Pdp1,hth,EcR,sima,foxo,mld,pan* |
| protein kinase binding | GO:MF | 0.009 | *RhoU,Msp300,spri,Mob2,l(3)80Fj,SNF4Agamma,l(2)gl,rl* |
| deacetylase activity | GO:MF | 0.009 | *serp,Cda5,sfl,Gug* |
| carbohydrate derivative binding | GO:MF | 0.01 | *Gasp,verm,serp,Cda4,Myo10A,Cht7,bt,Cht6,Cda5,RhoU,Myo81F,kis,sick,tkv,betaTub56D,Strn-Mlck,Egfr,CG31145,sli,Drak,Src42A,Dyrk2,Dbp80,Nuak,InR,Gprk1,Hsc70-3,SNF4Agamma,for,PMCA,sgg,rl,rdgA,Hipk,CG31140,Tlk,mnb,FER,PAN3,Pka-C1* |
| RNA polymerase II-specific DNA-binding transcription factor binding | GO:MF | 0.01 | *tai,lilli,Sarm,Gug,E(bx)* |
| GTPase regulator activity | GO:MF | 0.01 | *RhoGEF64C,CG30440,cv-c,RapGAP1,spri,RhoGAP19D,kst,siz,l(2)gl,Evi5,raskol* |
| adenyl ribonucleotide binding | GO:MF | 0.01 | *Myo10A,bt,Myo81F,kis,sick,tkv,Strn-Mlck,Egfr,CG31145,Drak,Src42A,Dyrk2,Dbp80,Nuak,InR,Gprk1,Hsc70-3,SNF4Agamma,for,PMCA,sgg,rl,rdgA,Hipk,CG31140,Tlk,mnb,FER,PAN3,Pka-C1* |
| protein homodimerization activity | GO:MF | 0.01 | *grh,bru3,CadN,Shrm,rdx,EcR,bun,psq,Ten-m* |
| nuclear receptor binding | GO:MF | 0.012 | *tai,Gug,E(bx)* |
| ATP binding | GO:MF | 0.018 | *Myo10A,bt,Myo81F,kis,sick,tkv,Strn-Mlck,Egfr,CG31145,Drak,Src42A,Dyrk2,Dbp80,Nuak,InR,Gprk1,Hsc70-3,for,PMCA,sgg,rl,rdgA,Hipk,CG31140,Tlk,mnb,FER,PAN3,Pka-C1* |
| translation repressor activity | GO:MF | 0.018 | *heph,brat,eIF4EHP,Gyf* |
| transcription coactivator binding | GO:MF | 0.02 | *hth,tai,EcR* |
| protein domain specific binding | GO:MF | 0.023 | *Ubx,Tl,fz2,Dys,fra,psq,InR* |
| purine nucleotide binding | GO:MF | 0.025 | *Myo10A,bt,RhoU,Hmgcr,Myo81F,kis,sick,tkv,betaTub56D,Strn-Mlck,Egfr,CG31145,Drak,Src42A,Dyrk2,Dbp80,Nuak,InR,Gprk1,Hsc70-3,SNF4Agamma,for,Parp,PMCA,sgg,rl,rdgA,Hipk,CG31140,Tlk,mnb,FER,PAN3,Pka-C1* |
| GTPase activator activity | GO:MF | 0.025 | *cv-c,RapGAP1,spri,RhoGAP19D,l(2)gl,Evi5,raskol* |
| identical protein binding | GO:MF | 0.026 | *grh,bru3,Tl,CadN,Shrm,rdx,EcR,bun,psq,InR,Ten-m* |
| chitin deacetylase activity | GO:MF | 0.026 | *serp,Cda5* |
| structural constituent of muscle | GO:MF | 0.026 | *bt,rols,Dys* |
| transmembrane receptor protein kinase activity | GO:MF | 0.028 | *tkv,Egfr,Src42A,InR* |
| lipid binding | GO:MF | 0.032 | *apolpp,kmr,cv-c,baz,Amph,EcR,unc-13,Cip4,Moe,Gprk1,kst,FER* |
| signaling receptor binding | GO:MF | 0.033 | *apolpp,uif,ed,CG41520,Pvf3,EcR,spri,mew,Spn,sli,Src42A,Gprk1,FER* |
| RNA polymerase II cis-regulatory region sequence-specific DNA binding | GO:MF | 0.035 | *vvl,grn,bi,grh,Ubx,Pdp1,zfh2,hth,EcR,CrebA,foxo,Eip75B,mld,Chd64,pan,E(bx)* |
| hormone binding | GO:MF | 0.037 | *mgl,EcR,InR* |
| phospholipid binding | GO:MF | 0.038 | *kmr,baz,Amph,unc-13,Cip4,Moe,Gprk1,kst,FER* |
| transcription regulator activity | GO:MF | 0.039 | *vvl,grn,bi,grh,Ubx,Pdp1,ab,zfh2,hth,tai,EcR,lola,sima,CrebA,mam,lilli,foxo,Eip75B,mld,cic,Gug,pan,CG14073,mnb,E(bx)* |
| cis-regulatory region sequence-specific DNA binding | GO:MF | 0.04 | *vvl,grn,bi,grh,Ubx,Pdp1,zfh2,hth,EcR,CrebA,foxo,Eip75B,mld,Chd64,pan,E(bx)* |
| mRNA regulatory element binding translation repressor activity | GO:MF | 0.04 | *heph,eIF4EHP,Gyf* |
| ATP-dependent diacylglycerol kinase activity | GO:MF | 0.045 | *rdgA,CG31140* |
| enzyme binding | GO:MF | 0.049 | *RhoU,Shrm,ec,Gbs-70E,Msp300,rdx,spri,Mob2,l(3)80Fj,tral,SNF4Agamma,l(2)gl,rl,CG42748,pan,Evi5,Diap1* |

### 1.8. Gene Ontology Enrichment Analysis: NB & neurons & GMC

Significant GO terms (p < 0.05) with associated genes

| GO Term | Source | P-value | Genes |
| --- | --- | --- | --- |
| animal organ morphogenesis | GO:BP | 4.30e-13 | *14-3-3epsilon,chn,Cam,CadN,Gbeta13F,br,ctp,zfh2,Galphao,fz2,CG43867,sbb,Atpalpha,Hrb98DE,Ubx,Lar,kis,bsk,msi,tinc,spen,Wnt4,Nrg,nrv2,caps,CtBP,ct,Fas2,Gug,mam,osa,hth,rin,RhoGAP19D,ena,Ggamma1,shi,tsh,psq* |
| regulation of RNA metabolic process | GO:BP | 2.74e-11 | *fne,scrt,trv,CG13928,14-3-3epsilon,chn,bru3,Imp,chinmo,HmgD,l(3)neo38,br,Antp,elav,zfh2,Tet,Pep,Dsp1,Pits,Sumo,Ssdp,HmgZ,sbb,Hrb98DE,Ubx,kis,fs(1)h,abd-A,gro,spen,Dora,CtBP,tou,REPTOR,ct,mei-P26,nonA,Gug,mam,osa,Rbp1-like,hth,mnb,CG32767,crp,tsh,psq,SC35* |
| regulation of metabolic process | GO:BP | 3.45e-10 | *fne,scrt,trv,CG13928,14-3-3epsilon,chn,bru3,Imp,brat,sm,chinmo,HmgD,l(3)neo38,br,wech,Antp,elav,zfh2,Tet,Pep,Dsp1,Pits,Sumo,Ssdp,HmgZ,sbb,Hrb98DE,Ubx,kek1,Rbp6,kis,fs(1)h,bsk,msi,abd-A,gro,CG17124,spen,Unr,Dora,CtBP,tou,REPTOR,CaMKII,ct,mei-P26,nonA,InR,Gug,mam,osa,Rbp1-like,hth,rin,mnb,tara,CG32767,crp,shi,tsh,psq,SC35,CkIIalpha* |
| negative regulation of biosynthetic process | GO:BP | 1.09e-06 | *scrt,CG13928,chn,bru3,brat,br,wech,Antp,elav,Dsp1,Pits,sbb,Hrb98DE,Ubx,kis,fs(1)h,msi,abd-A,gro,Unr,Dora,CtBP,CaMKII,ct,Gug,CG32767,tsh,psq* |
| trans-synaptic signaling | GO:BP | 1.77e-06 | *CG6329,X11Lbeta,nSyb,Rbp,cpx,cac,brp,cpo,Ank2,unc-13,Ten-a,Atpalpha,Sap47,Cadps,CaMKII,Fas2,shi,Syx1A* |
| cell adhesion involved in heart morphogenesis | GO:BP | 3.49e-06 | *Gbeta13F,Galphao,Nrg,nrv2,Ggamma1* |
| negative regulation of transcription by RNA polymerase II | GO:BP | 3.49e-06 | *scrt,chn,br,Antp,Dsp1,sbb,Ubx,kis,abd-A,gro,CtBP,ct,Gug,CG32767,tsh* |
| peripheral nervous system development | GO:BP | 6.08e-06 | *Appl,chn,Trim9,abd-A,spen,ct,Dscam1,hth* |
| short-term memory | GO:BP | 1.35e-05 | *Appl,brp,Ank2,kis,Fas2,shi* |
| specification of segmental identity, thorax | GO:BP | 2.29e-05 | *Antp,Ubx,tsh* |
| motor neuron axon guidance | GO:BP | 5.41e-05 | *fz2,Ten-a,Lar,Wnt4,Nrg,Sema1a,caps,Fas2* |
| signal release | GO:BP | 7.22e-05 | *nSyb,cpx,cac,brp,unc-13,Cadps,Flo2,InR,shi,Syx1A* |
| female gonad development | GO:BP | 1.00e-04 | *fz2,Wnt4,ct,InR* |
| retinal ganglion cell axon guidance | GO:BP | 1.00e-04 | *CadN,fz2,Lar,Wnt4* |
| axon target recognition | GO:BP | 1.38e-04 | *Liprin-gamma,CadN,sbb,Lar* |
| cell surface receptor signaling pathway | GO:BP | 1.53e-04 | *Imp,Gbeta13F,Sema2b,Galphao,fz2,Sumo,Ssdp,Trim9,kek1,Lar,bsk,gro,Prosap,spen,Wnt4,Sema1a,CtBP,Fas2,InR,Gug,mam,osa,tsh,CkIIalpha* |
| nucleobase-containing compound metabolic process | GO:BP | 2.17e-04 | *fne,scrt,trv,CG13928,14-3-3epsilon,chn,bru3,Imp,brat,sm,chinmo,HmgD,l(3)neo38,br,Antp,Srrm234,elav,zfh2,Tet,Pep,Dsp1,Pits,Sumo,Ssdp,HmgZ,sbb,Hrb98DE,Ubx,CG5065,kis,fs(1)h,abd-A,gro,spen,sdt,Dora,CtBP,tou,REPTOR,ct,mei-P26,nonA,Gug,mam,osa,Rbp1-like,hth,mnb,tara,CG32767,crp,tsh,psq,SC35* |
| secretory granule localization | GO:BP | 3.04e-04 | *unc-13,Cadps,shi* |
| midgut development | GO:BP | 4.35e-04 | *Antp,fz2,Ubx,abd-A,tsh* |
| muscle cell fate specification | GO:BP | 4.84e-04 | *Antp,Ubx,abd-A* |
| germ-line stem-cell niche homeostasis | GO:BP | 8.38e-04 | *Imp,fz2,Wnt4,InR* |
| female sex differentiation | GO:BP | 9.95e-04 | *fz2,Wnt4,ct,InR* |
| intracellular potassium ion homeostasis | GO:BP | 0.001 | *nrv3,Atpalpha,nrv2* |
| sodium ion export across plasma membrane | GO:BP | 0.001 | *nrv3,Atpalpha,nrv2* |
| intracellular sodium ion homeostasis | GO:BP | 0.001 | *nrv3,Atpalpha,nrv2* |
| specification of segmental identity, antennal segment | GO:BP | 0.001 | *Antp,hth* |
| female courtship behavior | GO:BP | 0.001 | *shep,Nrg* |
| positive regulation of cell proliferation involved in compound eye morphogenesis | GO:BP | 0.001 | *hth,tsh* |
| protein kinase C signaling | GO:BP | 0.001 | *CG31140,Flo1* |
| negative regulation of cell differentiation | GO:BP | 0.001 | *chn,brat,CadN,Sema2b,Sema1a,Fas2,osa* |
| import across plasma membrane | GO:BP | 0.002 | *nrv3,Mid1,cac,Atpalpha,nrv2* |
| sensory perception of sound | GO:BP | 0.002 | *nrv3,Cam,Ank2,Atpalpha,nrv2,ct* |
| regulation of tube diameter, open tracheal system | GO:BP | 0.002 | *Atpalpha,nrv2,Fas2* |
| homophilic cell adhesion via plasma membrane adhesion molecules | GO:BP | 0.002 | *CadN,Lar,caps,Fas2,Dscam1* |
| neuroblast fate determination | GO:BP | 0.002 | *brat,abd-A,spen,mam* |
| mushroom body development | GO:BP | 0.002 | *Appl,chinmo,bsk,Nrg,Fas2,Dscam1* |
| presynaptic dense core vesicle exocytosis | GO:BP | 0.003 | *unc-13,shi* |
| positive regulation of organ growth | GO:BP | 0.003 | *kis,InR,mnb* |
| positive regulation of neuron remodeling | GO:BP | 0.003 | *ctp,bsk,InR* |
| establishment or maintenance of cytoskeleton polarity involved in gastrulation | GO:BP | 0.005 | *Gbeta13F,Ggamma1* |
| synaptic target attraction | GO:BP | 0.005 | *Sema2b,Ten-a* |
| subsynaptic reticulum organization | GO:BP | 0.005 | *Prosap,ena* |
| formation of a compartment boundary | GO:BP | 0.005 | *ct,ena* |
| specification of animal organ identity | GO:BP | 0.005 | *Ubx,hth* |
| establishment of glial blood-brain barrier | GO:BP | 0.005 | *Galphao,Nrg,nrv2* |
| septate junction assembly | GO:BP | 0.005 | *Galphao,Atpalpha,Nrg,nrv2* |
| peptidyl-threonine phosphorylation | GO:BP | 0.005 | *CaMKII,mnb,CkIIalpha* |
| potassium ion import across plasma membrane | GO:BP | 0.005 | *nrv3,Atpalpha,nrv2* |
| positive regulation of transcription by RNA polymerase II | GO:BP | 0.006 | *chn,br,Antp,Tet,Ssdp,fs(1)h,abd-A,CtBP,tou,mam,hth,CG32767,tsh* |
| protein autophosphorylation | GO:BP | 0.006 | *CaMKII,InR,mnb,Tlk* |
| synaptic vesicle docking | GO:BP | 0.006 | *nSyb,unc-13,Syx1A* |
| apical protein localization | GO:BP | 0.006 | *Gbeta13F,sdt,Ggamma1* |
| RNA splicing, via transesterification reactions | GO:BP | 0.007 | *trv,bru3,Imp,sm,Srrm234,Pep,Hrb98DE,nonA,Rbp1-like,SC35* |
| mesodermal cell fate specification | GO:BP | 0.007 | *Ubx,abd-A* |
| neuroblast development | GO:BP | 0.007 | *brat,Antp,abd-A* |
| imaginal disc-derived leg morphogenesis | GO:BP | 0.008 | *Ubx,Gug,osa,hth,RhoGAP19D* |
| dendrite self-avoidance | GO:BP | 0.008 | *Sema2b,Fas2,Dscam1* |
| axonal fasciculation | GO:BP | 0.008 | *CadN,Fas2,Dscam1* |
| mesectoderm development | GO:BP | 0.01 | *Gbeta13F,Ggamma1* |
| positive regulation of muscle organ development | GO:BP | 0.01 | *Ubx,abd-A* |
| regulation of transcriptional start site selection at RNA polymerase II promoter | GO:BP | 0.01 | *Rbp1-like,SC35* |
| calcium ion import across plasma membrane | GO:BP | 0.01 | *Mid1,cac* |
| convergent extension involved in gastrulation | GO:BP | 0.01 | *Gbeta13F,Ggamma1* |
| negative regulation of axon extension involved in axon guidance | GO:BP | 0.01 | *Sema2b,Sema1a* |
| visual behavior | GO:BP | 0.012 | *cac,nonA,mnb* |
| regulation of short-term neuronal synaptic plasticity | GO:BP | 0.013 | *Sap47,shi* |
| cytoskeletal matrix organization at active zone | GO:BP | 0.013 | *Rbp,brp* |
| regulation of myosin II filament organization | GO:BP | 0.013 | *Gbeta13F,Ggamma1* |
| mesoderm formation | GO:BP | 0.014 | *Ubx,abd-A,mam* |
| wing disc dorsal/ventral pattern formation | GO:BP | 0.015 | *14-3-3epsilon,sbb,osa,tara* |
| positive regulation of filopodium assembly | GO:BP | 0.015 | *Lar,Flo2,ena* |
| regulation of alternative mRNA splicing, via spliceosome | GO:BP | 0.015 | *trv,bru3,Hrb98DE,Rbp1-like,SC35* |
| progression of morphogenetic furrow involved in compound eye morphogenesis | GO:BP | 0.016 | *chn,br* |
| proboscis extension reflex | GO:BP | 0.016 | *Ggamma1,shi* |
| dorsal appendage formation | GO:BP | 0.016 | *br,Sumo,bsk,shi* |
| regulation of gastrulation | GO:BP | 0.016 | *Gbeta13F,Ggamma1* |
| negative regulation of signal transduction | GO:BP | 0.016 | *Cam,Gbeta13F,kek1,Lar,gro,Prosap,scyl,CtBP,CaMKII,Fas2,Gug,mnb* |
| gastrulation | GO:BP | 0.017 | *Gbeta13F,Ubx,abd-A,mam,Ggamma1* |
| regulation of tube length, open tracheal system | GO:BP | 0.018 | *Atpalpha,nrv2,Fas2* |
| photoreceptor cell fate commitment | GO:BP | 0.019 | *14-3-3epsilon,br,msi,hth,rin* |
| calcium ion import | GO:BP | 0.019 | *Mid1,cac* |
| central complex development | GO:BP | 0.019 | *Ten-a,Nrg* |
| lateral inhibition | GO:BP | 0.019 | *mam,CkIIalpha* |
| female germ-line stem cell population maintenance | GO:BP | 0.022 | *fax,Hrb98DE,InR* |
| regulation of monoatomic ion transmembrane transport | GO:BP | 0.024 | *CG6329,cac,Cam,nrv2* |
| maintenance of presynaptic active zone structure | GO:BP | 0.026 | *brp,Ten-a* |
| cell fate commitment involved in pattern specification | GO:BP | 0.03 | *Ubx,abd-A,CtBP* |
| head involution | GO:BP | 0.034 | *scyl,ena,tsh* |
| myosin filament organization | GO:BP | 0.035 | *Gbeta13F,Ggamma1* |
| vesicle fusion | GO:BP | 0.035 | *nSyb,brp,aus,Syx1A* |
| positive regulation of cell-cell adhesion mediated by cadherin | GO:BP | 0.036 | *Flo1* |
| proteasomal ubiquitin-independent protein catabolic process | GO:BP | 0.036 | *stx* |
| establishment of thoracic bristle planar orientation | GO:BP | 0.036 | *spen* |
| postsynaptic density assembly | GO:BP | 0.036 | *Prosap* |
| peptidyl-threonine autophosphorylation | GO:BP | 0.036 | *CaMKII* |
| regulation of light-activated channel activity | GO:BP | 0.036 | *Cam* |
| detection of chemical stimulus involved in sensory perception of sour taste | GO:BP | 0.036 | *OtopLa* |
| negative regulation of RNA polymerase II transcription preinitiation complex assembly | GO:BP | 0.036 | *Dsp1* |
| male courtship behavior, veined wing generated song production | GO:BP | 0.036 | *cac,nonA* |
| R7 cell development | GO:BP | 0.036 | *CadN,Lar* |
| dense core granule priming | GO:BP | 0.036 | *unc-13* |
| morphogenesis of an epithelial sheet | GO:BP | 0.036 | *bsk,sdt* |
| negative regulation of mRNA 3'-end processing | GO:BP | 0.036 | *elav* |
| arachidonate metabolic process | GO:BP | 0.036 | *inaE* |
| regulation of non-canonical Wnt signaling pathway | GO:BP | 0.036 | *Prosap* |
| asymmetric neuroblast division | GO:BP | 0.036 | *brat,Gbeta13F,Ggamma1* |
| diacylglycerol catabolic process | GO:BP | 0.036 | *inaE* |
| positive regulation of ubiquitin-dependent endocytosis | GO:BP | 0.036 | *sdt* |
| response to electrical stimulus | GO:BP | 0.036 | *Appl* |
| regulation of hemocyte proliferation | GO:BP | 0.036 | *cpo,fz2,Wnt4* |
| negative regulation of presynaptic active zone assembly | GO:BP | 0.036 | *Spn* |
| positive regulation of fat cell proliferation | GO:BP | 0.036 | *InR* |
| positive regulation of border follicle cell migration | GO:BP | 0.036 | *kis,bsk,InR* |
| target-directed miRNA degradation | GO:BP | 0.036 | *Dora* |
| negative regulation of homophilic cell adhesion | GO:BP | 0.036 | *Lar* |
| ventral cord development | GO:BP | 0.036 | *brat,Antp,Galphao,Dscam1* |
| synaptic vesicle tethering involved in synaptic vesicle exocytosis | GO:BP | 0.036 | *unc-13* |
| stem cell development | GO:BP | 0.036 | *Rbp6* |
| dosage compensation complex assembly | GO:BP | 0.036 | *Unr* |
| positive regulation of receptor localization to synapse | GO:BP | 0.036 | *kis* |
| positive regulation of calcium ion transport into cytosol | GO:BP | 0.036 | *cac* |
| suture of dorsal opening | GO:BP | 0.036 | *ena* |
| dorsal vessel aortic cell fate commitment | GO:BP | 0.036 | *Ubx* |
| regulation of glutamate receptor clustering | GO:BP | 0.036 | *Rab2* |
| JUN phosphorylation | GO:BP | 0.036 | *bsk* |
| negative regulation of neuron remodeling | GO:BP | 0.036 | *Fas2* |
| positive regulation of synaptic assembly at neuromuscular junction | GO:BP | 0.038 | *cac,Ank2,mnb* |
| synaptic target inhibition | GO:BP | 0.041 | *fz2,Wnt4* |
| membrane fusion | GO:BP | 0.042 | *nSyb,brp,aus,Syx1A* |
| adult locomotory behavior | GO:BP | 0.044 | *cac,brp,shep,Atpalpha* |
| negative regulation of synaptic assembly at neuromuscular junction | GO:BP | 0.044 | *Liprin-gamma,Galphao,ena* |
| olfactory learning | GO:BP | 0.044 | *Fas2,mnb,shi* |
| determination of adult lifespan | GO:BP | 0.049 | *miple1,14-3-3epsilon,sm,Atpalpha,kis,InR* |
| positive regulation of actin filament polymerization | GO:BP | 0.049 | *chinmo,shi* |
| imaginal disc-derived male genitalia morphogenesis | GO:BP | 0.049 | *bsk,Fas2* |
| protein binding | GO:MF | 1.97e-11 | *Appl,stai,X11Lbeta,Ggamma30A,nSyb,miple1,Liprin-gamma,loaf,Rbp,Df31,14-3-3epsilon,chn,cpx,bru3,Cam,brat,fax,CadN,Gbeta13F,chinmo,Sema2b,br,wech,Antp,ctp,Ank2,Dsp1,Galphao,fz2,unc-13,Ten-a,Sumo,Ssdp,Spn,Ptp99A,sbb,tutl,His3.3B,tei,Trim9,Ubx,kek1,Lar,stx,kis,fs(1)h,bsk,abd-A,gro,Prosap,His4r,Wnt4,His3.3A,Unr,sdt,Nrg,Sema1a,caps,CtBP,tou,REPTOR,CG32264,CaMKII,mei-P26,Fas2,SKIP,InR,Dscam1,Rab2,Gug,osa,CG13917,hth,rin,mnb,CanA-14F,RhoGAP19D,CG32767,ena,Ggamma1,crp,shi,Tlk,Syx1A,tsh,psq,CkIIalpha* |
| mRNA binding | GO:MF | 2.77e-07 | *fne,bru3,Imp,brat,sm,shep,cpo,elav,Hrb98DE,Rbp6,msi,spen,Unr,nonA,Rbp1-like,rin,SC35* |
| nucleic acid binding | GO:MF | 1.42e-06 | *fne,scrt,trv,chn,bru3,Imp,brat,sm,shep,cpo,HmgD,l(3)neo38,br,Antp,elav,zfh2,Tet,Pep,Dsp1,CG31140,Ssdp,HmgZ,His3.3B,Hrb98DE,Ubx,Rbp6,kis,fs(1)h,msi,abd-A,His4r,spen,His3.3A,Unr,tou,REPTOR,ct,Rsf1,nonA,D1,osa,Rbp1-like,hth,rin,CG32767,B52,crp,tsh,psq,SC35* |
| poly(U) RNA binding | GO:MF | 1.48e-04 | *fne,shep,elav,Rbp6* |
| protein dimerization activity | GO:MF | 2.87e-04 | *Ggamma30A,Liprin-gamma,14-3-3epsilon,bru3,CadN,ctp,Ten-a,His3.3B,His4r,His3.3A,CtBP,REPTOR,Dscam1,hth,crp,psq* |
| protein domain specific binding | GO:MF | 3.42e-04 | *ctp,fz2,Ubx,Lar,gro,InR,ena,psq* |
| single-stranded RNA binding | GO:MF | 3.42e-04 | *fne,shep,elav,Pep,Hrb98DE,Rbp6* |
| translation repressor activity | GO:MF | 3.42e-04 | *brat,wech,elav,msi,Unr* |
| DNA binding | GO:MF | 4.31e-04 | *scrt,chn,HmgD,l(3)neo38,br,Antp,zfh2,Tet,Pep,Dsp1,Ssdp,HmgZ,His3.3B,Hrb98DE,Ubx,kis,fs(1)h,abd-A,His4r,His3.3A,tou,REPTOR,ct,D1,osa,hth,CG32767,crp,tsh,psq* |
| poly-pyrimidine tract binding | GO:MF | 4.69e-04 | *fne,shep,elav,Rbp6* |
| sequence-specific DNA binding | GO:MF | 0.001 | *scrt,chn,HmgD,l(3)neo38,br,Antp,zfh2,Ssdp,HmgZ,Hrb98DE,Ubx,fs(1)h,abd-A,REPTOR,ct,D1,hth,CG32767,crp,psq* |
| transcription regulator activity | GO:MF | 0.001 | *scrt,chn,l(3)neo38,br,Antp,zfh2,Pits,sbb,Ubx,fs(1)h,abd-A,gro,CtBP,REPTOR,ct,Gug,mam,hth,mnb,CG32767,crp,tsh* |
| DNA binding, bending | GO:MF | 0.002 | *HmgD,Dsp1,HmgZ* |
| transcription factor binding | GO:MF | 0.002 | *14-3-3epsilon,Dsp1,Ubx,abd-A,gro,CtBP,tou,Gug,hth* |
| transcription coregulator binding | GO:MF | 0.002 | *14-3-3epsilon,Ubx,CtBP,hth* |
| protein homodimerization activity | GO:MF | 0.002 | *Liprin-gamma,bru3,CadN,ctp,Ten-a,CtBP,Dscam1,psq* |
| heparin binding | GO:MF | 0.002 | *Appl,miple1,Lar,Sema1a* |
| RNA binding | GO:MF | 0.002 | *fne,trv,bru3,Imp,brat,sm,shep,cpo,elav,Pep,Hrb98DE,Rbp6,msi,spen,Unr,Rsf1,nonA,Rbp1-like,rin,B52,SC35* |
| mRNA 3'-UTR binding | GO:MF | 0.004 | *Imp,brat,shep,Hrb98DE,Unr* |
| protein-macromolecule adaptor activity | GO:MF | 0.004 | *nSyb,Gbeta13F,wech,Pits,sbb,gro,Prosap,Dora,CtBP,Gug,mam,mnb,Syx1A* |
| sequence-specific double-stranded DNA binding | GO:MF | 0.006 | *scrt,chn,HmgD,l(3)neo38,br,Antp,zfh2,HmgZ,Ubx,fs(1)h,abd-A,REPTOR,ct,D1,hth,CG32767,crp* |
| cytochrome-c oxidase activity | GO:MF | 0.006 | *mt:CoII,mt:CoIII* |
| transcription corepressor activity | GO:MF | 0.007 | *Pits,sbb,gro,CtBP,Gug* |
| molecular adaptor activity | GO:MF | 0.007 | *nSyb,Gbeta13F,wech,Pits,sbb,gro,Prosap,Dora,CtBP,Gug,mam,mnb,Syx1A* |
| transcription cis-regulatory region binding | GO:MF | 0.008 | *scrt,chn,HmgD,l(3)neo38,br,Antp,zfh2,HmgZ,Ubx,fs(1)h,abd-A,REPTOR,ct,hth,CG32767,crp* |
| minor groove of adenine-thymine-rich DNA binding | GO:MF | 0.008 | *HmgD,D1* |
| identical protein binding | GO:MF | 0.008 | *Liprin-gamma,bru3,CadN,ctp,Ten-a,CtBP,InR,Dscam1,psq* |
| G-protein beta-subunit binding | GO:MF | 0.008 | *Ggamma30A,Ggamma1* |
| DNA-binding transcription factor binding | GO:MF | 0.008 | *Dsp1,abd-A,gro,CtBP,tou,Gug* |
| double-stranded DNA binding | GO:MF | 0.01 | *scrt,chn,HmgD,l(3)neo38,br,Antp,zfh2,HmgZ,Ubx,fs(1)h,abd-A,REPTOR,ct,D1,hth,CG32767,crp* |
| mRNA regulatory element binding translation repressor activity | GO:MF | 0.011 | *elav,msi,Unr* |
| chemorepellent activity | GO:MF | 0.013 | *Sema2b,Sema1a* |
| glycosaminoglycan binding | GO:MF | 0.014 | *Appl,miple1,Lar,Sema1a* |
| cell adhesion molecule binding | GO:MF | 0.015 | *CadN,tei,Nrg,Fas2,Dscam1* |
| RNA polymerase II transcription regulatory region sequence-specific DNA binding | GO:MF | 0.018 | *scrt,chn,l(3)neo38,br,Antp,zfh2,Ubx,fs(1)h,abd-A,REPTOR,ct,hth,CG32767,crp* |
| DNA-binding transcription factor activity | GO:MF | 0.021 | *scrt,chn,l(3)neo38,br,Antp,zfh2,Ubx,fs(1)h,abd-A,REPTOR,ct,hth,CG32767,crp,tsh* |
| SNARE binding | GO:MF | 0.025 | *nSyb,cpx,unc-13,Syx1A* |
| protein heterodimerization activity | GO:MF | 0.027 | *Ggamma30A,14-3-3epsilon,Ten-a,His3.3B,His4r,His3.3A,REPTOR,hth* |
| syntaxin binding | GO:MF | 0.03 | *nSyb,cpx,unc-13* |
| DNA-binding transcription factor activity, RNA polymerase II-specific | GO:MF | 0.043 | *scrt,chn,l(3)neo38,Antp,zfh2,Ubx,abd-A,REPTOR,ct,hth,CG32767,crp,tsh* |
| semaphorin receptor binding | GO:MF | 0.043 | *Sema2b,Sema1a* |
| DNA-binding transcription repressor activity | GO:MF | 0.043 | *chn,br,CG32767,tsh* |
| signaling receptor binding | GO:MF | 0.044 | *miple1,Sema2b,Galphao,Spn,kek1,Lar,Prosap,Wnt4,Sema1a* |
| signaling receptor complex adaptor activity | GO:MF | 0.049 | *Gbeta13F,Prosap* |

### 1.9. Gene Ontology Enrichment Analysis: midgut

Significant GO terms (p < 0.05) with associated genes

| GO Term | Source | P-value | Genes |
| --- | --- | --- | --- |
| proton transmembrane transport | GO:BP | 4.76e-27 | *ATPsynC,Vha13,ATPsynF,Vha16-1,ATPsynbeta,Vha68-2,blw,ATPsynE,ATPsynO,ND-MLRQ,ATPsynG,ATPsynCF6,levy,ATPsynD,RFeSP,sun,ATPsynB,Vha55,ATPsyngamma,Vha26,Cyt-c1,Vha36-1,VhaAC39-1,Vha44,VhaM9.7-b,Vha14-1* |
| mitochondrial electron transport, NADH to ubiquinone | GO:BP | 2.14e-22 | *CG9034,ND-B14.5B,ND-B15,ND-B12,ND-PDSW,ND-B22,ND-24,ND-SGDH,ND-B16.6,ND-30,ND-B17,ND-18,ND-B18,ND-19,ND-13B,ND-MWFE,ND-B14,ND-15,ND-B14.7* |
| nucleoside triphosphate biosynthetic process | GO:BP | 3.70e-18 | *ATPsynC,ATPsynF,ATPsynbeta,Vha68-2,blw,ATPsynE,ATPsynO,ATPsynG,ATPsynCF6,ATPsynD,sun,ATPsyndelta,ATPsynB,sesB,awd,ATPsyngamma* |
| mitochondrial respiratory chain complex I assembly | GO:BP | 2.16e-14 | *ND-B14.5B,ND-B15,ND-B12,ND-PDSW,ND-B22,ND-SGDH,ND-B16.6,ND-18,ND-B18,ND-19,ND-B14,ND-15,ND-B14.7* |
| mitochondrial electron transport, ubiquinol to cytochrome c | GO:BP | 6.00e-13 | *UQCR-14,ox,RFeSP,UQCR-C2,UQCR-11,UQCR-Q,Cyt-c1,UQCR-6.4,Cyt-c-p* |
| mitochondrial electron transport, cytochrome c to oxygen | GO:BP | 1.98e-12 | *CG7630,cype,COX8,COX4,COX7A,COX5B,COX5A,levy,COX7C,Cyt-c-p* |
| mitochondrial respirasome assembly | GO:BP | 2.11e-12 | *COX6B,COX7A,ND-B14.5B,ND-B15,ND-B12,ND-PDSW,ND-B22,ND-SGDH,ND-B16.6,ND-18,ND-B18,ND-19,ND-B14,ND-15,ND-B14.7* |
| ribose phosphate metabolic process | GO:BP | 1.38e-11 | *ATPsynC,ATPsynF,ATPsynbeta,Vha68-2,blw,ATPsynE,ATPsynO,ATPsynG,ATPsynCF6,ATPsynD,sun,ATPsyndelta,ATPsynB,sesB,Vha55,awd,ATPsyngamma,Ak2* |
| proton motive force-driven mitochondrial ATP synthesis | GO:BP | 4.32e-05 | *ATPsynF,ATPsynbeta,ATPsynO,sun* |
| smooth septate junction assembly | GO:BP | 4.85e-05 | *Tsp2A,hoka,mesh* |
| snRNA pseudouridine synthesis | GO:BP | 4.85e-05 | *NHP2,Nop60B,CG7637* |
| septate junction assembly | GO:BP | 1.51e-04 | *Ssk,Tsp2A,hoka,mesh,l(2)gl,scrib* |
| tricarboxylic acid cycle | GO:BP | 0.003 | *SdhD,Idh3a,SdhB,SdhC,Scsalpha1* |
| establishment of endothelial barrier | GO:BP | 0.003 | *Tsp2A,hoka* |
| positive regulation of cytochrome-c oxidase activity | GO:BP | 0.003 | *ND-MLRQ,levy* |
| mitochondrial electron transport, succinate to ubiquinone | GO:BP | 0.003 | *SdhD,SdhB,SdhC* |
| sperm DNA decondensation | GO:BP | 0.004 | *P32,Nph,Nlp* |
| protein insertion into mitochondrial inner membrane | GO:BP | 0.005 | *CG34132,Tim8,CG6878* |
| vacuolar acidification | GO:BP | 0.007 | *Vha16-1,Vha55,VhaAC39-1* |
| determination of adult lifespan | GO:BP | 0.019 | *Sod1,levy,ATPsynD,sun,sesB,Trx-2,ND-SGDH,SdhC* |
| juvenile hormone mediated signaling pathway | GO:BP | 0.019 | *Fkbp39,Chd64* |
| epidermis morphogenesis | GO:BP | 0.019 | *l(2)gl,cno* |
| positive regulation of oxidoreductase activity | GO:BP | 0.019 | *ND-MLRQ,levy* |
| establishment or maintenance of polarity of larval imaginal disc epithelium | GO:BP | 0.019 | *Moe,scrib* |
| protein localization to endosome | GO:BP | 0.019 | *Moe,scrib* |
| post-translational protein targeting to membrane, translocation | GO:BP | 0.036 | *Sec61gamma,Sec61beta* |
| regulation of proton transport | GO:BP | 0.036 | *ND-MLRQ,levy* |
| SRP-dependent cotranslational protein targeting to membrane, translocation | GO:BP | 0.046 | *Sec61gamma,Sec61beta* |
| axon midline choice point recognition | GO:BP | 0.046 | *Trim9,eIF2beta,kra* |
| proton transmembrane transporter activity | GO:MF | 9.66e-29 | *ATPsynC,Vha13,ATPsynF,Vha16-1,ATPsynbeta,Vha68-2,blw,ATPsynE,ATPsynO,ATPsynG,ATPsynCF6,ATPsynD,RFeSP,sun,ATPsyndelta,ATPsynB,Vha55,ATPsyngamma,Vha26,Cyt-c1,Vha36-1,VhaAC39-1,ND-24,Vha44,ND-30,VhaM9.7-b,Vha14-1* |
| proton-transporting ATP synthase activity, rotational mechanism | GO:MF | 2.53e-20 | *ATPsynC,ATPsynF,ATPsynbeta,Vha68-2,blw,ATPsynE,ATPsynO,ATPsynG,ATPsynCF6,ATPsynD,sun,ATPsyndelta,ATPsynB,ATPsyngamma* |
| proton channel activity | GO:MF | 1.94e-19 | *ATPsynC,ATPsynF,ATPsynbeta,Vha68-2,blw,ATPsynE,ATPsynO,ATPsynG,ATPsynCF6,ATPsynD,sun,ATPsyndelta,ATPsynB,ATPsyngamma* |
| inorganic cation transmembrane transporter activity | GO:MF | 2.43e-16 | *Lfg,ATPsynC,Vha13,ATPsynF,Vha16-1,ATPsynbeta,Vha68-2,blw,ATPsynE,ATPsynO,ATPsynG,ATPsynCF6,ATPsynD,RFeSP,sun,ATPsyndelta,ATPsynB,Vha55,ATPsyngamma,Vha26,Cyt-c1,Vha36-1,VhaAC39-1,ND-24,Vha44,ND-30,VhaM9.7-b,Vha14-1* |
| monoatomic cation transmembrane transporter activity | GO:MF | 2.14e-15 | *Lfg,ATPsynC,Vha13,ATPsynF,Vha16-1,ATPsynbeta,Vha68-2,blw,ATPsynE,ATPsynO,ATPsynG,ATPsynCF6,ATPsynD,RFeSP,sun,ATPsyndelta,ATPsynB,Vha55,ATPsyngamma,Vha26,Cyt-c1,Vha36-1,VhaAC39-1,ND-24,Vha44,ND-30,VhaM9.7-b,Vha14-1* |
| monoatomic ion transmembrane transporter activity | GO:MF | 2.44e-13 | *Lfg,ATPsynC,Vha13,ATPsynF,Vha16-1,ATPsynbeta,Vha68-2,blw,ATPsynE,ATPsynO,porin,ATPsynG,ATPsynCF6,ATPsynD,RFeSP,sun,ATPsyndelta,ATPsynB,Vha55,ATPsyngamma,Vha26,Cyt-c1,Vha36-1,VhaAC39-1,ND-24,Vha44,ND-30,VhaM9.7-b,Vha14-1* |
| proton-transporting ATPase activity, rotational mechanism | GO:MF | 1.49e-12 | *Vha13,Vha16-1,ATPsynbeta,Vha68-2,Vha55,Vha26,Vha36-1,VhaAC39-1,Vha44,VhaM9.7-b,Vha14-1* |
| ATPase-coupled monoatomic cation transmembrane transporter activity | GO:MF | 2.51e-11 | *Vha13,Vha16-1,ATPsynbeta,Vha68-2,Vha55,Vha26,Vha36-1,VhaAC39-1,Vha44,VhaM9.7-b,Vha14-1* |
| oxidoreductase activity | GO:MF | 2.95e-11 | *CG3699,Gdh,COX6B,ND-MLRQ,cype,COX8,Sod1,COX7A,COX5A,RFeSP,Trx-2,Sod2,ND-B14.5B,ND-B15,ND-B12,Cyt-c1,ND-PDSW,ND-B22,SdhD,Idh3a,SdhB,ND-24,ND-SGDH,SdhC,ND-30,ND-B17,ND-B18,ND-19,ND-13B,scu,ND-B14,GstD1* |
| ligase activity | GO:MF | 3.78e-10 | *ATPsynC,ATPsynF,ATPsynbeta,Vha68-2,blw,ATPsynE,ATPsynO,ATPsynG,ATPsynCF6,ATPsynD,sun,ATPsyndelta,ATPsynB,ATPsyngamma,Gclm,Scsalpha1* |
| transporter activity | GO:MF | 4.74e-09 | *Lfg,ATPsynC,Vha13,ATPsynF,Vha16-1,ATPsynbeta,Vha68-2,blw,ATPsynE,ATPsynO,porin,ATPsynG,ATPsynCF6,ATPsynD,RFeSP,sun,ATPsyndelta,Tim8,ATPsynB,sesB,Vha55,ATPsyngamma,Vha26,Cyt-c1,Vha36-1,Sec61gamma,VhaAC39-1,ND-24,Vha44,ND-30,VhaM9.7-b,Tom7,Vha14-1* |
| transmembrane transporter activity | GO:MF | 4.74e-09 | *Lfg,ATPsynC,Vha13,ATPsynF,Vha16-1,ATPsynbeta,Vha68-2,blw,ATPsynE,ATPsynO,porin,ATPsynG,ATPsynCF6,ATPsynD,RFeSP,sun,ATPsyndelta,ATPsynB,sesB,Vha55,ATPsyngamma,Vha26,Cyt-c1,Vha36-1,Sec61gamma,VhaAC39-1,ND-24,Vha44,ND-30,VhaM9.7-b,Tom7,Vha14-1* |
| monoatomic cation channel activity | GO:MF | 2.26e-08 | *Lfg,ATPsynC,ATPsynF,ATPsynbeta,Vha68-2,blw,ATPsynE,ATPsynO,ATPsynG,ATPsynCF6,ATPsynD,sun,ATPsyndelta,ATPsynB,ATPsyngamma* |
| electron transfer activity | GO:MF | 1.65e-07 | *RFeSP,Cyt-c1,SdhB,ND-24,SdhC,ND-30,Cyt-c-p,Etfb* |
| ATPase-coupled transmembrane transporter activity | GO:MF | 1.65e-07 | *Vha13,Vha16-1,ATPsynbeta,Vha68-2,Vha55,Vha26,Vha36-1,VhaAC39-1,Vha44,VhaM9.7-b,Vha14-1* |
| active transmembrane transporter activity | GO:MF | 4.47e-07 | *Vha13,Vha16-1,ATPsynbeta,Vha68-2,RFeSP,sesB,Vha55,Vha26,Cyt-c1,Vha36-1,VhaAC39-1,ND-24,Vha44,ND-30,VhaM9.7-b,Vha14-1* |
| monoatomic ion channel activity | GO:MF | 9.31e-07 | *Lfg,ATPsynC,ATPsynF,ATPsynbeta,Vha68-2,blw,ATPsynE,ATPsynO,porin,ATPsynG,ATPsynCF6,ATPsynD,sun,ATPsyndelta,ATPsynB,ATPsyngamma* |
| channel activity | GO:MF | 3.67e-06 | *Lfg,ATPsynC,ATPsynF,ATPsynbeta,Vha68-2,blw,ATPsynE,ATPsynO,porin,ATPsynG,ATPsynCF6,ATPsynD,sun,ATPsyndelta,ATPsynB,ATPsyngamma* |
| succinate dehydrogenase activity | GO:MF | 5.57e-04 | *SdhD,SdhB,SdhC* |
| juvenile hormone response element binding | GO:MF | 0.001 | *Fkbp39,Chd64* |
| box H/ACA snoRNA binding | GO:MF | 0.004 | *NHP2,CG7637* |
| catalytic activity | GO:MF | 0.006 | *CG3699,ATPsynC,Gdh,St3,ATPsynF,ATPsynbeta,Vha68-2,Cyp1,blw,Cndp2,Hsp60A,COX6B,ATPsynE,Hsc70-4,ATPsynO,ND-MLRQ,cype,COX8,Sod1,ATPsynG,COX7A,Fkbp39,ATPsynCF6,COX5A,ATPsynD,RFeSP,Trim9,Pgant9,sun,ATPsyndelta,ATPsynB,awd,Trx-2,ATPsyngamma,Tina-1,Sod2,ND-B14.5B,ND-B15,ND-B12,GstE12,Cyt-c1,ND-PDSW,Fkbp12,ND-B22,Nop60B,SdhD,Idh3a,SdhB,ND-24,ND-SGDH,SdhC,Gclm,ND-30,ND-B17,UQCR-C1,Ak2,ND-B18,ND-19,ND-13B,p23,scu,ND-B14,RNASEK,GstD1,Scsalpha1* |
| ubiquinol-cytochrome-c reductase activity | GO:MF | 0.007 | *RFeSP,Cyt-c1* |
| ubiquinone binding | GO:MF | 0.007 | *SdhD,SdhB* |
| snoRNA binding | GO:MF | 0.011 | *NHP2,Nop56,CG7637* |
| succinate dehydrogenase (quinone) activity | GO:MF | 0.015 | *SdhD,SdhB* |
| superoxide dismutase activity | GO:MF | 0.015 | *Sod1,Sod2* |
| 2 iron, 2 sulfur cluster binding | GO:MF | 0.023 | *RFeSP,SdhB,ND-24* |
| protein-folding chaperone binding | GO:MF | 0.024 | *CG11267,Hsp60A,Hsc70-4,p23* |
| quinone binding | GO:MF | 0.039 | *SdhD,SdhB* |
| protein transporter activity | GO:MF | 0.041 | *Tim8,Sec61gamma,Tom7* |
| peptidyl-prolyl cis-trans isomerase activity | GO:MF | 0.041 | *Cyp1,Fkbp39,Fkbp12* |
| ATP-dependent protein folding chaperone | GO:MF | 0.043 | *CG11267,Hsp60A,Hsc70-4* |

### 1.10. Gene Ontology Enrichment Analysis: lymphgland

Significant GO terms (p < 0.05) with associated genes

| GO Term | Source | P-value | Genes |
| --- | --- | --- | --- |
| nucleobase-containing compound metabolic process | GO:BP | 1.57e-30 | *ham,kn,PCNA,dUTPase,Myc,jumu,srp,geminin,odd,Slbp,RnrS,RPA3,RPA2,Tgi,Chrac-14,Caf1-180,WRNexo,CG13096,Su(var)205,Mcm7,HipHop,Caf1-55,Ssrp,RnrL,N,hoip,RPA1,NHP2,jigr1,zfh1,rgn,dpa,Acf,Dek,Gnf1,dnk,nop5,Nop60B,Fib,Pde1c,Ndf,SmE,SmD1,Cdk1,SmD3,mtSSB,CG33217,SmB,Dp,CG8891,Uba2,La,dup,CG6712,CG7637,x16,CG1234,CycE,Rbm13,CG9107,Polr2F,REG,CG1542,RpLP0-like,CG42458,Polr2E,Nop56,gro,CG2199,HDAC1,CG32409,srl,Sem1,Nup358,His2Av,CG11583,Aos1,CG30122,Cwc25,SNRPG,CG13097,Polr2L,rgr,AdSS,drm,Rrp4,HmgD,XNP,CG11858,Hel25E,CG8545,Nopp140,CG32344,Polr1D,CG1142,Map60,CG3527,Polr2K,pit,Non3,CG12391,PAN3,CG1789,Nap1,Prp39,Polr2J,Snr1,U2A,ras,su(f),CG6937,CG10418,Cbp20,Ssb-c31a,CG30349,SmF,l(1)10Bb,Shmt,HmgZ,sqd,awd,Brd7-9,CG2034,sno,l(3)mbt,Sumo,Mtor,lin-28,MED22,Rrp47,mod,BEAF-32,CG12288,CG4866,MrgBP,Nnp-1,Sbat,Paics,Tis11,Prps,Pde9,CG4038,SmD2,dom,Samuel,RnpS1,LSm7,caz,Non2,CG4360,mago,ytr,rump,Hlc,apt,Pop5,mod(mdg4),Nurf-38,Taf13,CG9776,Cmpk,TfIIS,alpha-PheRS,ps,Ak2,kay,Polr2H,crp,CG4364,Acn,AspRS,e(r),Mes2,Atf3,Rpp25,Hsc70-4,Sf3b2,hang,SerRS,spoon,Cdk12,CG1677,tou,Rbp1,E(Pc),fs(1)h,Polr2A,FoxK,Xrp1,SC35,sov,lwr,Myd88,Caper,Psc,HDAC4,Sarm,Dg,Eip78C* |
| regulation of metabolic process | GO:BP | 6.18e-12 | *ham,kn,PCNA,Myc,jumu,srp,geminin,odd,Tgi,Chrac-14,Fkbp39,WRNexo,Su(var)205,Mcm7,Claspin,Caf1-55,Ssrp,N,hoip,jigr1,zfh1,PGRP-LC,dpa,Acf,Dek,Ndf,Dph5,Cdk1,mtSSB,Dp,Fdh,Uba2,La,dup,x16,CycE,Spn27A,REG,RpLP0-like,CG42458,gro,CG2199,HDAC1,srl,wds,Nup358,CycA,Aos1,Sps1,rgr,drm,Rrp4,HmgD,XNP,Hel25E,Map60,l(1)G0004,CG12391,PAN3,Prp39,Su(var)2-HP2,ex,Snr1,CG10418,Cbp20,Ssb-c31a,Shmt,HmgZ,sqd,Brd7-9,sno,l(3)mbt,Sumo,Mtor,lin-28,MED22,Rrp47,BEAF-32,MrgBP,Tis11,dom,Nedd8,Samuel,p23,lig,LSm7,caz,Non2,CG4360,Egfr,mago,rump,dod,scny,apt,mod(mdg4),Nurf-38,Dd,CG9776,CNBP,dnr1,eIF2gamma,ps,kay,Dcp-1,crp,Acn,Mes2,Atf3,raptor,msi,Hsc70-4,hang,Lk6,spoon,CG6878,Cdk12,CG1677,tou,Rbp1,numb,E(Pc),fs(1)h,FoxK,SC35,sov,SCAP,lwr,Myd88,CG32369,Caper,Psc,l(3)80Fj,Pomp,PRAS40,HDAC4,Pitslre,Pax,Dg,Eip78C* |
| RNA splicing, via transesterification reactions | GO:BP | 2.09e-09 | *hoip,SmE,SmD1,SmD3,SmB,x16,REG,CG30122,Cwc25,SNRPG,Hel25E,Prp39,U2A,CG10418,Cbp20,SmF,l(1)10Bb,sqd,mod,SmD2,dom,RnpS1,LSm7,caz,mago,ytr,ps,Acn,Hsc70-4,Sf3b2,CG1677,Rbp1,SC35,Caper* |
| regulation of RNA metabolic process | GO:BP | 2.07e-05 | *ham,kn,Myc,jumu,srp,odd,Tgi,Su(var)205,Caf1-55,N,jigr1,zfh1,Acf,Ndf,Cdk1,Dp,Uba2,x16,CG42458,gro,CG2199,HDAC1,srl,Aos1,rgr,drm,HmgD,Hel25E,Map60,CG12391,PAN3,Prp39,Snr1,CG10418,Ssb-c31a,HmgZ,sqd,Brd7-9,sno,l(3)mbt,Sumo,Mtor,lin-28,MED22,BEAF-32,MrgBP,Tis11,dom,Samuel,caz,Non2,CG4360,rump,apt,mod(mdg4),Nurf-38,CG9776,ps,kay,crp,Mes2,Atf3,hang,Cdk12,CG1677,tou,Rbp1,E(Pc),fs(1)h,FoxK,SC35,sov,lwr,Myd88,Caper,Psc,HDAC4,Eip78C* |
| tRNA transcription by RNA polymerase III | GO:BP | 1.36e-04 | *Polr2F,Polr2E,Polr2L,Polr1D,Polr2K,Polr2H* |
| negative regulation of biosynthetic process | GO:BP | 3.05e-04 | *srp,odd,Tgi,Chrac-14,Su(var)205,Caf1-55,N,zfh1,Acf,Spn27A,RpLP0-like,gro,CG2199,HDAC1,Nup358,drm,XNP,PAN3,Su(var)2-HP2,Cbp20,Ssb-c31a,Shmt,sqd,l(3)mbt,Mtor,lin-28,Tis11,dom,Samuel,p23,LSm7,CG4360,Egfr,rump,scny,apt,CG9776,dnr1,msi,Hsc70-4,numb,fs(1)h,FoxK,sov,lwr,Psc,HDAC4,Eip78C* |
| spliceosomal snRNP assembly | GO:BP | 8.73e-04 | *SmE,SmD1,SmD3,SNRPG,SmF,SmD2* |
| snRNA pseudouridine synthesis | GO:BP | 0.001 | *NHP2,Nop60B,CG7637* |
| negative regulation melanotic encapsulation of foreign target | GO:BP | 0.001 | *Uba2,Spn27A,lwr* |
| deoxyribonucleotide biosynthetic process | GO:BP | 0.001 | *dUTPase,RnrS,RnrL,dnk* |
| ribosomal large subunit export from nucleus | GO:BP | 0.001 | *emb,CG32409,Mys45A,Ns2* |
| eggshell chorion gene amplification | GO:BP | 0.001 | *PCNA,geminin,Caf1-55,Dp,dup,CycE* |
| negative regulation of signal transduction | GO:BP | 0.002 | *Myc,Uba2,gro,HDAC1,srl,His2Av,ex,Nedd8,lig,nmo,trol,Egfr,scny,apt,Dd,dnr1,Dcp-1,hang,numb,Patronin,lwr,PRAS40,raw,kek5,RapGAP1,Ptp4E,Sarm,pigs,hppy,nkd* |
| maturation of LSU-rRNA from tricistronic rRNA transcript (SSU-rRNA, 5.8S rRNA, LSU-rRNA) | GO:BP | 0.002 | *pit,Non3,CG6937,CG12288,CG4364* |
| germ cell migration | GO:BP | 0.003 | *srp,zfh1,mim,Tre1,mod(mdg4),lwr,ttv* |
| imaginal disc-derived wing vein specification | GO:BP | 0.003 | *kn,N,msk,Snr1,nmo,Egfr,Dd,Dys,Dg* |
| regulation of DNA-templated DNA replication initiation | GO:BP | 0.003 | *Mcm7,dpa,Cdk1,dup* |
| nuclear pore complex assembly | GO:BP | 0.003 | *Ran,emb,Nup358* |
| sperm DNA decondensation | GO:BP | 0.004 | *Nph,P32,Nlp,Nap1* |
| antimicrobial humoral immune response mediated by antimicrobial peptide | GO:BP | 0.004 | *PGRP-LC,Uba2,Aos1,Sumo,scny,dnr1,kay,lwr,Myd88,Pitslre* |
| DNA unwinding involved in DNA replication | GO:BP | 0.006 | *Mcm7,RPA1,dpa,mtSSB* |
| somatic stem cell population maintenance | GO:BP | 0.007 | *zfh1,CycE,His2Av,dom,scny* |
| protein peptidyl-prolyl isomerization | GO:BP | 0.009 | *CG14715,Fkbp59,CG2852,Cyp1,Fkbp12* |
| regulation of alternative mRNA splicing, via spliceosome | GO:BP | 0.009 | *x16,Hel25E,Prp39,CG10418,sqd,dom,ps,Rbp1,SC35,Caper* |
| second mitotic wave involved in compound eye morphogenesis | GO:BP | 0.01 | *N,Egfr,kay* |
| ventral cord development | GO:BP | 0.01 | *kn,Rcc1,glu,Surf6,CycE,Nup358,18w,Galphai,Psc* |
| regulation of transcriptional start site selection at RNA polymerase II promoter | GO:BP | 0.01 | *x16,Rbp1,SC35* |
| cell surface receptor signaling pathway | GO:BP | 0.011 | *ham,Myc,jumu,pyr,N,PGRP-LC,Swim,mew,Uba2,CycE,gro,Uch-L5,HDAC1,srl,Aos1,Gp150,sno,Sumo,lin-28,Nedd8,lig,nmo,trol,Egfr,apt,Dd,Dcp-1,raptor,numb,unc-5,lwr,Myd88,ttv,PRAS40,HDAC4,kek5,Pitslre,robo1,Ptp4E,Sarm,pigs,hppy,nkd* |
| dendrite guidance | GO:BP | 0.014 | *HDAC1,Snr1,dom,E(Pc),robo1* |
| snoRNA guided rRNA pseudouridine synthesis | GO:BP | 0.014 | *CG7637,CG4038* |
| energy homeostasis | GO:BP | 0.014 | *srl,ex* |
| DNA replication-dependent chromatin assembly | GO:BP | 0.015 | *Caf1-55,Acf,Nap1* |
| regulation of mRNA 3'-end processing | GO:BP | 0.015 | *x16,Rbp1,SC35* |
| animal organ morphogenesis | GO:BP | 0.017 | *kn,jumu,srp,odd,N,hoip,mew,stg,Pura,CycE,drm,msk,ex,Snr1,sno,SelR,nmo,trol,Egfr,Dd,CNBP,dnr1,app,RhoGAP15B,betaTub97EF,Ns2,kay,msi,kmr,numb,unc-5,CG43658,CadN,raw,rst,robo1,klar,kirre,Dys,Dg,Cdep,shot* |
| positive regulation of DNA replication | GO:BP | 0.018 | *PCNA,Acf,mtSSB,spoon* |
| ribosomal large subunit assembly | GO:BP | 0.022 | *RpLP0-like,CG11583,Non3,mRpL20* |
| protein sumoylation | GO:BP | 0.022 | *Uba2,Aos1,Sumo,lwr* |
| larval visceral muscle development | GO:BP | 0.026 | *pyr,klar,kirre* |
| DNA replication initiation | GO:BP | 0.028 | *Mcm7,dpa,Cdk1,dup,CG1234* |
| nucleoside triphosphate catabolic process | GO:BP | 0.029 | *dUTPase,CG8891* |
| regulation of glycolytic process | GO:BP | 0.029 | *N,Dg* |
| cortical microtubule organization | GO:BP | 0.029 | *Patronin,shot* |
| neuroblast fate specification | GO:BP | 0.029 | *N,nkd* |
| ommatidial rotation | GO:BP | 0.031 | *N,msk,nmo,Egfr,CadN* |
| negative regulation of Toll signaling pathway | GO:BP | 0.032 | *gro,Dcp-1,lwr* |
| regulation of exit from mitosis | GO:BP | 0.032 | *CycE,CycA,ex* |
| neuronal stem cell population maintenance | GO:BP | 0.032 | *RPA3,N,Snr1* |
| axonal fasciculation | GO:BP | 0.034 | *Fas3,zfh1,Hsc70-4,CadN* |
| negative regulation of cell population proliferation | GO:BP | 0.034 | *ham,ex,Snr1,l(3)mbt,dom,numb,Xrp1,lwr* |
| ribose phosphate metabolic process | GO:BP | 0.036 | *N,dnk,Pde1c,AdSS,ras,awd,Paics,Prps,Pde9,Cmpk,Ak2,Dg* |
| positive regulation of transcription by RNA polymerase II | GO:BP | 0.037 | *ham,kn,Myc,srp,odd,Su(var)205,N,Ndf,CG2199,srl,rgr,drm,Snr1,Ssb-c31a,sno,Mtor,apt,Nurf-38,kay,Cdk12,tou,fs(1)h,sov,Myd88* |
| muscle cell cellular homeostasis | GO:BP | 0.038 | *N,zfh1,robo1,Dys,Dg* |
| positive regulation of Toll signaling pathway | GO:BP | 0.039 | *Uba2,Aos1,Sumo,Pitslre* |
| NLS-bearing protein import into nucleus | GO:BP | 0.04 | *Rcc1,Nup358,Fs(2)Ket* |
| regulation of myoblast fusion | GO:BP | 0.04 | *rst,kirre,Pax* |
| protein insertion into mitochondrial inner membrane | GO:BP | 0.04 | *Tim8,CG34132,CG6878* |
| positive regulation of translation | GO:BP | 0.045 | *hoip,Dph5,La,CNBP,eIF2gamma,spoon* |
| negative regulation of peptidoglycan recognition protein signaling pathway | GO:BP | 0.046 | *Myc,His2Av,scny,dnr1,lwr* |
| GTP biosynthetic process | GO:BP | 0.047 | *ras,awd* |
| positive regulation of mitochondrial DNA replication | GO:BP | 0.047 | *mtSSB,spoon* |
| garland nephrocyte differentiation | GO:BP | 0.047 | *zfh1,kirre* |
| glial cell migration | GO:BP | 0.049 | *pyr,N,XNP,numb,unc-5* |
| nucleic acid binding | GO:MF | 1.70e-30 | *ham,kn,PCNA,Myc,jumu,srp,odd,Slbp,RPA3,RPA2,Chrac-14,Fkbp39,WRNexo,CG13096,Su(var)205,Mcm7,Caf1-55,Ssrp,Nph,hoip,RPA1,D1,NHP2,jigr1,zfh1,dpa,Acf,Dek,Gnf1,Rs1,nop5,Nop60B,Fib,Ndf,SmE,SmD1,SmD3,mtSSB,SmB,Dp,Surf6,baf,La,dup,CG9143,CG6712,CG7637,x16,CG9107,Polr2F,CG1542,CG42458,Polr2E,Nop56,CG2199,srl,His2Av,mbm,CG11583,CG11180,SNRPG,Polr2L,CG7006,drm,Rrp4,msk,HmgD,vig2,CG31301,XNP,CG11858,Hel25E,CG8545,CG32344,Polr1D,CG1142,CG3527,Polr2K,l(1)G0004,pit,Non3,CG12391,CG3918,Nlp,PAN3,Prp39,Polr2J,Su(var)2-HP2,Snr1,U2A,CG11563,su(f),CG6937,CG10418,CG4806,Cbp20,Ssb-c31a,SmF,Shmt,HmgZ,sqd,CG2034,B52,AIMP1,sno,l(3)mbt,Mtor,CG3224,lin-28,Rrp47,mod,BEAF-32,CG12288,CG4866,Sbat,Tis11,CG4038,SmD2,dom,RnpS1,eEF1delta,LSm7,caz,CG4360,mago,ytr,rump,Hlc,mRpL20,apt,His3.3A,Pop5,mod(mdg4),CNBP,mRpL16,TfIIS,eIF2gamma,Galphai,alpha-PheRS,ps,kay,crp,CG4364,Acn,AspRS,Atf3,Rpp25,msi,Sf3b2,hang,mRpS18A,SerRS,spoon,tou,Rbp1,fs(1)h,Non1,Polr2A,FoxK,Xrp1,SC35,Caper,Psc,Eip78C* |
| RNA binding | GO:MF | 1.99e-26 | *Slbp,CG13096,Su(var)205,Ssrp,Nph,hoip,NHP2,nop5,Nop60B,Fib,SmE,SmD1,SmD3,SmB,Surf6,La,CG9143,CG6712,CG7637,x16,CG1542,CG42458,Nop56,srl,CG11583,SNRPG,CG7006,Rrp4,msk,vig2,Hel25E,CG8545,CG32344,CG1142,CG3527,l(1)G0004,pit,Non3,Nlp,PAN3,Prp39,U2A,CG11563,su(f),CG6937,CG10418,CG4806,Cbp20,SmF,Shmt,sqd,CG2034,B52,AIMP1,lin-28,Rrp47,mod,CG12288,CG4866,Sbat,Tis11,CG4038,SmD2,RnpS1,LSm7,caz,mago,ytr,rump,Hlc,mRpL20,apt,Pop5,CNBP,mRpL16,eIF2gamma,alpha-PheRS,ps,CG4364,AspRS,Rpp25,msi,Sf3b2,hang,mRpS18A,SerRS,spoon,Rbp1,Non1,SC35,Caper* |
| histone binding | GO:MF | 9.85e-08 | *Caf1-180,Su(var)205,Mapmodulin,Caf1-55,Ssrp,Nph,Dek,Set,wds,P32,Nlp,Nap1,Df31,Brd7-9,sno,l(3)mbt,dom,Samuel,fs(1)h* |
| RNA polymerase II activity | GO:MF | 1.61e-07 | *Polr2F,Polr2E,Polr2L,Polr2K,Polr2J,Polr2H,Polr2A* |
| protein binding | GO:MF | 2.12e-07 | *dUTPase,Myc,srp,geminin,pyr,Slbp,RPA3,RPA2,Tgi,Chrac-14,ncd,Caf1-180,Fkbp39,Gmap,Su(var)205,HipHop,Mapmodulin,Caf1-55,Ssrp,Nph,N,Fas3,jigr1,PGRP-LC,glu,dpa,Acf,Dek,Set,Nxt1,Pde1c,Ran,SmE,Swim,Cdk1,emb,mew,Idgf6,side-V,Dp,Uba2,dup,CycE,gro,dpr17,HDAC1,CG32409,srl,wds,Sem1,Nup358,CycA,His2Av,P32,CG15019,Aos1,CG30122,CG13097,HP4,CG10565,drm,Gp150,msk,Hsp60A,Polr1D,Map60,Idgf4,Nlp,PAN3,Nap1,Prp39,Polr2J,Fkbp59,ex,Snr1,U2A,Karybeta3,su(f),CG10418,Cbp20,CG30349,SmF,Shmt,Df31,awd,Brd7-9,Ten-a,sno,l(3)mbt,Sumo,Mtor,MED22,mod,BEAF-32,Nplp2,mim,dom,Nedd8,Samuel,CAP,CG13926,p23,lig,nudC,18w,LSm7,nmo,trol,Egfr,side-IV,mago,Prosalpha2,dod,scny,apt,tutl,His3.3A,mod(mdg4),Fs(2)Ket,Taf13,CG5504,Arpc2,Prosbeta4,Prosbeta1,Phb1,Galphai,CG11267,Phb2,kay,Dcp-1,CG5515,crp,Acn,form3,Atf3,raptor,Prosbeta3,Prosbeta5,Hsc70Cb,Lrch,CCT6,Hsc70-4,Lk6,spoon,olf186-M,Cdk12,RhoGAP93B,tou,numb,fs(1)h,Prosalpha3,FoxK,Fkbp12,Nop17l,unc-5,Patronin,SCAP,lwr,Myd88,CG32369,mrj,l(3)80Fj,CadN,Tlk,ttv,rst,HDAC4,kek5,robo1,klar,Ptp4E,gukh,kirre,CG10011,Dys,Sarm,Dg,pigs,Cdep,shot,nkd* |
| mRNA binding | GO:MF | 2.09e-06 | *Slbp,Su(var)205,La,x16,srl,su(f),CG6937,CG4806,Cbp20,Shmt,sqd,lin-28,mod,CG12288,Tis11,RnpS1,caz,ytr,rump,apt,CNBP,ps,msi,hang,Rbp1,SC35,Caper* |
| DNA binding | GO:MF | 2.99e-06 | *ham,kn,PCNA,Myc,jumu,srp,odd,RPA3,RPA2,Chrac-14,Fkbp39,Su(var)205,Mcm7,Caf1-55,Ssrp,RPA1,D1,jigr1,zfh1,dpa,Acf,Dek,Gnf1,Ndf,mtSSB,Dp,Surf6,baf,La,dup,Polr2F,Polr2E,CG2199,His2Av,Polr2L,drm,HmgD,XNP,CG11858,Polr1D,Polr2K,CG12391,Polr2J,Su(var)2-HP2,Snr1,Ssb-c31a,HmgZ,sno,l(3)mbt,Mtor,Rrp47,mod,BEAF-32,Tis11,dom,CG4360,apt,His3.3A,mod(mdg4),TfIIS,Galphai,kay,crp,Atf3,hang,tou,fs(1)h,Polr2A,FoxK,Xrp1,Psc,Eip78C* |
| DNA-directed 5'-3' RNA polymerase activity | GO:MF | 6.86e-06 | *Polr2F,Polr2E,Polr2L,Polr1D,Polr2K,Polr2J,Polr2H,Polr2A* |
| RNA polymerase I activity | GO:MF | 1.15e-05 | *Polr2F,Polr2E,Polr2L,Polr1D,Polr2K,Polr2H* |
| RNA polymerase III activity | GO:MF | 1.15e-05 | *Polr2F,Polr2E,Polr2L,Polr1D,Polr2K,Polr2H* |
| rRNA binding | GO:MF | 2.06e-04 | *La,CG6712,CG11583,CG3527,Non3,CG12288,CG4866,mRpL20,mRpL16,mRpS18A* |
| peptidyl-prolyl cis-trans isomerase activity | GO:MF | 3.87e-04 | *Fkbp39,CG14715,CG11858,Fkbp59,CG2852,Cyp1,dod,Fkbp12* |
| chromatin binding | GO:MF | 4.96e-04 | *PCNA,Chrac-14,Rcc1,Su(var)205,Caf1-55,Ssrp,Nph,N,glu,Set,Ndf,CG1234,Nlp,Nap1,sno,l(3)mbt,Mtor,BEAF-32,Samuel,caz,mod(mdg4),fs(1)h,Xrp1,Psc* |
| snoRNA binding | GO:MF | 8.50e-04 | *NHP2,nop5,CG7637,Nop56,CG4866,CG4038* |
| FK506 binding | GO:MF | 9.95e-04 | *Fkbp39,CG14715,Fkbp59* |
| ubiquitin activating enzyme binding | GO:MF | 9.95e-04 | *Uba2,Aos1,lwr* |
| box H/ACA snoRNA binding | GO:MF | 9.95e-04 | *NHP2,CG7637,CG4038* |
| protein domain specific binding | GO:MF | 0.001 | *Ssrp,N,gro,ex,Snr1,mim,mod(mdg4),Dcp-1,Myd88,rst,Dys,nkd* |
| ATP binding | GO:MF | 0.007 | *ncd,Mcm7,RnrL,glu,dpa,Gnf1,dnk,Rs1,Cdk1,Uba2,CG9143,Sps1,XNP,Hel25E,CG32344,Hsp60A,pit,PAN3,awd,Paics,Prps,dom,nmo,Egfr,Hlc,CG5504,nmd,Sam-S,Cmpk,CG11267,alpha-PheRS,Ak2,AspRS,CG1703,Hsc70Cb,CCT6,Hsc70-4,SerRS,Lk6,Cdk12,numb,FoxK,lwr,Tlk,Pitslre,CG31145,hppy* |
| adenyl ribonucleotide binding | GO:MF | 0.008 | *ncd,Mcm7,RnrL,glu,dpa,Gnf1,dnk,Rs1,Cdk1,Uba2,CG9143,Sps1,XNP,Hel25E,CG32344,Hsp60A,pit,PAN3,awd,Paics,Prps,dom,nmo,Egfr,Hlc,CG5504,nmd,Sam-S,Cmpk,CG11267,alpha-PheRS,Ak2,AspRS,CG1703,Hsc70Cb,CCT6,Hsc70-4,SerRS,Lk6,Cdk12,numb,FoxK,lwr,Tlk,Pitslre,CG31145,hppy* |
| protein-folding chaperone binding | GO:MF | 0.008 | *Su(var)205,CG10565,Hsp60A,CG13926,p23,CG11267,Hsc70-4,mrj* |
| heat shock protein binding | GO:MF | 0.008 | *Su(var)205,CG10565,CG13926,p23,CG5504,Hsc70-4,lwr* |
| purine nucleotide binding | GO:MF | 0.009 | *ncd,Mcm7,RnrL,glu,dpa,Gnf1,dnk,Rs1,Ran,Ndf,Cdk1,Uba2,CG9143,Sps1,AdSS,XNP,Hel25E,CG32344,Hsp60A,pit,PAN3,awd,Paics,Prps,dom,nmo,Egfr,Hlc,CG5504,nmd,Sam-S,Cmpk,eIF2gamma,Galphai,CG11267,alpha-PheRS,betaTub97EF,Ns2,Ak2,AspRS,Ttd14,CG1703,Hsc70Cb,CCT6,Hsc70-4,SerRS,Lk6,Cdk12,numb,Non1,FoxK,lwr,Tlk,Pitslre,CG31145,hppy* |
| SUMO activating enzyme activity | GO:MF | 0.018 | *Uba2,Aos1* |
| ribonucleoside-diphosphate reductase activity, thioredoxin disulfide as acceptor | GO:MF | 0.018 | *RnrS,RnrL* |
| nucleotide binding | GO:MF | 0.027 | *ncd,Mcm7,RnrL,glu,dpa,Gnf1,dnk,Rs1,Ran,Ndf,Cdk1,CG8891,Uba2,CG9143,Sps1,AdSS,XNP,Hel25E,CG32344,Hsp60A,pit,PAN3,ras,awd,Paics,Prps,dom,nmo,Egfr,Hlc,CG5504,nmd,Sam-S,Cmpk,eIF2gamma,Galphai,CG11267,alpha-PheRS,betaTub97EF,Ns2,Ak2,AspRS,Ttd14,CG1703,Hsc70Cb,CCT6,Hsc70-4,SerRS,Lk6,Cdk12,numb,Non1,FoxK,lwr,Tlk,Pitslre,CG31145,hppy* |
| helicase activity | GO:MF | 0.03 | *Mcm7,dpa,Rs1,CG9143,XNP,Hel25E,CG32344,pit,dom,Hlc* |
| small molecule binding | GO:MF | 0.032 | *ham,kn,dUTPase,srp,odd,RnrS,ncd,WRNexo,Mcm7,RnrL,N,CG3376,RPA1,zfh1,PGRP-LC,glu,dpa,Acf,Gnf1,dnk,Rs1,Pde1c,Ran,Ndf,Cdk1,Sply,CG8891,Fdh,Uba2,CG9143,x16,CG2199,HDAC1,alpha-Man-IIb,Nup358,CG42566,mbm,Sps1,Polr2L,AdSS,drm,XNP,Hel25E,CG32344,Hsp60A,Polr2K,pit,CG12391,CG3918,PAN3,ras,Shmt,awd,l(3)mbt,CG3224,lin-28,Tim8,BEAF-32,SelR,Paics,Tis11,Prps,Pde9,dom,CG34132,und,nmo,caz,Egfr,Hlc,mod(mdg4),Nurf-38,CG9776,CNBP,CG5504,nmd,CG8036,Sam-S,dnr1,Cmpk,TfIIS,RhoGAP15B,eIF2gamma,Galphai,CG11267,alpha-PheRS,betaTub97EF,Ns2,Ak2,CG8635,AspRS,Ttd14,CG2915,CG1703,Clic,Noa36,Hsc70Cb,CCT6,Hsc70-4,hang,SerRS,Lk6,Cdk12,CG1677,tou,numb,Non1,Polr2A,FoxK,sov,lwr,Myd88,CG32369,Psc,CadN,Tlk,ttv,HDAC4,Pitslre,Cad87A,Pax,Dys,Dg,CG31145,Eip78C,shot,hppy,nkd* |
| anion binding | GO:MF | 0.032 | *ncd,Mcm7,RnrL,glu,dpa,Gnf1,dnk,Rs1,Ran,Cdk1,Sply,Uba2,CG9143,Sps1,AdSS,XNP,Hel25E,CG32344,Hsp60A,pit,PAN3,Shmt,awd,Paics,Prps,dom,nmo,Egfr,Hlc,CG5504,nmd,CG8036,Sam-S,Cmpk,RhoGAP15B,eIF2gamma,Galphai,CG11267,alpha-PheRS,betaTub97EF,Ns2,Ak2,AspRS,Ttd14,CG1703,Hsc70Cb,CCT6,Hsc70-4,SerRS,Lk6,Cdk12,numb,Non1,FoxK,lwr,Myd88,Tlk,Pitslre,CG31145,hppy* |
| carbohydrate derivative binding | GO:MF | 0.04 | *ncd,Mcm7,RnrL,PGRP-LC,glu,dpa,Gnf1,dnk,Rs1,Ran,Cdk1,Idgf6,Uba2,CG9143,Sps1,AdSS,XNP,Hel25E,CG32344,Hsp60A,pit,Idgf4,PAN3,awd,Paics,Prps,dom,nmo,Egfr,Hlc,CG5504,nmd,Sam-S,Cmpk,eIF2gamma,Galphai,CG11267,alpha-PheRS,betaTub97EF,Ns2,Ak2,AspRS,Ttd14,CG1703,Hsc70Cb,CCT6,Hsc70-4,SerRS,Lk6,Cdk12,numb,Non1,FoxK,lwr,Tlk,Pitslre,robo1,CG31145,hppy* |
| nucleotidyltransferase activity | GO:MF | 0.044 | *Polr2F,Polr2E,Polr2L,Polr1D,Polr2K,Polr2J,Polr2H,Polr2A* |
| protein folding chaperone | GO:MF | 0.046 | *Hsp60A,CG11267,Hsc70Cb,CCT6,Hsc70-4,mrj* |
| isomerase activity | GO:MF | 0.047 | *ncd,Gale,Fkbp39,Nop60B,CG14715,CG11858,Fkbp59,CG2852,Cyp1,p23,dod,Fkbp12* |

### 1.11. Gene Ontology Enrichment Analysis: unknown

Significant GO terms (p < 0.05) with associated genes

| GO Term | Source | P-value | Genes |
| --- | --- | --- | --- |
| P-type potassium transmembrane transporter activity | GO:MF | 0.032 | *Atpalpha* |
| mitogen-activated protein kinase binding | GO:MF | 0.032 | *Lk6* |
| P-type sodium:potassium-exchanging transporter activity | GO:MF | 0.032 | *Atpalpha* |
| ATPase-coupled transmembrane transporter activity | GO:MF | 0.032 | *Atpalpha,Hsc70-3* |
| protein-transporting ATPase activity | GO:MF | 0.032 | *Hsc70-3* |
| eukaryotic initiation factor 4E binding | GO:MF | 0.033 | *eIF4G1* |
| P-type transmembrane transporter activity | GO:MF | 0.042 | *Atpalpha* |
| P-type ion transporter activity | GO:MF | 0.042 | *Atpalpha* |
| calcium/calmodulin-dependent protein kinase activity | GO:MF | 0.042 | *Lk6* |
| calcium-dependent protein serine/threonine kinase activity | GO:MF | 0.042 | *Lk6* |
| gap junction channel activity | GO:MF | 0.042 | *ogre* |
| RNA 7-methylguanosine cap binding | GO:MF | 0.042 | *eIF4G1* |
| misfolded protein binding | GO:MF | 0.044 | *Hsc70-3* |
| RNA cap binding | GO:MF | 0.044 | *eIF4G1* |
| adenyl ribonucleotide binding | GO:MF | 0.044 | *Atpalpha,Lk6,Hsc70-3* |
| ATP binding | GO:MF | 0.044 | *Atpalpha,Lk6,Hsc70-3* |
| active transmembrane transporter activity | GO:MF | 0.044 | *Atpalpha,Hsc70-3* |
| ATP hydrolysis activity | GO:MF | 0.044 | *Atpalpha,Hsc70-3* |
| transporter activity | GO:MF | 0.044 | *Atpalpha,Hsc70-3,ogre* |
| transmembrane transporter activity | GO:MF | 0.044 | *Atpalpha,Hsc70-3,ogre* |
| translation initiation factor binding | GO:MF | 0.049 | *eIF4G1* |
| protein transmembrane transporter activity | GO:MF | 0.049 | *Hsc70-3* |

### 1.12. Gene Ontology Enrichment Analysis: somatic muscle

Significant GO terms (p < 0.05) with associated genes

| GO Term | Source | P-value | Genes |
| --- | --- | --- | --- |
| mitochondrial translation | GO:BP | 4.01e-46 | *Thor,mEFTu1,mRpL52,mRpL12,mRpL17,mRpS29,mRpS28,mRpS25,mRpL45,mRpL55,mRpL44,mRpS18C,mRpS18A,mRpL51,mRpS10,CG12848,mRpS33,mRpL54,tko,mRpL37,mRpL49,mRpL10,mRpS9,mRpS22,mRpL2,mRpL19,mRpL48,mRpS26,mRpL23,mRpS5,mRpL42,bonsai,mRpL36,mRpS2,mRpS17,mRpL1,mRpL22,mRpL27,mRpS6,mRpL14,mRpL38,mRpL40,GlyRS,mRpL21,mRpL16,mRpL4,mRpS11,mRpL32,mRpL15,mRpL20,mRpS35,mRpL18,mRpL47,mRpL46,mRpL3,mRpS23,mRpL24,mRpS34,mRpL39,mRpL9* |
| cytoplasmic translation | GO:BP | 2.42e-32 | *128up,CG8635,RpS15Ab,RpLP0-like,MCTS1,RpL34a,CNBP,RpS5a,Rack1,RpL30,RpS14a,RpL38,RpL40,RpLP0,RpL23A,RpS2,RpL35A,RpS10b,RpS20,RpL32,RpL29,RpL12,sta,RpLP1,RpS24,RpS3,RpL27,RpL34b,RpL13,RpS21,RpL13A,RpS7,RpL10Ab,RpL24,RpLP2,RpS12,RpL9,RpS27,RpL18A,RpL7,RpS13,RpS4,RpS27A,RpS8,RpL23,RpS9,RpL36A,RpS18,RpS26,RpL26,RpL28,RpS23,RpS16,RpS15,RpL27A* |
| mitochondrial electron transport, NADH to ubiquinone | GO:BP | 2.69e-21 | *ND-30,ND-B17.2,ND-39,ND-19,CG9034,ND-ASHI,ND-13B,ND-24,ND-B14.5B,ND-B14,ND-42,ND-B12,ND-B15,ND-SGDH,ND-PDSW,ND-B14.5A,ND-18,ND-13A,ND-15,ND-20,ND-B22,ND-23,ND-B18,ND-B14.7,ND-B17,ND-49,ND-75* |
| mitochondrial respirasome assembly | GO:BP | 2.98e-11 | *ND-B17.2,ND-39,ND-19,ND-ASHI,ND-B14.5B,ND-B14,ND-B12,ND-B15,ND-SGDH,ND-PDSW,ND-18,ND-15,ND-20,ND-B22,ND-23,ND-B18,CG10340,ND-B14.7,COX7A,Cox17,CG10075,COX6B,l(3)neo43,CG17996* |
| mitochondrial respiratory chain complex I assembly | GO:BP | 3.33e-11 | *ND-B17.2,ND-39,ND-19,ND-ASHI,ND-B14.5B,ND-B14,ND-B12,ND-B15,ND-SGDH,ND-PDSW,ND-18,ND-15,ND-20,ND-B22,ND-23,ND-B18,ND-B14.7* |
| ribose phosphate metabolic process | GO:BP | 3.86e-10 | *Gapdh1,Eno,Tpi,Pgk,Pglym78,Pfk,Gapdh2,tn,rut,CG11811,Pgi,awd,ATPsyndelta,Ald1,HINT1,ATPsynO,ATPsyngamma,ATPsynB,sun,ATPsynD,blw,Uck,sesB,ATPsynE,ATPsynG,ATPsynCF6,ATPsynF,ATPsynbeta,PyK,Vha68-1,ATPsynC* |
| nucleoside triphosphate biosynthetic process | GO:BP | 5.11e-10 | *awd,ATPsyndelta,ATPsynO,ATPsyngamma,ATPsynB,sun,ATPsynD,blw,Uck,sesB,ATPsynE,ATPsynG,ATPsynCF6,ATPsynF,ATPsynbeta,Vha68-1,ATPsynC* |
| striated muscle cell differentiation | GO:BP | 1.58e-09 | *sns,lmd,wupA,Vrp1,up,sals,Mlp84B,sing,Mlc2,bt,Mef2,Prm,Tm2,tn,sls,Mp20,Mical,MnM,blow,sd,if,how,flr* |
| muscle cell development | GO:BP | 2.15e-07 | *wupA,up,sals,Mlp84B,Mlc2,bt,Mef2,Prm,Tm2,tn,sls,Mical,MnM,sd,if,how,flr* |
| sarcomere organization | GO:BP | 2.62e-06 | *wupA,up,sals,Mlp84B,bt,Tm2,sls,Mical,MnM,if,how,flr* |
| glycolytic process | GO:BP | 3.90e-06 | *Gapdh1,Eno,Tpi,Pgk,Pglym78,Pfk,Gapdh2,tn,Pgi,Ald1,PyK* |
| nucleoside diphosphate catabolic process | GO:BP | 5.41e-06 | *Gapdh1,Eno,Tpi,Pgk,Pglym78,Pfk,Gapdh2,tn,Pgi,Ald1,PyK* |
| proton transmembrane transport | GO:BP | 7.81e-06 | *Cyt-c1,Vha36-3,ATPsynO,RFeSP,ND-20,ATPsyngamma,ATPsynB,sun,ATPsynD,blw,ATPsynE,levy,ATPsynG,ATPsynCF6,ATPsynF,ATPsynbeta,Vha68-1,ATPsynC* |
| purine ribonucleotide catabolic process | GO:BP | 1.12e-05 | *Gapdh1,Eno,Tpi,Pgk,Pglym78,Pfk,Gapdh2,tn,Pgi,Ald1,HINT1,PyK* |
| nucleotide catabolic process | GO:BP | 1.35e-05 | *Gapdh1,Eno,Tpi,Pgk,Pglym78,Pfk,Gapdh2,CG17224,tn,Pgi,Ald1,HINT1,PyK* |
| tricarboxylic acid cycle | GO:BP | 7.78e-05 | *CG7430,Fum1,SdhC,SdhD,CG5214,Idh3g,Idh3a,SdhA,SdhB,Idh3b,mAcon1* |
| mitochondrial electron transport, cytochrome c to oxygen | GO:BP | 1.67e-04 | *COX5B,COX8,COX5A,COX7A,COX7C,levy,CG7630,COX4* |
| protein refolding | GO:BP | 3.36e-04 | *Hsc70-1,CG14207,Hsp67Bc,Hsc70-5,Hsp70Ab,Hsc70-4,Hsp68,Hsp26,Hsp27* |
| organic acid metabolic process | GO:BP | 7.03e-04 | *Ldh,Gapdh1,regucalcin,Eno,Tpi,Pgk,CG9674,pyd3,CG9743,SERCA,Ahcy,Gs1,CG44242,Pglym78,CG7430,Pfk,Gapdh2,dj-1beta,tn,Fum1,Pgi,ND-ACP,CG33090,Gcat,Cth,CG15771,CG5214,Ald1,Acbp1,Got1,Idh3g,Idh3a,muc,Pdhb,SdhA,Idh3b,Mcad,Hibch,PyK* |
| mitochondrial electron transport, ubiquinol to cytochrome c | GO:BP | 7.38e-04 | *Cyt-c1,UQCR-C2,ox,RFeSP,UQCR-14,UQCR-Q* |
| glucose catabolic process | GO:BP | 0.003 | *Eno,Pfk,PhKgamma* |
| 'de novo' protein folding | GO:BP | 0.003 | *Hsc70-1,CG5001,Ero1L,unc-45,Hsc70-5,Hsp70Ab,Hsc70-4,Hsp68* |
| response to oxygen levels | GO:BP | 0.005 | *Drat,Mrtf,fau,tko,CHES-1-like,Phb1,Tom40,Hsp70Ab,UGP,MFS14,Vha68-1,Tm1* |
| proton motive force-driven mitochondrial ATP synthesis | GO:BP | 0.007 | *ATPsynO,sun,ATPsynF,ATPsynbeta* |
| protein insertion into mitochondrial outer membrane | GO:BP | 0.009 | *Tom7,CG7639,mge* |
| fructose 1,6-bisphosphate metabolic process | GO:BP | 0.009 | *Pfk,fbp,Ald1* |
| chaperone cofactor-dependent protein refolding | GO:BP | 0.01 | *Hsc70-1,CG5001,Ero1L,Hsc70-5,Hsp70Ab,Hsc70-4,Hsp68* |
| protein import into mitochondrial matrix | GO:BP | 0.01 | *Tom7,Tom20,Hsc70-5,CG6878,Tom40,mge,CG7394* |
| mitochondrial electron transport, succinate to ubiquinone | GO:BP | 0.011 | *SdhC,SdhD,SdhA,SdhB* |
| determination of adult lifespan | GO:BP | 0.013 | *Tpi,Thor,GlyP,rut,Tsp42Ef,SdhC,mld,ND-SGDH,Sam-S,ND-20,AGBE,sun,ATPsynD,sesB,Hsp68,levy,lt,Hsp26,Hsp27* |
| adult somatic muscle development | GO:BP | 0.014 | *wupA,sing,coro,Cf2,ewg* |
| gluconeogenesis | GO:BP | 0.017 | *Tpi,Pgk,fbp,Pgi* |
| muscle cell cellular homeostasis | GO:BP | 0.017 | *wupA,up,Pgk,tn,mbl,sesB,Rack1* |
| ribosomal large subunit assembly | GO:BP | 0.018 | *RpLP0-like,mRpL20,Non3,RpLP0,RpL23A* |
| positive regulation of protein ubiquitination | GO:BP | 0.031 | *Ndfip,SCCRO4,fzr,FipoQ,Rack1* |
| muscle attachment | GO:BP | 0.032 | *tx,ab,sls,if,Ilk,parvin,how* |
| lipid phosphorylation | GO:BP | 0.033 | *CG31140,rdgA,Mulk* |
| substrate adhesion-dependent cell spreading | GO:BP | 0.033 | *Tig,if,Ilk,parvin* |
| canonical glycolysis | GO:BP | 0.035 | *Eno,Pfk* |
| positive regulation of protein lipidation | GO:BP | 0.035 | *stv,Hsp67Bc* |
| positive regulation of translation | GO:BP | 0.038 | *CG13124,bol,mxt,Fmr1,hoip,La,CNBP,RpS9* |
| positive regulation of carbohydrate metabolic process | GO:BP | 0.042 | *GlyP,tn,GlyS,Rack1* |
| translational elongation | GO:BP | 0.043 | *mEFTu1,mRpL44,eEF1beta,Rack1,RpLP1,RpLP2* |
| regulation of monoatomic ion transmembrane transport | GO:BP | 0.046 | *Sh,SERCA,Ca-alpha1D,Ca-Ma2d,Rgk3,Rgk1,porin,levy,slo* |
| calcium import into the mitochondrion | GO:BP | 0.048 | *MCU,EMRE,porin* |
| isocitrate metabolic process | GO:BP | 0.048 | *Idh3g,Idh3a,Idh3b* |
| ribosomal small subunit assembly | GO:BP | 0.049 | *mRpS11,RpS5a,RpS14a,sta,RpS27,RpS15* |
| structural constituent of ribosome | GO:MF | 1.83e-93 | *mRpL52,mRpL12,mRpL17,mRpS29,mRpS28,mRpS25,mRpL45,mRpL55,mRpL44,mRpS18C,mRpS18A,mRpL51,mRpS10,mRpS33,RpS15Ab,mRpL54,tko,mRpL37,mRpL49,mRpL10,mRpS9,mRpS22,mRpL2,mRpL19,mRpL48,mRpS26,mRpL23,mRpS5,mRpL42,bonsai,mRpL36,mRpS2,mRpS17,mRpL1,mRpL22,mRpL27,mRpS6,RpLP0-like,mRpL14,mRpL38,mRpL40,mRpL21,mRpL16,mRpL4,mRpS11,mRpL32,mRpL15,mRpL20,mRpS35,CG4866,RpL34a,mRpL18,mRpL47,mRpL46,mRpL3,mRpS23,mRpL24,mRpS34,mRpL39,mRpL9,RpS5a,Rack1,RpL30,RpS14a,RpL38,RpL40,RpLP0,RpL23A,RpS2,RpL35A,RpS10b,RpS20,RpL32,RpL29,RpL12,sta,RpLP1,RpS24,RpS3,RpL27,RpL34b,RpL13,RpS21,RpL13A,RpS7,RpL10Ab,RpL24,RpLP2,RpS12,RpL9,RpS27,RpL18A,RpL7,RpS13,RpS4,RpS27A,RpS8,RpL23,RpS9,RpL36A,RpS18,RpS26,RpL26,RpL28,RpS23,RpS16,RpS15,RpL27A* |
| structural molecule activity | GO:MF | 3.69e-41 | *alphaTub85E,Strn-Mlck,Mlp84B,bt,Arpc3A,mRpL52,mRpL12,mRpL17,sls,betaTub60D,mRpS29,mRpS28,mRpS25,Cpr5C,mRpL45,Lam,mRpL55,mRpL44,mRpS18C,Arpc1,mRpS18A,mRpL51,mRpS10,mRpS33,RpS15Ab,betaTub97EF,mRpL54,tko,mRpL37,mRpL49,Nup153,mRpL10,mRpS9,mRpS22,mRpL2,mRpL19,mRpL48,mRpS26,mRpL23,mRpS5,mRpL42,bonsai,mRpL36,mRpS2,mRpS17,mRpL1,mRpL22,mRpL27,mRpS6,RpLP0-like,mRpL14,mRpL38,mRpL40,mRpL21,mRpL16,mRpL4,mRpS11,mRpL32,mRpL15,mRpL20,mRpS35,CG4866,RpL34a,Nup35,mRpL18,mRpL47,mRpL46,mRpL3,mRpS23,mRpL24,mRpS34,mRpL39,mRpL9,alphaTub84B,RpS5a,Rack1,RpL30,RpS14a,RpL38,RpL40,RpLP0,RpL23A,RpS2,RpL35A,RpS10b,RpS20,RpL32,RpL29,RpL12,sta,RpLP1,RpS24,RpS3,RpL27,RpL34b,RpL13,RpS21,RpL13A,RpS7,RpL10Ab,RpL24,RpLP2,RpS12,RpL9,RpS27,RpL18A,RpL7,RpS13,RpS4,RpS27A,RpS8,RpL23,RpS9,RpL36A,RpS18,RpS26,RpL26,RpL28,RpS23,RpS16,RpS15,RpL27A* |
| unfolded protein binding | GO:MF | 2.10e-14 | *Hsc70-1,CG5001,CCT1,CG10286,Pfdn4,Pfdn1,CCT3,CCT5,CCT4,Pfdn6,CCT6,CCT7,CCT2,Pfdn2,CG14207,Tim9a,Pfdn5,CCT8,Hsp67Bc,Hsc70-5,CG5504,Hsp70Ab,Hsc70-4,Hsp68,Nacalpha,Hsp83,Hsp26,Hsp27* |
| rRNA binding | GO:MF | 6.09e-14 | *mRpS18C,mRpS18A,CG12288,mRpS6,La,mRpL16,mRpS11,mRpL20,CG4866,mRpL18,Non3,RpS5a,RpS14a,RpLP0,RpL23A,RpL12,RpL9,RpS13,RpS4,RpL23,RpS9,RpS18* |
| RNA binding | GO:MF | 9.31e-13 | *EndoU,PheRS-m,CG4038,bru3,mRpL12,CG13124,Arc1,Nop60B,Fib,bol,Nop56,mxt,clu,yps,vig,NHP2,bin3,Fmr1,mod,mRpL44,mRpS18C,mbl,swm,mRpS18A,CG12288,Rbp9,Rb97D,eIF2A,LysRS,mRpS9,mRpL2,hoip,Rbp6,CG1542,mRpS5,CG13850,CG9775,CG13807,mRpL1,mRpS6,La,CG4045,CG1074,MCTS1,mRpL21,mRpL16,how,mRpS11,SLIRP2,mRpL20,CG8545,CG4866,mRpL18,CG11563,CNBP,mRpL24,Non3,CG5641,CG2021,AIMP1,l(1)G0004,Rpp25,RpS5a,RpL30,RpS14a,RpLP0,RpL23A,RpS2,RpS10b,RpS20,RpL12,RpS3,RpL13,RpL13A,RpL10Ab,RpL24,RpL9,RpS27,RpL7,RpS13,RpS4,RpL23,RpS9,RpS18,RpS26,RpL26,RpS16,RpS15* |
| proton-transporting ATP synthase activity, rotational mechanism | GO:MF | 3.21e-11 | *ATPsyndelta,ATPsynO,ATPsyngamma,ATPsynB,sun,ATPsynD,blw,ATPsynE,ATPsynG,ATPsynCF6,ATPsynF,ATPsynbeta,Vha68-1,ATPsynC* |
| proton channel activity | GO:MF | 2.83e-10 | *ATPsyndelta,ATPsynO,ATPsyngamma,ATPsynB,sun,ATPsynD,blw,ATPsynE,ATPsynG,ATPsynCF6,ATPsynF,ATPsynbeta,Vha68-1,ATPsynC* |
| protein folding chaperone | GO:MF | 1.51e-09 | *Hsc70-1,CCT1,unc-45,Pfdn1,CCT3,CCT5,CCT4,CCT6,CCT7,CCT2,Pfdn2,CCT8,Hsc70-5,Hsp70Ab,Hsc70-4,Hsp68,Hsp83* |
| proton transmembrane transporter activity | GO:MF | 1.53e-08 | *Cyt-c1,ND-30,ATPsyndelta,ND-24,Vha36-3,ATPsynO,RFeSP,ND-20,ND-23,ATPsyngamma,ATPsynB,sun,ND-49,ATPsynD,ND-75,blw,ATPsynE,ATPsynG,ATPsynCF6,ATPsynF,ATPsynbeta,Vha68-1,ATPsynC* |
| ATP-dependent protein folding chaperone | GO:MF | 1.98e-08 | *Hsc70-1,CCT1,CCT3,CCT5,CCT4,CCT6,CCT7,CCT2,CCT8,Hsc70-5,Hsp70Ab,Hsc70-4,Hsp68,Hsp83* |
| oxidoreductase activity | GO:MF | 1.63e-05 | *Asph,Ldh,Gapdh1,CG13284,CG9674,CG9743,Ero1L,Phyhd1,Cyp4aa1,CG7430,Gapdh2,dj-1beta,Rdh1,Drat,Mical,Cyt-c1,ND-30,ND-B17.2,ND-39,CG7322,ND-19,SdhC,Cyp4ac3,ND-ASHI,ND-13B,SdhD,CG5214,CG8888,ND-24,ND-B14.5B,ND-B14,ND-42,ND-B12,ND-B15,ND-SGDH,Idh3g,ND-PDSW,ND-B14.5A,Idh3a,RFeSP,ND-20,ND-B22,ND-23,ND-B18,ND-B17,ND-49,Pdhb,COX8,SdhA,CG9436,L2HGDH,ND-75,Sodh-2,COX5A,COX7A,SdhB,CG15717,Idh3b,CG31548,Mcad,COX6B,GstO2,CG31549,Nmdmc* |
| small ribosomal subunit rRNA binding | GO:MF | 2.42e-05 | *mRpS18C,mRpS18A,mRpS6,mRpS11,RpS14a,RpS13* |
| electron transfer activity | GO:MF | 1.50e-04 | *Cyt-c1,ND-30,SdhC,ND-24,RFeSP,ND-20,ND-23,ND-49,SdhA,ND-75,SdhB* |
| monoatomic cation channel activity | GO:MF | 1.52e-04 | *RyR,Sh,MCU,Ca-alpha1D,SK,Ca-Ma2d,CG4239,ATPsyndelta,EMRE,ATPsynO,ATPsyngamma,ATPsynB,sun,CG34396,ATPsynD,blw,ATPsynE,ATPsynG,ATPsynCF6,ATPsynF,ATPsynbeta,slo,Vha68-1,Ca-beta,ATPsynC* |
| inorganic cation transmembrane transporter activity | GO:MF | 1.63e-04 | *RyR,Sh,SERCA,MCU,Ca-alpha1D,SK,Ca-Ma2d,CG4239,Cyt-c1,ND-30,ATPsyndelta,ND-24,EMRE,Vha36-3,ATPsynO,RFeSP,ND-20,ND-23,ATPsyngamma,ATPsynB,sun,ND-49,CG34396,ATPsynD,ND-75,blw,ATPsynE,ATPsynG,ATPsynCF6,ATPsynF,ATPsynbeta,slo,Vha68-1,Ca-beta,ATPsynC* |
| monoatomic cation transmembrane transporter activity | GO:MF | 0.001 | *RyR,Sh,SERCA,MCU,Ca-alpha1D,SK,Ca-Ma2d,CG4239,Cyt-c1,ND-30,ATPsyndelta,ND-24,EMRE,Vha36-3,ATPsynO,RFeSP,ND-20,ND-23,ATPsyngamma,ATPsynB,sun,ND-49,CG34396,ATPsynD,ND-75,blw,ATPsynE,ATPsynG,ATPsynCF6,ATPsynF,ATPsynbeta,slo,Vha68-1,Ca-beta,ATPsynC* |
| actin binding | GO:MF | 0.002 | *wupA,Vrp1,CG1674,sals,Mhc,CG43897,Actn,Tm2,Arpc3A,coro,sls,Mp20,Mical,Arpc1,parvin,CG6891,tsr,flr,chic,Tm1* |
| cytoskeletal protein binding | GO:MF | 0.003 | *Mlc1,wupA,Vrp1,CG1674,up,sals,Mlp84B,Mhc,M7BP,Mlc2,CG43897,Actn,cana,Unc-76,Tm2,Arpc3A,coro,sls,unc-45,Mp20,Mical,mgr,awd,Arpc1,ens,ringer,Nin,parvin,CG6891,RpL34a,tsr,flr,chic,Tm1,RpL23,Hsp26* |
| ligase activity | GO:MF | 0.003 | *PheRS-m,LUBEL,Gs1,ATPsyndelta,Nadsyn,LysRS,ATPsynO,ATPsyngamma,ATPsynB,sun,ATPsynD,blw,ATPsynE,GlyRS,ATPsynG,ATPsynCF6,ATPsynF,ATPsynbeta,Vha68-1,ATPsynC* |
| structural constituent of cytoskeleton | GO:MF | 0.004 | *alphaTub85E,Strn-Mlck,Arpc3A,betaTub60D,Lam,Arpc1,betaTub97EF,alphaTub84B* |
| succinate dehydrogenase activity | GO:MF | 0.006 | *SdhC,SdhD,SdhA,SdhB* |
| NADH dehydrogenase (ubiquinone) activity | GO:MF | 0.007 | *ND-30,ND-24,ND-20,ND-23,ND-49,ND-75* |
| mRNA binding | GO:MF | 0.009 | *bru3,mRpL12,Arc1,bol,clu,yps,vig,Fmr1,mod,swm,CG12288,Rbp9,Rb97D,eIF2A,Rbp6,La,how,mRpS11,SLIRP2,CNBP,RpS5a,RpS14a,RpL13A,RpL24,RpS26* |
| iron-sulfur cluster binding | GO:MF | 0.009 | *CG9674,CG3420,MagR,ND-24,RFeSP,ND-20,ND-23,ND-75,SdhB,mAcon1,Nfs1* |
| NADH dehydrogenase activity | GO:MF | 0.016 | *ND-24,ND-20,ND-23,ND-49* |
| heat shock protein binding | GO:MF | 0.022 | *Hsc70-1,stv,Hsc70-5,CG5504,Hsp70Ab,Hsc70-4,Hsp68,CG13926* |
| quinone binding | GO:MF | 0.024 | *SdhD,ND-20,ND-49,SdhB* |
| isocitrate dehydrogenase (NAD+) activity | GO:MF | 0.025 | *Idh3g,Idh3a,Idh3b* |
| large ribosomal subunit rRNA binding | GO:MF | 0.025 | *RpLP0,RpL12,RpL23* |
| monoatomic ion channel activity | GO:MF | 0.027 | *RyR,Sh,MCU,Ca-alpha1D,SK,Ca-Ma2d,CG4239,ATPsyndelta,EMRE,ATPsynO,porin,ATPsyngamma,ATPsynB,sun,CG34396,ATPsynD,blw,ATPsynE,ATPsynG,ATPsynCF6,ATPsynF,ATPsynbeta,slo,Vha68-1,Ca-beta,ATPsynC* |
| channel activity | GO:MF | 0.027 | *RyR,Sh,MCU,Eglp4,Ca-alpha1D,SK,Ca-Ma2d,CG4239,ATPsyndelta,EMRE,ATPsynO,porin,ATPsyngamma,ATPsynB,sun,CG34396,ATPsynD,Tom40,blw,ATPsynE,ATPsynG,ATPsynCF6,ATPsynF,ATPsynbeta,slo,Vha68-1,Ca-beta,ATPsynC* |
| 2 iron, 2 sulfur cluster binding | GO:MF | 0.028 | *CG3420,MagR,ND-24,RFeSP,ND-75,SdhB* |
| actin filament binding | GO:MF | 0.028 | *Vrp1,sals,Mhc,Tm2,Arpc3A,coro,Mp20,Arpc1,CG6891,tsr,flr,Tm1* |
| misfolded protein binding | GO:MF | 0.037 | *Hsc70-1,Hsc70-5,Hsp70Ab,Hsc70-4,Hsp68* |
| succinate dehydrogenase (quinone) activity | GO:MF | 0.039 | *SdhD,SdhA,SdhB* |
| ATP-dependent diacylglycerol kinase activity | GO:MF | 0.039 | *CG31140,rdgA,Mulk* |
| monoatomic ion transmembrane transporter activity | GO:MF | 0.039 | *RyR,Sh,SERCA,MCU,Ca-alpha1D,SK,Ca-Ma2d,CG4239,Cyt-c1,ND-30,ATPsyndelta,ND-24,EMRE,Vha36-3,ATPsynO,RFeSP,porin,ND-20,ND-23,ATPsyngamma,ATPsynB,sun,ND-49,CG34396,ATPsynD,ND-75,blw,ATPsynE,ATPsynG,ATPsynCF6,ATPsynF,ATPsynbeta,slo,Vha68-1,Ca-beta,ATPsynC* |
| 2-hydroxyglutarate dehydrogenase activity | GO:MF | 0.041 | *Ldh,L2HGDH* |
| histone deacetylase binding | GO:MF | 0.041 | *SmydA-8,SmydA-1,Mef2,Smyd4-3,Smyd4-2* |
| ribosome binding | GO:MF | 0.042 | *clu,CG1354,eIF2A,CG3776,Nacalpha,Rack1,sta,RpS21* |
| nucleic acid binding | GO:MF | 0.043 | *lmd,EndoU,PheRS-m,gol,CG32813,sug,tx,ab,Snoo,Mef2,mEFTu1,jim,CG4038,CG31140,bru3,mRpL12,so,CG13124,Cf2,Arc1,Nop60B,Fib,bol,TFAM,neur,Nop56,tant,E(spl)m7-HLH,mxt,Nt5c,clu,yps,Kah,vig,NHP2,bin3,Fmr1,mod,mRpL44,mRpS18C,mbl,swm,mRpS18A,CG12288,mld,Rbp9,sd,Rb97D,mRpL49,eIF2A,LysRS,CHES-1-like,Nup153,mRpS9,mRpL2,hoip,Rbp6,CG1542,mRpS5,CG13850,CG9775,CG13807,mRpL1,net,mRpS6,mtSSB,La,CG4045,CG1074,Rs1,MCTS1,mRpL21,mRpL16,how,mRpS11,SLIRP2,mRpL20,CG8545,Parp,CG4866,Nup35,mRpL18,CG11563,CNBP,CG3726,mRpL24,Non3,CG5641,CG2021,AIMP1,bigmax,Polr1D,ewg,l(1)G0004,Rpp25,eEF1beta,RpS5a,RpL30,RpS14a,RpLP0,RpL23A,RpS2,RpS10b,RpS20,RpL12,RpS3,RpL13,RpL13A,RpL10Ab,RpL24,RpL9,RpS27,RpL7,RpS13,RpS4,RpL23,RpS9,RpS18,RpS26,RpL26,RpS16,RpS15* |

### 1.13. Gene Ontology Enrichment Analysis: cardioblasts

Significant GO terms (p < 0.05) with associated genes

| GO Term | Source | P-value | Genes |
| --- | --- | --- | --- |
| animal organ morphogenesis | GO:BP | 6.13e-37 | *disco-r,wb,sli,Syn2,CG43658,AdamTS-B,mthl5,how,rau,ed,tup,Rbfox1,bab2,mew,Mkp3,shg,GEFmeso,bnl,Timp,Smyd4-3,tin,flr,CG34347,jumu,Rip11,tinc,sqa,cindr,Galphao,rib,spg,Traf4,Grip,Src64B,CG43867,fz2,rst,Pura,pyd,Dys,Dg,baz,Nhe2,Ptpmeg,vn,sdk,flw,spi,pk,14-3-3zeta,S,unc-5,Dad,capt,raw,tsr,drl,fz,dome,Pak,kirre,Vang,crol,sn,Snr1,Egfr,Hrb87F,mam,Acer,Mtl,dock,stg,stck,kibra,Ten-m,tx,rin,RhoGAP19D,NetB,CNBP,gbb,CtBP,Mad,Mmp2,RasGAP1,mbl,ec,if,arm,Gug,ttk,sbb,exd,p38b,Ccm3,put,CadN,drk,Rok,Khc,mys,Stat92E,Usp47,Mnn1,Cka,mav,sd,Past1,ewg,EloC,sfl,Fak,ci,Ilk,Galphas,cdi,Cdep,tay* |
| RNA splicing, via transesterification reactions | GO:BP | 8.37e-19 | *how,bru2,Rbfox1,Pep,sqd,SmD3,Hel25E,Cpsf6,CG10418,hfp,SmB,SmF,SmE,tsu,Prp40,CG7483,yps,HnRNP-K,CG30122,x16,Hrb87F,SNRPG,Psi,RnpS1,Bin1,snf,CG14641,ytr,mbl,Phf5a,LSm7,Ref1,Rm62,CG11360,SmD2,lark,SF2,CG42724,U2af50,CG3198,qkr58E-1,Sf3b2,Acn,Caper,snRNP-U1-70K,mago,Bx42,SC35,CG9344,REG* |
| cell surface receptor signaling pathway | GO:BP | 3.22e-16 | *Tl,sli,wgn,AdamTS-B,mthl5,rau,ed,mew,Mkp3,htl,shg,GEFmeso,bnl,mib2,FER,Uba2,jumu,Galphao,spg,Ehbp1,Traf4,Src64B,Krn,fz2,Ack-like,Ptpmeg,Sumo,vn,HDAC1,flw,spi,14-3-3zeta,unc-5,Dad,lig,stumps,drl,fz,dome,nkd,CYLD,Vang,crol,Egfr,kek5,mam,dock,stck,gro,gbb,CtBP,Mad,Mmp2,RasGAP1,Fmr1,TER94,if,Nedd8,arm,Gug,ttk,sgl,p38b,Mob4,put,drk,Ptp4E,Ndfip,mys,Stat92E,Usp47,mav,SkpA,Mkrn1,ewg,EloC,sfl,C3G,ci,Cirl,Ilk,cdi,tay,HUWE1* |
| regulation of metabolic process | GO:BP | 6.17e-13 | *H15,mid,disco-r,Tl,disco,how,bru2,tup,CG16779,Rbfox1,bab2,htl,bnl,Timp,Smyd4-3,Mef2,tin,Tet,Hand,Uba2,ab,jumu,Pep,FipoQ,rib,Setd3,trsn,Traf4,Su(var)205,CG42672,Pits,Dg,tara,SmydA-1,sqd,Non2,CG18766,Sumo,Hel25E,HDAC1,TH1,Cpsf6,spi,Bap60,14-3-3zeta,Dad,CG10418,hfp,HP1b,lig,CG9705,drl,Bgb,l(3)neo38,Sps1,XNP,CG7483,yps,crol,Snr1,Lk6,tou,HnRNP-K,Egfr,x16,Hrb87F,CG8209,mam,gro,rump,kibra,Psi,Pop2,Psc,tx,rin,CG8963,Dek,Bin1,snf,CNBP,CG17124,CG14641,CG12391,dod,CtBP,Mad,Rpn2,Fmr1,TER94,mbl,ssx,Nedd8,arm,Gug,ttk,Brd7-9,LSm7,sbb,Ndf,exd,crp,Eip78C,FoxK,Rm62,CG11360,Rpn1,CG32772,p38b,lark,SF2,NK7.1,CG32767,Ccm3,Nipped-A,Rpt5,put,p23,vih,U2af50,CG3198,qkr58E-1,Helz,Ndfip,Rok,hzg,mri,MED11,MED22,Stat92E,Usp47,ko,Mnn1,Pfdn5,Acn,Cka,CoRest,SkpA,Caper,sd,snRNP-U1-70K,Nurf-38,Rpt1,mago,Ythdf,Mkrn1,SC35,gpp,Usp2,Ntf-2,ewg,Chro,DCP2,Atg13,Fak,ci,Ufd4,CG13124,REG* |
| regulation of alternative mRNA splicing, via spliceosome | GO:BP | 4.56e-12 | *how,bru2,Rbfox1,sqd,Hel25E,Cpsf6,CG10418,hfp,HnRNP-K,x16,Hrb87F,Psi,snf,CG14641,mbl,Rm62,CG11360,SF2,U2af50,qkr58E-1,Caper,snRNP-U1-70K,SC35* |
| striated muscle cell differentiation | GO:BP | 4.60e-10 | *kon,DAAM,how,mib2,Mef2,tin,flr,rst,Dg,tmod,sals,wupA,kirre,dock,if,Rok,mys,bves,Pak3,sd,C3G* |
| regulation of RNA metabolic process | GO:BP | 2.71e-09 | *H15,mid,disco-r,Tl,disco,how,bru2,tup,CG16779,Rbfox1,bab2,Mef2,tin,Tet,Hand,Uba2,ab,jumu,Pep,rib,Setd3,Su(var)205,Pits,sqd,Non2,CG18766,Sumo,Hel25E,HDAC1,TH1,Cpsf6,Bap60,14-3-3zeta,Dad,CG10418,hfp,HP1b,CG9705,Bgb,l(3)neo38,crol,Snr1,tou,HnRNP-K,x16,Hrb87F,mam,gro,rump,kibra,Psi,Pop2,Psc,tx,Bin1,snf,CG14641,CG12391,CtBP,Mad,Fmr1,mbl,arm,Gug,ttk,Brd7-9,sbb,Ndf,exd,crp,Eip78C,FoxK,Rm62,CG11360,CG32772,SF2,NK7.1,CG32767,Nipped-A,U2af50,qkr58E-1,hzg,MED11,MED22,Stat92E,ko,Mnn1,Pfdn5,CoRest,Caper,sd,snRNP-U1-70K,Nurf-38,Ythdf,SC35,ewg,Chro,DCP2,ci* |
| negative regulation of signal transduction | GO:BP | 3.09e-09 | *Tl,AdamTS-B,ed,Strn-Mlck,Mkp3,Uba2,Src64B,Patronin,chrb,Ptpmeg,HDAC1,flw,Dad,raw,lig,nkd,CYLD,crol,Egfr,kek5,dock,stck,gro,His2Av,CtBP,Mmp2,RasGAP1,Fmr1,TER94,Nedd8,Gug,p38b,Mob4,Ptp4E,Ndfip,JMJD6,Mnn1,Cka,SkpA,Usp2,EloC,ci,Ufd4,cdi,tay,HUWE1* |
| dorsal closure | GO:BP | 5.30e-08 | *ed,tup,FER,jumu,rib,Traf4,Ack-like,pyd,spi,raw,Pak,Egfr,Mtl,stck,gbb,arm,put,Rok,mys,Cka* |
| muscle cell development | GO:BP | 1.84e-07 | *kon,DAAM,how,htl,Mef2,flr,Dys,Dg,tmod,sals,wupA,if,mys,sd,C3G* |
| imaginal disc-derived wing vein specification | GO:BP | 6.71e-07 | *AdamTS-B,Rbfox1,Mkp3,GEFmeso,Dys,Dg,S,Snr1,Egfr,tx,gbb,Past1,EloC,sfl* |
| gastrulation | GO:BP | 8.54e-07 | *sli,htl,shg,Traf4,Dtg,baz,spi,stumps,mam,stg,arm,sgl,Rok,mys,Prosalpha6,T48,sfl* |
| cellular response to growth factor stimulus | GO:BP | 1.04e-06 | *rau,htl,bnl,Src64B,Sumo,Dad,stumps,kek5,gro,gbb,Mad,Mmp2,RasGAP1,sgl,p38b,put,drk,Ptp4E,mav,sfl* |
| nucleobase-containing compound metabolic process | GO:BP | 1.38e-06 | *H15,mid,disco-r,Tl,disco,how,bru2,tup,CG16779,Rbfox1,bab2,Mef2,tin,Tet,Hand,Uba2,ab,jumu,Pep,rib,Hmgs,Setd3,trsn,Su(var)205,Pits,Dg,tara,sqd,SmD3,Non2,CG18766,CG12065,Sumo,Hel25E,HDAC1,TH1,Cpsf6,Bap60,14-3-3zeta,Dad,CG10418,hfp,HP1b,SmB,SmF,CG9705,Bgb,l(3)neo38,SmE,XNP,tsu,Prp40,koi,CG7483,yps,crol,Snr1,tou,HnRNP-K,CG30122,x16,Hrb87F,SNRPG,mam,gro,rump,kibra,Psi,Pop2,Psc,TfIIS,RnpS1,tx,His2Av,CG3732,Dek,Bin1,snf,CG14641,CG12391,CG4022,CtBP,Mad,CG13096,ytr,Fmr1,e(r),mbl,ssx,arm,Gug,ttk,Brd7-9,sgl,Phf5a,LSm7,Taf13,sbb,Ndf,exd,Ref1,crp,Eip78C,FoxK,Rm62,CG11360,CG32772,SmD2,lark,SF2,NK7.1,CG32767,Nipped-A,CG42724,Polr2E,U2af50,CG3198,qkr58E-1,hzg,Sem1,hrg,Polr2K,Taf10b,Sf3b2,MED11,MED22,Stat92E,ko,Mnn1,Pfdn5,Acn,CoRest,SkpA,Caper,sd,pyd3,snRNP-U1-70K,Nurf-38,mago,Ythdf,Bx42,SC35,gpp,Hmgcr,ewg,Chro,EloC,thoc7,CG9344,DCP2,ci,REG,CG13807,HUWE1* |
| ommatidial rotation | GO:BP | 2.58e-06 | *shg,spi,pk,S,fz,Vang,Egfr,rin,ec,CadN* |
| sarcomere organization | GO:BP | 8.52e-06 | *kon,DAAM,how,flr,Dg,sals,wupA,if,mys,C3G* |
| tracheal outgrowth, open tracheal system | GO:BP | 2.37e-05 | *Timp,rib,stumps,Egfr,Mmp2,ttk,drk* |
| maintenance of epithelial integrity, open tracheal system | GO:BP | 5.61e-05 | *mew,Mkp3,shg,Egfr,if,mys* |
| head involution | GO:BP | 1.12e-04 | *tup,shg,rib,Ack-like,pyd,chrb,Mtl,stck,put* |
| negative regulation of JNK cascade | GO:BP | 1.58e-04 | *flw,raw,stck,JMJD6,Mnn1,Cka,SkpA,HUWE1* |
| negative regulation of biosynthetic process | GO:BP | 1.59e-04 | *mid,bru2,CG16779,Rbfox1,Smyd4-3,tin,ab,trsn,Su(var)205,Pits,SmydA-1,sqd,HDAC1,TH1,spi,Bap60,HP1b,CG9705,XNP,crol,Egfr,gro,rump,Psi,Pop2,Psc,Bin1,CtBP,Mad,Fmr1,ssx,arm,Gug,ttk,LSm7,sbb,Eip78C,FoxK,Rm62,CG32767,p23,Helz,Stat92E,CoRest,SkpA,sd,snRNP-U1-70K,Ythdf,gpp,Usp2,Chro,DCP2,ci* |
| negative regulation of epidermal growth factor receptor signaling pathway | GO:BP | 2.51e-04 | *AdamTS-B,ed,Mkp3,Ptpmeg,RasGAP1,Gug,Ptp4E,tay* |
| positive regulation of MAP kinase activity | GO:BP | 5.02e-04 | *Sumo,Ccm3,Cka* |
| apposition of dorsal and ventral imaginal disc-derived wing surfaces | GO:BP | 5.48e-04 | *how,mew,mam,stck,if,Ilk* |
| positive regulation of hippo signaling | GO:BP | 5.55e-04 | *ed,RtGEF,Src64B,pyd,14-3-3zeta,lig,kibra,alpha-Spec* |
| salivary gland morphogenesis | GO:BP | 6.57e-04 | *mew,shg,tin,rib,Src64B,fz2,raw,drl,fz,Mtl,if* |
| homophilic cell adhesion via plasma membrane adhesion molecules | GO:BP | 7.66e-04 | *Fas3,ed,Fas1,shg,rst,sdk,fz,kirre,CadN* |
| ventral cord development | GO:BP | 9.26e-04 | *htl,Timp,tin,Galphao,18w,Psc,gbb,Mad,Mmp2,if,mys* |
| branch fusion, open tracheal system | GO:BP | 9.41e-04 | *ed,shg,bnl,pyd,arm,ttk* |
| regulation of actin filament depolymerization | GO:BP | 0.001 | *flr,tmod,alpha-Spec,beta-Spec,Svil* |
| positive regulation of ERK1 and ERK2 cascade | GO:BP | 0.002 | *spg,vn,spi,Egfr,drk,Usp47* |
| epidermal cell differentiation | GO:BP | 0.002 | *flr,Galphao,pk,tsr,fz,Vang,sn,Mtl,Rok* |
| negative regulation of synaptic assembly at neuromuscular junction | GO:BP | 0.002 | *Galphao,Src64B,Dad,Fmr1,Mob4,SkpA,ewg,Fak* |
| muscle cell cellular homeostasis | GO:BP | 0.002 | *mib2,Dys,Dg,capt,wupA,TER94,mbl* |
| regulation of chromosome organization | GO:BP | 0.002 | *Bub3,cana,Non2,Bap60,CG15237,vih,Stat92E,SkpA,Chro,BuGZ* |
| negative regulation of hippo signaling | GO:BP | 0.002 | *Strn-Mlck,Src64B,Patronin,p38b,Mob4,Cka,SkpA,ci* |
| segment polarity determination | GO:BP | 0.002 | *fz2,fz,nkd,Egfr,arm,sgl,sfl,ci* |
| glial cell migration | GO:BP | 0.003 | *sli,how,htl,bnl,unc-5,XNP,NetB* |
| imaginal disc-derived wing vein morphogenesis | GO:BP | 0.003 | *Dys,vn,Snr1,Egfr,gbb,Mad,Gug,tay* |
| germarium-derived oocyte fate determination | GO:BP | 0.003 | *baz,14-3-3zeta,BicD,tsu,Fmr1,alpha-Spec* |
| female germline ring canal stabilization | GO:BP | 0.004 | *Src64B,Msp300,Rok* |
| notum cell fate specification | GO:BP | 0.004 | *tup,vn,Egfr* |
| mesoderm migration involved in gastrulation | GO:BP | 0.004 | *sli,htl,sgl,sfl* |
| maintenance of presynaptic active zone structure | GO:BP | 0.004 | *Dys,Ten-m,alpha-Spec,beta-Spec* |
| determination of digestive tract left/right asymmetry | GO:BP | 0.004 | *shg,tin,dome,Stat92E* |
| spliceosomal snRNP assembly | GO:BP | 0.004 | *SmD3,SmF,SmE,SNRPG,SmD2* |
| oocyte microtubule cytoskeleton polarization | GO:BP | 0.004 | *14-3-3zeta,capt,BicD,Khc,mago* |
| heterophilic cell-cell adhesion via plasma membrane cell adhesion molecules | GO:BP | 0.005 | *Fas3,mew,kirre,if,glec,mys* |
| somatic stem cell population maintenance | GO:BP | 0.005 | *shg,mam,His2Av,Mad,arm* |
| female gonad development | GO:BP | 0.005 | *bab2,fz2,tsr,Hrb87F* |
| negative regulation of microtubule depolymerization | GO:BP | 0.005 | *Patronin,tacc,alpha-Spec,beta-Spec* |
| chondroitin sulfate biosynthetic process | GO:BP | 0.006 | *Chpf,Chsy,sgl* |
| stem cell fate commitment | GO:BP | 0.006 | *spi,S,Egfr* |
| regulation of transcriptional start site selection at RNA polymerase II promoter | GO:BP | 0.006 | *x16,SF2,SC35* |
| photoreceptor cell fate determination | GO:BP | 0.006 | *spi,S,Egfr* |
| ventral furrow formation | GO:BP | 0.006 | *shg,Traf4,arm,Prosalpha6,T48* |
| hemocyte migration | GO:BP | 0.008 | *shg,sn,Mtl,if,mys* |
| stomatogastric nervous system development | GO:BP | 0.008 | *spi,S* |
| positive regulation of torso signaling pathway | GO:BP | 0.008 | *Src64B,14-3-3zeta* |
| long-term strengthening of neuromuscular junction | GO:BP | 0.008 | *alpha-Spec,beta-Spec* |
| negative regulation of canonical Wnt signaling pathway | GO:BP | 0.008 | *nkd,gro,CtBP,Mmp2,TER94,SkpA,HUWE1* |
| R7 cell development | GO:BP | 0.009 | *rau,Ten-m,ttk,CadN* |
| positive regulation of epidermal growth factor receptor signaling pathway | GO:BP | 0.009 | *rau,GEFmeso,Src64B,Usp47* |
| dorsal/ventral axis specification, ovarian follicular epithelium | GO:BP | 0.009 | *sqd,lig,Egfr,rin* |
| regulation of mRNA 3'-end processing | GO:BP | 0.009 | *x16,SF2,SC35* |
| dendrite guidance | GO:BP | 0.011 | *tup,HDAC1,drl,Snr1,NetB* |
| negative regulation of sister chromatid segregation | GO:BP | 0.011 | *Bub3,CG15237,Chro,BuGZ* |
| positive regulation of transcription by RNA polymerase II | GO:BP | 0.012 | *H15,mid,Tl,disco,tup,Mef2,tin,Tet,Hand,Setd3,Su(var)205,HP1b,Snr1,tou,mam,tx,CtBP,Mad,arm,ttk,Ndf,exd,CG32767,Stat92E,ko,sd,Nurf-38,ci* |
| response to cytokine | GO:BP | 0.013 | *Tl,wgn,Traf4,dome,CYLD,HUWE1* |
| primary branching, open tracheal system | GO:BP | 0.014 | *Mkp3,bnl,put* |
| oenocyte development | GO:BP | 0.014 | *how,spi,exd* |
| ectodermal digestive tract morphogenesis | GO:BP | 0.014 | *mew,tin,raw* |
| gonadal mesoderm development | GO:BP | 0.014 | *htl,shg,tin* |
| imaginal disc-derived wing expansion | GO:BP | 0.014 | *disco-r,Timp,arm* |
| positive regulation of sevenless signaling pathway | GO:BP | 0.014 | *spg,Src64B,Usp47* |
| axon midline choice point recognition | GO:BP | 0.015 | *sli,drl,alpha-Spec,beta-Spec,RhoGAP93B* |
| motor neuron axon guidance | GO:BP | 0.017 | *tup,fz2,unc-5,fend,Ten-m,NetB,Mmp2,Ptp4E,ko* |
| regulation of mitotic sister chromatid separation | GO:BP | 0.017 | *Bub3,CG15237,vih,Chro,BuGZ* |
| positive regulation of border follicle cell migration | GO:BP | 0.018 | *shg,Krn,vn,spi,Egfr,ttk* |
| determination of genital disc primordium | GO:BP | 0.019 | *spi,S,Egfr* |
| zonula adherens assembly | GO:BP | 0.019 | *shg,baz,arm* |
| larval visceral muscle development | GO:BP | 0.019 | *htl,Msp300,kirre* |
| mesodermal cell fate determination | GO:BP | 0.019 | *htl,mam* |
| neuroblast fate specification | GO:BP | 0.019 | *mid,nkd* |
| negative regulation of muscle organ development | GO:BP | 0.019 | *Him,Dad* |
| negative regulation of dendrite morphogenesis | GO:BP | 0.019 | *Fmr1,CadN* |
| regulation of Malpighian tubule diameter | GO:BP | 0.019 | *rib,raw* |
| negative regulation of mRNA splicing, via spliceosome | GO:BP | 0.019 | *Psi,snRNP-U1-70K* |
| negative regulation of insulin receptor signaling pathway | GO:BP | 0.019 | *Tl,Mkp3,dock,Fmr1,SkpA* |
| myoblast fate specification | GO:BP | 0.019 | *htl,ttk* |
| negative regulation of mitotic nuclear division | GO:BP | 0.02 | *Bub3,CG15237,Chro,BuGZ* |
| germ-band shortening | GO:BP | 0.024 | *tup,raw,Egfr,Mtl* |
| negative regulation of cell adhesion | GO:BP | 0.024 | *disco-r,Timp,baz,arm* |
| cell projection assembly | GO:BP | 0.025 | *sprt,kon,DAAM,shg,bnl,baz,pico,vn,Unc-115a,tsr,sn,Egfr,Mtl,Fmr1,alphaTub84D,mys,Past1,Flo2* |
| negative regulation of transcription by RNA polymerase II | GO:BP | 0.027 | *mid,ab,Su(var)205,HDAC1,TH1,HP1b,gro,Psc,CtBP,Mad,arm,Gug,ttk,sbb,CG32767,CoRest,Chro,ci* |
| R8 cell fate specification | GO:BP | 0.029 | *kibra,gbb,Mad,put* |
| regulation of myoblast fusion | GO:BP | 0.031 | *rst,kirre,Rok* |
| epithelial cell proliferation involved in Malpighian tubule morphogenesis | GO:BP | 0.031 | *spi,S,Egfr* |
| cell elongation involved in imaginal disc-derived wing morphogenesis | GO:BP | 0.031 | *shg,Dad,arm* |
| negative regulation of actin filament polymerization | GO:BP | 0.033 | *tmod,alpha-Spec,beta-Spec,Svil* |
| fasciculation of motor neuron axon | GO:BP | 0.033 | *mid,Mmp2* |
| nephrocyte diaphragm assembly | GO:BP | 0.033 | *pyd,kirre* |
| protein localization to adherens junction | GO:BP | 0.033 | *arm,Shrm* |
| synaptic target attraction | GO:BP | 0.033 | *Ten-m,NetB* |
| outflow tract morphogenesis | GO:BP | 0.033 | *sli,shg* |
| digestive tract mesoderm development | GO:BP | 0.033 | *wb,sli* |
| mushroom body development | GO:BP | 0.037 | *DAAM,ab,Src64B,Ptpmeg,tsr,drl,Fmr1,robl* |
| female sex differentiation | GO:BP | 0.038 | *bab2,fz2,tsr,Hrb87F* |
| mRNA splice site recognition | GO:BP | 0.038 | *bru2,hfp,CG3198* |
| histoblast morphogenesis | GO:BP | 0.038 | *Dad,stg,Mad* |
| R8 cell development | GO:BP | 0.038 | *pyd,kibra,CadN* |
| chorion-containing eggshell pattern formation | GO:BP | 0.038 | *H15,mid,Egfr* |
| retinal ganglion cell axon guidance | GO:BP | 0.038 | *fz2,Dad,CadN* |
| germ cell migration | GO:BP | 0.039 | *htl,shg,tin,Fpps,Hmgcr* |
| negative regulation of cell population proliferation | GO:BP | 0.039 | *Bap60,hfp,Snr1,kibra,Mmp2,Fmr1,Mnn1,sd* |
| cell fate commitment involved in pattern specification | GO:BP | 0.043 | *tup,vn,fz,Egfr,CtBP* |
| positive regulation of translation | GO:BP | 0.045 | *CG8963,CNBP,Fmr1,lark,Mkrn1,CG13124* |
| haltere development | GO:BP | 0.047 | *vn,drl,Egfr* |
| positive regulation of axon guidance | GO:BP | 0.047 | *pk,fz,Vang* |
| cell-matrix adhesion | GO:BP | 0.049 | *mew,if,mys,Ilk* |
| cell adhesion mediated by integrin | GO:BP | 0.049 | *mew,tx,if,mys* |
| larval heart development | GO:BP | 0.05 | *Hand,mys* |
| pericentric heterochromatin formation | GO:BP | 0.05 | *Su(var)205,XNP* |
| axis elongation | GO:BP | 0.05 | *baz,18w* |
| negative regulation of sevenless signaling pathway | GO:BP | 0.05 | *RasGAP1,cdi* |
| chaperone-mediated protein complex assembly | GO:BP | 0.05 | *Pfdn6,p23* |
| positive regulation of cytoplasmic translation | GO:BP | 0.05 | *CNBP,lark* |
| embryonic digestive tract morphogenesis | GO:BP | 0.05 | *tin,gbb* |
| muscle cell fate determination | GO:BP | 0.05 | *how,tup* |
| protein binding | GO:MF | 1.57e-40 | *mid,Tl,CG45263,disco,CAP,sli,sprt,Syn2,wgn,ndl,kon,DAAM,rau,Fas3,ed,Nlg1,tup,LRP1,Rbfox1,sick,Fas1,bab2,mew,Strn-Mlck,htl,Spn,shg,Magi,GEFmeso,bnl,Timp,mib2,zip,Smyd4-3,Mef2,RtGEF,FER,a,flr,Hand,Uba2,side-V,ab,Rip11,CG9135,sqa,cindr,Galphao,FipoQ,rib,spg,Setd3,trsn,Ehbp1,Oatp74D,Traf4,Grip,Src64B,Krn,Su(var)205,CG7029,fz2,CG42672,rst,Bub3,cana,CG34417,gukh,CG43102,Ack-like,CG5886,pyd,Dys,Dg,Patronin,SmydA-1,baz,Evi5,Plp,Df31,CG18766,pns,Ptpmeg,pico,Sumo,vn,sdk,Unc-115a,tmod,HDAC1,flw,spi,pk,Bap60,Eb1,14-3-3zeta,unc-5,CG3408,Dad,CG10418,capt,hfp,HP1b,tsr,SmF,lig,Msp300,fbp,stumps,emb,drl,Bgb,sals,fz,SmE,BicD,tsu,Prp40,HP4,wupA,dome,nkd,tacc,Pak,kirre,CG7483,yps,Vang,sn,Snr1,Lk6,tou,HnRNP-K,His3.3B,CG30122,Egfr,kek5,CG14207,CG42673,CG8209,Meltrin,Septin2,Mtl,dock,18w,stck,gro,kibra,Psi,Pop2,ReepA,RyR,Ten-m,stx,tx,rin,His2Av,CG8963,RhoGAP19D,Dek,NetB,Bin1,snf,dod,Rpn9,gbb,cass,CtBP,Mad,CCT8,toc,Fmr1,Nph,TER94,ec,Lrch,lbk,if,Nedd8,CG11658,arm,Gug,ttk,Brd7-9,CCT4,LSm7,Taf13,sbb,CG6966,Mapmodulin,exd,crp,FoxK,Rm62,alpha-Spec,Shrm,Rpn8,Set,CCT3,SF2,CG32767,CG6891,Pfdn6,Ccm3,Nipped-A,Rpt5,put,Rpn7,p23,His3.3A,CadN,CG42724,beta-Spec,drk,U2af50,Ptp4E,Hou,Rpn6,Helz,Ndfip,Rok,CCT7,Sem1,mri,CG16974,alphaTub84D,Taf10b,MED11,MED22,Khc,mys,Stat92E,Sobp,CCT1,Pfdn5,Acn,Cka,CoRest,Con,mav,SkpA,Pak3,CCT6,Mhcl,Ras64B,sd,snRNP-U1-70K,Svil,CCT2,CCT5,mago,Pfdn1,Ythdf,Bx42,CG10011,Actn,Past1,Hmgcr,Prosbeta3,Roc2,Prosbeta5,Vinc,Pgam5,Ntf-2,Chro,EloC,smash,RhoGAP93B,CG9344,DCP2,Atg13,BuGZ,C3G,Arl1,ci,SK,eIF3i,Ufd4,CG13124,Nlp,Ilk,mgr,robl,Galphas,zormin,Slip1,Cdep,Fkbp12,HUWE1* |
| RNA binding | GO:MF | 2.47e-10 | *how,bru2,Rbfox1,Pep,trsn,Su(var)205,sqd,SmD3,Hel25E,TH1,Cpsf6,CG10418,hfp,SmB,SmF,CG9705,SmE,tsu,Prp40,CG7483,yps,HnRNP-K,x16,Hrb87F,SNRPG,rump,Psi,RnpS1,rin,CG3732,CG8963,Rsf1,snf,CNBP,CG14641,cass,CG13096,ytr,Fmr1,Nph,mbl,ssx,Capr,Phf5a,LSm7,Ref1,Rm62,CG11360,B52,SmD2,lark,SF2,U2af50,CG3198,qkr58E-1,Helz,hrg,Sf3b2,Caper,snRNP-U1-70K,mago,Ythdf,Mkrn1,SC35,CG31712,CG9344,DCP2,eIF3i,CG13124,Nlp,CG13807* |
| cytoskeletal protein binding | GO:MF | 1.27e-09 | *sprt,DAAM,Spn,shg,mib2,zip,flr,Setd3,cana,gukh,Dys,Patronin,Ptpmeg,Unc-115a,tmod,Bap60,Eb1,capt,tsr,Msp300,sals,BicD,wupA,tacc,sn,ReepA,Ten-m,toc,crp,alpha-Spec,Shrm,CG6891,beta-Spec,alphaTub84D,Khc,Mhcl,Svil,Actn,Vinc,smash,DCP2,BuGZ,mgr,zormin,Cdep* |
| mRNA binding | GO:MF | 1.37e-09 | *how,bru2,Rbfox1,Su(var)205,sqd,Cpsf6,CG9705,tsu,CG7483,yps,HnRNP-K,x16,Hrb87F,rump,Psi,RnpS1,rin,snf,CNBP,CG14641,ytr,Fmr1,ssx,Ref1,Rm62,lark,SF2,U2af50,CG3198,qkr58E-1,Caper,snRNP-U1-70K,Ythdf,Mkrn1,SC35* |
| nucleic acid binding | GO:MF | 4.18e-08 | *H15,mid,disco,how,bru2,tup,CG32813,Rbfox1,bab2,Mef2,tin,Tet,Hand,ab,jumu,Pep,rib,trsn,Su(var)205,sqd,SmD3,Hel25E,TH1,Cpsf6,Bap60,Dad,CG10418,hfp,SmB,SmF,CG9705,PDCD-5,Bgb,l(3)neo38,SmE,XNP,tsu,Prp40,CG31140,CG7483,yps,crol,Snr1,tou,HnRNP-K,His3.3B,x16,Hrb87F,SNRPG,rump,Psi,Pop2,baf,Psc,TfIIS,RnpS1,tx,rin,His2Av,CG3732,CG8963,Rsf1,Dek,snf,CNBP,CG14641,CG12391,cass,Mad,CG13096,ytr,Fmr1,Nph,mbl,ssx,ttk,Capr,Phf5a,CG31301,LSm7,Ndf,exd,Ref1,crp,Eip78C,FoxK,Rm62,CG11360,CG32772,B52,SmD2,lark,D1,SF2,NK7.1,CG32767,His3.3A,Polr2E,U2af50,CG3198,qkr58E-1,Helz,hrg,Polr2K,Sf3b2,Stat92E,ko,Mnn1,Acn,Caper,sd,snRNP-U1-70K,mago,Ythdf,Mkrn1,Bx42,SC35,ewg,CG31712,CG9344,DCP2,Nt5c,BuGZ,ci,eIF3i,CG13124,Nlp,CG13807* |
| actin binding | GO:MF | 6.68e-08 | *sprt,DAAM,Spn,zip,flr,Setd3,gukh,Dys,Unc-115a,capt,tsr,Msp300,sals,wupA,sn,alpha-Spec,Shrm,CG6891,beta-Spec,Mhcl,Svil,Actn,Vinc,DCP2,zormin* |
| cell adhesion molecule binding | GO:MF | 1.14e-06 | *CG45263,Fas3,ed,Fas1,mew,cindr,rst,pyd,Dg,sdk,kirre,dock,if,arm,CadN,mys* |
| enzyme binding | GO:MF | 1.57e-06 | *DAAM,rau,Magi,GEFmeso,Timp,Smyd4-3,Mef2,Uba2,Rip11,CG9135,spg,Traf4,Su(var)205,CG7029,SmydA-1,Evi5,pns,pico,Sumo,capt,Msp300,stumps,emb,drl,BicD,dome,Pak,Lk6,CG42673,Mtl,dock,kibra,Fmr1,ec,Nedd8,arm,Shrm,Ccm3,CG42724,Rok,Cka,Pak3,Pgam5,ci* |
| transcription factor binding | GO:MF | 1.76e-06 | *mid,tup,Rbfox1,HDAC1,Bap60,14-3-3zeta,Bgb,dome,Snr1,tou,HnRNP-K,gro,tx,Bin1,dod,CtBP,Mad,arm,Gug,exd,Sobp,sd,ci* |
| actin filament binding | GO:MF | 2.69e-06 | *Spn,zip,flr,Dys,Unc-115a,tsr,Msp300,sals,sn,alpha-Spec,Shrm,CG6891,beta-Spec,Mhcl,Svil,Vinc,DCP2* |
| protein domain specific binding | GO:MF | 2.95e-06 | *Tl,cindr,fz2,CG42672,rst,Dys,pico,Bap60,nkd,Pak,Snr1,gro,ttk,Ndfip,sd,C3G,Arl1* |
| molecular adaptor activity | GO:MF | 5.03e-05 | *CG16779,RtGEF,Setd3,Su(var)205,gukh,Pits,Plp,HDAC1,Bap60,stumps,Bgb,BicD,koi,sn,Snr1,mam,Septin2,dock,gro,kibra,Bin1,CtBP,Mad,Fmr1,arm,Gug,sbb,exd,CG42724,drk,MED11,MED22,CoRest,SkpA,Ythdf,EloC* |
| DNA-binding transcription factor binding | GO:MF | 1.45e-04 | *tup,HDAC1,Bap60,dome,Snr1,tou,gro,tx,Bin1,CtBP,Mad,arm,Gug,exd,ci* |
| epidermal growth factor receptor binding | GO:MF | 4.10e-04 | *ed,Krn,vn,spi,drk* |
| mRNA 3'-UTR binding | GO:MF | 5.07e-04 | *how,bru2,Rbfox1,sqd,CG9705,Hrb87F,rump,Fmr1,ssx,Mkrn1* |
| protein-macromolecule adaptor activity | GO:MF | 6.26e-04 | *CG16779,Setd3,Su(var)205,gukh,Pits,HDAC1,Bap60,stumps,Bgb,BicD,koi,sn,Snr1,mam,dock,gro,Bin1,CtBP,Mad,Fmr1,arm,Gug,sbb,exd,CG42724,drk,MED11,MED22,CoRest,Ythdf,EloC* |
| protein-containing complex binding | GO:MF | 6.28e-04 | *mew,Spn,zip,flr,Galphao,Su(var)205,Dys,Patronin,Unc-115a,tsr,Msp300,sals,BicD,sn,His2Av,if,Ndf,alpha-Spec,Shrm,CG6891,put,beta-Spec,JMJD6,mys,Cka,Mhcl,Caper,Svil,Usp2,Vinc,CG31712,DCP2,Galphas* |
| growth factor receptor binding | GO:MF | 8.58e-04 | *ed,bnl,Krn,vn,spi,stumps,drk* |
| ATP-dependent protein folding chaperone | GO:MF | 0.001 | *CCT8,CCT4,CCT3,CCT7,CCT1,CCT6,CCT2,CCT5* |
| protein folding chaperone | GO:MF | 0.001 | *CCT8,CCT4,CCT3,CCT7,CCT1,CCT6,CCT2,CCT5,Pfdn1* |
| signaling receptor binding | GO:MF | 0.001 | *sli,ed,Nlg1,mew,Spn,bnl,FER,Galphao,Grip,Src64B,Krn,Ack-like,vn,spi,Dad,stumps,Vang,dock,gbb,if,drk,mys,Stat92E,mav,Galphas,Fkbp12* |
| unfolded protein binding | GO:MF | 0.002 | *CG14207,CCT8,CCT4,CCT3,Pfdn6,CCT7,CCT1,Pfdn5,CCT6,CCT2,CCT5,Pfdn1* |
| small GTPase binding | GO:MF | 0.002 | *DAAM,rau,Magi,GEFmeso,Rip11,CG9135,spg,Evi5,pns,emb,BicD,Pak,Rok,Pak3* |
| kinase binding | GO:MF | 0.003 | *Msp300,stumps,drl,dome,Lk6,Mtl,dock,kibra,arm,Shrm,Ccm3,Cka,Pgam5,ci* |
| GTPase binding | GO:MF | 0.003 | *DAAM,rau,Magi,GEFmeso,Rip11,CG9135,spg,Evi5,pns,emb,BicD,Pak,Rok,Pak3* |
| transcription coregulator activity | GO:MF | 0.003 | *CG16779,Setd3,Pits,HDAC1,Bap60,Bgb,Snr1,mam,gro,Bin1,CtBP,Mad,arm,Gug,sbb,exd,CG42724,MED11,MED22,CoRest* |
| PDZ domain binding | GO:MF | 0.003 | *fz2,CG42672,rst,nkd* |
| identical protein binding | GO:MF | 0.004 | *Tl,rau,shg,trsn,rst,14-3-3zeta,fbp,stumps,drl,Meltrin,Septin2,Ten-m,snf,CtBP,Fmr1,ttk,Shrm,CadN,Pak3,ci* |
| protein homodimerization activity | GO:MF | 0.007 | *rau,shg,trsn,14-3-3zeta,drl,Septin2,Ten-m,CtBP,Fmr1,ttk,Shrm,CadN,Pak3,ci* |
| transcription corepressor activity | GO:MF | 0.008 | *CG16779,Pits,HDAC1,gro,Bin1,CtBP,Gug,sbb,CoRest* |
| protein dimerization activity | GO:MF | 0.009 | *rau,mew,shg,Mef2,Hand,trsn,14-3-3zeta,drl,dome,His3.3B,Septin2,Ten-m,tx,His2Av,gbb,CtBP,Fmr1,if,ttk,Taf13,exd,crp,Shrm,His3.3A,CadN,mys,Pak3,ci* |
| Wnt receptor activity | GO:MF | 0.011 | *fz2,drl,fz* |
| purine nucleotide binding | GO:MF | 0.012 | *Sur,sick,Strn-Mlck,w,htl,Act87E,Nuak,zip,FER,Uba2,sqa,Galphao,Src64B,CG3961,cana,Ack-like,betaTub60D,Hel25E,drl,Sps1,XNP,Pak,CG31140,CG7483,Lk6,Egfr,Septin2,Mtl,alphaTub84B,CtBP,CCT8,TER94,sgl,CCT4,Ndf,FoxK,Rm62,p38b,CCT3,Rpt5,put,vih,Rok,CCT7,hrg,alphaTub84D,Khc,CCT1,bves,Pak3,CCT6,Mhcl,Ras64B,Rpt1,CCT2,CCT5,Past1,Hmgcr,Fak,Arl1,Ilk,Galphas,cdi* |
| non-membrane spanning protein tyrosine kinase activity | GO:MF | 0.016 | *FER,Src64B,Ack-like,Fak* |
| single-stranded RNA binding | GO:MF | 0.019 | *how,Pep,Hrb87F,snf,CNBP,ssx,U2af50* |
| pre-mRNA binding | GO:MF | 0.019 | *snf,CG14641,U2af50,Ythdf,CG31712* |
| N-acetylgalactosaminyl-proteoglycan 3-beta-glucuronosyltransferase activity | GO:MF | 0.023 | *Chpf,Chsy* |
| alpha-catenin binding | GO:MF | 0.023 | *arm,Vinc* |
| extracellular matrix protein binding | GO:MF | 0.023 | *if,mys* |
| transforming growth factor beta receptor binding | GO:MF | 0.023 | *gbb,Fkbp12* |
| U1 snRNP binding | GO:MF | 0.023 | *Caper,CG31712* |
| transcription regulator activity | GO:MF | 0.024 | *H15,mid,tup,CG16779,bab2,Mef2,tin,Hand,ab,jumu,rib,Setd3,Pits,HDAC1,Bap60,Bgb,l(3)neo38,crol,Snr1,mam,gro,tx,Bin1,CG12391,CtBP,Mad,arm,Gug,ttk,sbb,exd,crp,Eip78C,FoxK,CG32772,NK7.1,CG32767,CG42724,MED11,MED22,Stat92E,ko,CoRest,sd,ewg,ci* |
| structural constituent of muscle | GO:MF | 0.024 | *Syn2,Tina-1,Dys,Dg* |
| extracellular matrix binding | GO:MF | 0.025 | *mew,Dg,if* |
| histone binding | GO:MF | 0.025 | *Su(var)205,Df31,HP1b,Dek,Nph,Brd7-9,Mapmodulin,Set,Stat92E,Chro,Nlp* |
| adenyl ribonucleotide binding | GO:MF | 0.027 | *Sur,sick,Strn-Mlck,w,htl,Act87E,Nuak,zip,FER,Uba2,sqa,Src64B,CG3961,cana,Ack-like,Hel25E,drl,Sps1,XNP,Pak,CG31140,CG7483,Lk6,Egfr,CCT8,TER94,CCT4,FoxK,Rm62,p38b,CCT3,Rpt5,put,vih,Rok,CCT7,hrg,Khc,CCT1,bves,Pak3,CCT6,Mhcl,Rpt1,CCT2,CCT5,Past1,Fak,Ilk,cdi* |
| RNA polymerase II-specific DNA-binding transcription factor binding | GO:MF | 0.027 | *tup,Bap60,dome,Snr1,Mad,Gug* |
| protein tyrosine kinase activity | GO:MF | 0.027 | *htl,FER,Src64B,Ack-like,Egfr,Fak,cdi* |
| ATP binding | GO:MF | 0.036 | *Sur,sick,Strn-Mlck,w,htl,Act87E,Nuak,zip,FER,Uba2,sqa,Src64B,CG3961,cana,Ack-like,Hel25E,drl,Sps1,XNP,Pak,CG31140,CG7483,Lk6,Egfr,CCT8,TER94,CCT4,FoxK,Rm62,p38b,CCT3,Rpt5,put,vih,Rok,CCT7,hrg,Khc,CCT1,Pak3,CCT6,Mhcl,Rpt1,CCT2,CCT5,Past1,Fak,Ilk,cdi* |
| protein kinase binding | GO:MF | 0.039 | *Msp300,stumps,drl,dome,Lk6,Mtl,dock,Ccm3,Cka,ci* |
| transcription coactivator activity | GO:MF | 0.039 | *Setd3,Bap60,Bgb,Snr1,mam,CtBP,Mad,arm,exd,MED11* |
| myosin light chain kinase activity | GO:MF | 0.047 | *Strn-Mlck,sqa* |
| cytokine receptor activity | GO:MF | 0.047 | *Tl,dome* |
| promoter-specific chromatin binding | GO:MF | 0.047 | *Psc,ttk,Taf10b* |
| glucuronosyl-N-acetylgalactosaminyl-proteoglycan 4-beta-N-acetylgalactosaminyltransferase activity | GO:MF | 0.047 | *Chpf,Chsy* |
| N6-methyladenosine-containing RNA reader activity | GO:MF | 0.047 | *Fmr1,Ythdf* |

### 1.14. Gene Ontology Enrichment Analysis: Notch cluster

Significant GO terms (p < 0.05) with associated genes

| GO Term | Source | P-value | Genes |
| --- | --- | --- | --- |
| regulation of metabolic process | GO:BP | 6.97e-15 | *fne,HmgD,CG13928,Sumo,HmgZ,Pep,scrt,Imp,Su(var)205,Dsp1,14-3-3epsilon,Hel25E,elav,l(3)neo38,Tet,CNBP,sqd,akirin,wech,brat,Hrb98DE,CtBP,Hrb87F,eIF4A,gro,SF2,Lam,rin,Hrb27C,dap,Cpsf6,glo,Rm62,lola,Non2,fs(1)h,E(spl)m7-HLH,lolal,HnRNP-K,zfh2,SC35,mts,lark,Hsc70-4,bel,gus,Ndf,Rbp1-like,Pabp2,Sin3A,Pits* |
| regulation of RNA metabolic process | GO:BP | 4.13e-14 | *fne,HmgD,CG13928,Sumo,HmgZ,Pep,scrt,Imp,Su(var)205,Dsp1,14-3-3epsilon,Hel25E,elav,l(3)neo38,Tet,sqd,akirin,Hrb98DE,CtBP,Hrb87F,eIF4A,gro,SF2,Hrb27C,Cpsf6,Rm62,lola,Non2,fs(1)h,E(spl)m7-HLH,lolal,HnRNP-K,zfh2,SC35,Ndf,Rbp1-like,Pabp2,Sin3A,Pits* |
| RNA splicing, via transesterification reactions | GO:BP | 1.51e-12 | *Pep,Imp,Hel25E,sqd,Hrb98DE,Hrb87F,eIF4A,SF2,Hrb27C,Cpsf6,SmB,Rm62,Ref1,HnRNP-K,SC35,lark,Hsc70-4,Rbp1-like,Pabp2* |
| nucleobase-containing compound metabolic process | GO:BP | 1.59e-10 | *fne,HmgD,His2Av,CG13928,Sumo,HmgZ,Pep,scrt,Imp,Su(var)205,Dsp1,14-3-3epsilon,Hel25E,elav,l(3)neo38,Tet,sqd,akirin,brat,Hrb98DE,CtBP,Hrb87F,eIF4A,gro,Tctp,SF2,Hrb27C,dap,Cpsf6,SmB,glo,Rm62,Ref1,lola,Non2,fs(1)h,E(spl)m7-HLH,lolal,HnRNP-K,zfh2,SC35,lark,Hsc70-4,Ndf,Rbp1-like,Pabp2,Sin3A,Pits* |
| regulation of alternative mRNA splicing, via spliceosome | GO:BP | 7.20e-10 | *Hel25E,sqd,Hrb98DE,Hrb87F,eIF4A,SF2,Cpsf6,Rm62,HnRNP-K,SC35,Rbp1-like* |
| negative regulation of biosynthetic process | GO:BP | 1.08e-07 | *CG13928,scrt,Su(var)205,Dsp1,elav,sqd,wech,brat,Hrb98DE,CtBP,gro,Lam,Hrb27C,Rm62,fs(1)h,E(spl)m7-HLH,lolal,Hsc70-4,bel,Pabp2,Sin3A,Pits* |
| animal organ morphogenesis | GO:BP | 2.08e-05 | *14-3-3epsilon,CNBP,Hrb98DE,CtBP,Hrb87F,Cam,Lam,rin,dap,ctp,lola,Gbeta13F,chic,lolal,zfh2,tsr,mts,gus,eff* |
| positive regulation of mRNA splicing, via spliceosome | GO:BP | 5.13e-05 | *Hrb98DE,Hrb87F,Hrb27C* |
| regulation of transcriptional start site selection at RNA polymerase II promoter | GO:BP | 9.50e-05 | *SF2,SC35,Rbp1-like* |
| male germ-line stem cell population maintenance | GO:BP | 0.002 | *His2Av,chic,bel* |
| pole plasm oskar mRNA localization | GO:BP | 0.003 | *sqd,Hrb27C,glo,chic* |
| positive regulation of translation | GO:BP | 0.003 | *CNBP,Hrb98DE,Hrb27C,lark* |
| positive regulation of cytoplasmic translation | GO:BP | 0.004 | *CNBP,lark* |
| mitotic cytokinesis | GO:BP | 0.004 | *Act5C,Act42A,chic,tsr* |
| negative regulation of oskar mRNA translation | GO:BP | 0.009 | *elav,Hrb27C* |
| negative regulation of RNA splicing | GO:BP | 0.009 | *sqd,Hrb98DE* |
| negative regulation of transcription by RNA polymerase II | GO:BP | 0.009 | *scrt,Su(var)205,Dsp1,CtBP,gro,E(spl)m7-HLH,Sin3A* |
| nucleosome assembly | GO:BP | 0.01 | *Df31,His3.3A,His3.3B,Set,His4r* |
| sperm DNA decondensation | GO:BP | 0.013 | *Nph,Nlp* |
| positive regulation of axon regeneration | GO:BP | 0.013 | *Imp,chic* |
| negative regulation of signal transduction | GO:BP | 0.013 | *His2Av,CtBP,eIF4A,gro,Cam,Gbeta13F,mts,Sin3A,eff* |
| asymmetric neuroblast division | GO:BP | 0.015 | *brat,Gbeta13F,mts* |
| mRNA export from nucleus | GO:BP | 0.016 | *Hel25E,sqd,Ref1* |
| blastoderm segmentation | GO:BP | 0.017 | *sqd,Hrb27C,glo,fs(1)h,chic,gus* |
| histoblast morphogenesis | GO:BP | 0.017 | *dap,chic* |
| female gonad development | GO:BP | 0.017 | *Hrb87F,tsr* |
| negative regulation of canonical Wnt signaling pathway | GO:BP | 0.019 | *CtBP,gro,Sin3A* |
| photoreceptor cell fate commitment | GO:BP | 0.019 | *14-3-3epsilon,rin,lola,mts* |
| dorsal/ventral axis specification, ovarian follicular epithelium | GO:BP | 0.021 | *sqd,rin* |
| negative regulation of smoothened signaling pathway | GO:BP | 0.026 | *Gbeta13F,mts,eff* |
| female germ-line stem cell asymmetric division | GO:BP | 0.027 | *eIF4A,eff* |
| positive regulation of cytokinesis, actomyosin contractile ring assembly | GO:BP | 0.031 | *Lam* |
| negative regulation of RNA polymerase II transcription preinitiation complex assembly | GO:BP | 0.031 | *Dsp1* |
| negative regulation of mRNA 3'-end processing | GO:BP | 0.031 | *elav* |
| regulation of light-activated channel activity | GO:BP | 0.031 | *Cam* |
| mRNA alternative polyadenylation | GO:BP | 0.031 | *Cpsf6* |
| positive regulation of FACT complex assembly | GO:BP | 0.031 | *Su(var)205* |
| positive regulation of spindle assembly | GO:BP | 0.031 | *Lam* |
| IRES-dependent translational initiation of linear mRNA | GO:BP | 0.031 | *CNBP* |
| mitotic nuclear membrane reassembly | GO:BP | 0.031 | *Lam* |
| late endosomal microautophagy | GO:BP | 0.031 | *Hsc70-4* |
| regulatory ncRNA-mediated post-transcriptional gene silencing | GO:BP | 0.031 | *Rm62,Hsc70-4,bel* |
| positive regulation of peptidoglycan recognition protein signaling pathway | GO:BP | 0.031 | *akirin,lola* |
| negative regulation of eclosion | GO:BP | 0.031 | *lark* |
| larval somatic muscle development | GO:BP | 0.031 | *akirin,lola,Sin3A* |
| cortical actin cytoskeleton organization | GO:BP | 0.036 | *Lam,chic,tsr* |
| somatic stem cell population maintenance | GO:BP | 0.036 | *His2Av,chic* |
| negative regulation of neuroblast proliferation | GO:BP | 0.036 | *brat,mts* |
| axonal fasciculation | GO:BP | 0.036 | *Nrt,Hsc70-4* |
| maintenance of protein location in cell | GO:BP | 0.036 | *Act5C,chic* |
| meiotic cell cycle | GO:BP | 0.036 | *Lam,Ran,dap,chic,tsr,eff* |
| positive regulation of Toll signaling pathway | GO:BP | 0.039 | *Sumo,mts* |
| positive regulation of Ras protein signal transduction | GO:BP | 0.041 | *Sumo,14-3-3epsilon* |
| female sex differentiation | GO:BP | 0.041 | *Hrb87F,tsr* |
| establishment of mitotic spindle orientation | GO:BP | 0.041 | *Ran,ctp* |
| asymmetric stem cell division | GO:BP | 0.044 | *eIF4A,eff* |
| mRNA binding | GO:MF | 1.76e-15 | *fne,Imp,Su(var)205,elav,CNBP,sqd,brat,Hrb98DE,Hrb87F,SF2,rin,Hrb27C,Cpsf6,glo,Rm62,Ref1,HnRNP-K,SC35,lark,Rbp1-like,Pabp2* |
| nucleic acid binding | GO:MF | 1.76e-15 | *fne,HmgD,His2Av,HmgZ,His3.3A,Pep,scrt,Imp,Su(var)205,D1,Dsp1,His3.3B,Hel25E,B52,elav,His4r,l(3)neo38,Tet,CNBP,sqd,brat,Hrb98DE,Hrb87F,eIF4A,SF2,rin,Hrb27C,Cpsf6,SmB,glo,Rm62,Ref1,lola,fs(1)h,Nph,E(spl)m7-HLH,lolal,Rsf1,HnRNP-K,zfh2,SC35,Nlp,lark,bel,eIF3f1,Ndf,Rbp1-like,Pabp2* |
| RNA binding | GO:MF | 5.84e-14 | *fne,Pep,Imp,Su(var)205,Hel25E,B52,elav,CNBP,sqd,brat,Hrb98DE,Hrb87F,eIF4A,SF2,rin,Hrb27C,Cpsf6,SmB,glo,Rm62,Ref1,Nph,Rsf1,HnRNP-K,SC35,Nlp,lark,bel,Rbp1-like,Pabp2* |
| single-stranded RNA binding | GO:MF | 1.58e-08 | *fne,Pep,elav,CNBP,Hrb98DE,Hrb87F,Hrb27C,Pabp2* |
| protein binding | GO:MF | 2.65e-07 | *Df31,His2Av,Sumo,His3.3A,Su(var)205,fax,Dsp1,14-3-3epsilon,His3.3B,Act5C,Set,His4r,akirin,wech,Act42A,brat,CtBP,eIF4A,gro,Nrt,Cam,Tctp,SF2,Lam,rin,Ran,Hrb27C,dap,ctp,glo,Rm62,lola,Fkbp12,fs(1)h,Gbeta13F,chic,Nph,E(spl)m7-HLH,lolal,Mapmodulin,HnRNP-K,Jupiter,tsr,mts,Nlp,Hsc70-4,gus,eIF3f1,Vap33,Pabp2,Sin3A,eff* |
| poly(G) binding | GO:MF | 9.62e-06 | *Hrb98DE,Hrb87F,Hrb27C* |
| mRNA 3'-UTR binding | GO:MF | 2.47e-05 | *Imp,sqd,brat,Hrb98DE,Hrb87F,Hrb27C* |
| DNA binding | GO:MF | 3.93e-05 | *HmgD,His2Av,HmgZ,His3.3A,Pep,scrt,Su(var)205,D1,Dsp1,His3.3B,His4r,l(3)neo38,Tet,Hrb98DE,Hrb87F,SF2,Hrb27C,lola,fs(1)h,E(spl)m7-HLH,lolal,zfh2,Ndf* |
| histone binding | GO:MF | 1.27e-04 | *Df31,Su(var)205,Set,fs(1)h,Nph,Mapmodulin,Nlp* |
| translation regulator activity | GO:MF | 1.62e-04 | *elav,CNBP,wech,brat,eIF4A,Hrb27C,eIF3f1* |
| DNA binding, bending | GO:MF | 3.00e-04 | *HmgD,HmgZ,Dsp1* |
| translation repressor activity | GO:MF | 3.93e-04 | *elav,wech,brat,Hrb27C* |
| RNA helicase activity | GO:MF | 8.12e-04 | *Hel25E,eIF4A,rin,Rm62,bel* |
| ATP-dependent activity, acting on RNA | GO:MF | 8.29e-04 | *Hel25E,eIF4A,rin,Rm62,bel* |
| structural constituent of cytoskeleton | GO:MF | 8.97e-04 | *betaTub56D,alphaTub84B,Lam,Jupiter* |
| protein dimerization activity | GO:MF | 0.003 | *His2Av,His3.3A,14-3-3epsilon,His3.3B,His4r,CtBP,ctp,E(spl)m7-HLH,lolal,Sin3A* |
| satellite DNA binding | GO:MF | 0.003 | *Su(var)205,D1* |
| minor groove of adenine-thymine-rich DNA binding | GO:MF | 0.003 | *HmgD,D1* |
| transcription coregulator binding | GO:MF | 0.004 | *14-3-3epsilon,CtBP,E(spl)m7-HLH* |
| chromatin binding | GO:MF | 0.006 | *Su(var)205,Set,Lam,fs(1)h,Nph,Nlp,Ndf,Sin3A* |
| helicase activity | GO:MF | 0.007 | *Hel25E,eIF4A,rin,Rm62,bel* |
| transcription factor binding | GO:MF | 0.007 | *Dsp1,14-3-3epsilon,CtBP,gro,E(spl)m7-HLH,HnRNP-K* |
| transcription corepressor activity | GO:MF | 0.007 | *CtBP,gro,Sin3A,Pits* |
| poly(U) RNA binding | GO:MF | 0.014 | *fne,elav* |
| mRNA 5'-UTR binding | GO:MF | 0.014 | *Hrb98DE,Hrb27C* |
| protein-macromolecule adaptor activity | GO:MF | 0.024 | *Su(var)205,akirin,wech,CtBP,gro,Gbeta13F,Sin3A,Pits* |
| protein domain specific binding | GO:MF | 0.026 | *gro,ctp,lolal,Vap33* |
| protein heterodimerization activity | GO:MF | 0.026 | *His2Av,His3.3A,14-3-3epsilon,His3.3B,His4r,Sin3A* |
| poly-pyrimidine tract binding | GO:MF | 0.029 | *fne,elav* |
| DNA-binding transcription factor binding | GO:MF | 0.029 | *Dsp1,CtBP,gro,E(spl)m7-HLH* |
| sequence-specific DNA binding | GO:MF | 0.032 | *HmgD,HmgZ,scrt,Su(var)205,D1,l(3)neo38,Hrb98DE,Hrb87F,fs(1)h,E(spl)m7-HLH,zfh2* |
| single-stranded DNA binding | GO:MF | 0.032 | *Pep,Dsp1,Hrb27C* |
| molecular adaptor activity | GO:MF | 0.035 | *Su(var)205,akirin,wech,CtBP,gro,Gbeta13F,Sin3A,Pits* |
| type I transforming growth factor beta receptor binding | GO:MF | 0.04 | *Fkbp12* |
| CRD domain binding | GO:MF | 0.04 | *gro* |
| HMG box domain binding | GO:MF | 0.04 | *gro* |

### 1.15. Gene Ontology Enrichment Analysis: dead

Significant GO terms (p < 0.05) with associated genes

| GO Term | Source | P-value | Genes |
| --- | --- | --- | --- |
| proton transmembrane transport | GO:BP | 2.84e-12 | *mt:ND5,mt:ND4,mt:ATPase6,mt:CoII,mt:Cyt-b,mt:CoIII,mt:CoI* |
| electron transport coupled proton transport | GO:BP | 3.31e-08 | *mt:ND5,mt:ND4,mt:CoI* |
| mitochondrial electron transport, NADH to ubiquinone | GO:BP | 3.13e-07 | *mt:ND3,mt:ND5,mt:ND6,mt:ND2* |
| mitochondrial electron transport, cytochrome c to oxygen | GO:BP | 3.60e-06 | *mt:CoII,mt:CoIII,mt:CoI* |
| electron transfer activity | GO:MF | 1.19e-23 | *mt:ND3,mt:ND5,mt:ND4,mt:CoII,mt:Cyt-b,mt:CoIII,mt:CoI,mt:ND6,mt:ND1,mt:ND2* |
| proton transmembrane transporter activity | GO:MF | 6.95e-23 | *mt:ND3,mt:ND5,mt:ND4,mt:ATPase6,mt:CoII,mt:Cyt-b,mt:CoIII,mt:CoI,mt:ND6,mt:ND1,mt:ND2* |
| inorganic cation transmembrane transporter activity | GO:MF | 1.81e-17 | *mt:ND3,mt:ND5,mt:ND4,mt:ATPase6,mt:CoII,mt:Cyt-b,mt:CoIII,mt:CoI,mt:ND6,mt:ND1,mt:ND2* |
| monoatomic cation transmembrane transporter activity | GO:MF | 4.28e-17 | *mt:ND3,mt:ND5,mt:ND4,mt:ATPase6,mt:CoII,mt:Cyt-b,mt:CoIII,mt:CoI,mt:ND6,mt:ND1,mt:ND2* |
| monoatomic ion transmembrane transporter activity | GO:MF | 7.30e-16 | *mt:ND3,mt:ND5,mt:ND4,mt:ATPase6,mt:CoII,mt:Cyt-b,mt:CoIII,mt:CoI,mt:ND6,mt:ND1,mt:ND2* |
| active transmembrane transporter activity | GO:MF | 9.41e-16 | *mt:ND3,mt:ND5,mt:ND4,mt:CoII,mt:Cyt-b,mt:CoIII,mt:CoI,mt:ND6,mt:ND1,mt:ND2* |
| NADH dehydrogenase (ubiquinone) activity | GO:MF | 1.08e-14 | *mt:ND3,mt:ND5,mt:ND4,mt:ND6,mt:ND1,mt:ND2* |
| oxidoreductase activity, acting on NAD(P)H | GO:MF | 4.04e-13 | *mt:ND3,mt:ND5,mt:ND4,mt:ND6,mt:ND1,mt:ND2* |
| transmembrane transporter activity | GO:MF | 4.58e-13 | *mt:ND3,mt:ND5,mt:ND4,mt:ATPase6,mt:CoII,mt:Cyt-b,mt:CoIII,mt:CoI,mt:ND6,mt:ND1,mt:ND2* |
| transporter activity | GO:MF | 8.22e-13 | *mt:ND3,mt:ND5,mt:ND4,mt:ATPase6,mt:CoII,mt:Cyt-b,mt:CoIII,mt:CoI,mt:ND6,mt:ND1,mt:ND2* |
| oxidoreductase activity | GO:MF | 7.76e-12 | *mt:ND3,mt:ND5,mt:ND4,mt:CoII,mt:Cyt-b,mt:CoIII,mt:CoI,mt:ND6,mt:ND1,mt:ND2* |
| cytochrome-c oxidase activity | GO:MF | 1.35e-09 | *mt:CoII,mt:CoIII,mt:CoI* |
| catalytic activity | GO:MF | 2.01e-05 | *mt:ND3,mt:ND5,mt:ND4,mt:ATPase6,mt:CoII,mt:Cyt-b,mt:CoIII,mt:CoI,mt:ND6,mt:ND1,mt:ND2* |
| ubiquinol-cytochrome-c reductase activity | GO:MF | 0.007 | *mt:Cyt-b* |
| ubiquinone binding | GO:MF | 0.007 | *mt:ND4* |
| NADH dehydrogenase activity | GO:MF | 0.015 | *mt:ND5* |
| quinone binding | GO:MF | 0.016 | *mt:ND4* |
| proton-transporting ATP synthase activity, rotational mechanism | GO:MF | 0.035 | *mt:ATPase6* |
| copper ion binding | GO:MF | 0.037 | *mt:CoII* |
| proton channel activity | GO:MF | 0.037 | *mt:ATPase6* |

### 1.16. Gene Ontology Enrichment Analysis: dying bap-PC

Significant GO terms (p < 0.05) with associated genes

| GO Term | Source | P-value | Genes |
| --- | --- | --- | --- |
| animal organ morphogenesis | GO:BP | 5.74e-15 | *hbs,rpr,hid,by,tin,prc,pk,trol,stg,dally,cv-2,ftz-f1,bdl,Nedd4,Fas2,CG42674,Nrg,rdx,caps,heph,stck,fz,dnr1,RhoGEF64C,Grip,ex,LamC,Spn88Ea,unc-5,Cip4,Moe,Frl,foxo,Dif,flw,bun,Rac2,tup,NetB,Lac,corto,trn,shot,sqh,arr,Mob2,unk,cindr,Fhos,Hipk,Jra,Vps4,if,cora,Amph,Ilk,parvin,Mkp3,ATP6AP2,alphaSnap,Rab5,Gprk2,RhoGAP19D,cic,kibra,ics,puc,ci* |
| negative regulation of signal transduction | GO:BP | 2.22e-07 | *Sulf1,CG7378,trbl,trol,dally,scyl,Nedd4,Fas2,apt,rdx,stck,wdp,dnr1,chrb,ex,nkd,Atg9,RapGAP1,foxo,flw,sqh,ken,Ubc6,Hipk,Rheb,hppy,cact,Mkp3,raskol,Gprk2,BI-1,Cul1,ics,puc,ci* |
| cell surface receptor signaling pathway | GO:BP | 2.13e-06 | *Sulf1,hbs,egr,trbl,trol,dally,cv-2,Mp,Nedd4,Fas2,apt,RalGPS,rdx,stck,wdp,fz,nkd,unc-5,foxo,Dif,flw,Rac2,Ehbp1,HDAC4,Toll-7,sqh,arr,ken,Fit1,scb,Hipk,Rheb,Vps4,if,hppy,Myd88,Ilk,cact,Mkp3,ATP6AP2,E(spl)mbeta-HLH,E(spl)m3-HLH,Gprk2,cic,Cul1,Pitslre,ics,puc,ci* |
| motor neuron axon guidance | GO:BP | 8.02e-06 | *Sulf1,trol,dally,Mp,Fas2,Nrg,caps,unc-5,Rac2,tup,NetB,trn,cher* |
| larval heart development | GO:BP | 6.85e-05 | *prc,loh,Hand,scb* |
| larval midgut cell programmed cell death | GO:BP | 9.23e-05 | *rpr,hid,Atg9,Atg18a,Atg8a,Atg17,Ubi-p63E* |
| apposition of dorsal and ventral imaginal disc-derived wing surfaces | GO:BP | 1.94e-04 | *by,stck,if,Ilk,parvin,ics* |
| biological process involved in interspecies interaction between organisms | GO:BP | 3.12e-04 | *egr,GILT2,Nrg,dnr1,Fer2LCH,Spn88Ea,vir-1,drpr,Fer1HCH,Atg18a,DIP1,wun,foxo,Dif,mtd,Rac2,Toll-7,Rab1,Fit1,CG5390,Rab2,CBP,Rab7,Jra,Vps4,Myd88,cact,Rab5,Vps60,Gprk2,shrb,Cul1,Pitslre,puc* |
| viral process | GO:BP | 4.70e-04 | *Rab1,Rab2,Rab7,Jra,Vps4,Rab5,Vps60,shrb* |
| striated muscle cell differentiation | GO:BP | 5.47e-04 | *hbs,tin,Nedd4,Rac2,Bsg,tmod,Tm2,Zasp52,if,cher,hts* |
| regulation of metabolic process | GO:BP | 0.001 | *hbs,rpr,hid,egr,Eip74EF,Spn43Ab,tin,bap,CG5001,trbl,chinmo,CG9650,ps,CG3726,ftz-f1,Atf3,Spn31A,stv,Nedd4,Glut4EF,Pur-alpha,Hand,apt,rdx,heph,dnr1,CG9932,CG32369,ex,LamC,Spn88Ea,Syx17,Svip,UbcE2H,Atg18a,DIP1,foxo,Dif,Octbeta2R,tai,bun,Atg8a,CG3662,tup,Wbp2,PAN3,REPTOR-BP,Paip2,hzg,REPTOR,HDAC4,cwo,maf-S,corto,Toll-7,ken,AGO1,CG6770,Mob2,tna,Ubc6,BNIP3,Sox14,Smg6,Hipk,Rheb,CG14073,Atg17,Jra,Pdp1,Max,Myd88,CG13124,cact,Not1,gce,Su(Tpl),E(spl)mbeta-HLH,Fdx2,E(spl)m3-HLH,sordd1,Gprk2,BI-1,cic,CG12769,BtbVII,Cul1,kibra,Pitslre,ics,puc,ctrip,ci* |
| wing disc dorsal/ventral pattern formation | GO:BP | 0.001 | *Sulf1,dally,AnxB9,drpr,Rab7,CG14073,Su(Tpl),cic* |
| synaptic vesicle fusion to presynaptic active zone membrane | GO:BP | 0.001 | *Syx1A,Snap24,alphaSnap,Snap29* |
| juvenile hormone mediated signaling pathway | GO:BP | 0.002 | *Chd64,ftz-f1,gce* |
| segment polarity determination | GO:BP | 0.002 | *dally,rdx,fz,nkd,arr,AGO1,ci* |
| vacuole organization | GO:BP | 0.003 | *Atg9,Svip,Atg18a,Atg8a,Rab1,Rab7,Atg17,Vps4* |
| cell adhesion involved in heart morphogenesis | GO:BP | 0.003 | *prc,Nrg,Lac,cora* |
| muscle cell development | GO:BP | 0.003 | *Nedd4,Bsg,tmod,Tm2,Zasp52,if,cher,hts* |
| homophilic cell adhesion via plasma membrane adhesion molecules | GO:BP | 0.004 | *hbs,bdl,Fas2,caps,fz,Lac,Bsg* |
| catabolic process | GO:BP | 0.004 | *SP1029,rpr,hid,Eip74EF,CG8353,Idgf2,trbl,CG17896,CG6847,Nedd4,rdx,CG8360,dnr1,ex,Syx17,Atg9,Svip,CG6966,rudhira,UbcE2H,Atg18a,DIP1,foxo,CG33090,Rab18,Atg8a,PAN3,Plc21C,Toll-7,gzl,Rab1,Vps37B,AGO1,Rnf11,Ubi-p5E,Ubc6,CG11658,Smg6,Gale,Rab7,Rheb,Atg17,Vps4,Ubi-p63E,Not1,sordd1,shrb,BI-1,Snap29,Cul1,CG7461,ctrip,CtsF* |
| negative regulation of smoothened signaling pathway | GO:BP | 0.005 | *Sulf1,Nedd4,rdx,flw,sqh,Gprk2,Cul1* |
| septate junction assembly | GO:BP | 0.005 | *CG44325,Nrg,CG9628,wun,Lac,pasi2* |
| head involution | GO:BP | 0.005 | *hid,scyl,stck,chrb,Rac2,tup* |
| synaptic vesicle priming | GO:BP | 0.006 | *Snap24,alphaSnap,Snap29,unc-13* |
| dorsal appendage formation | GO:BP | 0.006 | *jvl,bun,Rac2,Jra,cact,cic,puc* |
| behavioral response to ethanol | GO:BP | 0.006 | *Fas2,apt,CG17734,rut,scb,hppy,cher* |
| positive regulation of neuroblast proliferation | GO:BP | 0.006 | *trol,bun,Rheb,E(spl)mbeta-HLH,E(spl)m3-HLH* |
| iron ion import across plasma membrane | GO:BP | 0.007 | *Fer2LCH,Fer1HCH* |
| detoxification of iron ion | GO:BP | 0.007 | *Fer2LCH,Fer1HCH* |
| cell surface pattern recognition receptor signaling pathway | GO:BP | 0.007 | *Toll-7,Myd88* |
| positive regulation of substrate adhesion-dependent cell spreading | GO:BP | 0.007 | *stck,Fhos* |
| imaginal disc-derived male genitalia morphogenesis | GO:BP | 0.009 | *rpr,Fas2,Rab5,puc* |
| mushroom body development | GO:BP | 0.01 | *chinmo,ftz-f1,Fas2,Nrg,Frl,bun,shot,orion* |
| antimicrobial humoral immune response mediated by antimicrobial peptide | GO:BP | 0.01 | *dnr1,Dif,Toll-7,Myd88,cact,Gprk2,Cul1,Pitslre* |
| membrane fusion | GO:BP | 0.011 | *Syx17,Svip,Syx1A,Snap24,Rab7,alphaSnap,Rab5,Snap29* |
| signal release | GO:BP | 0.013 | *Rph,CG9650,Esyt2,Syx17,Syx1A,Snap24,Rab1,Rnf11,alphaSnap,Snap29,unc-13* |
| positive regulation of integrin-mediated signaling pathway | GO:BP | 0.015 | *Mp,stck* |
| autophagy of mitochondrion | GO:BP | 0.015 | *Atg9,Atg18a,Atg8a,Ubc6,Atg17* |
| positive regulation of JNK cascade | GO:BP | 0.015 | *rpr,egr,rdx,Atg9,hppy* |
| cortical microtubule organization | GO:BP | 0.015 | *Moe,shot* |
| positive regulation of transcription by RNA polymerase II | GO:BP | 0.015 | *Eip74EF,tin,bap,ftz-f1,Hand,apt,CG9932,foxo,Dif,tai,tup,Wbp2,maf-S,Sox14,Jra,Pdp1,Max,Myd88,gce,Su(Tpl),CG12769,ci* |
| cytidine deamination | GO:BP | 0.015 | *CG8353,CG8360* |
| intracellular sequestering of iron ion | GO:BP | 0.015 | *Fer2LCH,Fer1HCH* |
| female germline ring canal formation, actin assembly | GO:BP | 0.015 | *cher,hts* |
| dorsal closure | GO:BP | 0.016 | *stck,Rac2,tup,shot,scb,Jra,cora,Rab5,puc* |
| genital disc morphogenesis | GO:BP | 0.016 | *rpr,Fas2,Rab5,puc* |
| sarcomere organization | GO:BP | 0.018 | *Bsg,Tm2,if,cher,hts* |
| glycophagy | GO:BP | 0.019 | *Atg9,Atg18a,Atg17* |
| engulfment of apoptotic cell | GO:BP | 0.023 | *egr,drpr,Rac2* |
| epidermal cell differentiation | GO:BP | 0.024 | *pk,fz,Cip4,sqh,cora,ATP6AP2* |
| long-term memory | GO:BP | 0.024 | *mol,HDAC4,arr,Mob2,cher,ATP6AP2,Pkc98E* |
| embryonic anterior midgut (ectodermal) morphogenesis | GO:BP | 0.025 | *tin,puc* |
| regulation of basement membrane organization | GO:BP | 0.025 | *Ndg,Ehbp1* |
| chorion micropyle formation | GO:BP | 0.025 | *Jra,puc* |
| organelle fusion | GO:BP | 0.026 | *Syx17,Svip,Syx1A,Snap24,Rab7,alphaSnap,Rab5,Snap29* |
| regulation of RNA metabolic process | GO:BP | 0.027 | *Eip74EF,tin,bap,CG5001,chinmo,CG9650,ps,CG3726,ftz-f1,Atf3,Glut4EF,Pur-alpha,Hand,apt,CG9932,foxo,Dif,Octbeta2R,tai,bun,tup,Wbp2,PAN3,REPTOR-BP,hzg,REPTOR,HDAC4,cwo,maf-S,ken,AGO1,CG6770,tna,Ubc6,Sox14,Smg6,CG14073,Jra,Pdp1,Max,Myd88,cact,Not1,gce,Su(Tpl),E(spl)mbeta-HLH,E(spl)m3-HLH,BI-1,cic,CG12769,BtbVII,kibra,ci* |
| late endosome to vacuole transport via multivesicular body sorting pathway | GO:BP | 0.028 | *CG30423,Vps60,shrb* |
| maintenance of epithelial integrity, open tracheal system | GO:BP | 0.028 | *Lac,if,Mkp3* |
| receptor clustering | GO:BP | 0.028 | *Fur1,Grip,Rab2* |
| positive regulation of axon guidance | GO:BP | 0.028 | *pk,fz,Rheb* |
| negative regulation of cell population proliferation | GO:BP | 0.03 | *wdp,ex,foxo,CG6770,Rab5,Pkc98E,kibra* |
| cortical actin cytoskeleton organization | GO:BP | 0.033 | *Moe,Frl,flw,Fhos,ghi,vib* |
| ubiquitin-dependent protein catabolic process via the multivesicular body sorting pathway | GO:BP | 0.033 | *Vps37B,Vps4,shrb* |
| positive regulation of smoothened signaling pathway | GO:BP | 0.033 | *Sulf1,trol,dally,Hipk,Gprk2* |
| positive regulation of neuron remodeling | GO:BP | 0.033 | *Sox14,orion,Cul1* |
| protein localization to endosome | GO:BP | 0.037 | *Grip,Moe* |
| embryonic digestive tract morphogenesis | GO:BP | 0.037 | *tin,puc* |
| response to tumor cell | GO:BP | 0.037 | *egr,Myd88* |
| actin crosslink formation | GO:BP | 0.037 | *IRSp53,cher* |
| negative regulation of insulin receptor signaling pathway | GO:BP | 0.038 | *trbl,foxo,Mkp3,Cul1* |
| regulation of tube length, open tracheal system | GO:BP | 0.038 | *Fas2,fz,sqh,pasi2* |
| short-term memory | GO:BP | 0.038 | *Fas2,rut,scb,ATP6AP2* |
| negative regulation of antimicrobial peptide production | GO:BP | 0.039 | *dnr1,cact,Cul1* |
| positive regulation of axon extension | GO:BP | 0.039 | *pk,fz,shot* |
| positive regulation of catalytic activity | GO:BP | 0.039 | *hbs,rpr,hid,rk,Sirup,RalGPS,RapGAP1,RhoGAP93B,RhoGAP18B,Atg17,Fdx2* |
| heterophilic cell-cell adhesion via plasma membrane cell adhesion molecules | GO:BP | 0.042 | *hbs,Nrt,scb,if* |
| establishment of glial blood-brain barrier | GO:BP | 0.045 | *Nrg,cora,pasi2* |
| wing disc anterior/posterior pattern formation | GO:BP | 0.045 | *Sulf1,dally,ci* |
| protein localization to phagophore assembly site | GO:BP | 0.05 | *Atg9,Atg18a* |
| protein localization involved in establishment of planar polarity | GO:BP | 0.05 | *Shrm,ATP6AP2* |
| positive regulation of cysteine-type endopeptidase activity | GO:BP | 0.05 | *rpr,hid* |
| detection of temperature stimulus involved in sensory perception of pain | GO:BP | 0.05 | *pain,Ncc69* |
| ommatidial rotation | GO:BP | 0.05 | *hbs,pk,fz,Frl* |
| heart formation | GO:BP | 0.05 | *Nedd4,ci* |
| clathrin-dependent endocytosis involved in vitellogenesis | GO:BP | 0.05 | *Rab7,Rab5* |
| protein binding | GO:MF | 1.96e-11 | *hbs,rpr,rdo,CG17839,hid,egr,by,CG10359,CG8353,rk,Rph,IRSp53,CG4393,CG5001,pain,Idgf2,trbl,pk,trol,CG45263,chinmo,dally,Chd64,cv-2,CG3726,ftz-f1,Drat,bdl,Atf3,CG31076,stv,Cp110,Nedd4,Fas2,AnxB9,mlt,Pur-alpha,olf186-M,Hand,apt,Nrg,rdx,caps,stck,wdp,fz,CG8360,CG32369,Grip,ex,nkd,unc-5,Syx17,Liprin-gamma,Cip4,rut,l(3)05822,jvl,drpr,Moe,mol,Nrt,CG6966,Frl,Syx1A,Atg18a,DIP1,Oatp74D,foxo,Plp,Dif,tai,flw,Rab18,bun,Atg8a,Rac2,tup,PAN3,REPTOR-BP,Paip2,NetB,REPTOR,Ehbp1,Lac,HDAC4,Snap24,RhoGAP93B,cwo,maf-S,corto,Toll-7,trn,shot,Septin2,Rab1,sqh,arr,ken,RhoGAP18B,Fit1,AGO1,Mob2,scb,Wdr62,unk,CG3402,cindr,Bsg,Rab2,Ubi-p5E,Ubc6,tmod,Tm2,BNIP3,slo,Zasp52,CG11658,Smg6,LRP1,Shrm,Fhos,orion,Hipk,Rheb,CG14073,pns,Atg17,Jra,if,Max,Vha13,Myd88,LTV1,cora,toc,nudE,Amph,CG30423,CG13124,Ubi-p63E,Ilk,cact,Not1,cher,parvin,hts,gce,Su(Tpl),ATP6AP2,E(spl)mbeta-HLH,alphaSnap,Rab5,E(spl)m3-HLH,sordd1,Rab4,shrb,BI-1,RhoGAP19D,cic,NKAIN,BtbVII,Snap29,Cul1,kibra,milt,ics,puc,sbr,ci,unc-13* |
| cell adhesion molecule binding | GO:MF | 1.48e-04 | *hbs,CG45263,bdl,Fas2,Nrg,Moe,Nrt,Fit1,scb,cindr,Bsg,if* |
| calcium-dependent phospholipid binding | GO:MF | 0.002 | *AnxB10,Rph,AnxB9,Esyt2,AnxB11* |
| phospholipid binding | GO:MF | 0.002 | *rpr,AnxB10,Rph,IRSp53,AnxB9,Esyt2,Cip4,drpr,Moe,Atg18a,AnxB11,Myd88,vib,Amph,Gprk2,unc-13* |
| lipid binding | GO:MF | 0.002 | *rpr,AnxB10,Rph,IRSp53,CG17896,AnxB9,Esyt2,Cip4,drpr,Moe,Atg18a,AnxB11,Myd88,vib,Amph,gce,Clic,Gprk2,CG7461,unc-13* |
| protein dimerization activity | GO:MF | 0.006 | *rpr,trbl,ftz-f1,Hand,rdx,Liprin-gamma,tai,bun,REPTOR-BP,REPTOR,Lac,cwo,maf-S,corto,Septin2,scb,Shrm,Jra,if,Max,gce,E(spl)mbeta-HLH,E(spl)m3-HLH,ci* |
| cell adhesion mediator activity | GO:MF | 0.006 | *hbs,bdl,Fas2,Nrt,Bsg* |
| MAP kinase phosphatase activity | GO:MF | 0.014 | *CG7378,Mkp3,puc* |
| actin binding | GO:MF | 0.019 | *by,Chd64,Moe,Frl,shot,Tm2,Zasp52,Shrm,Fhos,cora,cher,parvin,hts* |
| DNA-binding transcription activator activity | GO:MF | 0.02 | *Eip74EF,ftz-f1,Hand,apt,foxo,Dif,Pdp1,gce,CG12769,ci* |
| diacylglycerol-dependent serine/threonine kinase activity | GO:MF | 0.024 | *PKD,Pkcdelta,Pkc98E* |
| identical protein binding | GO:MF | 0.024 | *rpr,CG8353,trbl,Pur-alpha,rdx,CG8360,Liprin-gamma,bun,REPTOR-BP,Lac,corto,Septin2,BNIP3,Shrm,ci* |
| MAP kinase tyrosine phosphatase activity | GO:MF | 0.024 | *Mkp3,puc* |
| DNA-binding transcription factor activity | GO:MF | 0.024 | *Eip74EF,tin,bap,CG9650,ftz-f1,Atf3,Glut4EF,Pur-alpha,Hand,apt,CG9932,foxo,Dif,Octbeta2R,tup,REPTOR-BP,REPTOR,cwo,maf-S,ken,Sox14,Jra,Pdp1,Max,gce,E(spl)mbeta-HLH,E(spl)m3-HLH,cic,CG12769,ci* |
| protein homodimerization activity | GO:MF | 0.024 | *rpr,trbl,rdx,Liprin-gamma,bun,REPTOR-BP,Lac,corto,Septin2,Shrm,ci* |
| RNA polymerase II transcription regulatory region sequence-specific DNA binding | GO:MF | 0.024 | *tin,bap,CG9650,Chd64,ftz-f1,Atf3,Glut4EF,Pur-alpha,Hand,apt,CG9932,foxo,Dif,tup,REPTOR,cwo,maf-S,ken,Sox14,Jra,Pdp1,gce,E(spl)mbeta-HLH,E(spl)m3-HLH,cic,CG12769,ci* |
| protein tyrosine/threonine phosphatase activity | GO:MF | 0.024 | *Mkp3,puc* |
| cis-regulatory region sequence-specific DNA binding | GO:MF | 0.029 | *tin,bap,CG9650,Chd64,ftz-f1,Atf3,Glut4EF,CG9932,foxo,Dif,tup,cwo,maf-S,ken,Sox14,Jra,Pdp1,gce,Su(Tpl),E(spl)mbeta-HLH,E(spl)m3-HLH,CG12769,ci* |
| GTPase activity | GO:MF | 0.029 | *Rab18,Rac2,eEF1alpha1,Septin2,Rab1,Rab2,Non1,CG2017,Rab7,Rheb,Rab5,Rab4* |
| DNA-binding transcription activator activity, RNA polymerase II-specific | GO:MF | 0.033 | *Eip74EF,ftz-f1,Hand,apt,foxo,Dif,Pdp1,CG12769,ci* |
| sequence-specific double-stranded DNA binding | GO:MF | 0.035 | *tin,bap,CG9650,Chd64,ftz-f1,Atf3,Glut4EF,Pur-alpha,Hand,apt,CG9932,foxo,Dif,tup,REPTOR,cwo,maf-S,ken,Sox14,Jra,Pdp1,Max,gce,Su(Tpl),E(spl)mbeta-HLH,E(spl)m3-HLH,cic,CG12769,ci* |
| calcium ion binding | GO:MF | 0.035 | *AnxB10,AnxB9,Nrg,Esyt2,regucalcin,Ndg,AnxB11,Plc21C,shot,sqh,CBP,LRP1,Clic,unc-13* |
| transcription cis-regulatory region binding | GO:MF | 0.035 | *tin,bap,CG9650,Chd64,ftz-f1,Atf3,Glut4EF,Pur-alpha,Hand,apt,CG9932,foxo,Dif,tup,REPTOR,cwo,maf-S,ken,Sox14,Jra,Pdp1,gce,Su(Tpl),E(spl)mbeta-HLH,E(spl)m3-HLH,cic,CG12769,ci* |
| cytoskeletal protein binding | GO:MF | 0.035 | *by,Chd64,Nedd4,AnxB9,mlt,Moe,Frl,shot,sqh,tmod,Tm2,Zasp52,Shrm,Fhos,cora,toc,nudE,cher,parvin,hts,milt* |
| RNA polymerase II cis-regulatory region sequence-specific DNA binding | GO:MF | 0.038 | *tin,bap,CG9650,Chd64,ftz-f1,Atf3,Glut4EF,CG9932,foxo,Dif,tup,cwo,maf-S,ken,Sox14,Jra,Pdp1,gce,E(spl)mbeta-HLH,E(spl)m3-HLH,CG12769,ci* |
| cytidine deaminase activity | GO:MF | 0.04 | *CG8353,CG8360* |
| proton-transporting ATPase activity, rotational mechanism | GO:MF | 0.04 | *VhaM9.7-b,Vha36-1,VhaSFD,Vha13,VhaPPA1-1* |
| GTP binding | GO:MF | 0.04 | *Rab18,Rac2,eEF1alpha1,Septin2,Rab1,Rab2,Non1,CG2017,Rab7,Rheb,Rab5,AdSS,Rab4* |
| calcium-dependent protein serine/threonine kinase activity | GO:MF | 0.04 | *PKD,Pkcdelta,Pkc98E* |
| MAP kinase tyrosine/serine/threonine phosphatase activity | GO:MF | 0.04 | *Mkp3,puc* |
| sequence-specific DNA binding | GO:MF | 0.04 | *Eip74EF,tin,bap,CG9650,Chd64,ftz-f1,Atf3,Glut4EF,Pur-alpha,Hand,apt,CG9932,foxo,Dif,tup,REPTOR,cwo,maf-S,ken,Sox14,Smg6,Jra,Pdp1,Max,gce,Su(Tpl),E(spl)mbeta-HLH,E(spl)m3-HLH,cic,CG12769,ci* |
| molecular adaptor activity | GO:MF | 0.044 | *by,rdx,koi,Syx17,Syx1A,Plp,tai,Wbp2,Snap24,shot,Septin2,tna,CG14073,Atg17,Myd88,Not1,alphaSnap,Snap29,Cul1,kibra,milt* |
| SNARE binding | GO:MF | 0.044 | *Syx17,Syx1A,Snap24,alphaSnap,Snap29,unc-13* |
| guanyl ribonucleotide binding | GO:MF | 0.049 | *Rab18,Rac2,eEF1alpha1,Septin2,Rab1,Rab2,Non1,CG2017,Rab7,Rheb,Rab5,AdSS,Rab4* |
| guanyl nucleotide binding | GO:MF | 0.049 | *Rab18,Rac2,eEF1alpha1,Septin2,Rab1,Rab2,Non1,CG2017,Rab7,Rheb,Rab5,AdSS,Rab4* |
| actin filament binding | GO:MF | 0.049 | *by,Chd64,Frl,Tm2,Shrm,Fhos,cher,hts* |

### 1.17. Gene Ontology Enrichment Analysis: plasmatocytes

Significant GO terms (p < 0.05) with associated genes

| GO Term | Source | P-value | Genes |
| --- | --- | --- | --- |
| endoplasmic reticulum to Golgi vesicle-mediated transport | GO:BP | 1.15e-09 | *KdelR,bai,Sec24CD,CHOp24,ergic53,Sar1,eca,p24-1,Rab1,Sec13,epsilonCOP,CG3652,Sec23,Bet1,Sec22,loj,deltaCOP,CG13887* |
| protein localization to endoplasmic reticulum | GO:BP | 3.82e-08 | *KdelR,TRAM,Spase22-23,Spase12,Sec61beta,CG8860,Spase25,CG5885,Sec61alpha,SrpRbeta,SrpRalpha,CG13887* |
| signal release | GO:BP | 2.01e-06 | *nemy,cpx,bai,CHOp24,Syb,Chc,eca,Scamp,Snx3,p24-1,Arf1,Rab1,AdipoR,Cdc42,shi,Sec22,Syx1A,Arf6,sky* |
| animal organ morphogenesis | GO:BP | 2.29e-06 | *Col4a1,Arpc3B,Pvr,sty,LanB2,srp,pnt,LanA,sigmar,fog,RhoGAP16F,nuf,LanB1,sn,ct,Ten-m,Cip4,Mmp2,pnr,ftz-f1,trol,dap,ena,mbc,Fkbp14,Grasp65,Chc,kuz,Rab11,Acer,aop,Arf1,Ggamma1,Past1,if,Nhe2,sqh,Slik,msn,Cdc42,Frl,shi,shn,Doa,sgg,Stat92E,Arf6,deltaCOP,ATP6AP2,GEFmeso,px,Septin1* |
| biological process involved in interspecies interaction between organisms | GO:BP | 1.21e-05 | *Glt,Npc2a,St3,Pvr,Sr-CI,crq,RhoL,MP1,spz,LanA,senju,LanB1,Spn27A,Fer1HCH,Fer2LCH,cathD,TM9SF2,Mmp2,sau,pnr,Trx-2,Sar1,ush,Rab11,Arf1,drpr,Rab1,CBP,GILT2,Sec61alpha,Rab8,Tmep,Cdc42,shi,Rab7,NUCB1,Doa,Stat92E,Vps29,Vps60* |
| Golgi organization | GO:BP | 1.35e-05 | *sau,bai,Sec24CD,Grasp65,CHOp24,ergic53,Surf4,eca,p24-1,Sec23,Bet1,Sec22,loj,cnn* |
| glycosylation | GO:BP | 1.64e-05 | *GlcAT-P,Pgant9,C1GalTA,senju,CG9171,Gfat2,CG32276,wol,CG8668,CG33303,Ost48,OstDelta,Ostgamma,Alg11,kud,Stt3B,Dad1* |
| viral process | GO:BP | 2.24e-05 | *sau,Sar1,Rab11,Rab1,Sec61alpha,Rab8,shi,Rab7,Vps29,Vps60* |
| signal peptide processing | GO:BP | 8.15e-05 | *Spp,twr,Spase22-23,Spase12,Spase25* |
| chitin-based larval cuticle pattern formation | GO:BP | 3.23e-04 | *CrebA,Sar1,Sec13,SrpRbeta,Sec23,sgg* |
| substrate adhesion-dependent cell spreading | GO:BP | 3.57e-04 | *Tig,LanB2,LanA,LanB1,if* |
| positive regulation of cuticle pigmentation | GO:BP | 5.41e-04 | *Sec24CD,Sar1,Sec23* |
| basement membrane assembly | GO:BP | 5.41e-04 | *SPARC,Pxn,LanB2,LanB1* |
| SRP-dependent cotranslational protein targeting to membrane, translocation | GO:BP | 9.67e-04 | *TRAM,Sec61beta,CG8860,Sec61alpha* |
| dorsal closure | GO:BP | 0.001 | *Col4a1,Pvr,srp,ena,mbc,Rab11,aop,Ggamma1,Sec61alpha,msn,Cdc42,shn* |
| formation of a compartment boundary | GO:BP | 0.002 | *ct,ena,sqh* |
| positive regulation of apoptotic cell clearance | GO:BP | 0.002 | *srp,prtp,CaBP1* |
| positive regulation of border follicle cell migration | GO:BP | 0.002 | *Pvr,Catsup,mbc,kuz,aop,sqh,slbo* |
| lipid catabolic process | GO:BP | 0.003 | *CG5191,CG3376,bwa,Hexo2,CG5112,CG8839,Hnf4,yip2,fdl,CG7461,CG4598,Lsd-2,sws,Mtpbeta,whd* |
| anterior Malpighian tubule development | GO:BP | 0.004 | *SPARC,vkg,Pvr,mbc* |
| hemocyte migration | GO:BP | 0.005 | *Pvr,RhoL,sn,if,Cdc42* |
| organelle fusion | GO:BP | 0.006 | *Gs1,Snx6,Vamp7,Syb,Rab11,Rab8,Bet1,Sec22,Rab7,Syx1A* |
| muscle cell development | GO:BP | 0.006 | *Col4a1,sty,pnt,C1GalTA,LanB1,Bsg,aop,if* |
| imaginal disc fusion, thorax closure | GO:BP | 0.006 | *Arpc3B,Pvr,Mmp2,pnr,mbc,msn* |
| negative regulation of fibroblast growth factor receptor signaling pathway | GO:BP | 0.006 | *sty,Mmp2,trol* |
| membrane fusion | GO:BP | 0.006 | *Snx6,Vamp7,Syb,Rab11,Rab8,Bet1,Sec22,Rab7,Syx1A* |
| retrograde vesicle-mediated transport, Golgi to endoplasmic reticulum | GO:BP | 0.006 | *sau,zetaCOP,epsilonCOP,Sec22,deltaCOP* |
| negative regulation of neurotransmitter secretion | GO:BP | 0.009 | *cpx,sky* |
| iron ion import across plasma membrane | GO:BP | 0.009 | *Fer1HCH,Fer2LCH* |
| positive regulation of extracellular matrix constituent secretion | GO:BP | 0.009 | *Sec24CD,Sec23* |
| detoxification of iron ion | GO:BP | 0.009 | *Fer1HCH,Fer2LCH* |
| post-translational protein targeting to membrane, translocation | GO:BP | 0.009 | *Sec61beta,CG8860,Sec61alpha* |
| positive regulation of nuclear-transcribed mRNA catabolic process, deadenylation-dependent decay | GO:BP | 0.009 | *Tis11,pum* |
| protein processing | GO:BP | 0.01 | *Nepl19,sel,Mmp2,cer,Spp,twr,Spase22-23,Spase12,Spase25,CG8320* |
| endoplasmic reticulum calcium ion homeostasis | GO:BP | 0.012 | *BI-1,CG1840,SelG* |
| protein N-linked glycosylation via asparagine | GO:BP | 0.013 | *CG33303,Ost48,Ostgamma,Stt3B* |
| cell surface receptor signaling pathway | GO:BP | 0.017 | *Col4a1,Ilp4,NtR,nAChRbeta3,sgl,Pvr,sty,ham,crq,pnt,spz,Ilp6,senju,Akap200,Mmp2,trol,mbc,Fkbp14,pigs,pum,CycG,kuz,ush,aop,ken,if,sqh,AdipoR,msn,Sema1b,Cdc42,Dyrk2,shn,Doa,sgg,Stat92E,ATP6AP2,GEFmeso* |
| endoplasmic reticulum unfolded protein response | GO:BP | 0.018 | *CG32276,BI-1,Xbp1,Der-1* |
| protein K69-linked ufmylation | GO:BP | 0.02 | *Ufm1,Ufc1* |
| ganglioside catabolic process | GO:BP | 0.02 | *Hexo2,fdl* |
| intracellular sequestering of iron ion | GO:BP | 0.02 | *Fer1HCH,Fer2LCH* |
| pericardial nephrocyte differentiation | GO:BP | 0.021 | *pnt,pnr,kuz* |
| galactose metabolic process | GO:BP | 0.021 | *Gale,Gal,Galk* |
| negative regulation of catalytic activity | GO:BP | 0.022 | *sty,Spn27A,dap,cer,BI-1,sws,sgg* |
| retrograde transport, endosome to Golgi | GO:BP | 0.024 | *Snx6,Snx3,CG2747,Vps29,CG13784* |
| male meiosis cytokinesis | GO:BP | 0.024 | *sau,Rab11,Arf1,Rab1,Arf6* |
| midgut development | GO:BP | 0.024 | *LanB2,srp,if,shn,cnn* |
| mesoderm migration involved in gastrulation | GO:BP | 0.026 | *sgl,fog,wol* |
| negative regulation of stem cell differentiation | GO:BP | 0.026 | *pnt,C1GalTA,Stat92E* |
| intracellular protein transmembrane transport | GO:BP | 0.031 | *TRAM,Sec61beta,CG8860,Sec61alpha* |
| wing disc dorsal/ventral pattern formation | GO:BP | 0.032 | *pnt,nuf,drpr,shn,Rab7,CG14073* |
| positive regulation of dendrite morphogenesis | GO:BP | 0.032 | *Sar1,Rab1,Sec23* |
| engulfment of apoptotic cell | GO:BP | 0.032 | *NimC4,crq,drpr* |
| fatty acid beta-oxidation | GO:BP | 0.032 | *yip2,CG7461,CG4598,Mtpbeta,whd* |
| L-proline biosynthetic process | GO:BP | 0.033 | *P5CS,P5cr-2* |
| positive regulation of synapse pruning | GO:BP | 0.033 | *Sar1,Rab1* |
| negative regulation of viral entry into host cell | GO:BP | 0.033 | *Sar1,Rab1* |
| regulation of COPII vesicle coating | GO:BP | 0.033 | *Sar1,Sec23* |
| COPII-coated vesicle cargo loading | GO:BP | 0.033 | *Sec24CD,Sec23* |
| subsynaptic reticulum organization | GO:BP | 0.033 | *ena,Past1* |
| cardiac muscle cell development | GO:BP | 0.033 | *Col4a1,LanB1* |
| gastrulation | GO:BP | 0.034 | *sgl,srp,fog,Dtg,wol,Ggamma1,sqh,SelG* |
| trans-synaptic signaling | GO:BP | 0.034 | *CG8501,NtR,nAChRbeta3,CG9338,nemy,Manf,Ten-m,cpx,Syb,pum,Chc,Scamp,Arf1,Sod1,Cdc42,shi,Syx1A,Arf6,sky* |
| negative regulation of protein kinase activity | GO:BP | 0.034 | *sty,dap,sws,sgg* |
| negative regulation of apoptotic signaling pathway | GO:BP | 0.034 | *pnt,Men,BI-1,CG2918* |
| positive regulation of establishment of protein localization | GO:BP | 0.038 | *Sar1,Rab11,Snx3,Rab1,AdipoR* |
| intracellular iron ion homeostasis | GO:BP | 0.044 | *Fer1HCH,Fer2LCH,Cisd2* |
| positive regulation of nurse cell apoptotic process | GO:BP | 0.046 | *Atg13,E2f1* |
| juvenile hormone mediated signaling pathway | GO:BP | 0.046 | *ftz-f1,Chd64* |
| protein ufmylation | GO:BP | 0.046 | *Ufm1,Ufc1* |
| response to tumor cell | GO:BP | 0.046 | *Pvr,spz* |
| maternal specification of dorsal/ventral axis, oocyte, germ-line encoded | GO:BP | 0.046 | *sel,bai* |
| cortical actin cytoskeleton organization | GO:BP | 0.048 | *sau,Rab11,Slik,Cdc42,Frl,shi* |
| cargo receptor activity | GO:MF | 0.004 | *Sr-CI,crq,LpR2,bai,CHOp24,eca,p24-1,drpr,loj* |
| protein-containing complex binding | GO:MF | 0.017 | *SPARC,Tig,Arpc3B,Gel,LpR2,sn,Akap200,bif,pigs,pum,Rab11,Trf2,CG6891,Rab1,if,zip,Sec61alpha,Alh,SrpRbeta,HSPC300,Frl,Rab7,SrpRalpha,Der-1,Chd64* |
| signal recognition particle binding | GO:MF | 0.017 | *SrpRbeta,SrpRalpha,Der-1* |
| lipid binding | GO:MF | 0.017 | *Npc2a,CG5958,Npc2g,NimC4,CG5973,CG3246,CG9338,Gel,Cip4,sau,Snx6,CG7461,Snx3,drpr,AnxB11,Orp8,Clic,sky,CG3262* |
| actin filament binding | GO:MF | 0.025 | *Arpc3B,Gel,sn,Akap200,bif,pigs,CG6891,zip,Frl,Chd64* |
| fatty acid amide hydrolase activity | GO:MF | 0.025 | *CG5191,CG5112,CG8839* |
| GTPase activity | GO:MF | 0.034 | *RhoL,Sar1,Rab11,Arf1,Rab1,Rab8,SrpRbeta,Cdc42,shi,Rab7,SrpRalpha,Arf6,Septin1* |
| protein disulfide isomerase activity | GO:MF | 0.035 | *prtp,ERp60,CaBP1,Pdi* |
| SNAP receptor activity | GO:MF | 0.037 | *Vamp7,Syb,Bet1,Sec22,Syx1A* |
| procollagen galactosyltransferase activity | GO:MF | 0.046 | *Plod,CG31915* |
| guanyl nucleotide binding | GO:MF | 0.049 | *RhoL,Sar1,Rab11,Arf1,Past1,Rab1,Rab8,SrpRbeta,Cdc42,shi,Rab7,SrpRalpha,Arf6,Septin1* |
| GTP binding | GO:MF | 0.049 | *RhoL,Sar1,Rab11,Arf1,Past1,Rab1,Rab8,SrpRbeta,Cdc42,shi,Rab7,SrpRalpha,Arf6,Septin1* |
| GTPase activating protein binding | GO:MF | 0.049 | *Cip4,Rab11,Cdc42* |
| guanyl ribonucleotide binding | GO:MF | 0.049 | *RhoL,Sar1,Rab11,Arf1,Past1,Rab1,Rab8,SrpRbeta,Cdc42,shi,Rab7,SrpRalpha,Arf6,Septin1* |
| signaling receptor binding | GO:MF | 0.049 | *Nplp2,Idgf6,Ilp4,Tig,spz,LanA,fog,Ilp6,Akap200,wake,miple2,CG8507,ITP,CG30423,if,Sema1b,Stat92E,ATP6AP2* |
| actin binding | GO:MF | 0.049 | *Arpc3B,Gel,sn,Akap200,bif,ena,pigs,CG6891,zip,Frl,shi,Chd64* |
| cysteine-type endopeptidase inhibitor activity | GO:MF | 0.049 | *Cys,CG8066,cer* |

### 1.18. Gene Ontology Enrichment Analysis: longitudinal glia - naz

Significant GO terms (p < 0.05) with associated genes

| GO Term | Source | P-value | Genes |
| --- | --- | --- | --- |
| endoplasmic reticulum to Golgi vesicle-mediated transport | GO:BP | 2.97e-10 | *boca,opm,KdelR,CG5510,Bet1,Sar1,eca,cni,bai,CHOp24,p24-1,CG32069,epsilonCOP,CG13887,Rab1,Sec13,CG32576,Arl1,Sec23,Tango1,Sec16,CG3652,deltaCOP* |
| protein localization to endoplasmic reticulum | GO:BP | 1.20e-09 | *KdelR,Sec61gamma,CG11857,Spase25,CG5885,Spase12,Spase22-23,Sec61beta,Sec61alpha,CG13887,Sec63,CG8860,SrpRbeta,Sgt,Sec16,Srp9* |
| animal organ morphogenesis | GO:BP | 5.58e-07 | *Dll,zfh2,Dr,fng,rau,fog,ct,pnt,sty,bnl,pros,Mkp3,EMC3,Rbfox1,Pcyt1,rho,scrib,spi,Abl,Fas2,Atpalpha,aos,CG43658,Rab11,betaTub97EF,par-1,puc,Ras85D,Dad,Frl,caps,GEFmeso,Rac2,Spn88Ea,mav,Rala,Rap1,ATP6AP2,Arf6,Gbeta13F,dos,sli,Rab5,Arf1,14-3-3epsilon,Pura,fra,hth,Hs6st,Ggamma1,Nrg,ctp,sina,Exn,EloC,myo,p120ctn,Chc,Rab6,jagn,Grasp65,Fkbp14,Cbl,Rac1,alphaSnap,svr,gish,Atx2,nrv2,Hipk,Vps25,stck,Ccm3,RasGAP1,fz2,deltaCOP,CSN8,heph* |
| Golgi organization | GO:BP | 8.72e-07 | *Tango5,opm,CG5510,Bet1,eca,bai,CHOp24,p24-1,Surf4,Grasp65,Arl1,Vti1b,Snap29,Pld,Sec23,Ccm3,Tango1,Sec16,Pkc98E,fwd* |
| tail-anchored membrane protein insertion into ER membrane | GO:BP | 1.56e-06 | *EMC1,EMC7,EMC5,EMC6,EMC3,EMC8-9,EMC4* |
| signal release | GO:BP | 7.02e-06 | *opm,sky,Snx3,veli,Rap1,eca,Arf6,bai,CHOp24,Arl8,ric8a,p24-1,Arf1,Rab1,Exn,Chc,AdipoR,Syx1A,Scamp,Cadps,alphaSnap,Arl1,Syb,Snap29* |
| fatty acid beta-oxidation | GO:BP | 8.04e-06 | *yip2,CG4598,Mtpalpha,Arc42,Ech1,Etfb,Mtpbeta,CG7461,Mcad,wal,whd,Echs1* |
| cell surface receptor signaling pathway | GO:BP | 2.78e-05 | *stumps,fng,rau,pnt,Prosap,Buffy,sty,Trim9,htl,Sema2b,bnl,Mkp3,boca,wgn,Eaat1,pum,shv,rho,spi,Fas2,aos,for,par-1,puc,pen-2,Ras85D,Dad,GEFmeso,Rac2,lili,mav,Rala,Rap1,ATP6AP2,Gbeta13F,dos,sli,edl,ttv,Rheb,fra,pigs,sina,kek1,EloC,myo,AdipoR,par-6,lqf,Fkbp14,Mob4,Cbl,Rac1,sgl,gish,ben,Sik3,Hipk,Usp10,CkIIbeta,Nedd8,Sema1a,stck,pbl,Pp1alpha-96A,RasGAP1,Pp1-87B,Tsp86D,fz2,Ubr3,Ack-like,DENR* |
| catabolic process | GO:BP | 3.09e-05 | *GLaz,Ace,Gs2,Pde9,Ssadh,Gabat,Rchy1,Buffy,Trim9,Sod1,yip2,EMC6,Ntan1,Hnf4,AP-2alpha,CG18135,CG4598,Sting,Tango5,Mtpalpha,Arc42,pum,CRMP,Ech1,Pglym78,Shmt,aus,Ald1,CG15111,CG1640,Sec61gamma,Etfb,Der-1,Ras85D,Lamp1,olf413,Apt1,CtsB,Tpi,Pgk,Oscillin,Mtpbeta,ifc,Ubc7,CG6567,Arl8,Xbp1,Sec61beta,Gapdh2,CG6766,PhKgamma,Svip,Rheb,CG12384,GlyS,BI-1,Sec61alpha,CG7461,Mcad,ctp,sina,Eno,Rab1,EloC,Ubqn,Chc,lqf,Desat1,sordd1,wal,Cbl,Lsd-2,Sik3,Klp98A,Pgi,CG6878,CG1440,Usp10,Vti1b,Nedd8,Vps25,HINT1,zda,Snap29,Prosbeta5,Pld,Pde8,Sec16,whd,CG5676,PI31,Atg1,Vha100-1,Echs1,Atg3,EndoA,Prosbeta6,Ubr3,Rpt4* |
| proton transmembrane transport | GO:BP | 6.35e-05 | *VhaM9.7-a,Vha14-1,Atpalpha,Vha13,Vha26,Vha16-1,Vha68-2,VhaM9.7-c,ATP6AP2,VhaAC39-1,VhaPPA1-1,VhaM9.7-b,Vha44,Vha36-1,Vha55,VhaSFD,Vha100-1* |
| trans-synaptic signaling | GO:BP | 8.35e-05 | *Ace,Sod1,Ten-a,CG5541,Csas,wgn,Eaat1,FMRFaR,pum,sky,Manf,Abl,Fas2,Atpalpha,bou,CG12290,veli,be,Pgk,Ent2,Rap1,Arf6,Arl8,ric8a,Arf1,stas,Acsl,Exn,Chc,Syx1A,Scamp,Cadps,alphaSnap,Arl1,Syb,Snap29,lap,Atg1* |
| post-translational protein targeting to membrane, translocation | GO:BP | 1.57e-04 | *Sec61gamma,Sec61beta,Sec61alpha,Sec63,CG8860* |
| terminal branching, open tracheal system | GO:BP | 1.68e-04 | *bnl,Vha13,Vha26,Ras85D,VhaAC39-1,Rheb,VhaPPA1-1,par-6,Vha100-1* |
| organic acid metabolic process | GO:BP | 2.06e-04 | *Gs2,Ssadh,Gabat,Baldspot,Acbp2,yip2,CG4598,e,Mtpalpha,Arc42,Ech1,Pglym78,Shmt,Ald1,CG7920,CG1640,Etfb,Mdh1,Tpi,Pgk,Oscillin,Mtpbeta,PH4alphaEFB,Gapdh2,Hacd2,Acsl,scu,Sc2,CG7461,Mcad,Eno,Got1,AdipoR,Desat1,wal,sgl,CG1764,Pgi,CG1440,whd,Acbp1,Echs1* |
| fatty acid beta-oxidation using acyl-CoA dehydrogenase | GO:BP | 3.47e-04 | *Arc42,Etfb,CG7461,Mcad,wal* |
| long-term memory | GO:BP | 3.61e-04 | *fabp,orb2,CG5541,pum,for,vsg,be,ATP6AP2,CG4612,Scamp,Tob,svr,Atx2,Pkc98E* |
| motor neuron axon guidance | GO:BP | 7.90e-04 | *Ten-a,fend,Abl,Fas2,CG42327,for,caps,Rac2,Sdc,fra,Nrg,Rac1,Sema1a,fz2* |
| dorsal closure | GO:BP | 0.001 | *rho,scrib,spi,Abl,Rab11,puc,Ras85D,Rac2,Rap1,Rab5,Ggamma1,Sec61alpha,Rac1,stck,Inx3,Ack-like* |
| mesoderm migration involved in gastrulation | GO:BP | 0.002 | *fog,htl,sli,sgl,pbl* |
| negative regulation of epidermal growth factor-activated receptor activity | GO:BP | 0.002 | *aos,kek1,Cbl* |
| ommatidial rotation | GO:BP | 0.002 | *pnt,sty,spi,Abl,aos,Ras85D,Frl,Rap1* |
| nucleotide catabolic process | GO:BP | 0.002 | *Pde9,Pglym78,Ald1,Tpi,Pgk,Gapdh2,Eno,Pgi,HINT1,Pde8* |
| membrane fusion | GO:BP | 0.002 | *aus,Rab11,Snx6,Bet1,Arl8,Rab5,Svip,Syx1A,alphaSnap,Syb,Vti1b,Snap29,Pld* |
| gastrulation | GO:BP | 0.002 | *stumps,fog,htl,spi,Abl,Ras85D,Gbeta13F,sli,ric8a,Ggamma1,Rab35,sgl,Pld,pbl* |
| apical protein localization | GO:BP | 0.003 | *Vha26,Gbeta13F,Rheb,Ggamma1,Rab35,par-6* |
| positive regulation of filopodium assembly | GO:BP | 0.003 | *Rab35,par-6,Rac1,kug,HSPC300,cpa,Arp3* |
| dsRNA transport | GO:BP | 0.003 | *AP-2mu,Vha16-1,CG8671,Chc,Arl1,VhaSFD,egh* |
| endoplasmic reticulum unfolded protein response | GO:BP | 0.004 | *Der-1,Hsc70-3,Xbp1,CG6766,BI-1,Atf6* |
| viral process | GO:BP | 0.004 | *Rab11,Sar1,Arf4,Rab5,Vps29,Sec61alpha,Rab1,EloC,Rab35* |
| SRP-dependent cotranslational protein targeting to membrane, translocation | GO:BP | 0.005 | *Sec61gamma,Sec61beta,Sec61alpha,CG8860* |
| regulation of COPII vesicle coating | GO:BP | 0.005 | *Sar1,Sec23,Sec16* |
| peripheral nervous system development | GO:BP | 0.006 | *repo,ct,Trim9,pros,spi,Ras85D,fra,hth,Rac1,egh* |
| glycolytic process | GO:BP | 0.007 | *Pglym78,Ald1,Tpi,Pgk,Gapdh2,Eno,Pgi* |
| positive regulation of lamellipodium assembly | GO:BP | 0.007 | *Abl,par-6,HSPC300,Arp3* |
| signal peptide processing | GO:BP | 0.007 | *Spase25,twr,Spase12,Spase22-23* |
| positive regulation of establishment of protein localization | GO:BP | 0.008 | *Snx3,Abl,Rab11,Sar1,Rab1,Rab35,AdipoR,Sec16* |
| nucleoside diphosphate catabolic process | GO:BP | 0.009 | *Pglym78,Ald1,Tpi,Pgk,Gapdh2,Eno,Pgi* |
| organelle fusion | GO:BP | 0.01 | *aus,Rab11,Snx6,Bet1,Arl8,Rab5,Svip,Syx1A,alphaSnap,Syb,Vti1b,Snap29,Pld* |
| secretory granule organization | GO:BP | 0.011 | *Chc,Arl1,Tango1,AP-1gamma* |
| dorsal closure, amnioserosa morphology change | GO:BP | 0.011 | *Rab11,Rab5,Rac1,Ack-like* |
| protein stabilization | GO:BP | 0.012 | *orb2,Cnx99A,Snx3,ric8a,Tob,Hipk,CG7945,CSN8* |
| protein N-linked glycosylation via asparagine | GO:BP | 0.012 | *Ostgamma,Dpm1,Ost48,CG33303,Stt3B* |
| behavioral response to ethanol | GO:BP | 0.014 | *pum,rho,Shmt,spi,Fas2,vsg,Gbs-70E,Spn27A,Arf6* |
| lipid catabolic process | GO:BP | 0.014 | *GLaz,yip2,Hnf4,CG18135,CG4598,Mtpalpha,Arc42,Ech1,CG15111,Etfb,Mtpbeta,CG7461,Mcad,wal,Lsd-2,Sik3,Pld,whd,Echs1* |
| retrograde transport, endosome to Golgi | GO:BP | 0.014 | *Snx3,Snx1,Snx6,Vps29,Rab6,CG32576,Vti1b* |
| regulation of tube diameter, open tracheal system | GO:BP | 0.015 | *Fas2,Atpalpha,nrv2,deltaCOP* |
| negative regulation of stem cell differentiation | GO:BP | 0.015 | *pnt,C1GalTA,aos,lqf* |
| axon midline choice point recognition | GO:BP | 0.016 | *Trim9,Abl,sli,fra,Rac1,Sema1a* |
| oocyte microtubule cytoskeleton polarization | GO:BP | 0.019 | *Rab11,14-3-3epsilon,Chc,Rab6,heph* |
| chitin-based larval cuticle pattern formation | GO:BP | 0.019 | *Sar1,Sec13,SrpRbeta,Sec23,fz2* |
| mesectoderm development | GO:BP | 0.019 | *fog,Gbeta13F,Ggamma1* |
| intracellular monoatomic ion homeostasis | GO:BP | 0.019 | *zyd,CG11367,CG17593,CG10470,emei,Atpalpha,Vha16-1,Vha68-2,VhaAC45,VhaAC39-1,Vha55,BI-1,nrv2,CG1840,Vha100-1* |
| photoreceptor cell fate determination | GO:BP | 0.019 | *rho,spi,Ras85D* |
| stem cell fate commitment | GO:BP | 0.019 | *rho,spi,Ras85D* |
| negative regulation of lipophagy | GO:BP | 0.019 | *ifc,Rheb* |
| negative regulation of epithelial cell proliferation | GO:BP | 0.019 | *scrib,Vps25* |
| negative regulation of myotube cell migration | GO:BP | 0.019 | *Rac1,pbl* |
| peripheral nervous system neuron axonogenesis | GO:BP | 0.019 | *Trim9,fra* |
| 5-phosphoribose 1-diphosphate biosynthetic process | GO:BP | 0.019 | *Prps,CG2246* |
| cell adhesion involved in heart morphogenesis | GO:BP | 0.019 | *Gbeta13F,Ggamma1,Nrg,nrv2* |
| gamma-aminobutyric acid catabolic process | GO:BP | 0.019 | *Ssadh,Gabat* |
| protein localization to endoplasmic reticulum exit site | GO:BP | 0.019 | *CG13887,Sec16* |
| female courtship behavior | GO:BP | 0.019 | *shep,Nrg* |
| head involution | GO:BP | 0.02 | *pum,Rac2,Sec61alpha,wal,Rac1,stck,Ack-like* |
| glucose homeostasis | GO:BP | 0.02 | *Hnf4,Ald1,Tpi,Gapdh2,Eno,AdipoR,Desat1,Pgi* |
| glycosylation | GO:BP | 0.024 | *fng,NANS,Ostgamma,C1GalTA,Dpm1,kud,Dad1,CG33774,OstDelta,Ost48,CG33303,Dpm3,Stt3B,ttv,Alg11* |
| vacuolar acidification | GO:BP | 0.025 | *Vha16-1,VhaAC39-1,Vha55,Vha100-1* |
| clathrin-dependent synaptic vesicle endocytosis | GO:BP | 0.028 | *fwe,lap,EndoA* |
| autophagosome-lysosome fusion | GO:BP | 0.028 | *aus,Arl8,Svip* |
| regulation of myosin II filament organization | GO:BP | 0.028 | *fog,Gbeta13F,Ggamma1* |
| lymph gland plasmatocyte differentiation | GO:BP | 0.028 | *pnt,htl,Ras85D* |
| regulation of adherens junction organization | GO:BP | 0.028 | *Abl,Rac2,Rac1* |
| peptidyl-tyrosine phosphorylation | GO:BP | 0.031 | *Abl,aos,kek1,Cbl,Ack-like* |
| positive regulation of epidermal growth factor receptor signaling pathway | GO:BP | 0.031 | *rau,GEFmeso,Rala,edl* |
| sulfur compound metabolic process | GO:BP | 0.031 | *CG5065,GstE11,GstD1,GstE1,Chpf,GstE2,GstD3,Dpck,GstS1,Hmgs,ttv,Hs6st,Acsl,scu,sgl,CG1440,ScsbetaA,GstE12* |
| cellular response to growth factor stimulus | GO:BP | 0.032 | *stumps,rau,pnt,sty,htl,bnl,Ras85D,Dad,lili,mav,lqf,sgl,pbl,RasGAP1* |
| negative regulation of catalytic activity | GO:BP | 0.034 | *sty,aos,Spn27A,BI-1,kek1,endos,par-6,Cbl,conu* |
| negative regulation of signal transduction | GO:BP | 0.034 | *fng,pnt,Prosap,sty,Mkp3,pum,Fas2,CG2918,aos,par-1,puc,Ras85D,Dad,Rala,Rap1,Gbeta13F,edl,Rheb,BI-1,pigs,kek1,EloC,Mob4,Cbl,Sik3,Hipk,Nedd8,stck,pbl,raskol,Pp1alpha-96A,RasGAP1,Atg1,cpa* |
| melanotic encapsulation of foreign target | GO:BP | 0.036 | *Rac2,Spn27A,Nrg,Exn,Eb1,Rac1* |
| retrograde vesicle-mediated transport, Golgi to endoplasmic reticulum | GO:BP | 0.036 | *CG11857,epsilonCOP,Rab6,zetaCOP,deltaCOP* |
| regulation of gastrulation | GO:BP | 0.038 | *Gbeta13F,Ggamma1,Rab35* |
| primary branching, open tracheal system | GO:BP | 0.038 | *pnt,bnl,Mkp3* |
| dorsal closure, elongation of leading edge cells | GO:BP | 0.038 | *Rab11,Rac2,Rac1* |
| regulation of lipid transport | GO:BP | 0.038 | *LRP1,Npc1a,lilli* |
| ectodermal digestive tract morphogenesis | GO:BP | 0.038 | *puc,wal,Rac1* |
| vacuolar proton-transporting V-type ATPase complex assembly | GO:BP | 0.038 | *ATP6AP2,VhaAC45,CG5969* |
| L-leucine import across plasma membrane | GO:BP | 0.043 | *mnd,CD98hc* |
| positive regulation of integrin-mediated signaling pathway | GO:BP | 0.043 | *shv,stck* |
| positive regulation of cuticle pigmentation | GO:BP | 0.043 | *Sar1,Sec23* |
| glucose catabolic process | GO:BP | 0.043 | *PhKgamma,Eno* |
| mitotic cleavage furrow ingression | GO:BP | 0.043 | *AP-2alpha,Arf1* |
| positive regulation of sequestering of triglyceride | GO:BP | 0.043 | *Acsl,Lsd-2* |
| immune response-regulating cell surface receptor signaling pathway involved in phagocytosis | GO:BP | 0.043 | *Rac2,Rac1* |
| myoblast fate specification | GO:BP | 0.043 | *htl,Ras85D* |
| positive regulation of protein exit from endoplasmic reticulum | GO:BP | 0.043 | *Sar1,Sec16* |
| secondary branching, open tracheal system | GO:BP | 0.043 | *pnt,bnl* |
| protein retention in ER lumen | GO:BP | 0.043 | *KdelR,CG11857* |
| leg disc proximal/distal pattern formation | GO:BP | 0.045 | *Dll,pnt,rho,Ras85D* |
| proton-transporting ATPase activity, rotational mechanism | GO:MF | 5.23e-09 | *VhaM9.7-a,Vha14-1,Vha13,Vha26,Vha16-1,Vha68-2,VhaM9.7-c,VhaAC39-1,VhaPPA1-1,VhaM9.7-b,Vha44,Vha36-1,Vha55,VhaSFD,Vha100-1* |
| ATPase-coupled monoatomic cation transmembrane transporter activity | GO:MF | 1.86e-08 | *VhaM9.7-a,Vha14-1,Atpalpha,Vha13,Vha26,Vha16-1,Vha68-2,VhaM9.7-c,VhaAC39-1,VhaPPA1-1,VhaM9.7-b,Vha44,Vha36-1,Vha55,VhaSFD,Vha100-1* |
| membrane insertase activity | GO:MF | 4.31e-06 | *EMC1,EMC7,EMC5,EMC6,EMC3,EMC8-9,EMC4* |
| ATPase-coupled transmembrane transporter activity | GO:MF | 1.63e-04 | *CG32091,VhaM9.7-a,Vha14-1,Atpalpha,Vha13,Vha26,Vha16-1,Hsc70-3,Vha68-2,VhaM9.7-c,VhaAC39-1,VhaPPA1-1,VhaM9.7-b,Vha44,Vha36-1,Vha55,VhaSFD,Vha100-1* |
| GTPase activity | GO:MF | 0.001 | *EMC10,Rab11,betaTub97EF,Ras85D,Rac2,Sar1,Rala,Rap1,Arf6,Arl8,Arf4,Rab5,Arf1,Rheb,Rab1,Arl4,Rab35,Rab6,Rac1,Arl1,SrpRbeta* |
| oxidoreductase activity, acting on a sulfur group of donors | GO:MF | 0.002 | *CG11007,Grx1,SelT,GILT2,CG9302,CG7484,ERp60,CaBP1,CG5554,CG8993,GILT3* |
| guanyl nucleotide binding | GO:MF | 0.002 | *Sting,EMC10,Rab11,for,betaTub97EF,Ras85D,Rac2,Sar1,Rala,Rap1,Arf6,Arl8,Arf4,Rab5,Arf1,Rheb,Rab1,Arl4,Rab35,Rab6,Rac1,Arl1,SrpRbeta,alphaTub84D* |
| guanyl ribonucleotide binding | GO:MF | 0.002 | *Sting,EMC10,Rab11,for,betaTub97EF,Ras85D,Rac2,Sar1,Rala,Rap1,Arf6,Arl8,Arf4,Rab5,Arf1,Rheb,Rab1,Arl4,Rab35,Rab6,Rac1,Arl1,SrpRbeta,alphaTub84D* |
| disulfide oxidoreductase activity | GO:MF | 0.003 | *CG11007,Grx1,SelT,CG9302,CG7484,ERp60,CaBP1,CG5554,CG8993* |
| inorganic cation transmembrane transporter activity | GO:MF | 0.003 | *Gat,zyd,CG5888,rumpel,pain,EMC5,bumpel,CG10470,ine,Eaat1,VhaM9.7-a,Irk3,Vha14-1,Atpalpha,CG10413,Vha13,Vha26,Vha16-1,Vha68-2,VhaM9.7-c,fwe,Nmda1,VhaAC39-1,VhaPPA1-1,VhaM9.7-b,Vha44,Vha36-1,Vha55,BI-1,VhaSFD,Zip102B,Vha100-1* |
| protein binding | GO:MF | 0.003 | *alrm,GLaz,stumps,mgl,CG5888,rau,fog,Fas1,Spn,pnt,EMC2A,Prosap,Buffy,Trim9,htl,Sema2b,pain,EMC1,ced-6,Sod1,CG9467,bnl,EMC2B,orb2,NimA,sel,Cnx99A,Plp,GCS2beta,CG18135,Ten-a,wrapper,boca,CG5541,RabGGTb,TBCB,Hmu,wgn,Calr,Rbfox1,opm,wake,AnxB9,EMC8-9,C1GalTA,shv,scrib,Shmt,spi,CG14441,CG7484,Clc,Liprin-gamma,mtgo,Trf2,Abl,CG8507,Gp93,CLIP-190,Fas2,CG2918,Wdfy2,LRP1,Lapsyn,zormin,aos,kud,Rab11,Etfb,form3,CG11999,CG3061,Der-1,ITP,par-1,puc,mfas,Vha13,mei-P26,Vha26,Ras85D,Dad,Frl,caps,P58IPK,GEFmeso,Meltrin,Gbs-70E,corn,Hsc70-3,jvl,Rac2,veli,loco,Chd64,Tpi,Oscillin,Bet1,CG10353,fwe,mav,Sar1,MSBP,Rala,Rap1,NKAIN,EndoB,ATP6AP2,cni,Gbeta13F,dos,Act5C,pr,sli,CG42346,Arl8,PH4alphaEFB,ric8a,Rab5,edl,PhKgamma,hts,ttv,Rheb,Hacd2,Unc-115a,CG12384,14-3-3epsilon,Taf10b,fra,epsilonCOP,hth,spt4,Ggamma1,SLIRP2,TfIIA-S,drpr,CG32264,BI-1,Nrg,pigs,Sc2,ctp,sina,kek1,CG11267,Rab1,Exn,EloC,myo,Ubqn,Sec13,p120ctn,Chc,Arpc5,par-6,Syx1A,Rab6,Eb1,Grasp65,Sh3beta,lqf,Tob,Desat1,sordd1,wal,robl,Cbl,Rac1,E2f1,Taf3,alphaSnap,conu,Lsd-2,Atx2,ben,Arl1,Syb,Klp98A,kug,CG6891,Pfdn4,CAP,Sgt,Maf1,Hipk,gce,Vti1b,Nedd8,Vps25,CG40228,Sema1a,cib,zda,HSPC300,Snap29,Prosbeta5,stck,lbk,pbl,fax,Ccm3,Lamtor3,alphaTub84D,Tango1,lap,CG7668,Sec16,Alg-2,ubl,Pp1alpha-96A,CG9344,CG30423,PI31,Pp1-87B,Atg1,Vha100-1,HnRNP-K,CG7945,cpa,fz2,CG2765,EndoA,lilli,Arpc4,Ubr3,Arp3,Ack-like,Rpt4,tei* |
| proton transmembrane transporter activity | GO:MF | 0.004 | *VhaM9.7-a,Vha14-1,Vha13,Vha26,Vha16-1,Vha68-2,VhaM9.7-c,VhaAC39-1,VhaPPA1-1,VhaM9.7-b,Vha44,Vha36-1,Vha55,VhaSFD,Vha100-1* |
| GDP binding | GO:MF | 0.005 | *Ras85D,Rala,Rap1,Arl8,Rheb* |
| lipid binding | GO:MF | 0.005 | *fabp,CG3246,Acbp2,AP-2mu,sky,AnxB9,Pcyt1,Snx3,Snx1,Snx6,Snx17,MSBP,Npc1a,EndoB,Npc2a,drpr,CG7461,lqf,Klp98A,AnxB11,gce,CG9205,Pld,pbl,lap,Acbp1,EndoA* |
| protein transmembrane transporter activity | GO:MF | 0.005 | *Sec61gamma,Hsc70-3,CG31229,Sec61alpha,Sec63,CG8860,Tim17b* |
| GTP binding | GO:MF | 0.006 | *EMC10,Rab11,betaTub97EF,Ras85D,Rac2,Sar1,Rala,Rap1,Arf6,Arl8,Arf4,Rab5,Arf1,Rheb,Rab1,Arl4,Rab35,Rab6,Rac1,Arl1,SrpRbeta,alphaTub84D* |
| monoatomic cation transmembrane transporter activity | GO:MF | 0.022 | *Gat,zyd,CG5888,rumpel,pain,bumpel,CG10470,ine,Eaat1,VhaM9.7-a,Irk3,Vha14-1,Atpalpha,CG10413,Vha13,Vha26,Vha16-1,Vha68-2,VhaM9.7-c,fwe,Nmda1,VhaAC39-1,VhaPPA1-1,VhaM9.7-b,Vha44,Vha36-1,Vha55,BI-1,VhaSFD,Zip102B,Vha100-1* |
| protein transporter activity | GO:MF | 0.034 | *Sec61gamma,Hsc70-3,CG31229,Sec61alpha,Sec63,CG8860,Tim17b* |
| enoyl-CoA hydratase activity | GO:MF | 0.034 | *CG4598,Mtpalpha,Hacd2,Echs1* |
| protein-disulfide reductase activity | GO:MF | 0.038 | *SelT,CG9302,ERp60,CaBP1,CG5554,CG8993* |
| active transmembrane transporter activity | GO:MF | 0.038 | *Gat,zyd,rumpel,bumpel,CG32091,ine,Eaat1,VhaM9.7-a,Vha14-1,Atpalpha,CG10413,Vha13,Vha26,Vha16-1,Hsc70-3,Vha68-2,VhaM9.7-c,VhaAC39-1,VhaPPA1-1,VhaM9.7-b,Vha44,Vha36-1,Vha55,VhaSFD,Vha100-1* |
| actin binding | GO:MF | 0.04 | *Spn,CLIP-190,zormin,form3,Frl,Chd64,hts,Unc-115a,CG32264,pigs,Arpc5,Rab6,CG6891,cib,cpa,Arpc4,Arp3* |
| clathrin adaptor activity | GO:MF | 0.045 | *AP-2alpha,AP-2mu,AP-2sigma,AP-1gamma* |

### 1.19. Gene Ontology Enrichment Analysis: visceral muscle

Significant GO terms (p < 0.05) with associated genes

| GO Term | Source | P-value | Genes |
| --- | --- | --- | --- |
| striated muscle cell differentiation | GO:BP | 1.07e-15 | *up,wupA,Mlc2,Mp20,Tm2,Prm,bt,Zasp52,Mlp84B,how,sals,Rho1,Dg,kon,rhea,Mef2* |
| muscle cell development | GO:BP | 3.13e-15 | *up,wupA,Mlc2,Tm2,Prm,bt,Zasp52,Mlp84B,how,sals,Dg,kon,rhea,Mef2* |
| animal organ morphogenesis | GO:BP | 4.21e-12 | *disco-r,NetA,up,dlp,vn,ed,Wnt4,how,wb,14-3-3zeta,flw,sqa,fz2,hth,Galphao,Rho1,tsr,pyd,sli,Dg,cv-c,stck,ics,CG43658,rhea,dally,cic,bab2,Rip11,Usp47* |
| sarcomere organization | GO:BP | 5.72e-12 | *up,wupA,Tm2,bt,Mlp84B,how,sals,Dg,kon,rhea* |
| female gonad development | GO:BP | 3.27e-05 | *Wnt4,fz2,tsr,bab2* |
| glial cell migration | GO:BP | 1.38e-04 | *NetA,how,Pvf3,Rho1,sli* |
| muscle attachment | GO:BP | 1.56e-04 | *CAP,how,Dg,kon,rhea* |
| apposition of dorsal and ventral imaginal disc-derived wing surfaces | GO:BP | 1.86e-04 | *how,stck,ics,rhea* |
| female sex differentiation | GO:BP | 3.09e-04 | *Wnt4,fz2,tsr,bab2* |
| non-canonical Wnt signaling pathway | GO:BP | 3.24e-04 | *Wnt4,fz2,Galphao* |
| cell surface receptor signaling pathway | GO:BP | 9.15e-04 | *dlp,vn,ed,Wnt4,14-3-3zeta,Pvf3,flw,fz2,Galphao,Rho1,sli,stck,ics,Lmpt,dally,cic,Usp47* |
| cortical actin cytoskeleton organization | GO:BP | 0.001 | *flw,Galphao,Rho1,tsr,cv-c* |
| synaptic target inhibition | GO:BP | 0.002 | *Wnt4,fz2,sli* |
| negative regulation of signal transduction | GO:BP | 0.003 | *Strn-Mlck,ed,flw,Rho1,Hr4,cv-c,stck,ics,Lmpt,CaMKII,dally* |
| digestive tract mesoderm development | GO:BP | 0.003 | *wb,sli* |
| motor neuron axon guidance | GO:BP | 0.003 | *NetA,Wnt4,fz2,Rho1,dally* |
| muscle thin filament assembly | GO:BP | 0.005 | *up,bt* |
| positive regulation of canonical Wnt signaling pathway | GO:BP | 0.006 | *dlp,flw,dally,Usp47* |
| positive regulation of fibroblast growth factor receptor signaling pathway | GO:BP | 0.008 | *dlp,dally* |
| dorsal closure | GO:BP | 0.009 | *ed,Rho1,pyd,cv-c,stck* |
| negative regulation of JNK cascade | GO:BP | 0.011 | *flw,stck,ics* |
| muscle organ morphogenesis | GO:BP | 0.011 | *up,ed* |
| flight | GO:BP | 0.013 | *Mlc2,Msp300* |
| muscle cell cellular homeostasis | GO:BP | 0.014 | *up,wupA,Dg* |
| tracheal pit formation in open tracheal system | GO:BP | 0.015 | *Rho1,cv-c* |
| positive regulation of hippo signaling | GO:BP | 0.016 | *ed,14-3-3zeta,pyd* |
| head involution | GO:BP | 0.017 | *pyd,cv-c,stck* |
| regulation of myoblast fusion | GO:BP | 0.018 | *Mp20,Rho1* |
| protein autophosphorylation | GO:BP | 0.018 | *bt,sqa,CaMKII* |
| retinal ganglion cell axon guidance | GO:BP | 0.021 | *Wnt4,fz2* |
| maintenance of epithelial integrity, open tracheal system | GO:BP | 0.023 | *cv-c,rhea* |
| segment polarity determination | GO:BP | 0.026 | *dlp,fz2,dally* |
| imaginal disc-derived wing vein morphogenesis | GO:BP | 0.03 | *vn,cv-c,dally* |
| leg disc proximal/distal pattern formation | GO:BP | 0.033 | *disco,vn* |
| negative regulation of Rho protein signal transduction | GO:BP | 0.035 | *cv-c* |
| carbohydrate storage | GO:BP | 0.035 | *Mef2* |
| negative regulation of synaptic vesicle recycling | GO:BP | 0.035 | *sky* |
| regulation of lamellipodium assembly | GO:BP | 0.035 | *Tm1,tsr* |
| basement membrane assembly involved in embryonic body morphogenesis | GO:BP | 0.035 | *Ndg* |
| peptidyl-threonine phosphorylation | GO:BP | 0.035 | *bt,CaMKII* |
| negative regulation of semaphorin-plexin signaling pathway | GO:BP | 0.035 | *dally* |
| induction of negative chemotaxis | GO:BP | 0.035 | *sli* |
| female gonad morphogenesis | GO:BP | 0.035 | *bab2* |
| regulation of cardiac muscle contraction by calcium ion signaling | GO:BP | 0.035 | *up* |
| cell communication by electrical coupling | GO:BP | 0.035 | *shakB* |
| negative regulation of R8 cell differentiation | GO:BP | 0.035 | *ed* |
| regulation of morphogenesis of an epithelium | GO:BP | 0.035 | *flw* |
| regulation of protein localization to membrane | GO:BP | 0.035 | *dlp,dally* |
| dorsal closure, leading edge cell differentiation | GO:BP | 0.035 | *Rho1* |
| gap junction assembly | GO:BP | 0.035 | *shakB* |
| peptidyl-threonine autophosphorylation | GO:BP | 0.035 | *CaMKII* |
| regulation of actin cytoskeleton organization by cell-cell adhesion | GO:BP | 0.035 | *ed* |
| cardiac muscle cell-cardiac muscle cell adhesion | GO:BP | 0.035 | *rhea* |
| wing disc dorsal/ventral pattern formation | GO:BP | 0.035 | *dlp,dally,cic* |
| negative regulation of MAPK cascade | GO:BP | 0.038 | *flw,stck,ics* |
| epidermal cell differentiation | GO:BP | 0.038 | *Galphao,Rho1,tsr* |
| salivary gland boundary specification | GO:BP | 0.041 | *hth,sli* |
| peripheral nervous system development | GO:BP | 0.042 | *vn,hth,Rho1* |
| cell projection assembly | GO:BP | 0.044 | *Tm1,vn,Rho1,tsr,kon,CaMKII* |
| germ-line stem-cell niche homeostasis | GO:BP | 0.048 | *Wnt4,fz2* |
| branch fusion, open tracheal system | GO:BP | 0.048 | *ed,pyd* |
| spiracle morphogenesis, open tracheal system | GO:BP | 0.048 | *Rho1,cv-c* |
| actin binding | GO:MF | 2.45e-10 | *zormin,wupA,Mp20,Mhc,Tm1,Tm2,Zasp52,Actn,sals,CG43897,Rho1,Msp300,tsr,CG1674,rhea* |
| protein binding | GO:MF | 4.97e-09 | *zormin,disco,NetA,up,wupA,Mlc2,Mp20,Mhc,Mlc1,Tm1,CAP,Tm2,Strn-Mlck,dlp,bt,Zasp52,vn,Mlp84B,bru3,ed,Fas3,Actn,Dhit,CG13551,Wnt4,14-3-3zeta,sals,Pvf3,CG34417,flw,sqa,CG43897,fz2,hth,Galphao,CG14687,Rho1,Msp300,tsr,CG42748,pyd,sli,Dg,stck,CG1674,ics,kon,CG13124,CG14207,rhea,miple2,CaMKII,CaMKI,Mef2,dally,SK,slbo,cic,sick,bab2,Rip11,SKIP* |
| cytoskeletal protein binding | GO:MF | 7.26e-09 | *zormin,up,wupA,Mlc2,Mp20,Mhc,Mlc1,Tm1,Tm2,Zasp52,Mlp84B,Actn,sals,CG43897,Rho1,Msp300,tsr,CG1674,rhea* |
| calcium ion binding | GO:MF | 2.54e-05 | *up,Mlc2,TpnC47D,Mlc1,Ndg,Actn,TpnC73F,Frq1,sli,Dg,CG31650* |
| actin filament binding | GO:MF | 2.54e-05 | *Mp20,Mhc,Tm1,Tm2,sals,Msp300,tsr,rhea* |
| actinin binding | GO:MF | 5.92e-04 | *Zasp52,Mlp84B,CG43897* |
| calcium/calmodulin-dependent protein kinase activity | GO:MF | 0.002 | *Strn-Mlck,CaMKII,CaMKI* |
| structural constituent of muscle | GO:MF | 0.004 | *bt,Mlp84B,Dg* |
| myosin light chain kinase activity | GO:MF | 0.004 | *Strn-Mlck,sqa* |
| cell adhesion molecule binding | GO:MF | 0.006 | *ed,Fas3,pyd,Dg,rhea* |
| muscle alpha-actinin binding | GO:MF | 0.012 | *Zasp52,CG43897* |
| growth factor activity | GO:MF | 0.012 | *vn,Pvf3,miple2* |
| alpha-actinin binding | GO:MF | 0.012 | *Zasp52,CG43897* |
| calmodulin binding | GO:MF | 0.012 | *CG14687,CaMKII,CaMKI,SK* |
| growth factor receptor binding | GO:MF | 0.012 | *vn,ed,Pvf3* |
| heparin binding | GO:MF | 0.012 | *vn,sli,miple2* |
| protein-containing complex binding | GO:MF | 0.013 | *Mp20,Mhc,Tm1,Tm2,sals,Galphao,Msp300,tsr,smal,rhea* |
| epidermal growth factor receptor binding | GO:MF | 0.023 | *vn,ed* |
| structural constituent of cytoskeleton | GO:MF | 0.023 | *Strn-Mlck,alphaTub84B,rhea* |
| signaling receptor binding | GO:MF | 0.023 | *vn,ed,Wnt4,Pvf3,Galphao,sli,rhea,miple2* |
| photoreceptor activity | GO:MF | 0.035 | *shakB,SK* |
| transcription coactivator binding | GO:MF | 0.035 | *14-3-3zeta,hth* |
| myosin heavy chain binding | GO:MF | 0.035 | *Mlc2,Mlc1* |
| protein serine/threonine kinase activity | GO:MF | 0.035 | *Strn-Mlck,bt,sqa,Nuak,CaMKII,CaMKI* |
| glycosaminoglycan binding | GO:MF | 0.041 | *vn,sli,miple2* |
| cytoskeletal motor activator activity | GO:MF | 0.05 | *Strn-Mlck* |
| laminin receptor activity | GO:MF | 0.05 | *Dg* |
| laminin binding | GO:MF | 0.05 | *Dg* |
| Wnt-protein binding | GO:MF | 0.05 | *dlp,fz2* |
| small molecule binding | GO:MF | 0.05 | *disco,up,Mlc2,Act57B,TpnC47D,Mlp60A,Mhc,Mlc1,Strn-Mlck,bt,Zasp52,Ndg,Mlp84B,Actn,TpnC73F,CG31140,CG33521,flw,sqa,Frq1,Galphao,Nuak,Rho1,Hr4,Act87E,sli,Dg,stck,Lmpt,alphaTub84B,CG32280,CG14207,CaMKII,CaMKI,sick,CG31650* |
| ornithine decarboxylase inhibitor activity | GO:MF | 0.05 | *Oda* |
| small conductance calcium-activated potassium channel activity | GO:MF | 0.05 | *SK* |
| enzyme regulator activity | GO:MF | 0.05 | *TpnC47D,Mlc1,Dhit,TpnC73F,Frq1,Oda,cv-c,CG43658,sky,CG17124* |

### 1.20. Gene Ontology Enrichment Analysis: fatbody

Significant GO terms (p < 0.05) with associated genes

| GO Term | Source | P-value | Genes |
| --- | --- | --- | --- |
| mitochondrial electron transport, NADH to ubiquinone | GO:BP | 2.06e-29 | *ND-B14.5B,ND-PDSW,ND-B12,ND-19,ND-SGDH,CG9034,ND-B14,ND-B22,ND-13B,ND-B16.6,ND-B15,ND-B14.5A,ND-B18,ND-15,ND-18,ND-MWFE,ND-30,ND-39,ND-B17.2,ND-B17,CG11752,ND-24,ND-ASHI,ND-B14.7,ND-23,ND-13A,ND-20,ND-42* |
| mitochondrial translation | GO:BP | 3.27e-19 | *mRpL55,mRpL13,mRpL12,mRpL27,mRpL18,mRpS7,mRpS14,mRpL49,bonsai,mRpL20,mRpL54,mRpL36,mRpL23,mRpS17,mRpL30,mRpS18A,mRpL35,mRpL42,tko,mRpS10,mRpS26,mRpL14,mRpL34,mRpS18C,mRpS33,mRpS16,CG12848,mEFTu1,mRpS25,mRpL22* |
| mitochondrial respiratory chain complex I assembly | GO:BP | 1.70e-16 | *ND-B14.5B,ND-PDSW,ND-B12,ND-19,ND-SGDH,ND-B14,ND-B22,ND-B16.6,ND-B15,ND-B18,ND-15,ND-18,ND-39,ND-B17.2,ND-ASHI,ND-B14.7,ND-23,ND-20* |
| mitochondrial respirasome assembly | GO:BP | 4.27e-12 | *ND-B14.5B,ND-PDSW,COX6B,ND-B12,ND-19,COX7A,ND-SGDH,ND-B14,ND-B22,ND-B16.6,ND-B15,ND-B18,ND-15,ND-18,ND-39,ND-B17.2,ND-ASHI,ND-B14.7,ND-23,ND-20* |
| nucleoside triphosphate biosynthetic process | GO:BP | 2.58e-11 | *ATPsynC,ATPsynF,awd,ATPsynE,ATPsynG,ATPsynCF6,ATPsynD,ATPsynO,blw,ATPsynB,sun,ATPsyngamma,ATPsyndelta,ATPsynbeta,sesB* |
| protein localization to endoplasmic reticulum | GO:BP | 4.82e-10 | *Sec61beta,Sec61gamma,CG5885,CG8860,Srp14,TRAM,Spase22-23,Sec61alpha,Spase25,Srp19,Srp9,Spase12,SrpRalpha* |
| mitochondrial electron transport, ubiquinol to cytochrome c | GO:BP | 1.01e-09 | *UQCR-14,ox,RFeSP,UQCR-Q,UQCR-11,Cyt-c1,UQCR-6.4,UQCR-C2,Cyt-c-p* |
| ribose phosphate metabolic process | GO:BP | 1.45e-09 | *ATPsynC,Ak2,Paics,ATPsynF,awd,ATPsynE,ATPsynG,ATPsynCF6,Adk2,ATPsynD,ATPsynO,Aprt,blw,ATPsynB,sun,ATPsyngamma,ATPsyndelta,Prps,ATPsynbeta,sesB,Cmpk,HINT1,AdSS* |
| mitochondrial electron transport, cytochrome c to oxygen | GO:BP | 6.48e-09 | *CG7630,COX5B,COX4,COX5A,COX8,levy,COX7C,cype,COX7A,Cyt-c-p* |
| proton transmembrane transport | GO:BP | 1.18e-08 | *ATPsynC,ATPsynF,ATPsynE,ATPsynG,ATPsynCF6,ATPsynD,ATPsynO,blw,levy,ND-MLRQ,ATPsynB,sun,RFeSP,Cyt-c1,ATPsyngamma,ATPsynbeta,ND-20* |
| protein import into mitochondrial matrix | GO:BP | 2.97e-06 | *Roe1,CG6878,Hsc70-5,Tom7,CG7394,mge,Tom20,Tom40,Tim23* |
| SRP-dependent cotranslational protein targeting to membrane, translocation | GO:BP | 2.70e-05 | *Sec61beta,Sec61gamma,CG8860,TRAM,Sec61alpha* |
| signal peptide processing | GO:BP | 5.78e-05 | *twr,Spase22-23,Spase25,Spp,Spase12* |
| organic acid metabolic process | GO:BP | 2.30e-04 | *CG6415,ppl,Acbp2,hll,Shmt,Had1,Hn,arg,CG11134,P5CS,Dhpr,Gdh,P5cr-2,PCB,Echs1,Mdh1,ND-ACP,Cth,yip2,scu,CG7920,Etfb,CG4598,Ahcy,PH4alphaEFB,CG15771,Fum1,Pdhb* |
| post-translational protein targeting to membrane, translocation | GO:BP | 5.05e-04 | *Sec61beta,Sec61gamma,CG8860,Sec61alpha* |
| snRNA pseudouridine synthesis | GO:BP | 5.33e-04 | *NHP2,CG7637,Nop60B* |
| proton motive force-driven mitochondrial ATP synthesis | GO:BP | 9.33e-04 | *ATPsynF,ATPsynO,sun,ATPsynbeta* |
| amino acid catabolic process | GO:BP | 0.002 | *CG6415,ppl,Shmt,Hn,arg,Dhpr,Gdh,Etfb,Ahcy* |
| protein localization to membrane | GO:BP | 0.002 | *Sec61beta,Sec61gamma,CG34132,CG5885,Tim8,CG6878,CG8860,Tom7,Srp14,mge,TRAM,Sec61alpha,Tim9a,Srp19,Srp9,SrpRalpha* |
| protein insertion into mitochondrial inner membrane | GO:BP | 0.004 | *CG34132,Tim8,CG6878,Tim9a* |
| positive regulation of cytochrome-c oxidase activity | GO:BP | 0.012 | *levy,ND-MLRQ* |
| snoRNA guided rRNA pseudouridine synthesis | GO:BP | 0.012 | *CG7637,CG4038* |
| amino acid biosynthetic process | GO:BP | 0.029 | *Shmt,Hn,CG11134,P5CS,P5cr-2,Cth* |
| glycine decarboxylation via glycine cleavage system | GO:BP | 0.032 | *CG6415,ppl* |
| sperm DNA decondensation | GO:BP | 0.032 | *P32,Nph,Nlp* |
| protein processing | GO:BP | 0.033 | *CG5390,sel,twr,UQCR-C1,Spase22-23,Spase25,Spp,Spase12,CG8320* |
| determination of adult lifespan | GO:BP | 0.045 | *Trx-2,ATPsynD,levy,Sod1,sun,Prx3,ND-SGDH,sesB,Sam-S,ND-20,foxo,SdhC* |
| structural constituent of ribosome | GO:MF | 2.47e-16 | *mRpL55,mRpL13,RpL23,mRpL12,RpS2,RpL34a,mRpL27,mRpL18,RpS5a,mRpS7,mRpS14,mRpL49,bonsai,mRpL20,mRpL54,mRpL36,RpLP0-like,mRpL23,mRpS17,mRpL30,mRpS18A,mRpL35,mRpL42,tko,mRpS10,mRpS26,mRpL14,mRpL34,mRpS18C,mRpS33,mRpS16,mRpS25,mRpL22* |
| oxidoreductase activity | GO:MF | 2.05e-14 | *glob1,Akr1B,Fmo-2,Aldh7A1,Had1,Hn,P5CS,GstE6,Ldsdh1,Dhpr,Fdh,Gdh,CG7322,P5cr-2,Trx-2,COX5A,COX8,CG31548,ND-B14.5B,Mdh1,ND-PDSW,ND-MLRQ,COX6B,Sod1,ND-B12,cype,RFeSP,Plod,ND-19,Prx3,COX7A,ND-SGDH,Cyt-c1,ND-B14,ND-B22,ND-13B,scu,ND-B15,ND-B14.5A,ND-B18,Aldh-III,GstT4,ND-30,ND-39,Sod2,CG15717,ND-B17.2,Grx5,ND-B17,CaBP1,ND-24,ND-ASHI,ND-23,PH4alphaEFB,ND-20,SdhD,SdhC,Pdhb,ND-42* |
| proton-transporting ATP synthase activity, rotational mechanism | GO:MF | 3.75e-13 | *ATPsynC,ATPsynF,ATPsynE,ATPsynG,ATPsynCF6,ATPsynD,ATPsynO,blw,ATPsynB,sun,ATPsyngamma,ATPsyndelta,ATPsynbeta* |
| proton channel activity | GO:MF | 2.38e-12 | *ATPsynC,ATPsynF,ATPsynE,ATPsynG,ATPsynCF6,ATPsynD,ATPsynO,blw,ATPsynB,sun,ATPsyngamma,ATPsyndelta,ATPsynbeta* |
| proton transmembrane transporter activity | GO:MF | 5.39e-10 | *ATPsynC,ATPsynF,ATPsynE,ATPsynG,ATPsynCF6,ATPsynD,ATPsynO,blw,ATPsynB,sun,RFeSP,Cyt-c1,ATPsyngamma,ATPsyndelta,ATPsynbeta,ND-30,ND-24,ND-23,ND-20* |
| protein transmembrane transporter activity | GO:MF | 9.56e-06 | *Sec61gamma,CG8860,Hsc70-5,Tom7,Tom20,Tom40,Sec61alpha,Tim23* |
| protein transporter activity | GO:MF | 9.56e-06 | *Sec61gamma,Tim8,CG8860,Hsc70-5,Tom7,Tom20,Tom40,Sec61alpha,Tim23* |
| ligase activity | GO:MF | 9.64e-06 | *hll,ATPsynC,Paics,PCB,ATPsynF,ATPsynE,ATPsynG,ATPsynCF6,ATPsynD,ATPsynO,blw,ATPsynB,sun,ATPsyngamma,ATPsyndelta,ATPsynbeta,Scsalpha1,AdSS* |
| structural molecule activity | GO:MF | 2.39e-05 | *mRpL55,mRpL13,RpL23,mRpL12,RpS2,RpL34a,His2Av,mRpL27,mRpL18,RpS5a,Tina-1,His3.3A,mRpS7,mRpS14,mRpL49,bonsai,mRpL20,mRpL54,mRpL36,RpLP0-like,mRpL23,mRpS17,mRpL30,mRpS18A,betaTub97EF,mRpL35,mRpL42,Sec13,tko,mRpS10,mRpS26,mRpL14,mRpL34,mRpS18C,mRpS33,mRpS16,mRpS25,mRpL22* |
| electron transfer activity | GO:MF | 3.67e-05 | *RFeSP,Cyt-c1,Etfb,ND-30,ND-24,ND-23,Cyt-c-p,ND-20,SdhC* |
| RNA binding | GO:MF | 8.47e-05 | *Shmt,hoip,fus,Nph,NHP2,mRpL13,RpL23,mRpL12,RpS2,Nop56,CG7637,Nop60B,mRpL18,RpS5a,eIF2beta,mRpS7,mRpL20,CNBP,Srp14,Nlp,nop5,mRpS18A,clu,mod,Surf6,CG1542,Dp1,La,Tudor-SN,mRpS18C,vig2,Fib,CG4038,SmE,Srp19,Srp9,SmB,Rbp1,SrpRalpha,eIF2alpha,SC35,SmD2* |
| catalytic activity | GO:MF | 2.22e-04 | *glob1,CG6415,Idgf4,Akr1B,SpdS,Idgf2,hll,Fmo-2,Aldh7A1,Shmt,Had1,DNaseII,UK114,CG3505,CG8745,CG8586,Gbp2,Hn,CG16758,arg,Pu,ATPsynC,CG11134,fbp,P5CS,Hsp60A,GstE6,Ldsdh1,Dhpr,Fdh,Snp,Gdh,Cyp1,CG5390,CG7322,Ak2,Paics,P5cr-2,PCB,ATPsynF,eas,Fkbp39,awd,ATPsynE,GstE12,ATPsynG,ATPsynCF6,Fkbp14,Taldo,Trx-2,CG5793,Adk2,Fkbp12,ATPsynD,Echs1,COX5A,COX8,CG31548,ND-B14.5B,ATPsynO,Mdh1,CG8036,Aprt,ND-PDSW,CG14715,blw,ND-MLRQ,ATPsynB,COX6B,Sod1,Nop60B,ND-B12,Tina-1,sun,cype,Cth,RFeSP,Plod,Nurf-38,ND-19,Prx3,COX7A,ND-SGDH,Cyt-c1,p23,ND-B14,ATPsyngamma,ND-B22,Hsc70-5,CG31915,yip2,ND-13B,Polr2L,scu,ATPsyndelta,CG7920,ND-B15,Prps,ND-B14.5A,Pdi,ND-B18,betaTub97EF,Aldh-III,ATPsynbeta,CG4598,twr,GstT4,Rs1,ND-30,ND-39,Tudor-SN,Sod2,Fib,CG15717,ND-B17.2,UQCR-C1,Grx5,prtp,mEFTu1,ND-B17,Cmpk,CaBP1,Polr2F,Scsalpha1,HINT1,Spase22-23,ND-24,Spase25,CG1703,ND-ASHI,ND-23,Sam-S,Cchl,Ahcy,PH4alphaEFB,ND-20,SdhD,Ran,CG15771,Fum1,Ostgamma,Spp,SdhC,Pdhb,SrpRalpha,AdSS,ND-42* |
| box H/ACA snoRNA binding | GO:MF | 7.43e-04 | *NHP2,CG7637,CG4038* |
| rRNA binding | GO:MF | 0.002 | *RpL23,mRpL18,RpS5a,mRpS7,mRpL20,mRpS18A,La,mRpS18C* |
| snoRNA binding | GO:MF | 0.004 | *NHP2,Nop56,CG7637,nop5,CG4038* |
| aldehyde dehydrogenase (NAD+) activity | GO:MF | 0.009 | *Aldh7A1,Fdh,Aldh-III* |
| iron-sulfur cluster binding | GO:MF | 0.009 | *RFeSP,Grx5,CG3420,ND-24,ND-23,IscU,ND-20,Cisd2* |
| inorganic cation transmembrane transporter activity | GO:MF | 0.012 | *ATPsynC,ATPsynF,ATPsynE,ATPsynG,ATPsynCF6,ATPsynD,ATPsynO,blw,ATPsynB,sun,RFeSP,Cyt-c1,ATPsyngamma,ATPsyndelta,ATPsynbeta,ND-30,ND-24,ND-23,ND-20* |
| 2 iron, 2 sulfur cluster binding | GO:MF | 0.013 | *RFeSP,CG3420,ND-24,IscU,Cisd2* |
| 7S RNA binding | GO:MF | 0.014 | *Srp14,Srp19,Srp9* |
| procollagen galactosyltransferase activity | GO:MF | 0.015 | *Plod,CG31915* |
| monoatomic cation channel activity | GO:MF | 0.02 | *ATPsynC,ATPsynF,ATPsynE,ATPsynG,ATPsynCF6,ATPsynD,ATPsynO,blw,ATPsynB,sun,ATPsyngamma,ATPsyndelta,ATPsynbeta* |
| vitamin binding | GO:MF | 0.024 | *CG5958,apolpp,Shmt,CG8745,CG8036,Cth,Plod,PH4alphaEFB* |
| monoatomic cation transmembrane transporter activity | GO:MF | 0.024 | *ATPsynC,ATPsynF,ATPsynE,ATPsynG,ATPsynCF6,ATPsynD,ATPsynO,blw,ATPsynB,sun,RFeSP,Cyt-c1,ATPsyngamma,ATPsyndelta,ATPsynbeta,ND-30,ND-24,ND-23,ND-20* |
| NADH dehydrogenase activity | GO:MF | 0.024 | *ND-24,ND-23,ND-20* |
| NADH dehydrogenase (ubiquinone) activity | GO:MF | 0.024 | *ND-30,ND-24,ND-23,ND-20* |
| peptidyl-prolyl cis-trans isomerase activity | GO:MF | 0.027 | *Cyp1,Fkbp39,Fkbp14,Fkbp12,CG14715* |
| isomerase activity | GO:MF | 0.029 | *Cyp1,Fkbp39,Fkbp14,Fkbp12,CG14715,Nop60B,p23,Pdi,CG4598,prtp,CaBP1* |
| FK506 binding | GO:MF | 0.03 | *Fkbp39,CG14715* |
| nucleobase-containing compound kinase activity | GO:MF | 0.045 | *Ak2,awd,Adk2,Cmpk* |
| oxidoreductase activity, acting on the aldehyde or oxo group of donors, NAD or NADP as acceptor | GO:MF | 0.049 | *Aldh7A1,P5CS,Fdh,Aldh-III,CG15717* |
| retinoid binding | GO:MF | 0.049 | *CG5958,apolpp* |
| 5S rRNA binding | GO:MF | 0.049 | *mRpL18,La* |
| mRNA binding | GO:MF | 0.049 | *Shmt,fus,mRpL13,mRpL12,RpS5a,eIF2beta,mRpS7,CNBP,clu,mod,Dp1,La,Rbp1,SC35* |
| ubiquinol-cytochrome-c reductase activity | GO:MF | 0.049 | *RFeSP,Cyt-c1* |

### 1.21. Gene Ontology Enrichment Analysis: surface glial cell

Significant GO terms (p < 0.05) with associated genes

| GO Term | Source | P-value | Genes |
| --- | --- | --- | --- |
| septate junction assembly | GO:BP | 8.81e-22 | *CG9628,moody,crok,Gli,cold,Lac,crim,Nrg,kune,sinu,pck,pasi2,pasi1,Tsf2,Cont,Nrx-IV,Galphai,nrv2,scrib,bou,Mcr,Galphao* |
| proton transmembrane transport | GO:BP | 3.19e-21 | *Vha16-1,Vha13,Vha100-2,Vha14-1,Vha68-2,Vha36-1,VhaAC39-1,Vha26,VhaPPA1-1,Vha55,VhaM9.7-b,Vha44,ATP6AP2,ATPsynC,ND-MLRQ,levy,ATPsynG,ATPsynbeta,ATPsynF,VhaSFD,ATPsynO,Cyt-c1,ATPsynCF6,RFeSP,ATPsynE,ATPsynD,sun,ATPsynB,ATPsyngamma,ND-20* |
| establishment of glial blood-brain barrier | GO:BP | 1.82e-15 | *moody,Nrg,kune,sinu,pck,pasi2,pasi1,Cont,cora,Nrx-IV,Galphai,nrv2,Galphao* |
| mitochondrial electron transport, NADH to ubiquinone | GO:BP | 1.97e-12 | *ND-B14.5A,ND-B22,ND-B16.6,ND-B14.5B,ND-B18,ND-18,ND-B15,ND-B14.7,ND-B14,CG11752,ND-MWFE,ND-30,CG9034,ND-PDSW,ND-ASHI,ND-20,ND-B12* |
| cell adhesion involved in heart morphogenesis | GO:BP | 3.78e-12 | *Gli,Lac,Nrg,sinu,Cont,cora,Nrx-IV,nrv2,Ggamma1,Galphao* |
| animal organ morphogenesis | GO:BP | 1.44e-11 | *svp,Gli,Mmp1,dve,Lac,Nrg,sinu,btsz,Fas2,M6,aos,Cont,Dg,14-3-3epsilon,cora,Nrx-IV,LamC,aPKC,nrv2,Ras85D,Rab11,Ggamma1,Exn,Wnt4,Rbfox1,scrib,cv-c,Rac2,Timp,caps,ATP6AP2,Frl,drk,zfh2,Rho1,dlp,Ubx,Mob2,Rab6,Arf1,Moe,EMC3,Mkp3,sdk,aop,fra,p120ctn,Egfr,Rap1,elB,spi,Rab5,Arf6,stck,emc,Rala,Rip11,flw,sqh,Cdep,ics,shot,Galphao,alphaSnap,Pkn* |
| dorsal closure | GO:BP | 3.35e-10 | *sinu,cora,Nrx-IV,Ras85D,Rab11,Ggamma1,scrib,cv-c,Rac2,Mcr,Rho1,aop,FER,Egfr,Rap1,spi,Rab5,stck,emc,shot,Pkn* |
| mitochondrial electron transport, ubiquinol to cytochrome c | GO:BP | 2.09e-09 | *Cyt-c-p,UQCR-11,UQCR-Q,UQCR-6.4,ox,UQCR-14,Cyt-c1,RFeSP,UQCR-C2* |
| regulation of tube length, open tracheal system | GO:BP | 1.27e-08 | *Mmp1,bark,kune,pasi2,pasi1,Fas2,nrv2,Mcr,Rho1,Egfr,sqh* |
| mitochondrial electron transport, cytochrome c to oxygen | GO:BP | 1.40e-08 | *Cyt-c-p,COX7C,levy,cype,COX4,COX8,COX7A,COX5A,COX5B,CG7630* |
| nucleoside triphosphate biosynthetic process | GO:BP | 1.41e-08 | *Vha68-2,sesB,ATPsynC,ATPsynG,ATPsynbeta,ATPsynF,ATPsynO,ATPsynCF6,ATPsynE,ATPsynD,sun,ATPsynB,ATPsyngamma* |
| mitochondrial respiratory chain complex I assembly | GO:BP | 2.55e-08 | *ND-B22,ND-B16.6,ND-B14.5B,ND-B18,ND-18,ND-B15,ND-B14.7,ND-B14,ND-PDSW,ND-ASHI,ND-20,ND-B12* |
| ribose phosphate metabolic process | GO:BP | 4.23e-08 | *Ald1,Tpi,Pglym78,Gapdh2,Dg,Vha68-2,sesB,Vha55,Pfk,ATPsynC,Pgk,ATPsynG,ATPsynbeta,ATPsynF,ATPsynO,ATPsynCF6,ATPsynE,ATPsynD,sun,ATPsynB,ATPsyngamma,Ak2* |
| mitochondrial respirasome assembly | GO:BP | 5.69e-07 | *CG12107,COX6B,ND-B22,COX7A,ND-B16.6,ND-B14.5B,ND-B18,ND-18,ND-B15,ND-B14.7,ND-B14,ND-PDSW,ND-ASHI,ND-20,ND-B12* |
| tail-anchored membrane protein insertion into ER membrane | GO:BP | 1.67e-06 | *EMC6,EMC5,EMC3,EMC8-9,EMC4,EMC7* |
| cortical actin cytoskeleton organization | GO:BP | 5.14e-06 | *moody,btsz,Galphai,Rab11,cv-c,Frl,ghi,Rho1,Moe,flw,Galphao,Arpc4* |
| protein localization to endoplasmic reticulum | GO:BP | 2.21e-05 | *Spase25,Sec61gamma,CG8860,CG5885,Spase12,KdelR,Sec61beta,CG13887,Spase22-23* |
| trans-synaptic signaling | GO:BP | 2.16e-04 | *svp,crok,crim,CG6583,Fas2,CG9336,cpo,Dg,Ent2,Manf,CG9338,sesB,Exn,Syb,cv-c,beta-Spec,bou,Pgk,Arf1,Snap29,alpha-Spec,Rap1,Arf6,Cadps,Sod1,alphaSnap* |
| photoreceptor cell fate commitment | GO:BP | 3.16e-04 | *svp,14-3-3epsilon,Ras85D,scrib,drk,aop,Egfr,Rap1,elB,spi,emc,Rala* |
| nerve maturation | GO:BP | 3.50e-04 | *Nrg,Cont,Nrx-IV* |
| maintenance of epithelial integrity, open tracheal system | GO:BP | 3.66e-04 | *Lac,M6,cv-c,Mkp3,Egfr* |
| glycolytic process | GO:BP | 3.72e-04 | *Ald1,Tpi,Pglym78,Gapdh2,Dg,Pfk,Pgk* |
| regulation of synaptic transmission, cholinergic | GO:BP | 3.89e-04 | *crok,crim,CG6583,CG9336,CG9338,bou* |
| regulation of potassium ion transmembrane transport | GO:BP | 3.89e-04 | *crok,crim,CG6583,CG9336,CG9338,nrv2* |
| behavioral response to ethanol | GO:BP | 4.28e-04 | *moody,Fas2,Akap200,Spn27A,cher,Egfr,spi,Arf6,vsg* |
| nucleoside diphosphate catabolic process | GO:BP | 4.52e-04 | *Ald1,Tpi,Pglym78,Gapdh2,Dg,Pfk,Pgk* |
| hemocyte migration | GO:BP | 5.03e-04 | *Pvf3,Ras85D,RhoL,Rac2,Rho1,Rap1* |
| proton motive force-driven mitochondrial ATP synthesis | GO:BP | 6.24e-04 | *ATPsynbeta,ATPsynF,ATPsynO,sun* |
| peripheral nervous system development | GO:BP | 6.24e-04 | *repo,Gli,egh,Ras85D,Trim9,Rho1,fra,Egfr,spi* |
| positive regulation of voltage-gated potassium channel activity | GO:BP | 9.74e-04 | *crok,crim,CG6583,CG9336,CG9338* |
| determination of adult lifespan | GO:BP | 9.74e-04 | *Tpi,14-3-3epsilon,sesB,Ras85D,GlyP,Trx-2,levy,aop,cher,VhaSFD,Egfr,ATPsynD,sun,Sod1,SdhC,ND-20* |
| signal peptide processing | GO:BP | 0.001 | *Spase25,Spase12,twr,Spase22-23* |
| nucleotide catabolic process | GO:BP | 0.001 | *Ald1,Tpi,Pglym78,Gapdh2,Dg,Pfk,CG17224,Pgk* |
| homophilic cell adhesion via plasma membrane adhesion molecules | GO:BP | 0.001 | *Lac,Fas1,Fas2,Cont,caps,Fas3,Bsg,sdk* |
| axon midline choice point recognition | GO:BP | 0.001 | *beta-Spec,Trim9,Sema1a,alpha-Spec,fra,shot* |
| tracheal outgrowth, open tracheal system | GO:BP | 0.001 | *Ras85D,Rac2,Timp,drk,Egfr* |
| regulation of monoatomic ion transmembrane transport | GO:BP | 0.001 | *crok,crim,CG6583,CG9336,CG9338,nrv2,Cnx99A,ND-MLRQ,levy* |
| motor neuron axon guidance | GO:BP | 0.002 | *Nrg,Fas2,Wnt4,Rac2,caps,for,Sema1a,Rho1,fra,cher* |
| terminal branching, open tracheal system | GO:BP | 0.002 | *Vha13,VhaAC39-1,Vha26,aPKC,Ras85D,VhaPPA1-1* |
| purine ribonucleotide catabolic process | GO:BP | 0.002 | *Ald1,Tpi,Pglym78,Gapdh2,Dg,Pfk,Pgk* |
| epidermal cell differentiation | GO:BP | 0.002 | *Gli,cora,scrib,ATP6AP2,Rho1,Rap1,sqh,Galphao* |
| positive regulation of oxidoreductase activity | GO:BP | 0.002 | *ND-MLRQ,levy,Fdx2* |
| symbiont entry into host cell | GO:BP | 0.003 | *Rab11,Rab5,Rab7,Arf4,Vps29,Vps60* |
| ommatidial rotation | GO:BP | 0.003 | *aos,Ras85D,Frl,Egfr,Rap1,spi* |
| R3/R4 cell fate commitment | GO:BP | 0.003 | *svp,scrib,aop,Rala* |
| spiracle morphogenesis, open tracheal system | GO:BP | 0.003 | *cv-c,Rho1,Egfr,spi,sqh* |
| melanotic encapsulation of foreign target | GO:BP | 0.003 | *Nrg,aPKC,RhoL,Exn,Rac2,Spn27A* |
| vacuolar acidification | GO:BP | 0.004 | *Vha16-1,Vha100-2,VhaAC39-1,Vha55* |
| stem cell fate commitment | GO:BP | 0.004 | *Ras85D,Egfr,spi* |
| photoreceptor cell fate determination | GO:BP | 0.004 | *Ras85D,Egfr,spi* |
| establishment of epithelial cell apical/basal polarity | GO:BP | 0.004 | *aPKC,scrib,Egfr* |
| lumen formation, open tracheal system | GO:BP | 0.005 | *btsz,Vha68-2,Rho1,Moe,Egfr,shot* |
| positive regulation of ERK1 and ERK2 cascade | GO:BP | 0.005 | *Ras85D,drk,Egfr,Rap1,spi* |
| positive regulation of wound healing | GO:BP | 0.005 | *Rac2,Rho1,Egfr,Pkn* |
| liquid clearance, open tracheal system | GO:BP | 0.006 | *bark,crim,pck,Vha13,Vha26* |
| peripheral nervous system neuron axonogenesis | GO:BP | 0.006 | *Trim9,fra* |
| tricellular tight junction assembly | GO:BP | 0.006 | *bark,M6* |
| positive regulation of cytochrome-c oxidase activity | GO:BP | 0.006 | *ND-MLRQ,levy* |
| long-term strengthening of neuromuscular junction | GO:BP | 0.006 | *beta-Spec,alpha-Spec* |
| gamma-aminobutyric acid catabolic process | GO:BP | 0.006 | *Ssadh,Gabat* |
| oenocyte delamination | GO:BP | 0.006 | *aos,aop* |
| adherens junction maintenance | GO:BP | 0.007 | *Rab11,Rho1,p120ctn* |
| post-translational protein targeting to membrane, translocation | GO:BP | 0.007 | *Sec61gamma,CG8860,Sec61beta* |
| Wnt protein secretion | GO:BP | 0.007 | *CHOp24,p24-1,bai,eca* |
| positive regulation of border follicle cell migration | GO:BP | 0.009 | *Gli,aop,Egfr,spi,sqh,spri* |
| signal release | GO:BP | 0.009 | *Exn,Syb,CHOp24,Arf1,p24-1,Snap29,Rap1,Arf6,bai,eca,Cadps,alphaSnap* |
| vacuolar proton-transporting V-type ATPase complex assembly | GO:BP | 0.009 | *CG5969,VhaAC45,ATP6AP2* |
| SRP-dependent cotranslational protein targeting to membrane, translocation | GO:BP | 0.009 | *Sec61gamma,CG8860,Sec61beta* |
| dorsal closure, spreading of leading edge cells | GO:BP | 0.009 | *Ras85D,Rho1,Egfr* |
| dorsal closure, elongation of leading edge cells | GO:BP | 0.009 | *Rab11,Rac2,FER* |
| striated muscle cell differentiation | GO:BP | 0.011 | *rost,Dg,hts,blow,Rac2,Rho1,Bsg,cher,Arf6* |
| pigment metabolic process | GO:BP | 0.012 | *Nrg,aPKC,RhoL,Exn,Rac2,CG5390,Cnx99A,Rho1,Spn27A,EMC3,roh* |
| intracellular monoatomic ion homeostasis | GO:BP | 0.013 | *Vha16-1,Vha100-2,zyd,Vha68-2,VhaAC39-1,nrv2,sesB,Vha55,VhaAC45,CG10470,CG17593* |
| endoplasmic reticulum to Golgi vesicle-mediated transport | GO:BP | 0.014 | *CHOp24,p24-1,KdelR,cni,bai,CG13887,eca,Sec13* |
| viral process | GO:BP | 0.015 | *Rab11,Rab5,Rab7,Arf4,Vps29,Vps60* |
| endoplasmic reticulum unfolded protein response | GO:BP | 0.015 | *Hsc70-3,CG32276,Xbp1,Der-1* |
| axon ensheathment in central nervous system | GO:BP | 0.016 | *Galphai,Galphao* |
| establishment of planar polarity of embryonic epithelium | GO:BP | 0.016 | *btsz,Rho1* |
| cortical microtubule organization | GO:BP | 0.016 | *Moe,shot* |
| maintenance of imaginal disc-derived wing hair orientation | GO:BP | 0.016 | *Gli,cora* |
| mitotic cleavage furrow ingression | GO:BP | 0.016 | *Arf1,AP-2alpha* |
| female germline ring canal formation, actin assembly | GO:BP | 0.016 | *hts,cher* |
| NADH metabolic process | GO:BP | 0.017 | *Mdh1,Pfk,Gdh* |
| tracheal pit formation in open tracheal system | GO:BP | 0.017 | *cv-c,Timp,Rho1* |
| dorsal closure, amnioserosa morphology change | GO:BP | 0.017 | *Rab11,FER,Rab5* |
| dendrite self-avoidance | GO:BP | 0.017 | *Fas2,Cont,Rho1,Bsg* |
| midgut development | GO:BP | 0.022 | *dve,cv-c,Gp93,Ubx,emc* |
| epithelial cell proliferation involved in Malpighian tubule morphogenesis | GO:BP | 0.022 | *svp,Egfr,spi* |
| regulation of circadian sleep/wake cycle, sleep | GO:BP | 0.023 | *crok,crim,CG6583,CG9336,CG9338,bou* |
| chorion-containing eggshell pattern formation | GO:BP | 0.028 | *Ras85D,Egfr,emc* |
| negative regulation of microtubule depolymerization | GO:BP | 0.028 | *beta-Spec,alpha-Spec,shot* |
| fructose 1,6-bisphosphate metabolic process | GO:BP | 0.028 | *Ald1,Pfk* |
| netrin-activated signaling pathway | GO:BP | 0.028 | *Trim9,fra* |
| head involution | GO:BP | 0.029 | *cv-c,Rac2,Mcr,stck,emc* |
| haltere development | GO:BP | 0.034 | *aos,Ubx,Egfr* |
| tricarboxylic acid cycle | GO:BP | 0.04 | *Mdh1,Idh,SdhB,CG5214,SdhC* |
| dsRNA transport | GO:BP | 0.041 | *Vha16-1,egh,VhaSFD,Rab7* |
| cytokinesis, division site positioning | GO:BP | 0.043 | *Rho1,Pkn* |
| establishment or maintenance of polarity of larval imaginal disc epithelium | GO:BP | 0.043 | *scrib,Moe* |
| protein localization to endosome | GO:BP | 0.043 | *scrib,Moe* |
| germarium-derived oocyte fate determination | GO:BP | 0.045 | *14-3-3epsilon,aPKC,hts,alpha-Spec* |
| biological process involved in interspecies interaction between organisms | GO:BP | 0.046 | *Cyp6a20,Nrg,GILT1,RNASEK,aPKC,Ras85D,Rab11,RhoL,Exn,GlyP,Rac2,CG5390,Mcr,Rho1,Trx-2,Spn27A,Rab6,Arf1,FER,Rab5,NUCB1,Rala,Rab7,GILT2,Arf4,Vps29,Vps60,Pp1alpha-96A* |
| imaginal disc-derived wing vein specification | GO:BP | 0.046 | *aos,Dg,Ras85D,Rbfox1,Mkp3,Egfr* |
| rhodopsin biosynthetic process | GO:BP | 0.048 | *Cnx99A,EMC3,roh* |
| compound eye cone cell differentiation | GO:BP | 0.048 | *sdk,Egfr,emc* |
| ceramide biosynthetic process | GO:BP | 0.048 | *Spt-I,schlank,ifc* |
| positive regulation of cell-cell adhesion | GO:BP | 0.048 | *RhoL,Flo1,Rap1* |
| sensory perception of pain | GO:BP | 0.048 | *CG9336,Ncc69,pain* |
| short-term memory | GO:BP | 0.05 | *Fas2,ATP6AP2,drk,for* |
| membrane fusion | GO:BP | 0.05 | *Rab11,Syb,aus,Snap29,Rab5,Rab7,alphaSnap* |
| proton transmembrane transporter activity | GO:MF | 2.70e-18 | *Vha16-1,Vha13,Vha100-2,Vha14-1,Vha68-2,Vha36-1,VhaAC39-1,Vha26,VhaPPA1-1,Vha55,VhaM9.7-b,Vha44,ATPsynC,ATPsynG,ATPsynbeta,ATPsynF,VhaSFD,ATPsynO,Cyt-c1,ATPsynCF6,RFeSP,ATPsynE,ATPsynD,sun,ATPsynB,ATPsyngamma,ND-30,ND-20* |
| proton-transporting ATPase activity, rotational mechanism | GO:MF | 2.33e-11 | *Vha16-1,Vha13,Vha100-2,Vha14-1,Vha68-2,Vha36-1,VhaAC39-1,Vha26,VhaPPA1-1,Vha55,VhaM9.7-b,Vha44,ATPsynbeta,VhaSFD* |
| proton-transporting ATP synthase activity, rotational mechanism | GO:MF | 3.00e-11 | *Vha68-2,ATPsynC,ATPsynG,ATPsynbeta,ATPsynF,ATPsynO,ATPsynCF6,ATPsynE,ATPsynD,sun,ATPsynB,ATPsyngamma* |
| proton channel activity | GO:MF | 1.65e-10 | *Vha68-2,ATPsynC,ATPsynG,ATPsynbeta,ATPsynF,ATPsynO,ATPsynCF6,ATPsynE,ATPsynD,sun,ATPsynB,ATPsyngamma* |
| ATPase-coupled monoatomic cation transmembrane transporter activity | GO:MF | 5.92e-10 | *Vha16-1,Vha13,Vha100-2,Vha14-1,Vha68-2,Vha36-1,VhaAC39-1,Vha26,VhaPPA1-1,Vha55,VhaM9.7-b,Vha44,ATPsynbeta,VhaSFD* |
| inorganic cation transmembrane transporter activity | GO:MF | 7.34e-10 | *Vha16-1,Vha13,Vha100-2,zyd,Vha14-1,Vha68-2,Ncc69,Vha36-1,VhaAC39-1,Vha26,VhaPPA1-1,Vha55,VhaM9.7-b,Vha44,pain,CG10470,ATPsynC,EMC5,ATPsynG,ATPsynbeta,ATPsynF,VhaSFD,ATPsynO,Cyt-c1,ATPsynCF6,RFeSP,ATPsynE,ATPsynD,sun,ATPsynB,Nmda1,ATPsyngamma,ND-30,ND-20* |
| monoatomic cation transmembrane transporter activity | GO:MF | 7.15e-09 | *Vha16-1,Vha13,Vha100-2,zyd,Vha14-1,Vha68-2,Ncc69,Vha36-1,VhaAC39-1,Vha26,VhaPPA1-1,Vha55,VhaM9.7-b,Vha44,pain,CG10470,ATPsynC,Piezo,ATPsynG,ATPsynbeta,ATPsynF,VhaSFD,ATPsynO,Cyt-c1,ATPsynCF6,RFeSP,ATPsynE,ATPsynD,sun,ATPsynB,Nmda1,ATPsyngamma,ND-30,ND-20* |
| membrane insertase activity | GO:MF | 1.87e-06 | *EMC6,EMC5,EMC3,EMC8-9,EMC4,EMC7* |
| monoatomic ion transmembrane transporter activity | GO:MF | 1.92e-06 | *Vha16-1,Vha13,Vha100-2,zyd,Vha14-1,Vha68-2,Ncc69,Vha36-1,VhaAC39-1,Vha26,VhaPPA1-1,Vha55,VhaM9.7-b,Vha44,pain,CG10470,ATPsynC,Clic,Piezo,ATPsynG,ATPsynbeta,ATPsynF,VhaSFD,ATPsynO,Cyt-c1,ATPsynCF6,RFeSP,ATPsynE,ATPsynD,sun,ATPsynB,Nmda1,ATPsyngamma,ND-30,ND-20* |
| oxidoreductase activity | GO:MF | 2.70e-06 | *Cyp6a20,Ssadh,GILT1,Gapdh2,SelT,Mdh1,GstD1,ifc,Clic,Gdh,ND-MLRQ,ND-B14.5A,Trx-2,cype,ERp60,CG9743,Aldh-III,CG5554,L2HGDH,COX6B,Sc2,ND-B22,Idh,COX8,COX7A,COX5A,GstS1,Cyt-c1,SdhB,ND-B14.5B,RFeSP,GILT2,CG5214,ND-B18,ND-B15,spidey,ND-B14,CG11007,Sod1,ND-30,SdhC,ND-PDSW,ND-ASHI,ND-20,ND-B12* |
| ATPase-coupled transmembrane transporter activity | GO:MF | 4.08e-06 | *Vha16-1,Vha13,Vha100-2,Vha14-1,Vha68-2,Vha36-1,VhaAC39-1,Vha26,VhaPPA1-1,Vha55,VhaM9.7-b,Vha44,Hsc70-3,ATPsynbeta,VhaSFD* |
| actin binding | GO:MF | 1.05e-05 | *cora,hts,Akap200,zormin,beta-Spec,Chd64,Unc-115a,Frl,Rho1,Arpc5,Rab6,Moe,alpha-Spec,cher,CLIP-190,GMF,shot,Arpc4* |
| active transmembrane transporter activity | GO:MF | 1.52e-05 | *Vha16-1,Vha13,Vha100-2,zyd,Vha14-1,Vha68-2,Ncc69,Vha36-1,VhaAC39-1,Vha26,sesB,VhaPPA1-1,Vha55,VhaM9.7-b,Vha44,Hsc70-3,ATPsynbeta,VhaSFD,Oatp74D,Cyt-c1,RFeSP,ND-30,ND-20,Mpcp2* |
| transporter activity | GO:MF | 2.05e-05 | *CG3168,Vha16-1,CG32407,Vha13,Vha100-2,ogre,zyd,Ent2,Vha14-1,Vha68-2,Ncc69,Vha36-1,spin,VhaAC39-1,Vha26,sesB,VhaPPA1-1,Vha55,VhaM9.7-b,Vha44,Inx2,pain,CG10470,ATPsynC,Clic,Hsc70-3,mrva,EMC5,Piezo,Sec61gamma,ATPsynG,CG8860,ATPsynbeta,ATPsynF,VhaSFD,ATPsynO,Oatp74D,Cyt-c1,ATPsynCF6,RFeSP,ATPsynE,ATPsynD,sun,CG33298,ATPsynB,Nmda1,ATPsyngamma,ND-30,Tom20,ND-20,Mpcp2* |
| transmembrane transporter activity | GO:MF | 2.05e-05 | *CG3168,Vha16-1,Vha13,Vha100-2,ogre,zyd,Ent2,Vha14-1,Vha68-2,Ncc69,Vha36-1,spin,VhaAC39-1,Vha26,sesB,VhaPPA1-1,Vha55,VhaM9.7-b,Vha44,Inx2,pain,CG10470,ATPsynC,Clic,Hsc70-3,mrva,EMC5,Piezo,Sec61gamma,ATPsynG,CG8860,ATPsynbeta,ATPsynF,VhaSFD,ATPsynO,Oatp74D,Cyt-c1,ATPsynCF6,RFeSP,ATPsynE,ATPsynD,sun,ATPsynB,Nmda1,ATPsyngamma,ND-30,Tom20,ND-20,Mpcp2* |
| guanyl ribonucleotide binding | GO:MF | 2.08e-05 | *betaTub56D,alphaTub84B,Galphai,Ras85D,Rab11,RhoL,Rac2,Gdh,for,Rho1,Rab6,Arf1,Rap1,EMC10,Rab5,Arf6,Rala,Rab7,Arf4,Galphao* |
| guanyl nucleotide binding | GO:MF | 2.08e-05 | *betaTub56D,alphaTub84B,Galphai,Ras85D,Rab11,RhoL,Rac2,Gdh,for,Rho1,Rab6,Arf1,Rap1,EMC10,Rab5,Arf6,Rala,Rab7,Arf4,Galphao* |
| GTPase activity | GO:MF | 3.26e-05 | *betaTub56D,Galphai,Ras85D,Rab11,RhoL,Rac2,Rho1,Rab6,Arf1,Rap1,EMC10,Rab5,Arf6,Rala,Rab7,Arf4,Galphao* |
| electron transfer activity | GO:MF | 3.43e-05 | *Cyt-c-p,Fdx2,Cyt-c1,SdhB,RFeSP,ND-30,SdhC,ND-20,Etfb* |
| GTP binding | GO:MF | 4.17e-05 | *betaTub56D,alphaTub84B,Galphai,Ras85D,Rab11,RhoL,Rac2,Gdh,Rho1,Rab6,Arf1,Rap1,EMC10,Rab5,Arf6,Rala,Rab7,Arf4,Galphao* |
| cell adhesion mediator activity | GO:MF | 1.09e-04 | *Fas2,Cont,Dg,Nrt,Bsg,sdk* |
| cell adhesion molecule binding | GO:MF | 1.38e-04 | *Nrg,Fas1,Fas2,Cont,Dg,Nrt,Fas3,Moe,Bsg,sdk,p120ctn* |
| monoatomic cation channel activity | GO:MF | 3.23e-04 | *zyd,Vha68-2,pain,CG10470,ATPsynC,Piezo,ATPsynG,ATPsynbeta,ATPsynF,ATPsynO,ATPsynCF6,ATPsynE,ATPsynD,sun,ATPsynB,Nmda1,ATPsyngamma* |
| structural constituent of cytoskeleton | GO:MF | 5.30e-04 | *betaTub56D,alphaTub84B,LamC,beta-Spec,Arpc5,Jupiter,Arpc4* |
| GPI anchor binding | GO:MF | 5.76e-04 | *crok,crim,CG6583,CG9336,CG9338* |
| actin filament binding | GO:MF | 0.004 | *hts,Akap200,beta-Spec,Chd64,Unc-115a,Frl,Arpc5,alpha-Spec,cher,Arpc4* |
| cytoskeletal protein binding | GO:MF | 0.007 | *AnxB9,cora,aPKC,hts,Akap200,zormin,beta-Spec,Chd64,Unc-115a,Frl,Rho1,Arpc5,Rab6,Moe,alpha-Spec,Jupiter,cher,CLIP-190,GMF,sqh,Cdep,shot,Arpc4* |
| oxidoreductase activity, acting on a sulfur group of donors | GO:MF | 0.008 | *GILT1,SelT,Trx-2,ERp60,CG5554,GILT2,CG11007* |
| channel activity | GO:MF | 0.009 | *ogre,zyd,Vha68-2,Inx2,pain,CG10470,ATPsynC,Clic,Piezo,ATPsynG,ATPsynbeta,ATPsynF,ATPsynO,ATPsynCF6,ATPsynE,ATPsynD,sun,ATPsynB,Nmda1,ATPsyngamma* |
| ligase activity | GO:MF | 0.009 | *Vha68-2,ATPsynC,ATPsynG,Naprt,ATPsynbeta,ATPsynF,ATPsynO,ATPsynCF6,ATPsynE,ATPsynD,sun,ATPsynB,ATPsyngamma* |
| calcium ion binding | GO:MF | 0.01 | *Nrg,Dg,Calr,AnxB9,Cnx99A,Clic,alpha-Spec,Alg-2,AnxB11,CG17593,NUCB1,sqh,shot,CG31650,CG17493* |
| cell adhesion receptor activity | GO:MF | 0.012 | *Nrt,sdk* |
| lipid binding | GO:MF | 0.012 | *crok,Npc2b,crim,CG6583,fabp,CG9336,AnxB9,CG9338,cv-c,beta-Spec,CG8176,ATPsynC,Clic,Moe,FER,AnxB11,Acbp1* |
| monoatomic ion channel activity | GO:MF | 0.014 | *zyd,Vha68-2,pain,CG10470,ATPsynC,Clic,Piezo,ATPsynG,ATPsynbeta,ATPsynF,ATPsynO,ATPsynCF6,ATPsynE,ATPsynD,sun,ATPsynB,Nmda1,ATPsyngamma* |
| protein binding | GO:MF | 0.018 | *svp,CG10702,fipi,Lac,fax,Nrg,sinu,btsz,Fas1,Fas2,Vha13,Vha100-2,Tpi,aos,Cont,Usf,Dg,14-3-3epsilon,Calr,AnxB9,cora,Pvf3,RabGGTb,CG32354,Galphai,Vha26,Clc,aPKC,Ras85D,Rab11,Act5C,Ggamma1,Lapsyn,CAP,RhoL,hts,Akap200,Exn,zormin,Wnt4,CG1105,Rbfox1,RhoGAP18B,Pfk,Syb,scrib,GlyP,blow,Rac2,beta-Spec,Chd64,Unc-115a,Timp,caps,ATP6AP2,Frl,drk,Gp93,pain,Cnx99A,SKIP,Mcr,ATPsynC,krz,Gdh,Trim9,Nrt,Hsc70-3,Sema1a,Rho1,dlp,CG12384,Fas3,Ubx,Mob2,kud,Arpc5,Rab6,Moe,Fkbp12,CG5004,Snap29,alpha-Spec,Alg-2,Jupiter,Bsg,sdk,aop,FER,fra,ATPsynbeta,cher,CLIP-190,p120ctn,Egfr,Rap1,elB,Sc2,GMF,spi,Rab5,Der-1,cni,stck,emc,Rala,Rip11,flw,Tpr2,Oatp74D,EMC8-9,sqh,CG11267,CG14818,CG11999,Cdep,CG2918,Sec13,ics,sun,robl,spri,Pp1alpha-96A,shot,Sod1,Galphao,ND-30,Arpc4,alphaSnap,Pkn,TfIIA-S,Max,Etfb,CG40228* |
| epidermal growth factor receptor binding | GO:MF | 0.023 | *aos,drk,spi* |
| G protein activity | GO:MF | 0.023 | *Ras85D,Rap1,Rala* |
| acetylpyruvate hydrolase activity | GO:MF | 0.028 | *CG5793,CG6028* |
| serine C-palmitoyltransferase activity | GO:MF | 0.028 | *Spt-I,ghi* |
| ribonucleoside triphosphate phosphatase activity | GO:MF | 0.028 | *betaTub56D,Vha68-2,Galphai,Ras85D,Rab11,RhoL,Rac2,Gp93,Hsc70-3,Rho1,Rab6,Arf1,ATPsynbeta,Rap1,EMC10,Rab5,Arf6,Rala,Rab7,CG2918,Arf4,CG33298,Galphao* |
| GDP binding | GO:MF | 0.038 | *Ras85D,Rap1,Rala* |
| disulfide oxidoreductase activity | GO:MF | 0.039 | *SelT,Trx-2,ERp60,CG5554,CG11007* |
| cell-cell adhesion mediator activity | GO:MF | 0.047 | *Fas2,Cont,Bsg* |
| growth factor receptor binding | GO:MF | 0.048 | *aos,Pvf3,drk,spi* |
| protein transmembrane transporter activity | GO:MF | 0.048 | *Hsc70-3,Sec61gamma,CG8860,Tom20* |
| ubiquinol-cytochrome-c reductase activity | GO:MF | 0.048 | *Cyt-c1,RFeSP* |

### 1.22. Gene Ontology Enrichment Analysis: wing heart precursor

Significant GO terms (p < 0.05) with associated genes

| GO Term | Source | P-value | Genes |
| --- | --- | --- | --- |
| animal organ morphogenesis | GO:BP | 3.04e-21 | *rau,cv-2,AdamTS-A,Lac,Hs6st,bab2,Myo31DF,dally,tsh,CG34347,N,RhoGAP15B,unk,aos,cdi,robo2,klu,Fas2,cv-c,Dg,Mkp3,noc,fng,kibra,sinu,aop,Gprk2,RhoGAP71E,Dys,CG42788,Nrg,sbb,eve,mthl5,how,spg,prc,Cont,ptc,hth,ttk,cora,tup,dlg1,unc-5,Galphao,dos,stg,cic,numb,spi,ed,vn,pnut,l(2)gl,Mnn1,Shc,Rip11,Egfr,shn,myo,sdk,Rab5,sd,par-1,Mmp2,ics,Efa6,bun,RhoGEF2,klar,Hipk,stck,Rho1,Ype,spen,Cka,Ilk,Ras85D,RhoGAP68F,ci,dia,Nrx-IV,aPKC,crol,ec,gbb,Cdep,Slik,msi,ftz-f1,Asap,Pura,elB,raw,Cyfip,Stim,bdg* |
| cell surface receptor signaling pathway | GO:BP | 3.31e-20 | *rau,cv-2,Pvf3,hppy,dally,tsh,FER,wgn,N,htl,edl,aos,cdi,robo2,Mp,ken,Sulf1,Fas2,E(spl)m3-HLH,lin-28,Toll-7,E(spl)mbeta-HLH,gcl,Pdk1,Mkp3,noc,InR,fng,Ptp4E,aop,Gprk2,Sema1a,GluRIID,E(spl)m6-BFM,mthl5,CycG,spg,Usp12-46,ptc,ttk,dlg1,unc-5,Galphao,dos,cic,numb,spi,stumps,ed,vn,PRAS40,l(2)gl,Tnks,Cirl,CG45050,Akap200,nkd,Shc,Rheb,Egfr,Dyrk2,shn,Mnr,myo,par-1,Mmp2,ics,Ehbp1,Sema2a,Hipk,stck,Rho1,spen,pbl,Ilk,Ras85D,ci,Pitslre,aPKC,crol,gbb,RanBPM,Sirt1,RalGPS,lili,elB,mth,Rtf1,Akt,Lnk,Mob4,aph-1,Duba,Mkrn1,Graf,Hsp83* |
| septate junction assembly | GO:BP | 5.29e-15 | *Lac,CG9628,crim,crok,pasi2,pck,cold,sinu,Nrg,Cont,pasi1,kune,dlg1,Galphao,CG44325,l(2)gl,wun,Nrx-IV,bou* |
| negative regulation of signal transduction | GO:BP | 9.96e-14 | *hppy,dally,edl,aos,cdi,ken,Sulf1,Fas2,cv-c,gcl,Mkp3,fng,Ptp4E,aop,Gprk2,E(spl)m6-BFM,scyl,chrb,CycG,Usp12-46,ptc,LRR,dlg1,numb,ed,PRAS40,l(2)gl,Mnn1,nkd,Rheb,Egfr,Spred,Atg9,par-1,Mmp2,ics,Hr4,Ufd4,raskol,Hipk,stck,Rho1,pbl,Cka,Ras85D,ci,aPKC,PNUTS,crol,fmt,Ubc6,raw,Akt,Mob4,Duba,AMPKalpha,Graf* |
| establishment of glial blood-brain barrier | GO:BP | 3.00e-09 | *pasi2,pck,sinu,Nrg,Cont,pasi1,kune,cora,Galphao,Nrx-IV* |
| dorsal closure | GO:BP | 3.75e-08 | *FER,N,cv-c,sinu,aop,cora,tup,dlg1,spi,ed,l(2)gl,Egfr,shn,Rab5,stck,Rho1,Cka,Ras85D,Nrx-IV,gbb,raw* |
| positive regulation of neuroblast proliferation | GO:BP | 7.38e-08 | *N,klu,E(spl)m3-HLH,E(spl)mbeta-HLH,InR,Rheb,bun,Cka,aPKC,Mob4,Hsp83* |
| cell adhesion involved in heart morphogenesis | GO:BP | 9.35e-08 | *Lac,sinu,Nrg,prc,Cont,cora,Galphao,Nrx-IV* |
| regulation of metabolic process | GO:BP | 1.36e-07 | *sens-2,CG14431,CG9650,Pka-R2,bab2,Atf3,Antp,tsh,N,htl,Pax,Hand,edl,Eip78C,bru2,aos,CG43366,ken,BNIP3,Tet,klu,E(spl)m3-HLH,Dg,lin-28,CG3662,Toll-7,luna,E(spl)mbeta-HLH,gcl,noc,InR,Atg17,kibra,aop,zfh1,jigr1,Gprk2,Glut4EF,bwa,sbb,eve,how,chinmo,AGO1,CycG,Lk6,Usp12-46,Sox14,REPTOR-BP,hth,ttk,Trf2,REPTOR,tup,Oda,CG13124,CG8312,cic,Tob,numb,Pur-alpha,Edc3,spi,Mes2,slbo,Atg8a,elm,PRAS40,l(2)gl,Mnn1,Tnks,Rheb,Pink1,Egfr,tna,AGO2,UbcE2H,shn,ps,CG6770,CG12384,cer,sd,CG1764,par-1,ics,CG10543,Pits,Hr4,bun,Ufd4,CG4911,Smr,Hipk,Atg18a,Rho1,spen,Taf3,Cka,Ras85D,Mnt,maf-S,CG6701,ci,Pitslre,Not1,Psc,PNUTS,PAN3,CG32369,crol,dmpd,RanBPM,Sirt1,simj,Slik,msi,Crtc,CDK2AP1,ova,Adf1,Bruce,ko,ftz-f1,Lamtor1,Ubc6,Pcf11,elB,Abi,Svip,Rtf1,Max,Hmt4-20,Akt,CG10803,DnaJ-1,Smg5,CG2765,Cyfip,Daxx,Axud1,ClpX,sordd1,CG7879,AMPKalpha,Mkrn1,Clamp,rgr,CG12099,Hsp83* |
| cortical actin cytoskeleton organization | GO:BP | 2.31e-06 | *cv-c,Fim,Galphao,l(2)gl,RhoGEF2,fwd,Rho1,pbl,dia,Slik,mtm,Abi,Cyfip,Graf* |
| motor neuron axon guidance | GO:BP | 1.36e-05 | *dally,N,Mp,Sulf1,Fas2,Ptp4E,zfh1,Sema1a,Nrg,eve,tup,unc-5,Mmp2,Rho1,ko* |
| viral process | GO:BP | 2.35e-05 | *Rab2,shrb,AGO2,Rab5,Rab1,Rab8,Rab7,Vps60,Rab14,TSG101,Hlc* |
| positive regulation of border follicle cell migration | GO:BP | 4.94e-05 | *InR,aop,ttk,spi,slbo,vn,Egfr,par-1,spri,Akt* |
| gastrulation | GO:BP | 5.54e-05 | *Dtg,htl,Gprk2,RhoGAP71E,eve,stg,spi,stumps,RhoGEF2,Rho1,pbl,Ras85D,RhoGAP68F,cta,SelG* |
| determination of adult lifespan | GO:BP | 1.33e-04 | *N,CG42663,InR,aop,mthl5,Atg8a,Mnn1,Egfr,myo,bun,fwd,Ras85D,miple2,Sirt1,Indy,mth,SelG,VhaSFD,Daxx,Hsp27,Hsp26* |
| spiracle morphogenesis, open tracheal system | GO:BP | 1.83e-04 | *cv-c,Trf2,spi,Egfr,RhoGEF2,Rho1,ci* |
| imaginal disc-derived wing vein morphogenesis | GO:BP | 2.46e-04 | *dally,cv-c,Dys,vn,Egfr,shn,Hipk,spen,Ras85D,gbb* |
| imaginal disc-derived wing vein specification | GO:BP | 2.95e-04 | *cv-2,N,aos,Dg,Mkp3,Dys,cic,Shc,Egfr,Ras85D,gbb* |
| wing disc anterior/posterior pattern formation | GO:BP | 3.19e-04 | *dally,Sulf1,fng,ptc,shn,ci* |
| vacuole organization | GO:BP | 3.43e-04 | *Atg17,Atg8a,Atg9,Atg18b,Rab1,Rab7,Atg18a,Rab14,Lamtor1,Svip,Atg3* |
| tracheal outgrowth, open tracheal system | GO:BP | 4.59e-04 | *ttk,stumps,Shc,Egfr,Mmp2,Ras85D* |
| regulation of RNA metabolic process | GO:BP | 5.28e-04 | *sens-2,CG14431,CG9650,bab2,Atf3,Antp,tsh,N,Hand,edl,Eip78C,bru2,ken,Tet,klu,E(spl)m3-HLH,lin-28,luna,E(spl)mbeta-HLH,gcl,noc,kibra,aop,zfh1,jigr1,Glut4EF,sbb,eve,how,chinmo,AGO1,Sox14,REPTOR-BP,hth,ttk,Trf2,REPTOR,tup,CG8312,cic,Tob,Pur-alpha,Mes2,slbo,Mnn1,tna,AGO2,shn,ps,CG6770,CG12384,sd,CG10543,Pits,Hr4,bun,Smr,spen,Taf3,Mnt,maf-S,ci,Not1,Psc,PAN3,crol,dmpd,Sirt1,simj,Crtc,ova,Adf1,ko,ftz-f1,Ubc6,elB,Rtf1,Max,DnaJ-1,Daxx,Axud1,CG7879,Clamp,rgr,CG12099* |
| wing disc dorsal/ventral pattern formation | GO:BP | 5.67e-04 | *dally,drpr,N,Sulf1,fng,sbb,cic,shn,Rab7,CG1943* |
| cellular response to growth factor stimulus | GO:BP | 6.19e-04 | *rau,cv-2,dally,htl,Toll-7,Ptp4E,aop,stumps,PRAS40,Shc,shn,Mmp2,pbl,Ras85D,gbb,lili* |
| myoblast fate specification | GO:BP | 6.78e-04 | *htl,ttk,Ras85D* |
| nerve maturation | GO:BP | 6.78e-04 | *Nrg,Cont,Nrx-IV* |
| positive regulation of filopodium assembly | GO:BP | 7.23e-04 | *Mipp1,DAAM,pico,Flo2,aPKC,Abi,Cyfip* |
| cell fate commitment involved in pattern specification | GO:BP | 7.89e-04 | *N,klu,tup,dlg1,numb,vn,l(2)gl,Egfr* |
| glial cell migration | GO:BP | 9.46e-04 | *Pvf3,N,htl,how,unc-5,numb,orion,Rho1* |
| peripheral nervous system development | GO:BP | 9.77e-04 | *N,hth,ttk,Dscam1,spi,vn,Egfr,Rho1,spen,Ras85D* |
| positive regulation of TORC1 signaling | GO:BP | 0.001 | *hppy,Shc,Rheb,Ras85D,Lamtor1,Akt,Lnk* |
| homophilic cell adhesion via plasma membrane adhesion molecules | GO:BP | 0.001 | *CG13506,Lac,Fas2,Cont,Dscam1,Fas1,ed,sdk,Cad87A* |
| regulation of potassium ion transmembrane transport | GO:BP | 0.001 | *CG9338,crim,crok,CG9336,Neto,CG6583* |
| regulation of synaptic transmission, cholinergic | GO:BP | 0.001 | *CG9338,crim,crok,CG9336,CG6583,bou* |
| negative regulation of biosynthetic process | GO:BP | 0.002 | *Antp,tsh,N,edl,Eip78C,bru2,klu,lin-28,CG3662,E(spl)mbeta-HLH,gcl,noc,aop,zfh1,sbb,eve,AGO1,ttk,cic,Tob,numb,Edc3,spi,Pink1,Egfr,AGO2,shn,CG12384,cer,sd,Pits,Hr4,Smr,Ras85D,Mnt,CG6701,ci,Not1,Psc,PAN3,crol,Sirt1,simj,msi,Ubc6,elB,Max,Hmt4-20,DnaJ-1,Smg5,Daxx,Hsp83* |
| muscle cell development | GO:BP | 0.002 | *rols,htl,Dg,DAAM,aop,Dys,how,tmod,sd,dia* |
| establishment of mitotic spindle orientation | GO:BP | 0.002 | *cv-c,dlg1,nudE,l(2)gl,aPKC,Slik* |
| female germ-line stem cell population maintenance | GO:BP | 0.002 | *N,InR,fng,AGO1,fax,gbb,lili* |
| positive regulation of cell size | GO:BP | 0.002 | *Pvf3,hppy,Pdk1,InR,Rheb,Ras85D,Akt,MAPk-Ak2* |
| positive regulation of ERK1 and ERK2 cascade | GO:BP | 0.002 | *spg,spi,vn,Egfr,Ras85D,Lnk* |
| dorsal appendage formation | GO:BP | 0.003 | *N,Pax,ttk,cic,ed,jvl,Egfr,bun,RanBPM* |
| behavioral response to ethanol | GO:BP | 0.003 | *Pka-R2,hppy,Fas2,dlg1,mura,spi,Akap200,Egfr,Sirt1* |
| negative regulation of cell population proliferation | GO:BP | 0.003 | *kibra,Xrp1,numb,l(2)gl,Mnn1,CG6770,Rab5,sd,Mmp2,Axud1,Hsp83* |
| positive regulation of JNK cascade | GO:BP | 0.003 | *hppy,Mnn1,Tnks,Atg9,Alg-2,ALiX,Cka* |
| positive regulation of voltage-gated potassium channel activity | GO:BP | 0.003 | *CG9338,crim,crok,CG9336,CG6583* |
| pseudocleavage involved in syncytial blastoderm formation | GO:BP | 0.004 | *Eb1,RhoGEF2,Rho1,dia* |
| positive regulation of actin filament polymerization | GO:BP | 0.004 | *DAAM,chinmo,LRR,IRSp53,dia* |
| peptidyl-threonine phosphorylation | GO:BP | 0.004 | *Pdk1,Dyrk2,Hipk,Dyrk3,Slik* |
| catabolic process | GO:BP | 0.004 | *N,CG4829,Pax,CG11658,Dg,lin-28,Toll-7,Rnf11,gcl,InR,Atg17,CG9003,bwa,rudhira,AGO1,Usp12-46,Oda,Kaz,Edc3,Klp98A,Atg8a,Tnks,shrb,Cht2,Rheb,Pink1,gzl,CG12402,Ppa,AGO2,UbcE2H,CG12384,CG5445,Atg9,Mmp2,Atg18b,Rab1,Ufd4,CG4911,Rab7,CG7611,ALiX,Atg18a,ref(2)P,Rab18,Ras85D,Ubi-p5E,Ubr1,Not1,Ubi-p63E,PAN3,Vti1b,Snap29,Vps37B,dmpd,CG5961,RanBPM,CtsF,Bruce,Ubc6,Vps28,Svip,bmm,Akt,RNaseX25,Smg5,Plc21C,Atg3,Psa,ClpX,sordd1,AMPKalpha,CG17691* |
| notum cell fate specification | GO:BP | 0.004 | *tup,vn,Egfr* |
| muscle cell fate determination | GO:BP | 0.004 | *N,how,tup* |
| positive regulation of hippo signaling | GO:BP | 0.004 | *hppy,kibra,ptc,ed,l(2)gl,Hipk,alpha-Spec* |
| peptidyl-serine phosphorylation | GO:BP | 0.004 | *Pdk1,Lk6,Pink1,Dyrk2,Hipk,aPKC,Dyrk3,Akt,MAPk-Ak2* |
| biological process involved in interspecies interaction between organisms | GO:BP | 0.005 | *drpr,FER,N,htl,GILT2,GILT1,Toll-7,18w,Gprk2,Eb1,Nrg,Fer2LCH,dlg1,Dscam1,Rab2,shrb,Shc,Fer1HCH,Flo2,AGO2,sick,Rab5,Mmp2,Rab1,Rab8,Rab7,NUCB1,Atg18a,Rho1,spen,Vps60,Ras85D,wun,Rab14,Pitslre,dia,TSG101,aPKC,Hlc,Hsp27* |
| germarium-derived oocyte fate determination | GO:BP | 0.005 | *BicD,AGO1,nudE,par-1,alpha-Spec,aPKC* |
| glycophagy | GO:BP | 0.005 | *Atg17,Atg9,Atg18a,Atg3* |
| maintenance of presynaptic active zone structure | GO:BP | 0.005 | *Dys,Dyb,beta-Spec,alpha-Spec* |
| trans-synaptic signaling | GO:BP | 0.005 | *wgn,CG9338,crim,crok,CG9336,Fas2,cv-c,Dg,Neto,BicD,Dys,GluRIID,CG6583,Dyb,dlg1,Sap47,beta-Spec,Pink1,scramb2,alpha-Spec,Snap29,gbb,mth,Tomosyn,Akt,bou,Plc21C* |
| positive regulation of insulin receptor signaling pathway | GO:BP | 0.006 | *lin-28,CycG,Mob4,Mkrn1,Hsp83* |
| negative regulation of cell adhesion | GO:BP | 0.006 | *N,aos,aop,par-1,RanBPM* |
| gravitaxis | GO:BP | 0.006 | *w,dlg1,l(3)L1231,CG10543,Hipk,Cka* |
| regulation of tube length, open tracheal system | GO:BP | 0.006 | *Fas2,pasi2,pasi1,kune,Egfr,Rho1* |
| spindle localization | GO:BP | 0.006 | *cv-c,Eb1,dlg1,nudE,l(2)gl,aPKC,Slik* |
| striated muscle cell differentiation | GO:BP | 0.007 | *rols,Pax,Dg,DAAM,eve,how,tmod,sd,Rho1,blow,dia* |
| mushroom body development | GO:BP | 0.007 | *robo2,Fas2,DAAM,Nrg,chinmo,Dscam1,orion,myo,bun,ftz-f1* |
| stem cell fate commitment | GO:BP | 0.007 | *spi,Egfr,Ras85D* |
| protein localization to phagophore assembly site | GO:BP | 0.007 | *Atg9,Atg18b,Atg18a* |
| photoreceptor cell fate determination | GO:BP | 0.007 | *spi,Egfr,Ras85D* |
| inositol phosphate biosynthetic process | GO:BP | 0.007 | *IP3K1,Ip6k,IP3K2* |
| positive regulation of Notch signaling pathway | GO:BP | 0.007 | *fng,ed,Akap200,Mnr,Ehbp1,Hipk,Sirt1* |
| negative regulation of JNK cascade | GO:BP | 0.008 | *Mnn1,ics,stck,Cka,fmt,raw* |
| positive regulation of organ growth | GO:BP | 0.009 | *Pdk1,InR,Hipk,Akt* |
| maintenance of epithelial integrity, open tracheal system | GO:BP | 0.009 | *Lac,cv-c,Mkp3,Egfr* |
| detoxification of iron ion | GO:BP | 0.009 | *Fer2LCH,Fer1HCH* |
| specification of segmental identity, antennal segment | GO:BP | 0.009 | *Antp,hth* |
| long-term strengthening of neuromuscular junction | GO:BP | 0.009 | *beta-Spec,alpha-Spec* |
| regulation of polarized epithelial cell differentiation | GO:BP | 0.009 | *par-1,aPKC* |
| negative regulation of fusion cell fate specification | GO:BP | 0.009 | *aop,ed* |
| oenocyte delamination | GO:BP | 0.009 | *aos,aop* |
| iron ion import across plasma membrane | GO:BP | 0.009 | *Fer2LCH,Fer1HCH* |
| positive regulation of cell proliferation involved in compound eye morphogenesis | GO:BP | 0.009 | *tsh,hth* |
| germ cell repulsion | GO:BP | 0.009 | *wun2,wun* |
| ommatidial rotation | GO:BP | 0.009 | *N,aos,spi,Egfr,Ras85D,ec* |
| negative regulation of ERK1 and ERK2 cascade | GO:BP | 0.01 | *aos,Mkp3,Spred,Ufd4,Ubc6* |
| melanotic encapsulation of foreign target | GO:BP | 0.01 | *Eb1,Nrg,Shc,Flo2,dia,aPKC* |
| positive regulation of wound healing | GO:BP | 0.011 | *InR,Egfr,Rho1,dia* |
| behavioral response to starvation | GO:BP | 0.011 | *Galphao,AMPKalpha,Hsp27* |
| segment polarity determination | GO:BP | 0.012 | *dally,AGO1,ptc,nkd,Egfr,AGO2,ci* |
| negative regulation of hippo signaling | GO:BP | 0.012 | *par-1,Hipk,Cka,ci,aPKC,Akt,Mob4* |
| negative regulation of canonical Wnt signaling pathway | GO:BP | 0.012 | *Sulf1,aop,nkd,Mmp2,Rho1,pbl,Duba* |
| cholesterol homeostasis | GO:BP | 0.014 | *InR,Rheb,Lamtor1,Akt* |
| positive regulation of lipid storage | GO:BP | 0.014 | *InR,elm,bmm,Akt* |
| ventral cord development | GO:BP | 0.016 | *Antp,htl,robo2,18w,Galphao,Dscam1,Mmp2,Psc,gbb* |
| negative regulation of peptide hormone secretion | GO:BP | 0.016 | *CG9650,InR,Akt* |
| cold acclimation | GO:BP | 0.016 | *Hsp23,Hsp26,Hsp83* |
| dorsal closure, spreading of leading edge cells | GO:BP | 0.016 | *Egfr,Rho1,Ras85D* |
| larval midgut cell programmed cell death | GO:BP | 0.016 | *Atg17,Atg8a,Atg9,Atg18a,Ubi-p63E* |
| leg disc proximal/distal pattern formation | GO:BP | 0.017 | *tsh,vn,Egfr,Ras85D* |
| female germ-line stem cell asymmetric division | GO:BP | 0.017 | *dally,InR,shrb,ALiX* |
| negative regulation of secretion by cell | GO:BP | 0.017 | *CG9650,InR,Snap29,Akt* |
| eye-antennal disc morphogenesis | GO:BP | 0.02 | *bab2,N,aos,ptc,Egfr,Ras85D* |
| head involution | GO:BP | 0.02 | *tsh,cv-c,scyl,chrb,tup,stck* |
| regulation of monoatomic ion transmembrane transport | GO:BP | 0.021 | *CG9338,crim,crok,CG9336,Neto,CG6583,elm,Stim* |
| imaginal disc-derived male genitalia morphogenesis | GO:BP | 0.021 | *Myo31DF,N,Fas2,Rab5* |
| multivesicular body assembly | GO:BP | 0.021 | *Rab5,TSG101* |
| regulation of glycolytic process | GO:BP | 0.021 | *N,Dg* |
| mitotic actomyosin contractile ring assembly | GO:BP | 0.021 | *Rho1,pbl* |
| positive regulation of mitotic cell cycle | GO:BP | 0.021 | *N,stg,ci,PNUTS,crol,Abi* |
| neuroblast fate specification | GO:BP | 0.021 | *N,nkd* |
| glial cell fate determination | GO:BP | 0.021 | *N,spen* |
| maintenance of imaginal disc-derived wing hair orientation | GO:BP | 0.021 | *cora,spen* |
| specification of segmental identity, thorax | GO:BP | 0.021 | *Antp,tsh* |
| dendrite regeneration | GO:BP | 0.021 | *ttk,Akt* |
| positive regulation of integrin-mediated signaling pathway | GO:BP | 0.021 | *Mp,stck* |
| regulation of actomyosin contractile ring contraction | GO:BP | 0.021 | *RhoGEF2,Graf* |
| intracellular sequestering of iron ion | GO:BP | 0.021 | *Fer2LCH,Fer1HCH* |
| lateral inhibition | GO:BP | 0.021 | *N,Mmp2,Sirt1* |
| extracellular vesicle biogenesis | GO:BP | 0.021 | *Rab7,ALiX,Hsp83* |
| determination of genital disc primordium | GO:BP | 0.021 | *ptc,spi,Egfr* |
| positive regulation of catalytic activity | GO:BP | 0.022 | *RhoGAP15B,htl,InR,Atg17,spg,Ziz,Tbc1d8-9,Rho1,pbl,Cka,RalGPS,Asap,Abi,CG5521,aph-1* |
| negative regulation of Notch signaling pathway | GO:BP | 0.024 | *fng,E(spl)m6-BFM,CycG,Usp12-46,numb,l(2)gl* |
| terminal branching, open tracheal system | GO:BP | 0.024 | *noc,Rheb,Ras85D,TSG101,aPKC* |
| positive regulation of phosphatidylinositol 3-kinase/protein kinase B signal transduction | GO:BP | 0.024 | *Pdk1,InR,CycG,Lnk* |
| positive regulation of synaptic assembly at neuromuscular junction | GO:BP | 0.026 | *htl,dlg1,Rheb,RhoGAP68F,gbb,simj* |
| positive regulation of transcription by RNA polymerase II | GO:BP | 0.027 | *Antp,tsh,N,Hand,Tet,eve,Sox14,hth,ttk,tup,CG8312,slbo,shn,sd,CG10543,Taf3,maf-S,ci,Crtc,Adf1,ko,ftz-f1,Rtf1,Max,Axud1,Clamp,rgr,CG12099* |
| tracheal pit formation in open tracheal system | GO:BP | 0.028 | *cv-c,Mmp2,Rho1* |
| neuroblast development | GO:BP | 0.029 | *Antp,N,klu,numb* |
| asymmetric protein localization involved in cell fate determination | GO:BP | 0.029 | *dlg1,Klp98A,l(2)gl,aPKC* |
| germ-band shortening | GO:BP | 0.029 | *InR,tup,Egfr,raw* |
| apposition of dorsal and ventral imaginal disc-derived wing surfaces | GO:BP | 0.029 | *how,ics,stck,Ilk* |
| R8 cell fate specification | GO:BP | 0.034 | *kibra,l(2)gl,myo,gbb* |
| dendrite self-avoidance | GO:BP | 0.034 | *Fas2,Cont,Dscam1,Rho1* |
| protein refolding | GO:BP | 0.035 | *CG14207,Hsp70Ab,Hsp23,Hsp27,Hsp26* |
| regulation of myoblast fusion | GO:BP | 0.035 | *Pax,eve,Rho1* |
| epithelial cell proliferation involved in Malpighian tubule morphogenesis | GO:BP | 0.035 | *N,spi,Egfr* |
| regulation of stem cell division | GO:BP | 0.035 | *N,klu,chinmo* |
| germ-band extension | GO:BP | 0.035 | *eve,RhoGEF2,Rho1* |
| regulation of basement membrane organization | GO:BP | 0.036 | *Ndg,Ehbp1* |
| regulation of telomere maintenance | GO:BP | 0.036 | *Tnks,PNUTS* |
| posterior midgut invagination | GO:BP | 0.036 | *RhoGEF2,Rho1* |
| endosome organization | GO:BP | 0.039 | *Vps24,Rab5,ref(2)P,TSG101,Vps28* |
| germ-line stem-cell niche homeostasis | GO:BP | 0.039 | *N,lin-28,InR,RanBPM* |
| negative regulation of actin filament polymerization | GO:BP | 0.039 | *LRR,beta-Spec,tmod,alpha-Spec* |
| genital disc morphogenesis | GO:BP | 0.039 | *Myo31DF,N,Fas2,Rab5* |
| chaeta morphogenesis | GO:BP | 0.041 | *dally,N,unk,tup,Ype,spen* |
| muscle cell cellular homeostasis | GO:BP | 0.043 | *N,Dg,zfh1,Dys,Dyb* |
| ectopic germ cell programmed cell death | GO:BP | 0.044 | *Pink1,wun2,wun* |
| meiosis II cytokinesis | GO:BP | 0.044 | *fwd,pbl,dia* |
| cell projection assembly | GO:BP | 0.044 | *Mipp1,N,DAAM,Unc-115a,LRR,vn,pnut,pico,Egfr,Flo2,IRSp53,Efa6,Rho1,dia,aPKC,mtm,Abi,Cyfip* |
| negative regulation of cell size | GO:BP | 0.045 | *Sema1a,stg,pbl,AMPKalpha* |
| sensory organ precursor cell fate determination | GO:BP | 0.045 | *N,klu,numb,l(2)gl* |
| negative regulation of transcription by RNA polymerase II | GO:BP | 0.047 | *Antp,tsh,klu,E(spl)mbeta-HLH,aop,zfh1,sbb,eve,ttk,cic,shn,Smr,Mnt,ci,Psc,simj,Max,DnaJ-1* |
| midgut development | GO:BP | 0.048 | *Antp,tsh,cv-c,vn,shn* |
| germ cell migration | GO:BP | 0.048 | *htl,zfh1,wun2,stai,wun* |
| post-translational protein modification | GO:BP | 0.049 | *CG11658,Rnf11,gcl,Usp12-46,mura,Atg8a,Tnks,tna,gzl,UbcE2H,Ufd4,CG4911,CG7222,Ubi-p5E,Ubr1,Ubi-p63E,ec,CG5961,CG17754,Bruce,Ubc6,sordd1,Duba,Mkrn1,CG12099* |
| positive regulation of cell growth | GO:BP | 0.049 | *Pdk1,InR,Rheb,bun,Akt,Lnk* |
| protein binding | GO:MF | 1.74e-24 | *CG14431,rau,cv-2,Pvf3,Fili,CG13506,tei,Lac,Pka-R2,bab2,Atf3,Myo31DF,dally,Antp,tsh,rols,LRP1,drpr,FER,wgn,N,htl,unk,Hand,edl,CG11658,aos,robo2,CG10011,ken,BNIP3,Fas2,E(spl)m3-HLH,Dg,Toll-7,E(spl)mbeta-HLH,Chd64,Fim,gcl,Kal1,18w,Neto,noc,InR,BicD,Atg17,slim,DAAM,CG42748,kibra,mlt,sinu,CG9003,Ptp4E,CG14995,aop,jigr1,Dys,cib,side-IV,udd,CG42788,Eb1,Sema1a,CG7702,CG14696,E(spl)m6-BFM,Nrg,sbb,chinmo,AGO1,fax,CycG,Lk6,spg,Cont,REPTOR-BP,ptc,Unc-115a,Lrch,hth,ttk,Trf2,cora,LRR,REPTOR,tup,dlg1,unc-5,Galphao,Dscam1,pns,dos,Fas1,CG11882,CG13124,Oatp74D,Kaz,cic,Rab2,Tob,numb,Pur-alpha,Sh3beta,Wdr62,CG42709,Edc3,spi,nudE,CG32066,stumps,slbo,Klp98A,ed,beta-Spec,bif,vn,Atg8a,pnut,l(2)gl,jvl,Tnks,CG45050,Akap200,nkd,shrb,Shc,pico,Plp,Rip11,Con,Rheb,Pink1,Egfr,Meltrin,orion,CG12402,Ppa,AGO2,Rab4,shn,Spred,CG12384,myo,CG5445,stai,tmod,IRSp53,Mhcl,GMF,sick,sdk,Rab5,sd,par-1,Atg18b,Alg-2,ics,NKAIN,Efa6,Rab1,Ehbp1,Sema2a,LTV1,bun,Ufd4,CG4911,Rab8,RhoGEF2,CG7611,ALiX,klar,Smr,Hipk,Atg18a,stck,alpha-Spec,Rho1,mol,ref(2)P,Taf3,Rab18,Hil,pbl,sbr,CG1888,Cka,Ilk,Ras85D,Ubi-p5E,Mnt,miple2,CG4393,CG3402,CG30423,maf-S,blow,RhoGAP68F,ci,Not1,dia,Ubi-p63E,TfIIA-S,TSG101,aPKC,Fs(2)Ket,PNUTS,eIF3l,cta,PAN3,CG32369,Vti1b,fmt,ec,Snap29,gbb,CG1105,Cdep,dmpd,CG5961,RanBPM,Sirt1,CG40228,spri,simj,Crtc,CDK2AP1,olf186-M,CG17754,Bruce,CG14207,ftz-f1,Asap,mtm,Hsp70Ab,Ubc6,Dlic,Pcf11,CG5721,Pde11,elB,Vps28,mth,Abi,Rtf1,Max,Hmt4-20,Tomosyn,Tpr2,Akt,Lnk,DnaJ-1,Smg5,CG2765,Vinc,Cyfip,Stim,Ras64B,Daxx,ClpX,Faf2,sordd1,sowah,Graf,Hsp23,MAPk-Ak2,Hsp27,IP3K2,Hsp26,Hsp83* |
| enzyme binding | GO:MF | 1.72e-07 | *CG14431,rau,Pka-R2,N,BicD,Atg17,DAAM,CG42748,kibra,CG14696,E(spl)m6-BFM,AGO1,Lk6,spg,dlg1,pns,CG32066,stumps,Atg8a,pnut,l(2)gl,Shc,pico,Rip11,Spred,Rho1,ref(2)P,pbl,Cka,Ubi-p5E,RhoGAP68F,ci,dia,Ubi-p63E,Fs(2)Ket,PNUTS,fmt,ec,CG1105,RanBPM,CG40228,spri,CDK2AP1,Pcf11,Rtf1,Lnk,Cyfip,sordd1,MAPk-Ak2* |
| phospholipid binding | GO:MF | 1.35e-04 | *Myo31DF,drpr,FER,RhoGAP15B,CG9338,crim,crok,CG9336,Gprk2,CG6583,ptc,Klp98A,beta-Spec,IRSp53,Atg18b,Efa6,Atg18a,pbl,dia,Asap,CG1902,Graf* |
| GTPase regulator activity | GO:MF | 1.35e-04 | *RhoGAP15B,cv-c,RhoGAP71E,spg,pns,Ziz,l(2)gl,Tbc1d8-9,Efa6,RhoGEF2,raskol,pbl,RhoGAP68F,cta,Cdep,RalGPS,spri,Asap,Lamtor1,Pura,CG5521,Tomosyn,Graf* |
| protein domain specific binding | GO:MF | 3.48e-04 | *N,InR,aop,Dys,ttk,dos,Sh3beta,nkd,pico,CG12384,sd,Taf3,Ubc6,Abi,Hsp83* |
| kinase binding | GO:MF | 3.48e-04 | *Pka-R2,Atg17,kibra,Lk6,dlg1,stumps,Atg8a,l(2)gl,Shc,Spred,Rho1,ref(2)P,Cka,ci,spri,Lnk,MAPk-Ak2* |
| phosphatidylinositol binding | GO:MF | 3.48e-04 | *Myo31DF,RhoGAP15B,CG9338,crim,crok,CG9336,Gprk2,CG6583,ptc,Klp98A,beta-Spec,Atg18b,Atg18a,pbl,dia,Asap,CG1902* |
| protein kinase binding | GO:MF | 4.28e-04 | *Pka-R2,Atg17,Lk6,stumps,Atg8a,l(2)gl,Shc,Spred,Rho1,ref(2)P,Cka,ci,spri,Lnk,MAPk-Ak2* |
| cytoskeletal protein binding | GO:MF | 6.72e-04 | *Myo31DF,Chd64,Fim,BicD,DAAM,mlt,Dys,cib,Eb1,Unc-115a,cora,nudE,Klp98A,beta-Spec,bif,pnut,l(2)gl,Akap200,stai,tmod,Mhcl,GMF,Efa6,klar,alpha-Spec,Rho1,dia,aPKC,Cdep,CG17754,Tomosyn,CG2765,Vinc,Hsp26* |
| transcription regulator activity | GO:MF | 7.49e-04 | *sens-2,CG14431,CG9650,bab2,Atf3,Antp,tsh,N,Pax,Hand,edl,Eip78C,ken,klu,E(spl)m3-HLH,luna,E(spl)mbeta-HLH,aop,zfh1,Glut4EF,sbb,eve,Sox14,REPTOR-BP,hth,ttk,Trf2,REPTOR,tup,cic,Tob,Pur-alpha,slbo,tna,shn,sd,CG10543,Pits,Hr4,Smr,Mnt,maf-S,ci,crol,dmpd,Sirt1,Crtc,ova,Adf1,ko,ftz-f1,Max,Daxx,Axud1,Clamp,rgr* |
| epidermal growth factor receptor binding | GO:MF | 7.49e-04 | *aos,spi,ed,vn,Shc* |
| actin binding | GO:MF | 7.88e-04 | *Myo31DF,Chd64,Fim,DAAM,Dys,cib,Unc-115a,cora,beta-Spec,bif,pnut,Akap200,Mhcl,GMF,alpha-Spec,Rho1,dia,CG17754,Vinc* |
| DNA-binding transcription factor activity | GO:MF | 8.07e-04 | *sens-2,CG14431,CG9650,bab2,Atf3,Antp,tsh,Hand,edl,Eip78C,ken,klu,E(spl)m3-HLH,luna,E(spl)mbeta-HLH,aop,zfh1,Glut4EF,eve,Sox14,REPTOR-BP,hth,ttk,Trf2,REPTOR,tup,cic,Pur-alpha,slbo,shn,sd,CG10543,Hr4,Mnt,maf-S,ci,crol,ova,Adf1,ko,ftz-f1,Max,Axud1,Clamp,rgr* |
| protein tyrosine kinase activity | GO:MF | 8.07e-04 | *FER,htl,cdi,InR,Egfr,Dyrk2,par-1,Hipk,Pitslre,Dyrk3* |
| signaling receptor binding | GO:MF | 8.18e-04 | *cv-2,Pvf3,FER,aos,Sema1a,ptc,Galphao,numb,spi,stumps,ed,vn,Atg8a,Akap200,Shc,orion,myo,Sema2a,RhoGEF2,pbl,miple2,CG30423,cta,gbb,spri,olf186-M,Lnk,Hsp83* |
| growth factor receptor binding | GO:MF | 0.002 | *Pvf3,aos,spi,stumps,ed,vn,Shc* |
| small GTPase binding | GO:MF | 0.002 | *CG14431,rau,BicD,DAAM,spg,pns,CG32066,Rip11,pbl,RhoGAP68F,dia,Fs(2)Ket,RanBPM,spri,Cyfip* |
| sequence-specific DNA binding | GO:MF | 0.002 | *sens-2,CG14431,CG9650,Atf3,Antp,Hand,edl,Eip78C,ken,klu,E(spl)m3-HLH,luna,E(spl)mbeta-HLH,Chd64,aop,zfh1,jigr1,Glut4EF,eve,Sox14,hth,ttk,Trf2,REPTOR,tup,Xrp1,cic,Pur-alpha,slbo,Mnn1,shn,sd,CG10543,Hr4,Mnt,maf-S,ci,Psc,crol,ova,Adf1,ko,ftz-f1,Max,Smg5,Axud1,Clamp,CG12099* |
| sequence-specific double-stranded DNA binding | GO:MF | 0.002 | *sens-2,CG14431,CG9650,Atf3,Antp,Hand,Eip78C,ken,klu,E(spl)m3-HLH,luna,E(spl)mbeta-HLH,Chd64,aop,zfh1,jigr1,Glut4EF,eve,Sox14,hth,ttk,Trf2,REPTOR,tup,Xrp1,cic,Pur-alpha,slbo,Mnn1,shn,sd,CG10543,Hr4,Mnt,maf-S,ci,crol,ova,Adf1,ko,ftz-f1,Max,Clamp,CG12099* |
| GTPase binding | GO:MF | 0.002 | *CG14431,rau,BicD,DAAM,spg,pns,CG32066,Rip11,pbl,RhoGAP68F,dia,Fs(2)Ket,RanBPM,spri,Cyfip* |
| phosphatidylinositol phosphate binding | GO:MF | 0.002 | *Myo31DF,RhoGAP15B,Gprk2,ptc,Klp98A,beta-Spec,Atg18b,Atg18a,pbl,dia,Asap,CG1902* |
| transcription cis-regulatory region binding | GO:MF | 0.003 | *sens-2,CG14431,CG9650,Atf3,Antp,Hand,Eip78C,ken,klu,E(spl)m3-HLH,luna,E(spl)mbeta-HLH,Chd64,aop,zfh1,jigr1,Glut4EF,eve,Sox14,hth,ttk,Trf2,REPTOR,tup,cic,Pur-alpha,slbo,Mnn1,shn,sd,CG10543,Hr4,Mnt,maf-S,ci,crol,ova,Adf1,ko,ftz-f1,Clamp,CG12099* |
| GTPase activator activity | GO:MF | 0.003 | *RhoGAP15B,cv-c,RhoGAP71E,l(2)gl,Tbc1d8-9,raskol,pbl,RhoGAP68F,spri,Asap,CG5521,Tomosyn,Graf* |
| GTPase activity | GO:MF | 0.004 | *Galphao,Rab2,pnut,Rheb,Rab4,Rab5,Rab1,Non1,Rab8,Rab7,Rho1,Rab18,Ras85D,Rab14,cta,CG2017,Ras64B* |
| structural constituent of muscle | GO:MF | 0.004 | *rols,Dg,Dys,Tina-1,Dyb* |
| GPI anchor binding | GO:MF | 0.004 | *CG9338,crim,crok,CG9336,CG6583* |
| RNA polymerase II transcription regulatory region sequence-specific DNA binding | GO:MF | 0.004 | *sens-2,CG14431,CG9650,Atf3,Antp,Hand,Eip78C,ken,klu,E(spl)m3-HLH,luna,E(spl)mbeta-HLH,Chd64,aop,zfh1,Glut4EF,eve,Sox14,hth,ttk,Trf2,REPTOR,tup,cic,Pur-alpha,slbo,shn,sd,CG10543,Hr4,Mnt,maf-S,ci,crol,ova,ko,ftz-f1,Clamp* |
| cell adhesion molecule binding | GO:MF | 0.004 | *tei,rols,Fas2,Dg,Nrg,Cont,Dscam1,Fas1,ed,sdk,Abi* |
| double-stranded DNA binding | GO:MF | 0.004 | *sens-2,CG14431,CG9650,Atf3,Antp,Hand,Eip78C,ken,klu,E(spl)m3-HLH,luna,E(spl)mbeta-HLH,Chd64,aop,zfh1,jigr1,Glut4EF,eve,Sox14,hth,ttk,Trf2,REPTOR,tup,Xrp1,cic,Pur-alpha,slbo,Mnn1,shn,sd,CG10543,Hr4,CG5316,Mnt,maf-S,ci,crol,ova,Adf1,ko,ftz-f1,Max,Clamp,CG12099* |
| lipid binding | GO:MF | 0.005 | *Myo31DF,drpr,FER,RhoGAP15B,CG9338,crim,crok,CG9336,cv-c,Gprk2,CG6583,ptc,Klp98A,beta-Spec,IRSp53,Atg18b,Efa6,Atg18a,pbl,dia,Asap,CG1902,Graf* |
| inositol hexakisphosphate kinase activity | GO:MF | 0.005 | *IP3K1,Ip6k,IP3K2* |
| G protein activity | GO:MF | 0.006 | *Rab4,Ras85D,cta,Ras64B* |
| DNA-binding transcription factor activity, RNA polymerase II-specific | GO:MF | 0.008 | *sens-2,CG14431,Atf3,Antp,tsh,Hand,edl,Eip78C,klu,E(spl)m3-HLH,luna,E(spl)mbeta-HLH,aop,zfh1,eve,Sox14,hth,ttk,REPTOR,tup,cic,Pur-alpha,slbo,shn,sd,CG10543,Hr4,Mnt,maf-S,ci,crol,ova,Adf1,ko,ftz-f1,Axud1,Clamp* |
| cell adhesion mediator activity | GO:MF | 0.009 | *Fas2,Dg,Cont,Dscam1,sdk* |
| receptor tyrosine kinase binding | GO:MF | 0.009 | *stumps,Shc,spri,Lnk* |
| guanyl-nucleotide exchange factor activity | GO:MF | 0.016 | *spg,pns,Ziz,Efa6,RhoGEF2,pbl,Cdep,RalGPS,spri,Lamtor1,Pura* |
| small molecule binding | GO:MF | 0.016 | *Nep4,AdamTS-A,CG9650,Pka-R2,Ndg,hppy,Myo31DF,tsh,rols,LRP1,CG17646,FER,N,RhoGAP15B,htl,Pax,unk,Eip78C,cdi,w,ken,Sulf1,Tet,Dg,lin-28,CG34445,luna,Rnf11,Fim,CG3164,Pdk1,CG32486,noc,InR,fng,zfh1,Gprk2,Dys,Glut4EF,CG10737,bwa,Nrg,chinmo,Lk6,Unc-115a,Dyb,Atet,ttk,Fer2LCH,tup,Galphao,Dscam1,mura,Kaz,Rab2,numb,Klp98A,beta-Spec,Ypel,elm,pnut,rho-4,Tnks,CG2991,CG45050,nkd,Tbc1d8-9,Rheb,Pink1,Egfr,Meltrin,Fer1HCH,tna,gzl,AGO2,Dyrk2,UbcE2H,Rab4,shn,Mhcl,sick,Rab5,sd,par-1,Mmp2,Atg18b,Alg-2,CG10543,Rab1,Dph4,Hr4,Ufd4,Non1,CG33298,Rab8,Rab7,RhoGEF2,NUCB1,CG32280,Hipk,Atg18a,stck,CG5316,alpha-Spec,Rho1,Ype,ref(2)P,Taf3,Rab18,Hil,CG30195,Ilk,Ras85D,Ubr1,Rab14,ci,Pitslre,Act87E,dia,Psc,aPKC,PNUTS,cta,PAN3,CG32369,crol,Sirt1,CG40228,Dyrk3,Slik,Cad87A,CG14207,ftz-f1,Asap,Hlc,Hsp70Ab,Ubc6,Dlic,Pde11,elB,CG2017,IscU,Akt,CG10803,Mob4,Stim,CG3224,Plc21C,Ras64B,Psa,ClpX,sordd1,AMPKalpha,Mkrn1,Clamp,CG12099,MAPk-Ak2,Hsp83* |
| protein kinase activity | GO:MF | 0.018 | *hppy,FER,htl,cdi,Pdk1,InR,Gprk2,Lk6,Pink1,Egfr,Dyrk2,par-1,Hipk,Ilk,Pitslre,aPKC,Dyrk3,Slik,Akt,AMPKalpha,MAPk-Ak2* |
| actin filament binding | GO:MF | 0.018 | *Myo31DF,Chd64,Fim,Dys,Unc-115a,beta-Spec,bif,Akap200,Mhcl,alpha-Spec,Vinc* |
| enzyme regulator activity | GO:MF | 0.018 | *Pka-R2,RhoGAP15B,CG43366,cv-c,Kal1,Pdk1,Atg17,RhoGAP71E,CycG,spg,pns,Oda,Ziz,spi,l(2)gl,Tbc1d8-9,cer,Rab5,Efa6,RhoGEF2,raskol,pbl,Ras85D,RhoGAP68F,cta,fmt,Cdep,RalGPS,spri,Bruce,Asap,Lamtor1,Pura,CG5521,Tomosyn,Graf* |
| kinase activity | GO:MF | 0.018 | *hppy,FER,htl,cdi,IP3K1,Pdk1,Mkp3,InR,Gprk2,Lk6,dlg1,Akap200,Ip6k,Pink1,Egfr,Dyrk2,par-1,Hipk,fwd,Cka,Ilk,Pitslre,aPKC,Dyrk3,Slik,Abi,Akt,Mob4,AMPKalpha,MAPk-Ak2,IP3K2* |
| molecular adaptor activity | GO:MF | 0.02 | *N,Pax,gcl,BicD,Atg17,kibra,sbb,dos,Tob,stumps,pnut,Plp,tna,shn,Pits,Smr,Not1,Vti1b,Snap29,dmpd,Sirt1,Crtc,Lamtor1,Lnk,aph-1,Daxx,Faf2,Clamp,Graf* |
| protein tyrosine kinase binding | GO:MF | 0.022 | *stumps,Shc,spri,Lnk* |
| miRNA binding | GO:MF | 0.024 | *AGO1,AGO2* |
| transcription factor binding | GO:MF | 0.025 | *edl,hth,Trf2,tup,cic,shn,sd,Hipk,Taf3,ci,TfIIA-S,Sirt1,Crtc,ftz-f1,DnaJ-1* |
| GTP binding | GO:MF | 0.028 | *Galphao,Rab2,pnut,Rheb,Rab4,Rab5,Rab1,Non1,Rab8,Rab7,Rho1,Rab18,Ras85D,Rab14,cta,CG2017,Ras64B* |
| p53 binding | GO:MF | 0.029 | *Taf3,Sirt1,Ubc6* |
| RNA polymerase II cis-regulatory region sequence-specific DNA binding | GO:MF | 0.029 | *CG14431,CG9650,Atf3,Antp,Eip78C,ken,klu,E(spl)m3-HLH,luna,E(spl)mbeta-HLH,Chd64,aop,zfh1,Glut4EF,eve,Sox14,hth,ttk,tup,slbo,shn,sd,Hr4,Mnt,maf-S,ci,crol,ova,ftz-f1* |
| growth factor activity | GO:MF | 0.034 | *Pvf3,vn,myo,miple2,gbb* |
| cis-regulatory region sequence-specific DNA binding | GO:MF | 0.037 | *CG14431,CG9650,Atf3,Antp,Eip78C,ken,klu,E(spl)m3-HLH,luna,E(spl)mbeta-HLH,Chd64,aop,zfh1,Glut4EF,eve,Sox14,hth,ttk,tup,slbo,shn,sd,Hr4,Mnt,maf-S,ci,crol,ova,ftz-f1* |
| guanyl ribonucleotide binding | GO:MF | 0.037 | *Galphao,Rab2,pnut,Rheb,Rab4,Rab5,Rab1,Non1,Rab8,Rab7,Rho1,Rab18,Ras85D,Rab14,cta,CG2017,Ras64B* |
| guanyl nucleotide binding | GO:MF | 0.037 | *Galphao,Rab2,pnut,Rheb,Rab4,Rab5,Rab1,Non1,Rab8,Rab7,Rho1,Rab18,Ras85D,Rab14,cta,CG2017,Ras64B* |
| axon guidance receptor activity | GO:MF | 0.04 | *CG13506,robo2,Dscam1* |
| phosphotransferase activity, alcohol group as acceptor | GO:MF | 0.043 | *hppy,FER,htl,cdi,IP3K1,Pdk1,InR,Gprk2,Lk6,Pink1,Egfr,Dyrk2,par-1,Hipk,fwd,Ilk,Pitslre,aPKC,Dyrk3,Slik,Akt,AMPKalpha,MAPk-Ak2,IP3K2* |
| protein serine/threonine kinase activity | GO:MF | 0.046 | *hppy,cdi,Pdk1,Gprk2,Lk6,Pink1,Dyrk2,par-1,Hipk,Pitslre,aPKC,Dyrk3,Slik,Akt,AMPKalpha,MAPk-Ak2* |
| carbohydrate derivative binding | GO:MF | 0.046 | *cv-2,Pka-R2,hppy,Myo31DF,CG17646,FER,CG9338,htl,cdi,w,crim,Sulf1,crok,CG9336,CG3164,Pdk1,InR,Gprk2,Sema1a,Lk6,CG6583,Atet,Galphao,Rab2,numb,Klp98A,vn,pnut,galectin,Cht2,Rheb,Pink1,Egfr,Dyrk2,UbcE2H,Rab4,Mhcl,sick,Rab5,par-1,Rab1,Non1,CG33298,Rab8,Rab7,Hipk,Rho1,Rab18,Ilk,Ras85D,miple2,Rab14,Pitslre,Act87E,aPKC,cta,PAN3,Dyrk3,Slik,Hlc,Hsp70Ab,Ubc6,Dlic,CG2017,Akt,Ras64B,ClpX,AMPKalpha,MAPk-Ak2,Hsp83* |
| ubiquitin-like protein transferase activity | GO:MF | 0.048 | *Rnf11,mura,Kaz,CG2991,tna,gzl,UbcE2H,Pits,Ufd4,Ubr1,CG32369,Bruce,Ubc6,Atg3,sordd1,Mkrn1,CG12099* |
| cAMP response element binding protein binding | GO:MF | 0.048 | *ci,Crtc* |
| transmembrane receptor protein tyrosine kinase adaptor activity | GO:MF | 0.048 | *stumps,Lnk* |
| calmodulin binding | GO:MF | 0.048 | *Myo31DF,Lk6,beta-Spec,alpha-Spec,Cka,MAPk-Ak2,IP3K2* |
| G protein-coupled receptor kinase activity | GO:MF | 0.048 | *Gprk2,AMPKalpha* |
| inositol tetrakisphosphate kinase activity | GO:MF | 0.048 | *IP3K1,IP3K2* |
| calcium-dependent protein binding | GO:MF | 0.048 | *Alg-2,IP3K2* |
| ferrous iron binding | GO:MF | 0.048 | *Fer2LCH,Fer1HCH,IscU* |
| inositol-1,4,5-trisphosphate 3-kinase activity | GO:MF | 0.048 | *IP3K1,IP3K2* |

### 1.23. Gene Ontology Enrichment Analysis: sensory complex

Significant GO terms (p < 0.05) with associated genes

| GO Term | Source | P-value | Genes |
| --- | --- | --- | --- |
| septate junction assembly | GO:BP | 9.55e-14 | *vari,CG9628,crim,pck,sinu,Nrg,cold,kune,crok,scrib,bou* |
| animal organ morphogenesis | GO:BP | 3.29e-09 | *sv,qua,jv,crb,pros,trn,sinu,Nrg,Frl,pnt,rau,baz,cora,cv-c,kmr,Mob2,fog,par-1,ct,scrib,Dg,sano,how,eff* |
| establishment of glial blood-brain barrier | GO:BP | 1.39e-06 | *pck,sinu,Nrg,kune,cora* |
| zonula adherens assembly | GO:BP | 5.93e-06 | *crb,sdt,baz,scrib* |
| establishment or maintenance of polarity of embryonic epithelium | GO:BP | 1.03e-04 | *crb,sdt,scrib* |
| cell adhesion involved in heart morphogenesis | GO:BP | 8.43e-04 | *sinu,Nrg,cora* |
| apical protein localization | GO:BP | 0.002 | *crb,sdt,baz* |
| regulation of establishment of planar polarity | GO:BP | 0.003 | *pnt,baz,kmr* |
| regulation of synaptic transmission, cholinergic | GO:BP | 0.004 | *crim,crok,bou* |
| border follicle cell delamination | GO:BP | 0.004 | *baz,par-1* |
| epidermal cell differentiation | GO:BP | 0.004 | *qua,cora,par-1,scrib* |
| liquid clearance, open tracheal system | GO:BP | 0.005 | *crb,crim,pck* |
| regulation of R7 cell differentiation | GO:BP | 0.005 | *sv,pros,eff* |
| dorsal closure | GO:BP | 0.006 | *crb,sinu,cora,cv-c,scrib* |
| regulation of tube length, open tracheal system | GO:BP | 0.007 | *vari,kune,sano* |
| establishment or maintenance of polarity of follicular epithelium | GO:BP | 0.007 | *crb,scrib,Dg* |
| regulation of dendrite morphogenesis | GO:BP | 0.01 | *vvl,cv-c,babos* |
| cell dedifferentiation | GO:BP | 0.014 | *pros,pnt* |
| muscle attachment | GO:BP | 0.014 | *CAP,Dg,how* |
| pole plasm protein localization | GO:BP | 0.019 | *par-1,scrib* |
| axon ensheathment | GO:BP | 0.019 | *Nrg,how* |
| R3/R4 cell fate commitment | GO:BP | 0.019 | *pnt,scrib* |
| regulation of photoreceptor cell differentiation | GO:BP | 0.021 | *sv,pros,eff* |
| receptor clustering | GO:BP | 0.022 | *scrib,Fur1* |
| positive regulation of epidermal growth factor receptor signaling pathway | GO:BP | 0.024 | *rau,Rgl* |
| regulation of circadian sleep/wake cycle, sleep | GO:BP | 0.026 | *crim,crok,bou* |
| positive regulation of voltage-gated potassium channel activity | GO:BP | 0.029 | *crim,crok* |
| regulation of cell projection organization | GO:BP | 0.031 | *pros,vvl,baz,cv-c,babos* |
| negative regulation of signal transduction | GO:BP | 0.032 | *crb,SoxN,pnt,cv-c,par-1,CycG,eff,chrb* |
| centrosome localization | GO:BP | 0.032 | *crb,baz* |
| cell morphogenesis involved in Malpighian tubule morphogenesis | GO:BP | 0.034 | *crb* |
| G1 to G0 transition | GO:BP | 0.034 | *pros* |
| negative regulation of Rho protein signal transduction | GO:BP | 0.034 | *cv-c* |
| negative regulation of removal of superoxide radicals | GO:BP | 0.034 | *Vdup1* |
| peripheral nervous system development | GO:BP | 0.034 | *pros,vvl,ct* |
| zonula adherens maintenance | GO:BP | 0.034 | *crb* |
| positive regulation of ubiquitin-dependent endocytosis | GO:BP | 0.034 | *sdt* |
| calcium-independent cell-cell adhesion via plasma membrane cell-adhesion molecules | GO:BP | 0.034 | *Fas1* |
| salivary gland cavitation | GO:BP | 0.034 | *fog* |
| polar body extrusion after meiotic divisions | GO:BP | 0.034 | *spir* |
| establishment of centrosome localization | GO:BP | 0.034 | *baz* |
| negative regulation of cell adhesion | GO:BP | 0.036 | *baz,par-1* |
| cortical actin cytoskeleton organization | GO:BP | 0.036 | *crb,Frl,cv-c* |
| muscle cell development | GO:BP | 0.038 | *pnt,Dg,how* |
| spiracle morphogenesis, open tracheal system | GO:BP | 0.041 | *cv-c,ct* |
| regulation of potassium ion transmembrane transport | GO:BP | 0.041 | *crim,crok* |
| branch fusion, open tracheal system | GO:BP | 0.041 | *dnd,trn* |
| trachea development | GO:BP | 0.048 | *crb,pnt* |
| protein binding | GO:MF | 0.006 | *dnd,Vdup1,qua,crb,SoxN,vari,trn,CAP,CG45263,sinu,spir,Act5C,Nrg,bol,Frl,pnt,rau,CG14995,sdt,baz,Fas3,Jupiter,cora,fax,Act42A,Rgl,Mob2,CG13917,fog,par-1,scrib,Fas1,His3.3A,CG6118,Dg,His3.3B,CycG,Msp300,eff* |
| cell adhesion molecule binding | GO:MF | 0.006 | *CG45263,Nrg,Fas3,Fas1,Dg* |
| phosphatidylinositol binding | GO:MF | 0.025 | *qua,crim,baz,crok,kmr* |
| structural constituent of cytoskeleton | GO:MF | 0.031 | *betaTub56D,Jupiter,alphaTub84B* |

### 1.24. Gene Ontology Enrichment Analysis: crystal cell

Significant GO terms (p < 0.05) with associated genes

| GO Term | Source | P-value | Genes |
| --- | --- | --- | --- |
| organic acid metabolic process | GO:BP | 7.96e-16 | *CG18609,FASN3,CG7910,CG8534,FASN2,CG15531,bond,CG6660,CG9459,Desat1,eloF,CG4860,ACC,CG30008,Irp-1B,CROT,CG16904,CG7900,Sc2,Idh,Acbp2,CG6178,Acox3,Men,Fatp1,Mcad,CG7920,CG8814,Ech1,Hydr2,Hacd1,CG31523,Mfe2,Gdh,CG8839,Arc42,yip2,Shmt,CG17896,pdgy,Mtpalpha,Mdh2,Etfb,scu,Etf-QO,Hacd2,CG4598,Hibch,Echs1,CG7461,kdn,wal,Acsl,N,CG17691,Cth,whd* |
| very long-chain fatty acid biosynthetic process | GO:BP | 1.02e-09 | *CG18609,CG8534,bond,CG6660,CG9459,eloF,CG30008,CG16904,Sc2,Hacd1,CG31523,Hacd2* |
| lipid catabolic process | GO:BP | 2.99e-08 | *CG7910,Hnf4,CG4860,CROT,CG7900,Jheh2,Acox3,Mcad,Ech1,Hydr2,CG17292,Lsd-2,Mfe2,CG8839,Arc42,yip2,pdgy,Fitm,Mtpalpha,Etfb,foxo,Etf-QO,CG4598,Echs1,CG7461,wal,whd* |
| catabolic process | GO:BP | 2.49e-07 | *CG7910,Hnf4,Desat1,CG4860,CROT,CG7900,Jheh2,daw,Cat,Trc8,CG46385,Acox3,Mcad,Debcl,Ntan1,Ech1,Hydr2,CG17292,Sod1,Prosalpha4,Lsd-2,zda,Myc,Hsc70-4,Mfe2,Gdh,CG8839,Arc42,yip2,Shmt,CG17896,RpLP0-like,Apt1,pdgy,Fitm,Mtpalpha,p47,Etfb,foxo,Prosbeta1,Fkbp39,Ubi-p63E,Etf-QO,Prosalpha5,TER94,Pomp,Prosbeta4,EMC6,CG4598,Hibch,Prosalpha2,wds,Prosbeta6,Prosbeta2,Echs1,CG7461,BI-1,PAN3,Hsc70-5,Prosalpha7,dnc,eff,Rpn6,CG6567,REG,rump,wal,pelo,N,CG6878,Ubc7,CG6966,Rpt3,Prosbeta3,Prosbeta7,Prosbeta5,Prosalpha3,HINT1,CG17691,scny,EloC,sordd1,Pka-C1,whd,mbf1* |
| fatty acid elongation, monounsaturated fatty acid | GO:BP | 1.21e-06 | *CG18609,CG8534,bond,CG6660,CG9459,eloF,CG30008,CG16904,CG31523* |
| fatty acid elongation, polyunsaturated fatty acid | GO:BP | 1.21e-06 | *CG18609,CG8534,bond,CG6660,CG9459,eloF,CG30008,CG16904,CG31523* |
| fatty acid elongation, saturated fatty acid | GO:BP | 1.21e-06 | *CG18609,CG8534,bond,CG6660,CG9459,eloF,CG30008,CG16904,CG31523* |
| fatty acid beta-oxidation using acyl-CoA dehydrogenase | GO:BP | 4.63e-06 | *CG4860,Mcad,Arc42,Etfb,CG7461,wal* |
| sulfur compound metabolic process | GO:BP | 1.25e-05 | *CG17562,CG17560,FarO,ACC,GstE3,CG4020,GstT1,fbl,CG6178,GstT3,GstT4,Sam-S,Hs6st,ScsbetaG,Hmgs,pdgy,Fitm,scu,Pmvk,ScsbetaA,Acsl,Cth,Dpck* |
| cuticle hydrocarbon biosynthetic process | GO:BP | 1.08e-04 | *Cyp4g1,FASN2,Desat1,Cpr* |
| sphingolipid biosynthetic process | GO:BP | 1.43e-04 | *CG18609,CG8534,bond,CG6660,CG9459,eloF,CG30008,CG16904,Hacd1,CG31523,Hacd2* |
| tail-anchored membrane protein insertion into ER membrane | GO:BP | 4.10e-04 | *EMC4,EMC6,EMC5,EMC3,EMC8-9* |
| heme biosynthetic process | GO:BP | 4.88e-04 | *Alas,Pbgs,l(3)02640,FeCH,Urod,CG5532* |
| snRNA pseudouridine synthesis | GO:BP | 0.002 | *Nop60B,CG7637,NHP2* |
| tRNA transcription by RNA polymerase III | GO:BP | 0.004 | *Polr2E,Polr2F,Polr2L,Polr2K,Polr2H* |
| protoporphyrinogen IX biosynthetic process | GO:BP | 0.004 | *Alas,Pbgs,l(3)02640,Urod* |
| nucleobase-containing compound metabolic process | GO:BP | 0.004 | *CG17562,CG17560,fus,vvl,salm,Hnf4,FarO,peb,svp,ACC,CG34183,salr,CG12375,CG4020,Dr,CG33958,esg,spidey,Idh,mirr,NADK,fbl,r-l,CG6178,CG46385,Men,Ntan1,Trmt112,klu,Pde1c,Dhx15,Naxd,Myc,Atf3,Hsc70-4,SF2,ScsbetaG,Gdh,CG6712,Nop60B,Shmt,hoip,Hmgs,CG42813,CG12288,CG17896,CG32409,Rbm13,RpLP0-like,MED11,CG13096,pdgy,CG7637,pths,Nop56,CG33217,nop5,Pgls,Fitm,Mes2,NHP2,Xrp1,Tctp,CG9932,Nurf-38,Samuel,Lst8,foxo,Polr2E,TfIIA-L,Cipc,scu,cwo,cnc,mod,La,CG11583,CG5862,Cmpk,Non3,Rpi,CG6770,Nap1,CG4866,SmD3,Bka,CG1542,pit,Fib,Pmvk,psq,Nfs1,Hers,koi,Ip259,CG1789,BI-1,PAN3,CG8545,CG1234,Ak2,ras,dnc,Polr2F,Aprt,CG4364,CG11858,ScsbetaA,Polr2L,Polr2K,CG4038,REG,Taldo,rump,CG9344,Nnp-1,Acsl,pelo,crc,N,Sumo,SmD2,rgr,HINT1,Naprt,EloC,SerRS,poly,Phf5a,Polr2H,pan,mld,Dpck,mbf1* |
| proteasome assembly | GO:BP | 0.005 | *CG12321,Hsp83,PSMG1,Pomp,Rpn6* |
| determination of adult lifespan | GO:BP | 0.007 | *ImpL2,daw,Cat,Trx-2,Men,Tspo,Sam-S,Sod1,Mtpalpha,Trxr-1,Indy,foxo,cnc,Prosalpha5,eEF1alpha1,N,mld* |
| long-chain fatty-acyl-CoA metabolic process | GO:BP | 0.011 | *CG17562,CG17560,FarO,CG4020,Acsl* |
| chaperone cofactor-dependent protein refolding | GO:BP | 0.019 | *Hsc70-4,Tpr2,HIP,CG11267,HIP-R,Hsc70-5* |
| protein import into mitochondrial matrix | GO:BP | 0.019 | *Tom40,Tom7,Roe1,Hsc70-5,mge,CG6878* |
| multivesicular body fusion to apical plasma membrane | GO:BP | 0.028 | *Hsp83,Stip1* |
| succinyl-CoA metabolic process | GO:BP | 0.028 | *ScsbetaG,ScsbetaA* |
| snoRNA guided rRNA pseudouridine synthesis | GO:BP | 0.028 | *CG7637,CG4038* |
| regulation of bicoid mRNA localization | GO:BP | 0.028 | *cnc,Pka-C1* |
| 'de novo' protein folding | GO:BP | 0.028 | *Hsc70-4,Tpr2,HIP,CG11267,HIP-R,Hsc70-5* |
| positive regulation of entry into reproductive diapause | GO:BP | 0.028 | *ImpL2,foxo* |
| butyrate catabolic process | GO:BP | 0.028 | *CG4860,Arc42* |
| muscle tissue morphogenesis | GO:BP | 0.028 | *LamC,hoip* |
| ribosomal large subunit export from nucleus | GO:BP | 0.033 | *Mys45A,CG32409,Ns2* |
| maturation of LSU-rRNA from tricistronic rRNA transcript (SSU-rRNA, 5.8S rRNA, LSU-rRNA) | GO:BP | 0.038 | *CG12288,Non3,pit,CG4364* |
| positive regulation of neuroblast proliferation | GO:BP | 0.046 | *daw,esg,klu,Hsp83,N* |
| fatty acid elongase activity | GO:MF | 9.94e-06 | *CG18609,CG8534,bond,CG6660,CG9459,eloF,CG30008,CG16904,CG31523* |
| hydro-lyase activity | GO:MF | 9.94e-06 | *FASN3,FASN2,Irp-1B,Pbgs,CAHbeta,Hacd1,Naxd,Mfe2,Mtpalpha,CG6028,Hacd2,CG4598,Echs1* |
| catalytic activity | GO:MF | 2.74e-05 | *Cyp4g1,CG18609,CG17562,FASN3,CG7910,CG16799,CG8534,CG17560,FASN2,CG15531,bond,Fkbp59,CG6660,CG9459,FarO,Desat1,Cpr,CG14615,eloF,CG4860,ACC,Alas,CG34183,CG30008,GstE3,CG12375,CG4020,Irp-1B,GstT1,CROT,CG16904,Ect3,PPO2,CG12256,nvd,CG7900,Jheh2,CG33958,Sc2,spidey,Idh,Cat,NADK,Trc8,PHGPx,fbl,r-l,CG31689,CG6178,GstT3,CG46385,Acox3,Pdk,Trx-2,Mgstl,Idi,Dhrs4,Men,Fatp1,Pbgs,eas,Mcad,CAHbeta,l(3)02640,Ntan1,Hmu,FeCH,GstT4,CG7920,Hsp83,Sam-S,p23,Fur1,Ech1,CenG1A,Hydr2,CG17292,Sod1,dysc,Hacd1,Urod,Pde1c,CG17684,Dhx15,Naxd,zda,Hs6st,CG12279,Cyp1,CG31523,Hsc70-4,SpdS,Mfe2,CG9143,ScsbetaG,Gdh,CG3655,CG8839,Arc42,Agpat3,SmydA-8,Nop60B,betaTub56D,yip2,Shmt,Hmgs,CG42813,CG17896,Fkbp12,Apt1,pdgy,pths,alphaTub84B,CanB2,CG3164,Pgls,Fitm,Mtpalpha,Hsdl2,betaTub97EF,Trxr-1,Nurf-38,atl,Mdh2,Hsc70Cb,Polr2E,Naa20A,CaMKI,scu,prtp,Prosbeta1,Haspin,Fkbp39,Cmpk,Hsp60A,CG6028,Etf-QO,Rpi,CG13284,Tina-1,TER94,bbc,pit,Hacd2,Prosbeta4,Fib,Pmvk,CG4598,CG2991,Nfs1,Hibch,CG1307,Prosbeta2,CG4822,Echs1,Aldh-III,CG7461,SelT,SppL,Hsc70-5,CG8545,Ak2,ras,alphaTub84D,dnc,kdn,Polr2F,eEF1alpha1,eff,Aprt,CG11858,ScsbetaA,Polr2L,Polr2K,CG6567,Taldo,wal,Acsl,pelo,Ubc7,mahe,Non1,Rpt3,CCT4,CG9330,Prosbeta5,HINT1,Ns2,CG17691,Naprt,scny,CG8993,Cth,SerRS,cl,sordd1,Polr2H,Pka-C1,Sod2,CG1354,whd,Dpck,CCT7,CG9281,CCT6,CCT2* |
| ATP-dependent protein folding chaperone | GO:MF | 3.46e-05 | *Hsp83,Hsc70-4,CG11267,Hsc70Cb,Hsp60A,Hsc70-5,CCT4,CCT7,CCT6,CCT2* |
| lyase activity | GO:MF | 3.46e-05 | *Cyp4g1,FASN3,FASN2,Irp-1B,CG33958,r-l,Pbgs,CAHbeta,Hmu,FeCH,Hacd1,Urod,Naxd,Mfe2,Mtpalpha,CG6028,Tina-1,Hacd2,CG4598,Echs1,Cth* |
| enoyl-CoA hydratase activity | GO:MF | 4.74e-05 | *Hacd1,Mfe2,Mtpalpha,Hacd2,CG4598,Echs1* |
| protein folding chaperone | GO:MF | 4.74e-05 | *Hsp83,zda,Hsc70-4,CG11267,Hsc70Cb,Hsp60A,Hsc70-5,CCT4,CCT7,CCT6,CCT2* |
| oxidoreductase activity | GO:MF | 2.60e-04 | *Cyp4g1,CG17562,FASN3,CG17560,FASN2,CG15531,FarO,Desat1,Cpr,CG4860,CG4020,GstT1,PPO2,nvd,Sc2,spidey,Idh,Cat,PHGPx,CG6178,GstT3,Acox3,Trx-2,Mgstl,Dhrs4,Men,Mcad,GstT4,Sod1,Mfe2,Gdh,Arc42,CG17896,Mtpalpha,Hsdl2,Trxr-1,Mdh2,scu,Etf-QO,CG13284,Aldh-III,CG7461,SelT,ras,wal,CG17691,CG8993,cl,Sod2* |
| RNA binding | GO:MF | 4.12e-04 | *fus,CG12375,Dhx15,vig2,SF2,CG9143,CG6712,Nph,CG7006,Nop60B,vito,Shmt,hoip,CG12288,Nlp,CG13096,CG3594,CG7637,pths,CG4806,Nop56,nop5,NHP2,CG11563,mod,La,CG11583,Surf6,Non3,CG4866,SmD3,CG1542,pit,Srp14,Fib,l(1)G0004,PAN3,CG8545,CG11444,CG4364,CG4038,rump,AIMP1,CG9344,mahe,SmD2,Non1,mRpS18C,eIF3i,SerRS,Phf5a,mbf1* |
| membrane insertase activity | GO:MF | 4.37e-04 | *EMC4,EMC6,EMC5,EMC3,EMC8-9* |
| RNA polymerase II activity | GO:MF | 4.37e-04 | *Polr2E,Polr2F,Polr2L,Polr2K,Polr2H* |
| RNA polymerase III activity | GO:MF | 6.48e-04 | *Polr2E,Polr2F,Polr2L,Polr2K,Polr2H* |
| RNA polymerase I activity | GO:MF | 6.48e-04 | *Polr2E,Polr2F,Polr2L,Polr2K,Polr2H* |
| antioxidant activity | GO:MF | 6.48e-04 | *GstT1,Cat,PHGPx,GstT3,Mgstl,GstT4,Sod1,Trxr-1,SelT,cl,Sod2* |
| box H/ACA snoRNA binding | GO:MF | 0.002 | *CG7637,NHP2,CG4038* |
| 3-hydroxyacyl-CoA dehydratase activity | GO:MF | 0.002 | *Hacd1,Mfe2,Hacd2* |
| snoRNA binding | GO:MF | 0.002 | *CG7637,Nop56,nop5,NHP2,CG4866,CG4038* |
| rRNA binding | GO:MF | 0.003 | *CG6712,vito,CG12288,pths,La,CG11583,Non3,CG4866,mRpS18C* |
| thiolester hydrolase activity | GO:MF | 0.003 | *FASN3,FASN2,CG7920,Apt1,Hibch,CG6567* |
| acyl-CoA dehydrogenase activity | GO:MF | 0.003 | *CG4860,Mcad,Arc42,CG7461* |
| anion binding | GO:MF | 0.003 | *Cpr,CG4860,ACC,Alas,fbl,Acbp2,CG31689,Acox3,Pdk,Mcad,CG8814,Hsp83,Sam-S,CenG1A,Dhx15,Naxd,Hsc70-4,CG9143,ScsbetaG,Gdh,Arc42,betaTub56D,Shmt,CG17896,pths,alphaTub84B,CG3164,Droj2,Mtpalpha,betaTub97EF,Trxr-1,CG11267,atl,Hsc70Cb,CaMKI,Haspin,Cmpk,Hsp60A,TER94,pit,Pmvk,Nfs1,Slmap,CG4822,Myo81F,CG7461,PAN3,Hsc70-5,Ak2,alphaTub84D,eEF1alpha1,eff,Aprt,ScsbetaA,wal,Ubc7,mahe,Non1,Rpt3,CCT4,CG9330,Ns2,Cth,SerRS,Pka-C1,CG1354,Dpck,CCT7,CG9281,CCT6,CCT2* |
| heat shock protein binding | GO:MF | 0.003 | *p23,Hsc70-4,Tpr2,Stip1,HIP,Droj2,HIP-R,Hsc70-5* |
| protein-folding chaperone binding | GO:MF | 0.004 | *p23,Hsc70-4,Stip1,HIP,Droj2,CG11267,Hsp60A,HIP-R,Roe1* |
| acyltransferase activity, transferring groups other than amino-acyl groups | GO:MF | 0.004 | *CG18609,FASN3,CG8534,FASN2,bond,CG6660,CG9459,CG14615,eloF,Alas,CG30008,CROT,CG16904,CG31523,Agpat3,yip2,Naa20A,whd* |
| unfolded protein binding | GO:MF | 0.005 | *Hsp83,Hsc70-4,nudC,Droj2,CG11267,Roe1,Hsc70-5,CCT4,CCT7,CCT6,CCT2* |
| nucleotide binding | GO:MF | 0.005 | *Cpr,CG4860,ACC,Idh,fbl,CG31689,Acox3,Pdk,Men,Mcad,Hsp83,Sam-S,CenG1A,Dhx15,Naxd,Hsc70-4,CG9143,ScsbetaG,Gdh,Arc42,betaTub56D,pths,alphaTub84B,CG3164,Droj2,Mtpalpha,betaTub97EF,Trxr-1,CG11267,atl,Hsc70Cb,CaMKI,Haspin,Cmpk,Hsp60A,TER94,pit,Pmvk,Slmap,CG4822,Myo81F,CG7461,PAN3,Hsc70-5,Ak2,ras,alphaTub84D,eEF1alpha1,eff,Aprt,ScsbetaA,wal,Ubc7,mahe,Non1,Rpt3,CCT4,CG9330,Ns2,SerRS,Pka-C1,CG1354,Dpck,CCT7,CG9281,CCT6,CCT2* |
| monoacylglycerol lipase activity | GO:MF | 0.007 | *CG7910,CG7900,Hydr2,CG8839* |
| glutathione peroxidase activity | GO:MF | 0.008 | *GstT1,PHGPx,GstT3,Mgstl,GstT4* |
| fatty acid amide hydrolase activity | GO:MF | 0.01 | *CG7910,CG7900,CG8839* |
| oxidoreductase activity, acting on the aldehyde or oxo group of donors, NAD or NADP as acceptor | GO:MF | 0.012 | *CG17562,CG17560,FarO,CG4020,spidey,CG17896,Aldh-III* |
| DNA-directed 5'-3' RNA polymerase activity | GO:MF | 0.012 | *Polr2E,Polr2F,Polr2L,Polr2K,Polr2H* |
| fatty-acyl-CoA binding | GO:MF | 0.013 | *Acbp2,CG8814,CG17896,CG7461* |
| protein-disulfide reductase (NAD(P)H) activity | GO:MF | 0.016 | *Trxr-1,SelT,cl* |
| isomerase activity | GO:MF | 0.017 | *Fkbp59,Mgstl,Idi,p23,Ech1,zda,Cyp1,Nop60B,Fkbp12,prtp,Fkbp39,Rpi,CG4598,CG11858* |
| peroxidase activity | GO:MF | 0.018 | *GstT1,Cat,PHGPx,GstT3,Mgstl,GstT4,cl* |
| oxidoreductase activity, acting on peroxide as acceptor | GO:MF | 0.018 | *GstT1,Cat,PHGPx,GstT3,Mgstl,GstT4,cl* |
| purine nucleotide binding | GO:MF | 0.018 | *Cpr,ACC,Idh,fbl,CG31689,Pdk,Men,Hsp83,Sam-S,CenG1A,Dhx15,Naxd,Hsc70-4,CG9143,ScsbetaG,Gdh,betaTub56D,pths,alphaTub84B,CG3164,Droj2,Mtpalpha,betaTub97EF,CG11267,atl,Hsc70Cb,CaMKI,Haspin,Cmpk,Hsp60A,TER94,pit,Pmvk,Slmap,CG4822,Myo81F,PAN3,Hsc70-5,Ak2,alphaTub84D,eEF1alpha1,eff,Aprt,ScsbetaA,Ubc7,mahe,Non1,Rpt3,CCT4,CG9330,Ns2,SerRS,Pka-C1,CG1354,Dpck,CCT7,CG9281,CCT6,CCT2* |
| peptidyl-prolyl cis-trans isomerase activity | GO:MF | 0.018 | *Fkbp59,zda,Cyp1,Fkbp12,Fkbp39,CG11858* |
| oxidoreductase activity, acting on the CH-OH group of donors, NAD or NADP as acceptor | GO:MF | 0.018 | *FASN3,FASN2,nvd,spidey,Idh,Dhrs4,Men,Mfe2,Mtpalpha,Mdh2,scu,ras* |
| 17-beta-hydroxysteroid dehydrogenase (NAD+) activity | GO:MF | 0.019 | *Mfe2,scu* |
| 3-hydroxyacyl-CoA dehydrogenase activity | GO:MF | 0.021 | *Mfe2,Mtpalpha,scu* |
| protein-disulfide reductase activity | GO:MF | 0.025 | *Trx-2,Trxr-1,SelT,CG8993,cl* |
| oxidoreductase activity, acting on the CH-CH group of donors | GO:MF | 0.026 | *CG4860,Sc2,Acox3,Mcad,Arc42,CG7461* |
| acyltransferase activity | GO:MF | 0.026 | *CG18609,FASN3,CG8534,FASN2,bond,CG6660,CG9459,CG14615,eloF,Alas,CG30008,CROT,CG16904,Trc8,CG6178,CG31523,Agpat3,yip2,Hmgs,Naa20A,Tina-1,CG2991,kdn,eff,Ubc7,sordd1,whd* |
| threonine-type endopeptidase activity | GO:MF | 0.029 | *Prosbeta1,Prosbeta2,Prosbeta5* |
| alcohol-forming very long-chain fatty acyl-CoA reductase activity | GO:MF | 0.034 | *CG17562,CG17560,FarO,CG4020* |
| transferase activity, transferring alkyl or aryl (other than methyl) groups | GO:MF | 0.035 | *GstE3,GstT1,GstT3,Mgstl,l(3)02640,GstT4,Sam-S,SpdS* |
| oxidoreductase activity, acting on a sulfur group of donors, NAD(P) as acceptor | GO:MF | 0.039 | *Trxr-1,SelT,cl* |
| long-chain-fatty-acyl-CoA reductase activity | GO:MF | 0.039 | *FarO,spidey* |
| FK506 binding | GO:MF | 0.039 | *Fkbp59,Fkbp39* |
| Hsp70 protein binding | GO:MF | 0.039 | *Stip1,HIP,Droj2,HIP-R* |
| flavin adenine dinucleotide binding | GO:MF | 0.047 | *Cpr,CG4860,Acox3,Mcad,Arc42,Trxr-1,CG7461,wal* |
| oxidoreductase activity, acting on CH-OH group of donors | GO:MF | 0.048 | *FASN3,FASN2,nvd,spidey,Idh,Dhrs4,Men,Mfe2,Mtpalpha,Mdh2,scu,ras* |
| adenyl ribonucleotide binding | GO:MF | 0.048 | *ACC,fbl,CG31689,Pdk,Hsp83,Sam-S,Dhx15,Naxd,Hsc70-4,CG9143,ScsbetaG,Gdh,pths,CG3164,Droj2,CG11267,Hsc70Cb,CaMKI,Haspin,Cmpk,Hsp60A,TER94,pit,Pmvk,Slmap,CG4822,Myo81F,PAN3,Hsc70-5,Ak2,eff,Aprt,ScsbetaA,Ubc7,mahe,Rpt3,CCT4,CG9330,SerRS,Pka-C1,CG1354,Dpck,CCT7,CG9281,CCT6,CCT2* |
| pyrophosphatase activity | GO:MF | 0.049 | *CG31689,Hsp83,CenG1A,Hsc70-4,betaTub56D,CG42813,CG3164,Fitm,betaTub97EF,Nurf-38,atl,Hsc70Cb,Hsp60A,TER94,pit,CG4822,Hsc70-5,eEF1alpha1,Non1,Rpt3,CCT4,CG9330,Ns2,CG1354,CCT7,CG9281,CCT6,CCT2* |
