## Supplemental Table 7 for "A complete single-cell atlas of the embryonic *Drosophila* heart and the cell-type specific role of Tinman"

### 1.1. Gene Ontology Enrichment Analysis: Notch cluster

Significant GO terms (p < 0.05) with associated genes

| GO Term | Source | P-value | Genes |
| --- | --- | --- | --- |
| Wnt-protein binding | MF | 0.043 | *peg* |

### 1.2. Gene Ontology Enrichment Analysis: longitudinal glia - naz

Significant GO terms (p < 0.05) with associated genes

| GO Term | Source | P-value | Genes |
| --- | --- | --- | --- |
| intracellularly cAMP-activated cation channel activity | MF | 6.20e-04 | *CngA, CG42260* |
| intracellularly cGMP-activated cation channel activity | MF | 6.20e-04 | *CngA, CG42260* |
| intracellularly cyclic nucleotide-activated monoatomic cation channel activity | MF | 6.20e-04 | *CngA, CG42260* |
| cGMP binding | MF | 0.001 | *CngA, CG42260* |
| intracellularly ligand-gated monoatomic ion channel activity | MF | 0.003 | *CngA, CG42260* |
| peroxiredoxin activity | MF | 0.004 | *Prx2540-1, Prx2540-2* |
| peroxidase activity | MF | 0.035 | *Prx2540-1, Prx2540-2* |
| phosphoenolpyruvate carboxykinase activity | MF | 0.035 | *Pepck2* |
| phosphoenolpyruvate carboxykinase (GTP) activity | MF | 0.035 | *Pepck2* |
| oxidoreductase activity, acting on peroxide as acceptor | MF | 0.035 | *Prx2540-1, Prx2540-2* |
| monoatomic cation channel activity | MF | 0.04 | *nan, CngA, CG42260* |
| guanyl ribonucleotide binding | MF | 0.043 | *Pepck2, CngA, CG42260* |
| antioxidant activity | MF | 0.043 | *Prx2540-1, Prx2540-2* |
| guanyl nucleotide binding | MF | 0.043 | *Pepck2, CngA, CG42260* |
| glutathione hydrolase activity | MF | 0.045 | *CG4829* |
| double-stranded RNA adenosine deaminase activity | MF | 0.045 | *blanks* |

### 1.3. Gene Ontology Enrichment Analysis: plasmatocytes

Significant GO terms (p < 0.05) with associated genes

| GO Term | Source | P-value | Genes |
| --- | --- | --- | --- |
| peptidyl-glutamate ADP-deribosylation | BP | 0.005 | *CG33054, CG33056* |
| ADP-ribosylglutamate hydrolase activity | MF | 0.002 | *CG33054, CG33056* |
| purine nucleoside binding | MF | 0.002 | *CG33054, CG33056* |
| glutathione transferase activity | MF | 0.006 | *GstD9, GstE11, GstD1* |
| transferase activity, transferring alkyl or aryl (other than methyl) groups | MF | 0.026 | *GstD9, GstE11, GstD1* |
| hydrolase activity, hydrolyzing N-glycosyl compounds | MF | 0.029 | *CG33054, CG33056* |
| DDT-dehydrochlorinase activity | MF | 0.048 | *GstD1* |
| transcription regulator inhibitor activity | MF | 0.048 | *Dad* |

### 1.4. Gene Ontology Enrichment Analysis: eve-PC

Significant GO terms (p < 0.05) with associated genes

| GO Term | Source | P-value | Genes |
| --- | --- | --- | --- |
| motor neuron axon guidance | BP | 4.64e-06 | *side, fz2, beat-IIb, Mp, beat-IIa, caps, spz, tok, beat-IIIc, cher, Fas2, fend, NetB* |
| animal organ morphogenesis | BP | 9.61e-06 | *heph, sano, crb, Dg, AdamTS-A, fz, Timp, fz2, rau, ct, pk, bab2, bdl, caps, AdamTS-B, shg, pnt, Fhos, tok, dlp, fng, Myo81F, Acer, mav, Fas2, app, mbl, mbc, RhoGEF64C, LamC, wb, ed, stg, Tak1, Ubx, Src64B, tx, bnl, daw, hth, drl, sog, Scr, NetB* |
| cell surface receptor signaling pathway | BP | 5.26e-05 | *crb, PRAS40, scb, fz, kek1, CG15529, Toll-7, fz2, rau, spz6, Mp, AdamTS-B, GluRIIB, shg, Pvf3, Prosap, Buffy, pnt, spz, tok, sona, dlp, Rgl, fng, mav, Fas2, Roc1a, mbc, Dh31-R, E(spl)m6-BFM, ed, Tak1, Src64B, Cad96Ca, Ilp2, CG32447, GluRIIC, bnl, daw, apt, drl, sog* |
| homophilic cell adhesion via plasma membrane adhesion molecules | BP | 8.50e-04 | *fz, bdl, caps, shg, CG13506, Fas2, ed, Cad96Ca* |
| branch fusion, open tracheal system | BP | 0.004 | *caps, shg, ed, form3, bnl* |
| cellular response to growth factor stimulus | BP | 0.006 | *PRAS40, Toll-7, rau, pnt, tok, dlp, mav, Src64B, bnl, daw, sog* |
| positive regulation of fibroblast growth factor receptor signaling pathway | BP | 0.012 | *rau, dlp, Src64B* |
| positive regulation of circadian sleep/wake cycle, sleep | BP | 0.013 | *wake, mld, Acer, Sh* |
| ommatidial rotation | BP | 0.018 | *fz, pk, shg, pnt, Tak1* |
| cell competition in a multicellular organism | BP | 0.02 | *shg, spz, mbc, drpr* |
| striated muscle cell differentiation | BP | 0.025 | *Dg, DAAM, cher, Six4, mbc, mspo, Zasp52, blow* |
| midgut development | BP | 0.025 | *scb, fz, fz2, Ubx, Scr* |
| wing disc dorsal/ventral pattern formation | BP | 0.029 | *crb, pnt, dlp, fng, CG1943, drpr* |
| female germline ring canal formation, actin assembly | BP | 0.029 | *cher, Src64B* |
| secondary branching, open tracheal system | BP | 0.029 | *pnt, bnl* |
| imaginal disc-derived wing margin morphogenesis | BP | 0.034 | *heph, fz, fz2, ct, dlp, fng* |
| female gonad development | BP | 0.039 | *fz2, ct, bab2* |
| specification of animal organ identity | BP | 0.045 | *Ubx, hth* |
| growth factor activity | MF | 3.85e-05 | *spz6, Pvf3, spz, mav, Ilp2, bnl, daw, sog* |
| signaling receptor binding | MF | 0.008 | *CG5742, slow, crb, scb, kek1, wake, spz6, Pvf3, Prosap, Idgf4, spz, mav, plx, ed, Src64B, Ilp2, bnl, daw, sog, Tollo* |
| Wnt receptor activity | MF | 0.028 | *fz, fz2, drl* |
| growth factor receptor binding | MF | 0.037 | *kek1, Pvf3, Idgf4, ed, bnl* |
| cell adhesion molecule binding | MF | 0.046 | *Dg, scb, CG2082, bdl, plx, Fas2, ed, nrm* |
| metallopeptidase activity | MF | 0.046 | *Psa, AdamTS-A, Ppn, AdamTS-B, CG4678, tok, sona, Acer, gogo, CG30049, CG10939, CG4017, Zasp52* |

### 1.5. Gene Ontology Enrichment Analysis: bap-PC

Significant GO terms (p < 0.05) with associated genes

| GO Term | Source | P-value | Genes |
| --- | --- | --- | --- |
| animal organ morphogenesis | BP | 1.88e-09 | *hth, Traf4, odd, flz, CadN, Dys, CG43658, mthl5, sano, Timp, pk, tin, scrib, AdamTS-A, pnr, Dg, mew, Myo81F, nvy, shg, fw, Smyd4-3, elB, lbl, NetB, bdl, twi, RhoGEF64C, Pura, Hs6st, rst, sfl, trn, bab2, dally, Ubx, rho, srp, Pvr, caps, Wnt4, fzr, hbs, emc, CG16758, pyd, heph, RhoGAP71E, AdamTS-B, Doa, byn, ex, rau, Ten-m, msi* |
| homophilic cell adhesion via plasma membrane adhesion molecules | BP | 1.88e-09 | *CadN, Fas1, CG13506, Cad87A, shg, fw, bdl, rst, caps, hbs, Cad96Ca, CG34353, ImpL2, Fas3* |
| gastrulation | BP | 1.14e-07 | *Traf4, pyr, T48, bap, shg, sgl, tsl, twi, sfl, Ubx, srp, ths, RhoGAP71E, byn, Pld, srw* |
| motor neuron axon guidance | BP | 1.24e-06 | *fend, beat-IIIc, side, Mp, NetB, Ptp10D, trn, dally, beat-IIb, caps, Wnt4, for, ko, Ten-m* |
| mesodermal cell fate specification | BP | 9.83e-05 | *pyr, twi, Ubx, ths* |
| heterophilic cell-cell adhesion via plasma membrane cell adhesion molecules | BP | 3.14e-04 | *scb, beat-IIIc, mew, ItgaPS4, beat-IIb, hbs, Fas3* |
| cell surface receptor signaling pathway | BP | 5.59e-04 | *Traf4, Trim9, CG31183, pyr, Ser, mthl5, scb, E(spl)m6-BFM, spz6, Egfrap, mew, Pli, shg, elB, sgl, tsl, Mp, CG15529, sfl, Ptp10D, dally, rho, ItgaPS4, Pvr, pigs, egr, Wnt4, apt, hbs, Cad96Ca, ImpL2, wdp, for, ths, AdamTS-B, spict, Doa, Jafrac2, rau, Usp12-46, srw* |
| striated muscle cell differentiation | BP | 6.23e-04 | *mspo, tin, Dg, Zasp52, tmod, rst, Six4, rols, Whamy, hbs, sls* |
| larval salivary gland morphogenesis | BP | 0.002 | *AdamTS-A, trn, rho, caps* |
| mesoderm migration involved in gastrulation | BP | 0.003 | *pyr, sgl, sfl, byn* |
| midgut development | BP | 0.005 | *scb, mew, Ubx, srp, emc, byn* |
| R7 cell development | BP | 0.008 | *CadN, bdl, rau, Ten-m* |
| regulation of striated muscle tissue development | BP | 0.011 | *twi, rst, hbs* |
| myoblast migration | BP | 0.012 | *pyr, ths* |
| muscle cell development | BP | 0.02 | *Dys, Dg, Zasp52, twi, tmod, rols, sls* |
| intercellular transport | BP | 0.023 | *Inx2, shakB, ogre* |
| larval visceral muscle development | BP | 0.023 | *pyr, Msp300, ths* |
| dorsal closure | BP | 0.027 | *Traf4, scb, scrib, Mcr, rho, srp, Pvr, emc, pyd* |
| pericardial nephrocyte differentiation | BP | 0.03 | *pyr, tin, pnr* |
| actin filament network formation | BP | 0.031 | *spir, Fim* |
| branch fusion, open tracheal system | BP | 0.032 | *shg, trn, caps, pyd* |
| calcium-dependent cell-cell adhesion via plasma membrane cell adhesion molecules | BP | 0.036 | *CadN, Cad87A, shg, Cad96Ca* |
| glial cell migration | BP | 0.037 | *pyr, NetB, Pvr, fzr, ths* |
| regulation of myoblast fusion | BP | 0.038 | *mspo, rst, hbs* |
| head involution | BP | 0.041 | *scyl, shg, Mcr, emc, pyd* |
| retinal ganglion cell axon guidance | BP | 0.046 | *CadN, gogo, Wnt4* |
| anterior Malpighian tubule development | BP | 0.046 | *Ubx, Pvr, rols* |
| regulation of ubiquitin-protein transferase activity | BP | 0.046 | *CG6966, fzr, Fem-1* |
| cell adhesion molecule binding | MF | 3.66e-07 | *CG2082, CadN, Fas1, scb, Dg, mew, Tig, bdl, rst, ItgaPS4, rols, hbs, ImpL2, pyd, Fas3* |
| protein binding | MF | 0.003 | *CG2082, hth, Shab, Traf4, Con, Trim9, Sdic1, RhoGAP18B, pyr, CadN, Dys, CG31076, dpr6, Ser, Fas1, scb, E(spl)m6-BFM, Galphaf, CG5541, dpr19, Timp, pk, SKIP, spz6, Hsp70Bb, fs(1)Yb, Egfrap, side-IV, HP1e, ReepA, FANCI, esc, scrib, Ca-beta, pip, pnr, drpr, Dg, toc, CG13506, CG14483, Idgf4, jigr1, CG5001, Zasp52, mew, Pli, Myo81F, CG5023, Kal1, Fili, Tig, CG42259, shg, CG13252, Smyd4-3, side, Oatp74D, elB, Mcr, CG10011, brat, Hesr, tsl, NetB, bdl, EndoB, twi, olf186-M, CG17258, Spn, Snap25, CG6966, tmod, CG14441, CG10353, rst, a, Ptp10D, trn, bab2, dally, Ubx, CG8064, CG7702, CG32354, srp, CG6739, sim, ItgaPS4, Septin2, Pvr, pigs, beat-IIb, dsx, 18w, egr, rols, Sans, Whamy, caps, Wnt4, SmydA-8, apt, fzr, bif, hbs, RhoL, side-V, CG34353, pain, smash, emc, ImpL2, wdp, Msp300, pyd, spir, ths, Fim, nxf2, Jafrac2, byn, ex, sls, rau, CG43980, Sh, RabGGTa, Ten-m, CG8642, Nup50, Klp54D, fbp, Fem-1, Fas3* |

### 1.6. Gene Ontology Enrichment Analysis: dead

Significant GO terms (p < 0.05) with associated genes

| GO Term | Source | P-value | Genes |
| --- | --- | --- | --- |
| glutathione metabolic process | BP | 0.026 | *GstE11, NA* |
| sulfur compound metabolic process | BP | 0.026 | *GstE11, NA* |
| glutathione transferase activity | MF | 0.014 | *GstE11, NA, NA* |
| transferase activity, transferring alkyl or aryl (other than methyl) groups | MF | 0.014 | *GstE11, NA, NA* |

### 1.7. Gene Ontology Enrichment Analysis: visceral mesoderm

Significant GO terms (p < 0.05) with associated genes

| GO Term | Source | P-value | Genes |
| --- | --- | --- | --- |
| adenylate kinase activity | MF | 0.045 | *Ak1* |
| UMP kinase activity | MF | 0.045 | *Ak1* |
| UMP/dUMP kinase activity | MF | 0.045 | *Ak1* |

### 1.8. Gene Ontology Enrichment Analysis: cardioblasts

Significant GO terms (p < 0.05) with associated genes

| GO Term | Source | P-value | Genes |
| --- | --- | --- | --- |
| heparan sulfate proteoglycan biosynthetic process, polysaccharide chain biosynthetic process | BP | 0.01 | *sfl, sgl* |
| fibroblast growth factor receptor signaling pathway | BP | 0.01 | *sfl, pnt, sgl* |
| epithelial cell migration, open tracheal system | BP | 0.013 | *sfl, pnt, sgl* |
| mesoderm migration involved in gastrulation | BP | 0.014 | *sfl, sgl* |
| cell surface receptor signaling pathway | BP | 0.014 | *tnc, noc, E(spl)malpha-BFM, Ama, sfl, pnt, spz6, sgl, E(spl)m2-BFM* |
| cellular response to growth factor stimulus | BP | 0.014 | *tnc, sfl, pnt, sgl* |
| cGMP metabolic process | BP | 0.021 | *Pde1c, Pde9* |
| negative regulation of cell differentiation | BP | 0.032 | *salm, pnt, CadN* |
| stem cell development | BP | 0.032 | *Rbp6* |
| cGMP catabolic process | BP | 0.032 | *Pde9* |
| positive regulation of serine-type endopeptidase activity | BP | 0.032 | *pip* |
| ommatidial rotation | BP | 0.038 | *pnt, CadN* |
| cyclic nucleotide metabolic process | BP | 0.041 | *Pde1c, Pde9* |
| cell surface receptor protein tyrosine kinase signaling pathway | BP | 0.044 | *Ama, sfl, pnt, sgl* |
| maintenance of blood-brain barrier | BP | 0.044 | *Ama* |
| negative regulation of terminal cell fate specification, open tracheal system | BP | 0.044 | *salm* |
| myoblast fusion | BP | 0.044 | *Six4, lmd* |
| 3', 5'-cyclic-GMP phosphodiesterase activity | MF | 0.014 | *Pde1c, Pde9* |
| 3', 5'-cyclic-nucleotide phosphodiesterase activity | MF | 0.015 | *Pde1c, Pde9* |
| cyclic-nucleotide phosphodiesterase activity | MF | 0.017 | *Pde1c, Pde9* |
| heparan sulfate N-deacetylase activity | MF | 0.036 | *sfl* |
| UDP-glucose 6-dehydrogenase activity | MF | 0.036 | *sgl* |
| inositol-3-phosphate synthase activity | MF | 0.036 | *Inos* |
| [heparan sulfate]-glucosamine N-sulfotransferase activity | MF | 0.036 | *sfl* |
| sulfotransferase activity | MF | 0.036 | *sfl, pip* |
| natriuretic peptide receptor activity | MF | 0.036 | *CG14877* |
| proctolin receptor activity | MF | 0.036 | *Proc-R* |
| heterotrimeric G-protein binding | MF | 0.036 | *Octalpha2R* |
| calmodulin-activated 3', 5'-cyclic-GMP phosphodiesterase activity | MF | 0.036 | *Pde1c* |
| phosphoric diester hydrolase activity | MF | 0.044 | *Pde1c, Pde9* |
| D-erythro-sphingosine kinase activity | MF | 0.05 | *Sk1* |

### 1.9. Gene Ontology Enrichment Analysis: midgut

Significant GO terms (p < 0.05) with associated genes

| GO Term | Source | P-value | Genes |
| --- | --- | --- | --- |
| hemolymph coagulation | BP | 6.43e-04 | *PPO2, PPO1* |
| humoral immune response | BP | 0.001 | *PPO2, PPO1, Nplp2* |
| dopamine metabolic process | BP | 0.001 | *PPO2, PPO1* |
| defense response to Gram-positive bacterium | BP | 0.006 | *PPO2, PPO1* |
| defense response to fungus | BP | 0.007 | *PPO2, PPO1* |
| scab formation | BP | 0.011 | *PPO1* |
| wound healing | BP | 0.011 | *PPO2, PPO1* |
| pigment metabolic process | BP | 0.019 | *PPO2, PPO1* |
| cellular response to glucose stimulus | BP | 0.03 | *meep* |
| cellular response to monosaccharide stimulus | BP | 0.034 | *meep* |
| melanin biosynthetic process | BP | 0.036 | *PPO2* |
| positive regulation of insulin receptor signaling pathway | BP | 0.048 | *meep* |
| catechol oxidase activity | MF | 1.06e-04 | *PPO2, PPO1* |
| tyrosinase activity | MF | 1.06e-04 | *PPO2, PPO1* |
| long-chain fatty acyl-CoA binding | MF | 0.031 | *Acbp4* |
| aminoacylase activity | MF | 0.031 | *CG17109* |
| chitin binding | MF | 0.031 | *obst-H, CG7017* |
| monooxygenase activity | MF | 0.031 | *PPO2, PPO1* |
| fatty-acyl-CoA binding | MF | 0.047 | *Acbp4* |
| oxidoreductase activity, acting on paired donors, with incorporation or reduction of molecular oxygen | MF | 0.047 | *PPO2, PPO1* |

### 1.10. Gene Ontology Enrichment Analysis: fatbody

Significant GO terms (p < 0.05) with associated genes

| GO Term | Source | P-value | Genes |
| --- | --- | --- | --- |
| sex differentiation | BP | 0.002 | *Yp1, Yp2, Yp3* |
| lipid catabolic process | BP | 0.003 | *Yp1, Yp2, Yp3* |
| detection of peptidoglycan | BP | 0.017 | *PGRP-SD* |
| catabolic process | BP | 0.025 | *Yp1, Yp2, Yp3, PGRP-SD* |
| detection of biotic stimulus | BP | 0.029 | *PGRP-SD* |
| carboxylesterase activity | MF | 5.05e-05 | *Yp1, Yp2, Yp3* |
| lipase activity | MF | 2.12e-04 | *Yp1, Yp2, Yp3* |
| carboxylic ester hydrolase activity | MF | 6.45e-04 | *Yp1, Yp2, Yp3* |
| serine hydrolase activity | MF | 0.005 | *Yp1, Yp2, Yp3* |
| hydrolase activity, acting on ester bonds | MF | 0.013 | *Yp1, Yp2, Yp3* |
| N-acetylmuramoyl-L-alanine amidase activity | MF | 0.03 | *PGRP-SD* |
| peptidoglycan binding | MF | 0.032 | *PGRP-SD* |
| hydrolase activity | MF | 0.043 | *Yp1, Yp2, Yp3, PGRP-SD* |

### 1.11. Gene Ontology Enrichment Analysis: MM cell body glia

Significant GO terms (p < 0.05) with associated genes

| GO Term | Source | P-value | Genes |
| --- | --- | --- | --- |
| peroxiredoxin activity | MF | 0.023 | *Prx2540-1, Prx2540-2* |
| glutathione-dependent sulfide quinone oxidoreductase activity | MF | 0.037 | *CG14997* |
| L-ascorbate:sodium symporter activity | MF | 0.037 | *CG6293* |
| glucuronosyltransferase activity | MF | 0.037 | *GlcAT-S, Ugt35B1* |
| sulfide:quinone oxidoreductase activity | MF | 0.037 | *CG14997* |
| peptidoglycan transmembrane transporter activity | MF | 0.037 | *CG8046* |
| asioloorosomucoid beta-1, 3-glucuronosyltransferase activity | MF | 0.037 | *GlcAT-S* |
| ammonium channel activity | MF | 0.037 | *Rh50* |
| peroxidase activity | MF | 0.037 | *Prx2540-1, Prx2540-2* |
| galactosyl beta-1, 3 N-acetylgalactosamine beta-1, 3-glucuronosyltransferase activity | MF | 0.037 | *GlcAT-S* |
| N-acetyllactosamine beta-1, 3-glucuronosyltransferase activity | MF | 0.037 | *GlcAT-S* |
| oxidoreductase activity, acting on peroxide as acceptor | MF | 0.037 | *Prx2540-1, Prx2540-2* |
| hexosyltransferase activity | MF | 0.037 | *GlcAT-S, Ugt35B1, CG31690* |
| oxoglutarate dehydrogenase (succinyl-transferring) activity | MF | 0.049 | *CG1544* |
| galactosylgalactosylxylosylprotein 3-beta-glucuronosyltransferase activity | MF | 0.049 | *GlcAT-S* |
| transmembrane transporter activity | MF | 0.049 | *CG8046, CG6293, CG31100, Pzl, Rh50, CG1358* |
| glycosyltransferase activity | MF | 0.049 | *GlcAT-S, Ugt35B1, CG31690* |

### 1.12. Gene Ontology Enrichment Analysis: wing heart precursors

Significant GO terms (p < 0.05) with associated genes

| GO Term | Source | P-value | Genes |
| --- | --- | --- | --- |
| determination of genital disc primordium | BP | 3.49e-04 | *Abd-B, rho, pnt* |
| pericardial nephrocyte differentiation | BP | 4.26e-04 | *tin, pnr, pnt* |
| leg disc proximal/distal pattern formation | BP | 0.001 | *rho, Dll, pnt* |
| negative regulation of cardioblast cell fate specification | BP | 0.002 | *Abd-B, pnt* |
| specification of animal organ identity | BP | 0.002 | *Ubx, Dll* |
| cell fate determination | BP | 0.003 | *slou, Ubx, pnr, fz, rho* |
| neuroendocrine cell differentiation | BP | 0.004 | *tin, Abd-B* |
| midgut development | BP | 0.006 | *Abd-B, Ubx, fz* |
| cell fate commitment involved in pattern specification | BP | 0.006 | *Ubx, fz, rho* |
| gonadal mesoderm development | BP | 0.007 | *tin, Abd-B* |
| epithelial cell proliferation involved in Malpighian tubule morphogenesis | BP | 0.012 | *rho, pnt* |
| negative regulation of stem cell differentiation | BP | 0.012 | *Abd-B, pnt* |
| R3/R4 cell fate commitment | BP | 0.014 | *fz, pnt* |
| negative regulation of cell fate specification | BP | 0.018 | *Abd-B, pnt* |
| visual perception | BP | 0.024 | *Ggamma30A, rdgC, lovit* |
| cell redox homeostasis | BP | 0.03 | *Prx2540-1, CG12896* |
| negative regulation of striated muscle tissue development | BP | 0.033 | *Abd-B* |
| mitochondrial double-strand break repair via homologous recombination | BP | 0.033 | *mre11* |
| negative regulation of extracellular matrix disassembly | BP | 0.033 | *obst-A* |
| regulation of chitin-based cuticle tanning | BP | 0.033 | *rk* |
| embryonic heart tube development | BP | 0.033 | *tin, pnr* |
| positive regulation of extracellular matrix assembly | BP | 0.033 | *loh* |
| dorsal vessel aortic cell fate commitment | BP | 0.033 | *Ubx* |
| L-ascorbic acid transmembrane transport | BP | 0.033 | *CG6293* |
| external genitalia morphogenesis | BP | 0.033 | *Abd-B* |
| determination of ventral identity | BP | 0.033 | *Dll* |
| photoreceptor cell fate commitment | BP | 0.038 | *fz, rho, pnt* |
| muscle organ development | BP | 0.043 | *slou, Abd-B, Ubx* |
| ommatidial rotation | BP | 0.046 | *fz, pnt* |
| positive regulation of apoptotic process involved in morphogenesis | BP | 0.05 | *Abd-B* |
| male pigmentation | BP | 0.05 | *Abd-B* |
| germ cell migration | BP | 0.05 | *tin, Abd-B* |
| sucrose transport | BP | 0.05 | *lovit* |
| ammonium homeostasis | BP | 0.05 | *Rh50* |
| ammonium transmembrane transport | BP | 0.05 | *Rh50* |
| glutamine biosynthetic process | BP | 0.05 | *Gs2* |
| animal organ morphogenesis | BP | 0.05 | *tin, Abd-B, Ubx, pnr, fz, rho, Dll, pnt* |
